# A structural screen for candidate polyphosphate binding surfaces in human membrane associated proteins

**DOI:** 10.64898/2026.10.25.761654

**Authors:** Jianlin Wu, Zongchao Jia

## Abstract

Inorganic polyphosphate (polyP) can alter protein function through noncovalent binding, yet its partners among human membrane associated proteins remain poorly characterized. PolyP is recognized through two modes: a structure-based mode driven by charge complementarity with sites assembled by protein folding, and a more recently defined sequence-based mode at histidine and lysine rich tracts. The sequence determinant is well defined and directly searchable; the structure-based mode has few characterized examples and no systematic search method, because its residues leave no contiguous sequence signature. Broad experimental binding screens are especially demanding for integral membrane proteins, which require specialized conditions. We therefore developed a computational workflow that searches predicted structures for surfaces able to accommodate a phosphate chain. We designed custom scores that integrate surface electrostatics, docking results and structural confidence to prioritize candidates. From 19,481 completed docking searches on 5,648 AlphaFold models representing 2,925 UniProt protein entries, we nominated 82 entries as candidates for noncovalent polyP binding. Predicted polyP contacts in 18 candidate entries overlapped experimentally characterized functional residues, linking the predicted interfaces to established sites of ligand recognition, regulation and membrane attachment. The workflow ranks computational results rather than measuring binding, providing specific targets for experimental investigation.

## Introduction

Inorganic polyphosphate (polyP) is a linear chain of orthophosphate residues joined by phosphoanhydride bonds, found across bacteria, archaea and eukaryotes and long studied as a phosphate and energy reserve [1]. In human cells it is stored in platelet dense granules [2] and released upon platelet activation, where it accelerates clotting through interactions with coagulation proteins [3, 4]. PolyP also protects proteins from aggregation [1] and influences receptor signaling in mammalian cells [5, 6]. These activities are exerted through protein partners, and the recognition chemistry that selects those partners is only partly defined.

Two modes of noncovalent polyP recognition are now known, and they differ in what determines the interaction. The classical mode is structure-based: recognition arises from charge complementarity between the polyanion and a positively charged site assembled by the folded protein, whether that site is an exposed surface patch, a groove or an enclosed tunnel, and without any requirement for a particular sequence. The bacterial polyphosphate kinases provide the best structurally resolved examples. In *Escherichia coli* PPK1, which synthesizes polyP from ATP, the active site lies within a tunnel whose conserved, positively charged wall was proposed to engage the growing chain and to account for processive synthesis [7]. Structures of class I and class III PPK2 enzymes captured with bound polyphosphate show an extended chain accommodated against a basic surface adjacent to the nucleotide site [8]. These are catalytic sites rather than regulatory ones, but they establish the recognition chemistry of this mode: an extended phosphate chain is read by a complementary array of basic side chains contributed by parts of the fold that are distant in sequence.

The same recognition also operates at sites with no catalytic role, where polyP binding is regulatory rather than mechanistic. Thrombin provides the clearest characterized example in a human protein: polyP binds exosite II, a basic surface it shares partly with heparin [9]. A polyP site has also been proposed at an RNA binding region of the SARS-CoV-2 RNA dependent RNA polymerase [10], and polyP inhibits binding between the SARS-CoV-2 spike protein and ACE2 [11]. Beyond these, experimentally characterized examples of structure-based binding in human proteins remain few.

The second mode, described more recently, is sequence-based, and in contrast to the structure-based mode it now has a well defined structural feature for polyP binding. PolyP associates with histidine-rich tracts in human, yeast and bacterial proteins [12] and with lysine-rich tracts [13], and the polyacidic serine and lysine rich (PASK) clusters represent a related sequence determinant [14]. In each case a contiguous and compositionally distinctive stretch of sequence carries the signal, and these tracts are found largely in intrinsically disordered regions. A determinant of this kind can be searched directly, and a recent screen used lysine-rich sequences to identify human polyP interacting proteins [15].

The structure-based mode offers no equivalent handle, and this has a practical consequence for discovery. The basic residues that form a polyP binding site may be widely separated in sequence and brought together only by folding [16], leaving no contiguous signature for a sequence search to detect. The alternative is to search structures. An earlier in silico study took this approach, describing a positively charged semi-tunnel as a shared surface feature of the polyphosphate kinase families and testing it by docking short phosphate ligands, although it was applied to a few hundred structures selected for similarity to those enzymes [17]. Proteins that bind polyP through the structure-based mode therefore remain systematically underrepresented among known partners, and the small number of characterized examples may reflect the absence of a general search method rather than the rarity of the mode itself.

Membrane associated proteins are a population in which structure-based recognition is both plausible and poorly surveyed. Secreted polyP acts on cells from the outside, and proinflammatory responses to polyP in vascular endothelium require the receptor for advanced glycation end products and the P2Y1 purinergic receptor [6]. Negatively charged lipids control membrane attachment and signaling at the same surfaces: lipid binding supports myosin-1c attachment to membranes and the assembly of receptor–β-arrestin complexes [18, 19]. Surfaces that evolved to engage phosphorylated lipid headgroups present the kind of basic, phosphate accommodating geometry that could also engage a phosphate polymer. For human membrane associated proteins this defines a specific structural question: which proteins could bind polyP, where binding could occur, and which established functional sites might be involved. The relevant surfaces include both extracellular regions and regions facing the cytoplasm.

Examining this question experimentally across many proteins is demanding. Extraction can disrupt the stability or activity of integral membrane proteins, and conditions that preserve each protein often require individual optimization [20]. Preparing each protein separately makes broad experimental binding screens costly in time and resources.

Predicted structures remove the main obstacle to a structural search at scale. AlphaFold provides models for large protein sets [21, 22], and prospective screens of two human receptors using AlphaFold2 models yielded experimentally confirmed ligands with hit rates and affinities comparable to screens using experimental structures [23]. The structure-based question can therefore now be asked at the scale previously available only to sequence-based searches.

Here we screen human membrane associated proteins for surfaces able to engage polyP through the structure-based mode. Poisson–Boltzmann electrostatics calculated under near physiological temperature and ionic strength [24] located continuous positive surface patches [25], and each patch defined a search box in which AutoDock Vina docked polyP10, a ten phosphate model chain, scored with the AutoDock4 force field [26, 27, 28]. We developed two scoring formulae: the Patch Docking Index (PDI100) ranks docking results by electrostatic intensity and docking favorability, and the pLDDT Reliability Index (PRI) applies a penalty derived from AlphaFold model confidence across the search box. We then varied the index settings systematically to establish which results did not depend on a particular weighting.

Applying this screen to 8,808 AlphaFold models representing 4,033 reviewed human UniProt protein entries annotated to the cell or plasma membrane, we nominate 82 protein entries whose surfaces meet stringent combined criteria, report the functional classes overrepresented among them, and compare the predicted contacts with experimentally characterized recognition and regulatory sites. The screen ranks computational results rather than measuring binding: it was built to limit false positive nominations, not to recover every polyP binding protein, and proteins outside the selected set are not thereby excluded.

## Methodology

### Overview

We used a stringent computational screen to nominate human membrane protein candidates for noncovalent polyphosphate (polyP) binding. PolyP carries repeated negative charges. Structural studies of PPK1 and PPK2 support recognition of a phosphate chain by positively charged sites assembled by protein folding, motivating the use of positive surface regions as starting points for docking [7, 8]. The screen began with 8,808 AlphaFold models, including models of alternative isoforms, linked to reviewed UniProt protein entries annotated to the cell or plasma membrane. The workflow combined electrostatic mapping, polyP docking and local structural confidence to rank docking results and used protein location to describe the selected candidates (Fig. 1a,b,d).

**Figure 1.**
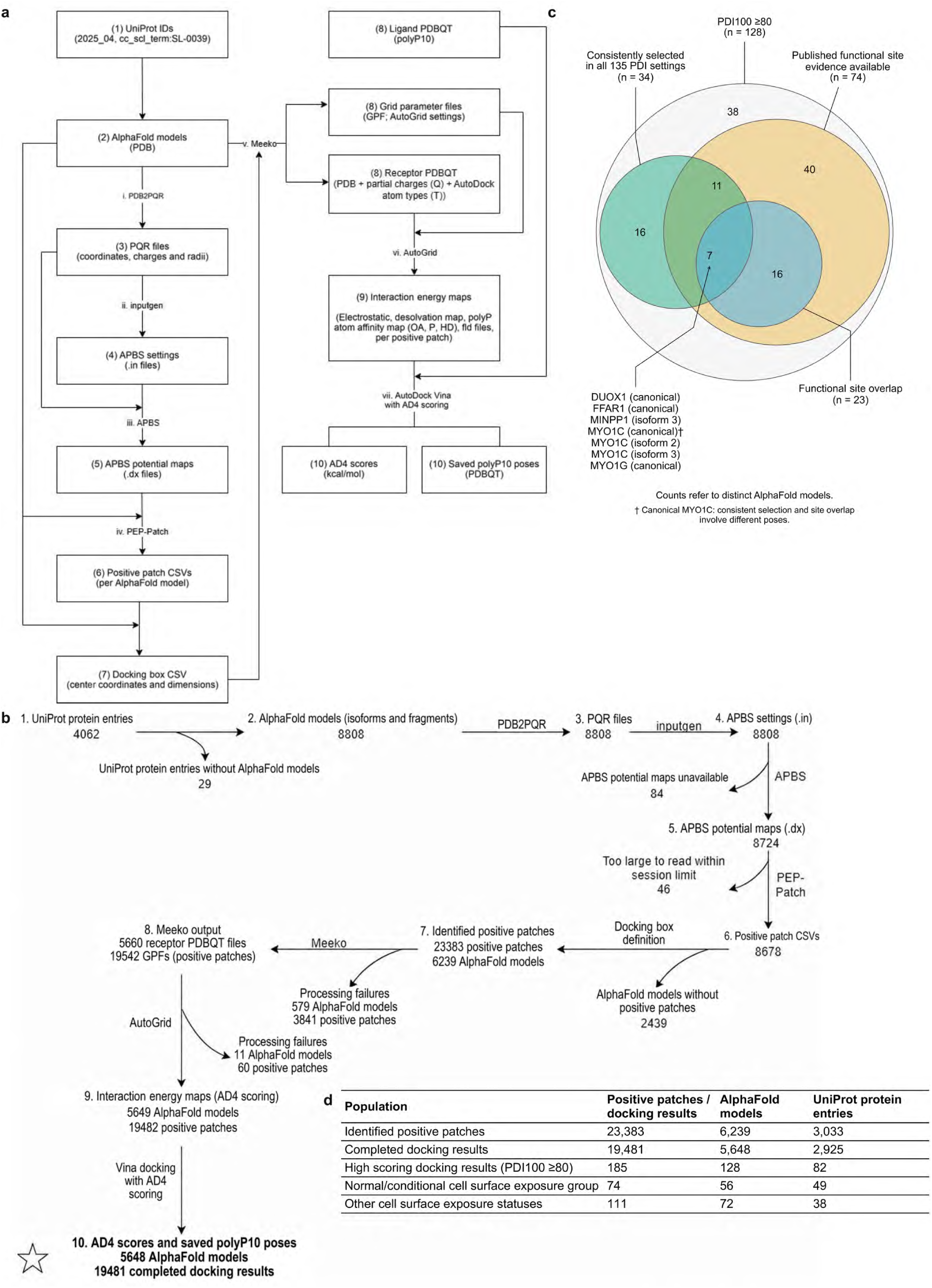
Electrostatic screening nominated candidate groups with complementary evidence. (a) Software and file dependencies. (b) Numbers processed and lost at each step through polyP10 docking. (c) Evidence groups among the 128 AlphaFold models with high scoring docking results. Consistent selection means that at least one docking result remained in the top 185 under all 135 PDI settings. Canonical MYO1C combined consistent selection and functional site overlap through different poses (†). (d) Screening and cell surface exposure counts; exposure did not determine selection. Models are distinct AlphaFold models; protein entries are UniProt protein entries. Five protein entries occur in both exposure groups.

PDB2PQR prepared the AlphaFold models for APBS electrostatic calculations [24], and PEP-Patch identified continuous positive patches that defined the docking box centers [25]. AutoDock Vina searched each box for poses of polyP10, a computationally manageable chain of ten phosphate residues [26, 27]. We retained the pose with the best AutoDock4 (AD4) score from each search.

The Patch Docking Index (PDI100) combined scaled measures of patch electrostatic intensity and docking favorability, with a penalty derived from AlphaFold confidence across the docking box. PDI100 orders computational results for candidate nomination; its ranking does not establish an order of biological binding affinity or binding probability. We selected docking results scoring PDI100 ≥80 for detailed analysis and used protein localization, topology and isoform information to describe the associated AlphaFold models. Nonselection is not evidence of weak binding or nonbinding.

We compared scores across isoforms to assess consistency and tested 135 combinations of PDI settings to determine how consistently each candidate docking result remained selected. Functional enrichment analysis identified functional classes overrepresented among UniProt protein entries with high scoring docking results. We compared predicted polyP contacts with experimentally identified recognition and regulatory sites to assess whether the contacts involved regions important for protein function. The pipeline was developed locally and run at scale on Queen’s CAC Frontenac. We also identified docking poses that remained selected under all 135 PDI settings and contacted mapped residues at experimentally characterized functional sites (Supplementary Methods, “Combining evidence within each AlphaFold model”).

### AlphaFold model retrieval

We first searched UniProt [29] for reviewed Homo sapiens proteins annotated with the UniProt vocabulary term SL-0039 (cell/plasma membrane). The search identified 4,062 UniProt protein entries, generally one entry per gene. We downloaded their main accession numbers (UniProt IDs), without isoform specific suffixes, as a list for subsequent 3D structure retrieval (Supplementary Data S7). The SL-0039 “cell/plasma membrane” annotation also covers proteins on intracellular membranes and proteins that face only the cytoplasmic side of the surface membrane. This inclusive membrane protein starting set was therefore not limited to extracellular facing proteins.

To standardize inputs across the multistep screening pipeline, we mapped the main UniProt IDs to entries in the AlphaFold Protein Structure Database and retrieved models exclusively from that database. Experimental PDB structures may contain missing regions, cover different protein segments, or represent different forms obtained under different experimental conditions. These differences can complicate structure preparation and introduce variability into subsequent analyses.

A UniProt protein entry can be linked to multiple AlphaFold models representing different isoforms of the same gene or protein fragments labeled with an “-F” suffix (F1, F2… Fn).

We therefore retrieved 8,808 AlphaFold models in PDB format using the 4062 main UniProt IDs. These models represented 4,033 UniProt protein entries; 29 UniProt protein entries had no corresponding AlphaFold model at the time of retrieval (Supplementary Data S7).

### AlphaFold model preparation with PDB2PQR

Structural studies provide a biological basis for examining electrostatic attraction between polyP and basic protein regions. PPK1 contains a positively charged tunnel proposed to engage polyP, while a ligand bound class I PPK2 structure shows polyP positioned by basic residues in a folded channel [7, 8]. These structural features motivated our calculation of electrostatic potentials under specified solution conditions and our use of positive surface patches to guide docking. Those calculations required charges and radii beyond the atomic positions in the retrieved AlphaFold PDB files.

Each AlphaFold model was processed with PDB2PQR 3.7.1 [24]. PROPKA estimated protonation states at pH 7.4 under near physiological conditions [30, 31]; PDB2PQR then added the corresponding hydrogen atoms and assigned partial charges and radii using PARSE. Implementation details are given in Supplementary Methods, “Preparing electrostatic maps”. The -- drop-water option removed any water molecules from the input structures, while APBS represented the surrounding water as a continuous medium. PDB2PQR produced 8,808 PQR files, one per AlphaFold model, for APBS calculations.

### Electrostatic calculations with APBS

APBS (Adaptive Poisson–Boltzmann Solver) [24] used the PQR coordinates, charges and radii to calculate electrostatic potential around each protein in solution. Solving the Poisson-Boltzmann equation accounted for water and dissolved ions, allowing strongly positive patches to be identified under the specified solution conditions.

APBS calculations used 310.15 K (37 °C) and 0.14 M monovalent ionic strength to approximate physiological conditions. The protein and solvent dielectric constants were 4.0 and 74.2, respectively. A 1.4 Å spherical probe defined the boundary between the protein and solvent.

APBS 1.3 calculated a .dx electrostatic potential map from each PQR file and its matching settings (.in) file. Of 8,808 AlphaFold models, 8,724 produced maps and 84 had no available map.

### Positive patch identification with PEP-Patch

The APBS .dx maps describe electrostatic potential throughout a 3D grid, but we needed to locate continuous positive regions on the protein surface as candidate polyP binding sites. We used PEP-Patch in electrostatics mode to identify and measure surface patches (Fig. 2b). Here, a positive patch is a continuous surface region where the potential exceeds a chosen cutoff.

**Figure 2.**
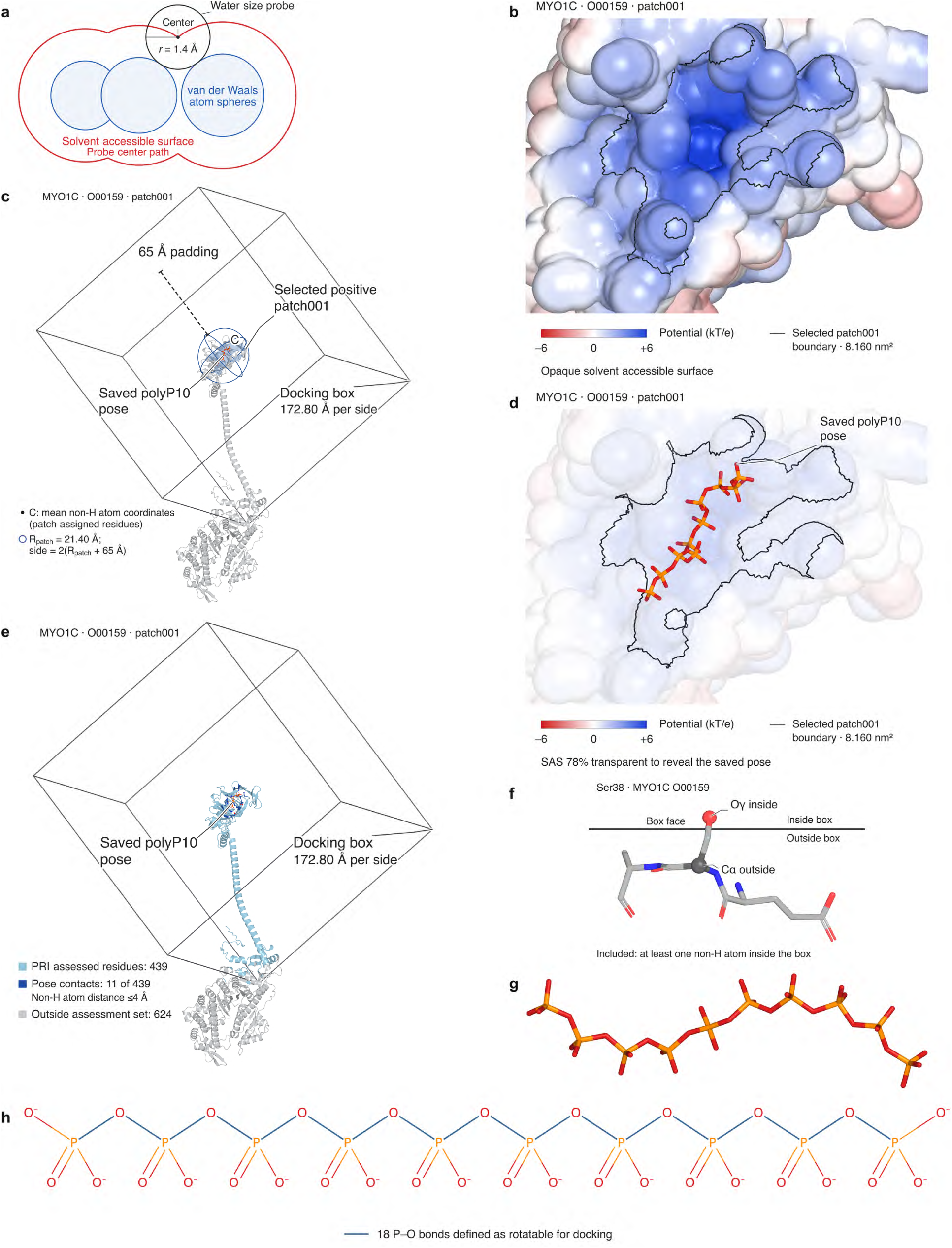
Positive surface patches defined polyP10 docking boxes. (a) The solvent accessible surface follows the center of a 1.4 Å probe around protein atoms. (b–e) Canonical MYO1C (O00159), patch001. (b) APBS potential mapped onto the surface; the selected positive patch is outlined. (c) C marks the mean coordinates of the nonhydrogen atoms in patch assigned residues. The patch extent, Rpatch = 21.40 Å, and 65 Å padding defined a 172.80 Å cubic box, with side = 2(Rpatch + 65 Å). (d) The same surface view as b, made transparent to reveal the saved polyP10 pose. Blue and red surfaces indicate positive and negative potential. (e) PRI was calculated from 439 residues with at least one nonhydrogen atom in the docking box; 11 contacted the saved pose within 4 Å between nonhydrogen atoms. The remaining 624 model residues lay outside the box. (f) In the separate patch012 box, Ser38 was included because Oγ lay inside although Cα lay outside. (g) Prepared polyP10 input conformation. Phosphorus is orange and oxygen red. (h) PolyP10 connectivity and formal charges; blue bonds mark the 18 P–O bonds defined as rotatable for docking.

PEP-Patch used each AlphaFold PDB file and its matching .dx map to group connected surface vertices above the potential cutoff into patches. PEP-Patch defined the solvent accessible surface using a water sized probe with a radius of 0.14 nm (1.4 Å; Fig. 2a). Mesh construction and output fields are described in Supplementary Methods, “Linking positive patches to docking boxes”.

The positive potential cutoff was set to +2.7 kT/e (approximately +72 mV at 310.15 K). In a point charge approximation, this gives an electrostatic energy of −2.7 kT for a single negative charge, such as that of a DNA backbone or internal polyP phosphate.

Only positive patches exceeding 0.7 nmZ, roughly the membrane footprint of one phospholipid molecule [32], were retained. These criteria prioritize strongly positive patches while excluding very small patches, without requiring the entire polyP chain to fit within a patch. These stringent cutoffs were chosen for candidate nomination, not as established binding requirements or evidence of nonbinding below either cutoff.

PEP-Patch could not read 46 AlphaFold models within CAC’s 6 h session limit because they were too large; it therefore analyzed 8678 AlphaFold models in total. PEP-Patch identified 23383 positive patches above threshold in 6239 of the 8678 AlphaFold models analyzed; the remaining 2439 models had no identified patch above the cutoff.

### Patch Coordinates and Vina Docking Box Definition

Each positive patch was assigned a 3D docking box within its AlphaFold model for AutoDock Vina to explore ligand positions and conformations (Fig. 2c). The boxes were defined by their center coordinates and dimensions in ångströms (Å).

A Python script defined each docking box by first matching patch IDs labeled positive in *_patch_list.csv to their residues in *_residue_list.csv. It then located each residue by its name and number in the corresponding AlphaFold PDB file and extracted the coordinates of its nonhydrogen atoms. The script calculated the box center coordinates as the mean x, y and z coordinates of these atoms.

The greatest distance from this center to any included atom defines the patch extent (R_patch_A). Padding was set to 65 Å beyond this extent in every direction. Each box is therefore cubic, with a distance from the center to a face of R_patch_A+65 Å and a full side length of 2×(R_patch_A+65 Å).

This 65 Å padding was chosen to accommodate fully extended polyP conformations and allow sampling of neighboring surface regions beyond the +2.7 kT/e positive patch boundary. Positive patches near the 0.7 nmZ area cutoff can be too small to accommodate the full polyP10 chain. They therefore guide box placement without confining the ligand to the above threshold region.

The script produced one CSV containing 23,383 patch centered docking boxes across 6,239 AlphaFold models. Each row linked a positive patch and its AlphaFold model to the box center coordinates (center_x, center_y, center_z) and dimensions (size_x, size_y, size_z), all in Å. Additional output fields are described in Supplementary Methods, “Linking positive patches to docking boxes”. Vina later used the reported center_* and size_* values directly as its docking box parameters.

### pLDDT Reliability Index (PRI)

We developed a custom scoring formula, the pLDDT Reliability Index (PRI), to downweight docking results from model regions with low structural confidence.

AlphaFold’s predicted local distance difference test (pLDDT) scores each residue from 0 to 100, with higher values indicating greater confidence in its local structure. In AlphaFold’s confidence guide, pLDDT ≥ 90 indicates very high confidence, often supporting reliable detailed structural interpretation; 70-90 indicates a confident prediction with a generally reliable backbone; 50-70 indicates low confidence and calls for caution; and < 50 indicates very low confidence, often associated with flexibility or disorder [22]. PRI uses the pLDDT scores stored in the B-factor column of the original AlphaFold PDB files.

We calculated PRI for the entire docking box, including regions outside the final polyP contacts (Fig. 2e). Vina samples poses throughout this box, so PRI summarizes confidence in all model regions available to the search rather than only in the residues contacted by the saved pose.

For each box, a residue’s pLDDT score was included in the PRI calculation if any of its nonhydrogen atoms lay within the boundaries defined by the docking box CSV’s center coordinates and dimensions. This inclusion rule captures side chains extending into the docking box even when their Cα atoms lie outside (Fig. 2f); each residue contributes only once.

To combine residue level confidence into one score for the docking box, each included residue’s pLDDT value xi was converted to a 0–1 weight Si, using pLDDT values of 50 and 90 as reference points:

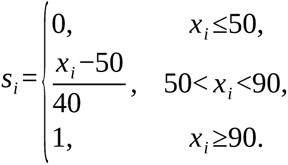

Residues scoring ≤50 receive 0 weight but remain in the calculation, lowering the box average. Weights increase linearly for intermediate pLDDT scores, allowing residues with moderate structural confidence to contribute without receiving full weight. Scores ≥90 receive full weight without additional rewards within this already high confidence range.

We calculated PRI by subtracting a variability term from the mean of the residue weights si. The mean *s̅* represents the overall confidence across N residues in the docking box, and the standard deviation, SD(s), measures how unevenly confidence is distributed across those residues.

Subtracting 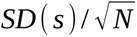 gives a larger deduction when residue confidence scores are more uneven. For the same variability in residue weights, a box containing more residues receives a smaller deduction because averages based on fewer residues are more sensitive to individual scores. Negative PRI values are set to zero, keeping PRI between 0 and 1.

The deduction lowers PRI for boxes that mix residues with high and low confidence, which a mean alone can obscure.

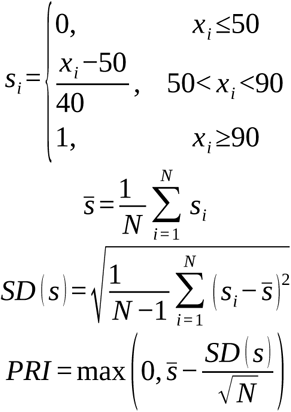

We calculated PRI for all 23,383 docking boxes and saved the values in a separate CSV under their AlphaFold model patch identifiers. These identifiers link PRI to subsequent docking results (Supplementary Data S7).

### Docking preparation with Meeko and AutoGrid

AutoDock Vina searched for ligand arrangements within a docking box on a receptor, and the AD4 scoring function estimated how energetically favorable each pose was. AD4 uses the AutoDock4/4.2 semiempirical free energy force field, which was calibrated against protein–ligand complexes with experimentally determined structures and binding energies [28]. Its explicit electrostatic term represents attraction between anionic polyP and cationic protein regions. This complements the positive patch measure by evaluating electrostatics within a ligand pose.

Meeko [33] converted the PQR structures to PDBQT files containing atomic coordinates, partial charges and the AutoDock atom types required for docking. AutoGrid4 generated the precalculated 3D interaction energy maps required for AD4 scoring. A custom Python script read the ligand atom types from the polyP10 PDBQT to specify the required maps.

A custom Python script used each Meeko generated Grid Parameter File (GPF) as a template to create a separate GPF for every positive patch on the corresponding AlphaFold model. The script inserted the patch’s docking box center coordinates and dimensions from the docking box CSV. AutoGrid used each patch specific GPF and its matching receptor PDBQT to generate the gridmap set required for AD4 scoring. File preparation and map construction are detailed in Supplementary Methods, “Searching polyP10 poses with AD4 maps”.

After Meeko and AutoGrid4 processing, 19482 of the 23383 positive patches identified by PEP-Patch above the 2.7 kT/e threshold were ready for docking, covering 5649 of the 6239 AlphaFold models. Processing failed for 3901 patches, leaving 590 AlphaFold models with no patches ready for docking.

### Vina docking simulation with AD4 scoring

AutoDock Vina searched each patch centered docking box for favorable polyP10 poses, and AD4 scored their predicted binding free energies. The AD4 scoring algorithm combines weighted contributions from van der Waals contacts, hydrogen bonding based on donor–acceptor geometry, electrostatic interactions, and desolvation, the energy changes caused by reduced solvent exposure during binding. These terms allow the docked polyP10 pose to be evaluated through several noncovalent interaction contributions, extending the electrostatic description supplied by the positive patch.

AD4 also includes a torsional penalty that increases with the number of rotatable bonds. For a flexible linear ligand such as polyP, a relatively favorable AD4 score therefore requires electrostatic, hydrogen bonding and van der Waals interactions strong enough to compensate for the loss of conformational freedom upon binding. The torsional rationale is expanded in Supplementary Methods, “Searching polyP10 poses with AD4 maps”.

Biological polyP can contain tens to thousands of phosphate units. Longer chains have more rotatable bonds and a much larger conformational search space, increasing the chance that Vina misses favorable poses within a limited search. Greater flexibility also increases the AD4 torsional entropy penalty, making very long polyP chains harder to sample reliably and score robustly.

We therefore selected polyP10 (Fig. 2g,h; Supplementary Data S7), containing ten phosphate units, as a shorter model to identify plausible local binding arrangements. The model retains a flexible phosphate chain with repeated negative charges that can interact with cationic protein surfaces. Its shorter length limits the range of possible conformations, enabling consistent, computationally manageable surfaceome scale screening.

A custom Python script ran Vina 1.2.5 separately for each patch, using its AutoGrid4 grid map set and the same polyP10 ligand PDBQT. Search exhaustiveness was set to 16, giving 16 independent searches from random starting arrangements. Each run saved only the best scoring pose found.

For each successful run, Vina wrote the retained polyP10 pose (Fig. 2d) as atomic coordinates in a PDBQT file and reported its AD4 score in kcal/mol. More negative scores indicate more favorable predicted binding under the AD4 scoring model. The script recorded the best score per patch in a CSV so the scores could be combined with the patch descriptors and PRI for ranking.

Completed docking results comprised one retained pose and AD4 score for each of 19,481 patches across 5,648 AlphaFold models (Supplementary_Structures.zip; Supplementary Data S7).

### Patch Docking Index (PDI100): integrated docking result ranking

We developed a custom scoring formula, the Patch Docking Index (PDI100), to rank patch level docking results for candidate nomination. Structural evidence for polyP recognition at basic protein sites [7, 8] informed its two interaction components: E measures patch electrostatic intensity, and D measures the favorability of a docked polyP10 pose. PRI serves a separate role as a structural confidence criterion, reducing the index for docking boxes with low AlphaFold confidence. The combined index selects results that meet stringent computational criteria; it does not classify every protein as a binder or nonbinder.

Before combination, the docking and electrostatic inputs were converted to a common 0–1 scale. The transformation preserves the ordering within each measure while compressing extreme values.

The first raw measure, E, represents the positive electrostatic signal per unit patch area. The PEP-Patch field patch_value_total is the sum of APBS potentials, in kT/e, assigned to the patch’s surface vertices. Larger patches can accumulate a greater total signal simply because they contain more vertices. To reduce this size dependence, the total was divided by the patch area:

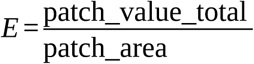

Here, patch_area is the patch’s surface area in nmZ. E was used to emphasize strongly positive regions that could attract anionic polyP, rather than reward patch size alone. E is a relative electrostatic intensity measure derived from APBS electrostatic potential, not a measurement of physical charge density or absolute surface charge.

The second measure, D, represents docking favorability. AD4 best_score is the saved polyP10 pose’s predicted binding energy score in kcal/mol, with more negative values indicating more favorable docking. Its sign was therefore reversed:

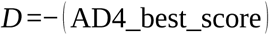

Reversing the sign converts a more negative docking energy to a larger *D*.

D and E have different units and represent different properties. We scaled each measure relative to its distribution, then applied a logistic transformation to place both on a common 0–1 scale. Larger raw values retain larger transformed values within each measure, while differences at the extremes are compressed.

We used a continuous transformation instead of bins that would, for example, assign every value above the 95th percentile a score of 1. We avoided min-max scaling because it can be dominated by extreme minimum or maximum values, compressing differences among most values in the dataset. Rank/quantile transforms were also avoided because they mainly preserve ordering rather than magnitude; values with neighboring ranks can receive similarly spaced scores even when their original numerical differences vary widely.

We therefore scaled D and E separately relative to each measure’s median and typical spread. For each measure x ∈{D,E}, we applied robust scaling analogous to a z score:

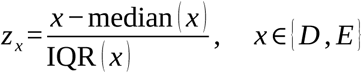

The notation x ∈{D,E} means that the same scaling is applied separately to D and E. Subtracting the median centers each measure around its typical value, so the median becomes z=0. The interquartile range (IQR) spans the middle 50% of measure values. Dividing the difference from the median by the IQR expresses how far the measure value lies above or below its median relative to this usual spread. A positive z therefore denotes a measure value above its median, and a negative z denotes one below it. Using the IQR rather than the full range makes the scale less sensitive to extreme values; differences between docking results are expressed in units of the IQR.

A logistic sigmoid then converts each z value into a continuous 0-1 score:

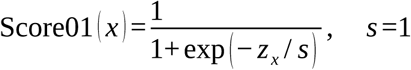

Here, exp(y) means the exponential function e^y^. The logistic sigmoid converts any positive or negative z value smoothly into the 0-1 range while keeping larger z values as larger scores. When the measure is at its median, z=0 and the score is 0.5. Values above the median give scores above 0.5, while values below it give scores below 0.5. The same change in measure value produces a larger score change near the median than at the extremes. Extreme values are gradually compressed toward 0 or 1, limiting their influence on the final PDI100.

The adjustable parameter s controls how sharply the score changes around the median. We manually set s=1, applying no additional rescaling beyond the preceding median/IQR standardization. The resulting scores are stored in the final CSV as:

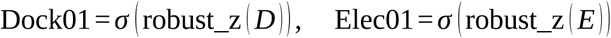

We combined Dock01 and Elec01 into a single score, Base, using a weighted geometric mean. This limits the compensation between a high Elec01 score and a low Dock01 score, or vice versa, that a simple weighted sum would permit:

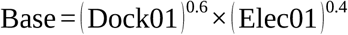

We manually set the ranking weights to 0.6 for docking and 0.4 for electrostatics. They sum to one and give docking modestly greater influence because it evaluates the complete ligand pose and several interaction terms, whereas E summarizes the patch’s positive potential. These weights were not fitted to experimental polyP binding measurements. We later assessed dependence of candidate selection on these weights and the adjustable sigmoid and PRI penalty settings across 135 parameter combinations (Results, “A subset of high scoring docking results remained selected across all tested PDI settings”).

Finally, PRI is applied to Base only as a penalty, reducing Base for docking boxes below the chosen structural confidence reference and leaving it unchanged at or above that reference. We manually set the PRI reference to 0.75. The pLDDT 70–90 confidence range provides context for favoring reliable backbone geometry, but PRI is not simply pLDDT divided by 100. For example, a region with pLDDT = 80 for every included residue would have a PRI = 0.75.

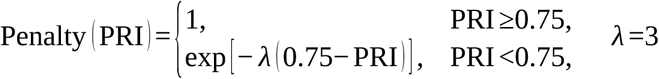

The manually adjustable parameter lambda controls how much the PRI penalty reduces Base when PRI is below the reference. We manually set lambda to 3, retaining approximately 86% of Base at PRI = 0.70 and 47% at PRI = 0.50. Thus, PRI values just below the reference cause only small reductions in Base, while lower PRI values cause progressively larger reductions, without an abrupt pass/fail boundary. At PRI values of 0.75 or above, the penalty multiplier remains one, leaving Base unchanged.

The final index is expressed on a 0-100 scale:

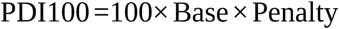

The factor of 100 improves readability without changing ranking or adding resolution; distinctions remain in the continuous scores and their decimal values. Higher PDI100 scores prioritize docking results that combine a strong positive electrostatic signal, favorable polyP10 docking and sufficient structural confidence under the chosen settings. This computational order supports candidate nomination without estimating biological binding affinity or probability.

### Compiling the Results

A custom Python script assembled all patch level results into one master spreadsheet for selection, sorting and interpretation (Fig. 3e; Supplementary Data S1). The source files held separate patch records (Fig. 3a,b), docking box records (Fig. 3c) and PRI records (Fig. 3d; Supplementary Data S7). Each row represents one positive patch above the PEP-Patch threshold on one AlphaFold model. Results were matched using *patch_id,* which combines the AlphaFold model identifier and patch number suffix. The table retained patches lacking downstream outputs because of computational failures, with unavailable parameter fields left blank to record processing losses. The table contains 23,383 rows, including 19,482 with numerical PDI100 values. One of these rows lacked an AD4 score and was excluded from completed result analyses (Supplementary Methods, “Defining completed and high scoring results”).

**Figure 3.**
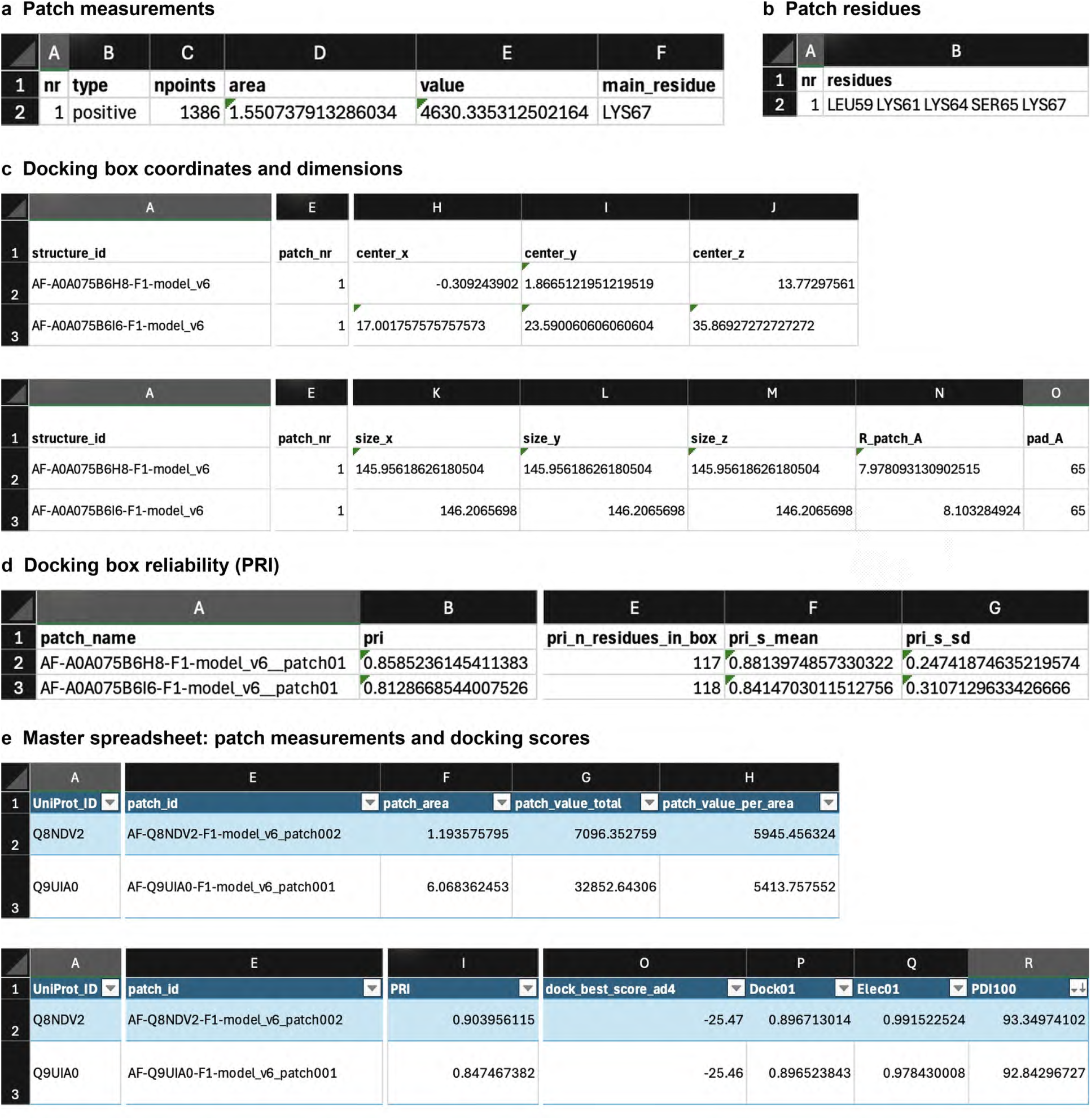
Patch level outputs were combined in the master spreadsheet. Excel excerpts retain the original column names and data. (a) Patch measurements from *_patch_list.csv. (b) Patch residue assignments from *_residue_list.csv. Both show positive patch 1 of AF-A0A075B6H8-F1-model_v6. (c) Docking box coordinates and dimensions. (d) PRI, residue count, and the mean and standard deviation of the residue weights. Panels c and d show patch 1 of the A0A075B6H8 and A0A075B6I6 models. (e) Separate master spreadsheet examples for GPR26 (Q8NDV2), patch002, and CYTH4 (Q9UIA0), patch001, showing the patch measurements and docking scores joined by patch_id. Selected column groups are displayed in separate strips, with identifiers repeated to link the same rows. Patch areas are in nmZ; docking box coordinates and dimensions are in Å.

The *patch_residues* column lists the residues that PEP-Patch assigned to each patch. From this list, *pct_HIS*, *pct_ARG* and *pct_LYS* were calculated as the percentages of residues that are histidine, arginine and lysine, respectively; *pct_HRK* is their combined percentage. These percentages describe the basic residue composition of each positive patch.

### High scoring docking results and protein location

We used a stringent PDI100 cutoff to nominate candidates from the 19,481 completed docking results and focus detailed analysis on the selected set.

We manually chose PDI100 ≥ 80 as a stringent cutoff defining a manageable set of high scoring docking results for detailed analysis. The cutoff selected the top 185 docking results in the PDI100 ranking across 128 AlphaFold models. Grouping AlphaFold models of alternative isoforms by their UniProt protein entry gave 82 unique UniProt protein entries. This selected set corresponds to the top ∼0.95% of the 19,481 completed docking results.

No PDI100 value equaled exactly 80, so PDI100 ≥ 80 and PDI100 >80 selected the same records (Supplementary Methods, “Defining completed and high scoring results”). These 185 high scoring docking results were recorded in the second worksheet of the master spreadsheet (Supplementary Data S1).

The settings, cutoffs and PDI100 were chosen to minimize false positive nominations by requiring strong combined electrostatic, docking and structural confidence evidence. The screen was never intended to recover every polyP binding protein or minimize false negatives. Results outside the shortlist were not classified as weak binders or nonbinders.

For each PDI100 ≥80 docking result, we reviewed the documented location and topology of the represented protein using the exact associated AlphaFold model Entry_ID. The inclusive UniProt SL-0039 cell/plasma membrane search captured proteins on intracellular membranes, such as mitochondrial or Golgi membranes, as well as plasma membrane proteins with or without extracellularly exposed regions. We manually reviewed UniProtKB and NCBI/RefSeq records for protein localization, membrane topology and isoform specific information. When these records did not clearly establish extracellular exposure, we consulted primary experimental or structural studies for additional evidence. We assessed AlphaFold models separately by Entry_ID because alternative isoforms can differ in expression, localization and topology (Supplementary Data S7).

For cell surface exposure, YES indicates that the represented protein has a documented extracellular facing region at the cell surface under its normal localization. CONDITIONAL indicates that this exposure is documented only under specific biological circumstances. For example, MCOLN1/TRPML1 is primarily lysosomal but can transiently reach the plasma membrane during lysosomal exocytosis [34]. Other CONDITIONAL cases depended on factors such as cell type, trafficking state, membrane assembly, or a documented cell surface population.

The normal/conditional cell surface exposure group comprised 74 of the 185 high scoring docking results: 69 associated with YES models and five with CONDITIONAL models. These 74 results spanned 56 AlphaFold models and represented 49 unique UniProt protein entries. Protein location provided descriptive context for all 185 results; it did not determine inclusion in the ranking sensitivity and functional site analyses. The cell surface exposure labels describe the represented protein or isoform, not access to the final polyP10 contact site (Supplementary Data S2).

### Code availability

The screening and analysis scripts, supporting tabular inputs and run instructions are supplied in Supplementary_Code.zip.

### Use of artificial intelligence

Codex (OpenAI) and Claude (Anthropic) assisted with Python script generation and organization of supplementary files.

## Results

### Stringent PDI selection nominated membrane protein candidates

Of the 8,678 AlphaFold models analyzed by PEP-Patch, 6,239 (71.9%) contained 23,383 positive patches above the study’s +2.7 kT/e and 0.7 nmZ thresholds. These models contained a median of three positive patches, and 77.5% had more than one, providing multiple starting regions for docking (Supplementary Data S1).

PolyP10 docking was completed for 19,481 of the 23,383 positive patches across 5,648 AlphaFold models and 2,925 UniProt protein entries (Fig. 1d; Supplementary Data S1). Each docking result records one patch centered search on one AlphaFold model. Models of different isoforms can belong to the same UniProt protein entry, and searches from different patches can reach the same surface region. We therefore counted docking results, AlphaFold models and UniProt protein entries separately.

The PDI100 ≥80 cutoff selected 185 docking results (the top 0.95% of completed results) across 128 AlphaFold models and 82 UniProt protein entries (Fig. 1d; Supplementary Table S7). This computational ranking nominates candidates; it does not establish an order of protein binding affinity or biological binding probability. Nonselection is not evidence of weak binding or nonbinding.

The PRI penalty reduced docking results meeting the ≥80 cutoff from 312 with Base ×100 to 185 with PDI100, moving 127 results (40.7%) below the cutoff (Fig. 4a). We also compared the top 185 docking results ranked by AD4 score, electrostatic intensity E, Base or PDI100. Comparing Base before the PRI penalty with PDI100 isolated PRI’s contribution at a fixed selection size: results with PRI <0.75 decreased from 86 to 18 (Fig. 4b; Supplementary Methods, “Isolating PRI’s effect on selection”). Adding PRI therefore reduced selection from lower confidence regions of the AlphaFold models.

**Figure 4.**
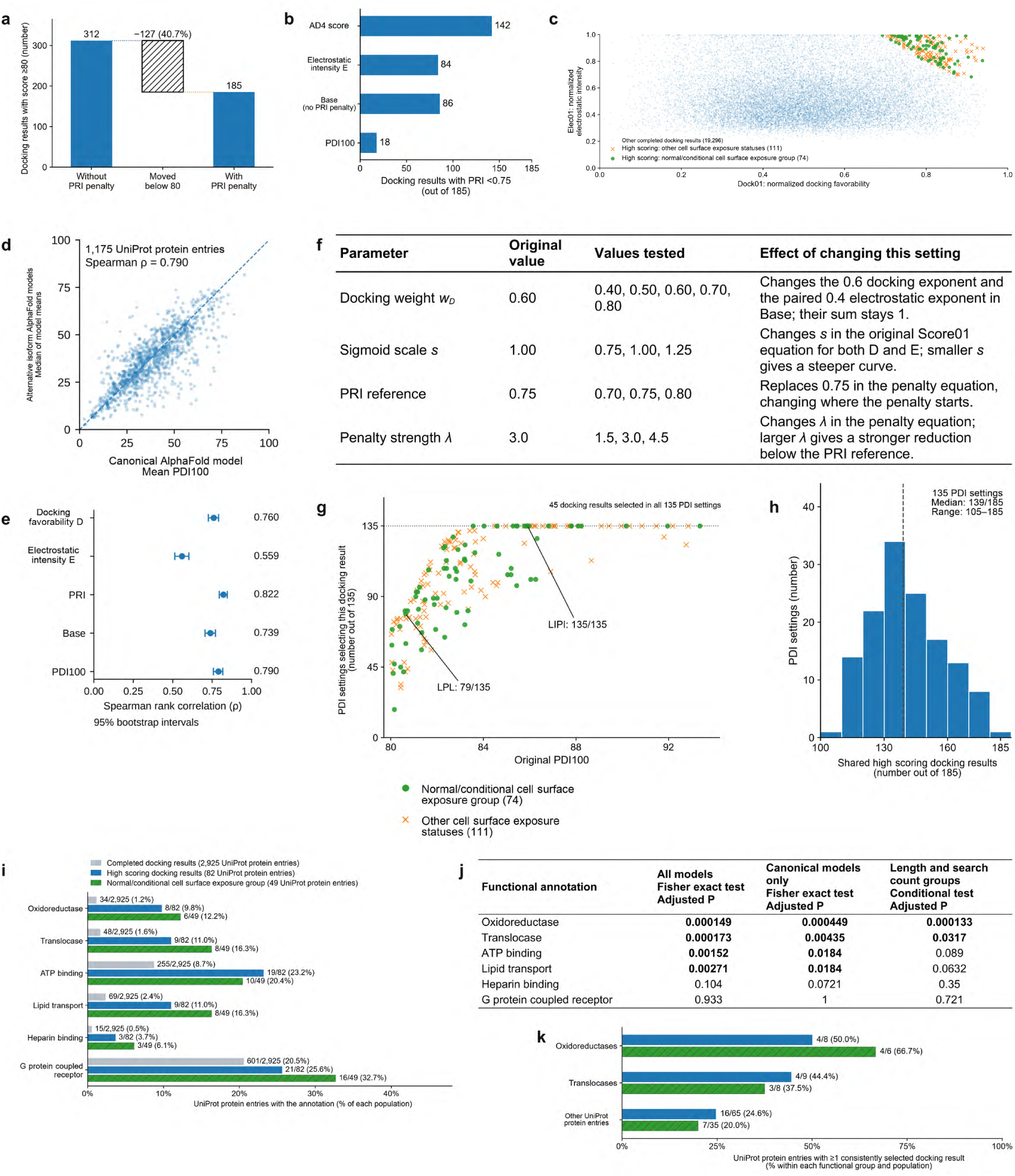
PDI selection, score consistency and functional representation. (a) PRI penalty effect on the ≥80 cutoff; hatching marks results moved below it. (b) Counts with PRI <0.75 among each metric’s top 185 docking results. (c) Normalized docking favorability versus electrostatic intensity, highlighting high scoring docking results. (d) Canonical model mean PDI100 versus median alternative isoform model means for 1,175 UniProt protein entries; dashed line, equality. (e) Spearman correlations for these paired summaries; 95% intervals from 2,000 paired bootstrap resamples. (f) Tested docking weights, sigmoid scales, PRI references and penalty strengths. Electrostatic weight remained 1−wD. (g) Number of PDI settings selecting each high scoring docking result (out of 135) versus its original PDI100. (h) Number of docking results shared between each top 185 selection and the original selection; ten count bins, dashed median. In c and g, circles/crosses distinguish normal/conditional cell surface exposure from other exposure statuses; pale points in c show other completed results. (i) Functional representation among entries with completed docking, high scoring results and normal/conditional cell surface exposure; annotations overlap. (j) Adjusted P values for enrichment among UniProt protein entries with high scoring docking results, compared with the remaining entries with completed docking. Fisher tests used all models or canonical models alone; conditional tests accounted for protein length and completed docking result count. Two sided tests used Benjamini–Hochberg adjustment [35] across 219 annotations per comparison; bold values indicate adjusted P <0.05. (k) Proportions of UniProt protein entries with ≥1 consistently selected docking result; functional groups are nonoverlapping within each population. Bar labels give counts and denominators. D = −AD4 score; E is positive patch electrostatic intensity; Base combines Dock01 and Elec01 before PRI.

The cell surface exposure review identified normal or conditional cell surface exposure for models contributing 74 of the 185 high scoring docking results, spanning 56 models and 49 UniProt protein entries (Figs. 1d and 4c). All 185 results were included in the ranking sensitivity and functional site analyses.

Related isoform models provided a further check of scoring consistency. Among UniProt protein entries with completed docking, 1,175 had a canonical model and at least one alternative isoform model. We averaged each score across each model’s completed docking results, then paired the canonical model mean with the median of the alternative model means, giving one comparison per UniProt protein entry (Supplementary Methods, “Comparing scores across isoform models”).

We compared these paired summaries using Spearman rank correlations for docking favorability D = −AD4 score, electrostatic intensity E, PRI, Base and PDI100 (Fig. 4d,e; Supplementary Table S1; Supplementary Data S5). Canonical model PDI100 means showed a strong positive association with the paired alternative isoform medians (ρ = 0.790; 95% bootstrap interval, 0.758–0.817). The same comparison for docking favorability gave ρ = 0.760. Favorable scores therefore tended to recur across related isoform models, including docking favorability before its integration into PDI100, supporting the internal consistency of our computational prioritization. We also examined associations between docking favorability and patch properties, and score variation between and within AlphaFold models (Supplementary Fig. S1; Supplementary Methods, “Separating patch size from composition”).

### A subset of high scoring docking results remained selected across all tested PDI settings

We tested how candidate selection depended on the manually chosen PDI settings. Across 135 parameter combinations, we varied four settings controlling the balance of docking, electrostatics and structural reliability. We tracked how often each of the original 185 high scoring docking results remained among the top 185. The original AlphaFold coordinates, isoform identities, docking boxes and saved polyP10 poses remained fixed.

The equations below show where each of the four settings in Fig. 4f enters PDI100.

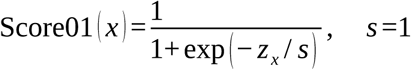

The sigmoid scale s was originally 1; each tested value was used in both transformations that convert D to Dock01 and E to Elec01.

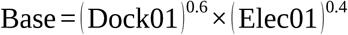

The docking weight wD was originally 0.6, with the electrostatic weight fixed at 1 − wD.

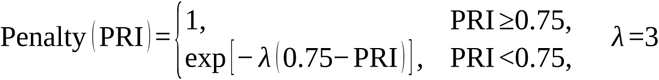

The PRI reference was originally 0.75 and the penalty strength λ was 3. Each tested reference replaced 0.75 in both conditions and the exponential term.

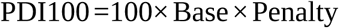

The final multiplication remained unchanged. Combining the tested values of the four settings in every possible way produced 135 combinations (5 × 3 × 3 × 3), including the original settings (Supplementary Data S3). The original D and E normalization references remained fixed across all 135 combinations (Supplementary Methods, “Defining completed and high scoring results”).

For each combination, we reranked all 19,481 completed results and selected the top 185. The cutoff of 80 was not reapplied because changing the score scale can change how many results exceed it. We tracked each of the original 185 high scoring docking results by its exact AlphaFold model and positive patch, including the 74 in the normal/conditional cell surface exposure group. For each result, the selection count records how many PDI combinations kept it in the top 185 (Fig. 4g; Supplementary Data S2; Supplementary Methods, “Tracking selection across 135 PDI settings”). 45 of the 185 high scoring docking results (24.3%) remained in the top 185 under all 135 combinations, spanning 34 AlphaFold models and 24 UniProt protein entries (Table 1). Within the normal/conditional cell surface exposure group, 19 of 74 results (25.7%) were consistently selected, covering 15 models and 14 entries. Among the other 111 results, 26 (23.4%) were consistently selected from 19 models and 13 entries. Of the 19 results in the exposure group, 18 came from canonical models and one from LIPI isoform 2; canonical LIPI also contributed a consistently selected docking result (Table 1).

**Table 1.** Docking results selected under all 135 PDI settings.

| Gene | UniProt ID | AlphaFold model | Cell surface exposure | Positive patch | Original PDI100 | Best–worst rank / 19,481 |
| --- | --- | --- | --- | --- | --- | --- |
| GPR26 | Q8NDV2 | AF-Q8NDV2-F1 | Yes | patch002 | 93.35 | 1–4 |
| CYTH4 | Q9UIA0 | AF-Q9UIA0-F1 | No | patch001 | 92.84 | 2–5 |
| MYO1G | B0I1T2 | AF-B0I1T2-F1 | No | patch001 | 92.18 | 1–23 |
| NOX1 | Q9Y5S8 | AF-Q9Y5S8-F1 | Yes | patch003 | 91.96 | 2–14 |
| MINPP1 | Q9UNW1 | AF-Q9UNW1-3-F1 | Review | patch001 | 91.47 | 2–171 |
| CYTH4 | Q9UIA0 | AF-Q9UIA0-F1 | No | patch002 | 90.84 | 4–14 |
| TAAR3P | Q9P1P4 | AF-Q9P1P4-F1 | Uncertain† | patch003 | 90.31 | 5–153 |
| NOX1 | Q9Y5S8 | AF-Q9Y5S8-F1 | Yes | patch002 | 90.16 | 5–20 |
| MYO1C | O00159 | AF-O00159-2-F1 | No | patch010 | 89.96 | 6–21 |
| MYO1C | O00159 | AF-O00159-2-F1 | No | patch017 | 89.42 | 4–35 |
| LIPI | Q6XZB0 | AF-Q6XZB0-6-F1 | Review | patch004 | 89.10 | 6–34 |
| GPR183 | P32249 | AF-P32249-F1 | Yes | patch001 | 88.23 | 4–149 |
| GNA12 | Q03113 | AF-Q03113-2-F1 | No | patch001 | 88.18 | 8–184 |
| ABCC2 | Q92887 | AF-Q92887-F1 | Yes | patch010 | 88.09 | 5–76 |
| DUOX2 | Q9NRD8 | AF-Q9NRD8-F1 | Yes | patch002 | 87.80 | 10–41 |
| FFAR1 | O14842 | AF-O14842-F1 | Yes | patch002 | 87.80 | 11–62 |
| NOX1 | Q9Y5S8 | AF-Q9Y5S8-3-F1 | Review | patch002 | 87.72 | 12–58 |
| MYO1C | O00159 | AF-O00159-3-F1 | No | patch010 | 87.72 | 10–47 |
| ABCC1 | P33527 | AF-P33527-8-F1 | Review | patch008 | 87.62 | 12–47 |
| NOD2 | Q9HC29 | AF-Q9HC29-2-F1 | No | patch002 | 87.10 | 13–97 |
| RGS6 | P49758 | AF-P49758-12-F1 | No | patch001 | 87.06 | 9–78 |
| NOX1 | Q9Y5S8 | AF-Q9Y5S8-3-F1 | Review | patch003 | 87.00 | 10–107 |
| LIPI | Q6XZB0 | AF-Q6XZB0-2-F1 | Yes | patch002 | 86.82 | 16–64 |
| MYO1C | O00159 | AF-O00159-F1 | No | patch003 | 86.75 | 18–88 |
| MYO1C | O00159 | AF-O00159-3-F1 | No | patch004 | 86.55 | 20–80 |
| LIPI | Q6XZB0 | AF-Q6XZB0-6-F1 | Review | patch002 | 86.25 | 19–88 |
| ABCC1 | P33527 | AF-P33527-2-F1 | Review | patch008 | 86.18 | 22–103 |
| RGS6 | P49758 | AF-P49758-14-F1 | No | patch001 | 86.04 | 17–108 |
| MYO1C | O00159 | AF-O00159-F1 | No | patch013 | 85.96 | 24–120 |
| ABCC2 | Q92887 | AF-Q92887-F1 | Yes | patch016 | 85.96 | 11–178 |
| LIPI | Q6XZB0 | AF-Q6XZB0-F1 | Yes | patch002 | 85.91 | 25–124 |
| NOD2 | Q9HC29 | AF-Q9HC29-F1 | No | patch002 | 85.90 | 16–154 |
| OPN5 | Q6U736 | AF-Q6U736-F1 | Yes | patch005 | 85.87 | 31–144 |
| SLC16A7 | O60669 | AF-O60669-F1 | Yes | patch003 | 85.85 | 23–91 |
| RGS6 | P49758 | AF-P49758-6-F1 | No | patch001 | 85.77 | 25–98 |
| DUOX1 | Q9NRD9 | AF-Q9NRD9-F1 | Yes | patch008 | 85.77 | 11–180 |
| DUOX1 | Q9NRD9 | AF-Q9NRD9-F1 | Yes | patch002 | 85.46 | 17–177 |
| ABCC8 | Q09428 | AF-Q09428-F1 | Yes | patch007 | 85.43 | 19–165 |
| SLC16A7 | O60669 | AF-O60669-F1 | Yes | patch002 | 85.28 | 22–140 |
| MYO1C | O00159 | AF-O00159-2-F1 | No | patch003 | 84.96 | 28–181 |
| NOX4 | Q9NPH5 | AF-Q9NPH5-F1 | Conditional | patch003 | 84.59 | 34–156 |
| NOX4 | Q9NPH5 | AF-Q9NPH5-8-F1 | Review | patch002 | 84.57 | 34–185 |
| TBCD | Q9BTW9 | AF-Q9BTW9-F1 | No | patch005 | 84.48 | 24–177 |
| ABCB6 | Q9NP58 | AF-Q9NP58-F1 | Yes | patch007 | 84.02 | 38–185 |
| ATP11C | Q8NB49 | AF-Q8NB49-F1 | Yes | patch009 | 83.55 | 40–184 |

All results remained in the top 185 of 19,481 completed results under all 135 settings. Models are AlphaFold version 6; append “-model\_v6\_” and the patch label to reconstruct patch\_id (Supplementary Data S2). YES/CONDITIONAL denote normal/conditional cell surface exposure; NO, intracellular or inward facing localization; REVIEW, unresolved exposure. These cell surface exposure labels do not establish access to the saved pose. Three UniProt protein entries have models in both exposure groups. † TAAR3P is a pseudogene with uncertain protein existence.

For these 45 results, top 185 membership was stable within the tested parameter ranges even when individual ranks changed. GPR26 patch002 ranked 1st–4th, whereas GPR183 patch001 ranked 4th– 149th. Across all 135 settings, each top 185 selection shared 105–185 results with the original selection, with a median of 139 (Fig. 4h; Supplementary Data S2 and S3).

### Oxidoreductase and translocase annotations were enriched among proteins with high scoring docking results

We tested functional enrichment by comparing annotations for the 82 UniProt protein entries with high scoring docking results against those for the remaining 2,843 of the 2,925 entries with completed docking. Each UniProt protein entry was counted once in the two sided Fisher exact tests. We also described these annotations for the 49 entries represented in the normal/conditional cell surface exposure group.

Oxidoreductases and translocases showed enrichment factors of 8.39 and 6.69, respectively (Fig. 4i). Oxidoreductases accounted for 8/82 UniProt protein entries with high scoring docking results versus 34/2,925 with completed docking; translocases accounted for 9/82 versus 48/2,925. ATP binding and lipid transport were also significantly overrepresented in the initial Fisher tests (adjusted P = 0.00152 and 0.00271). Heparin binding and G protein coupled receptor annotations were more frequent among selected entries (3.7% versus 0.5% and 25.6% versus 20.5%, respectively), but neither was significant after adjustment (P = 0.104 and 0.933; Fig. 4i,j; Supplementary Table S2).

We then tested whether the enrichment reflected differences in protein length or in the numbers of isoform models and completed docking searches per protein. First, tests restricted to canonical models compared 64 UniProt protein entries with high scoring canonical model results against the remaining 2,533 entries with completed canonical docking.

Second, we accounted for sequence length and completed docking result count across all 2,925 UniProt protein entries. We compared observed annotation counts with those expected under random selection within 15 groups, defined by three sequence length ranges and five search count ranges. All three analyses applied Benjamini–Hochberg adjustment across 219 functional annotations (Supplementary Methods, “Testing enrichment with sampling controls”; Supplementary Data S4).

Of the six categories shown, only oxidoreductases and translocases remained significantly enriched both when alternative isoform results were excluded and when sequence length and completed docking result count were accounted for. Adjusted P values after accounting for length and search count were 0.000133 and 0.0317, respectively (Fig. 4j).

Six of the eight oxidoreductases and eight of the nine translocases had high scoring docking results from models in the normal/conditional cell surface exposure group (Supplementary Tables S3, S4 and S8; Supplementary Data S8). All six oxidoreductases belonged to the NOX/DUOX family; the eight translocases comprised five ABC transporters and three phospholipid transporting ATPases. Twelve of these 14 proteins had normal cell surface exposure (YES); NOX4 and ATP9A had exposure under defined biological circumstances (CONDITIONAL).

Four of the eight oxidoreductases and four of the nine translocases had at least one docking result selected under all 135 PDI settings, compared with 16 of the other 65 UniProt protein entries (Fig. 4k). These eight entries—NOX1, NOX4, DUOX1, DUOX2, ABCB6, ABCC1, ABCC2 and ATP11C—contributed 15 of the 45 consistently selected docking results across 11 AlphaFold models (Table 1; Supplementary Table S3). Within the normal/conditional cell surface exposure group, four of six oxidoreductases and three of eight translocases contributed 10 of the 19 consistently selected docking results; 7 of the other 35 entries also had a consistently selected docking result. Enrichment in all three analyses and consistent selection across the tested PDI settings strengthen the computational basis for nominating these eight entries as polyP binding candidates.

FAD, NADP and flavoprotein annotations occurred in six, seven and six of the eight oxidoreductases, respectively. All nine translocases carried ATP binding annotations and eight also carried lipid transport annotations. The 82 UniProt protein entries with high scoring docking results also included 21 G protein coupled receptors and the lipases LPL, LIPI and LIPH (Fig. 4i; Supplementary Tables S7 and S8).

### Consistently selected polyP10 poses overlapped functional sites

Across the 185 high scoring docking results, mapped experimental residue sets were available for 113 saved polyP10 poses from 74 AlphaFold models and 44 UniProt protein entries. Of these 113 poses, 31 from 23 models and 18 entries contacted at least one residue in a specified experimental set (Table 2; Fig. 1c; Supplementary Data S6, S10 and S12). The sets comprised experimentally identified catalytic, ligand binding or regulatory sites. We mapped them to the exact AlphaFold model sequences and assessed overlap using a 4 Å nonhydrogen atom contact criterion, distinguishing structural contacts from mutation dependent functional effects. Mapping procedures and evidence coverage are given in Supplementary Methods, “Mapping experimental sites to saved poses”. Experimental context for all 18 entries is provided in Supplementary Notes, “PolyP poses overlapped functional sites” and “LPL contacts overlapped heparin affinity determinants”.

**Table 2.**
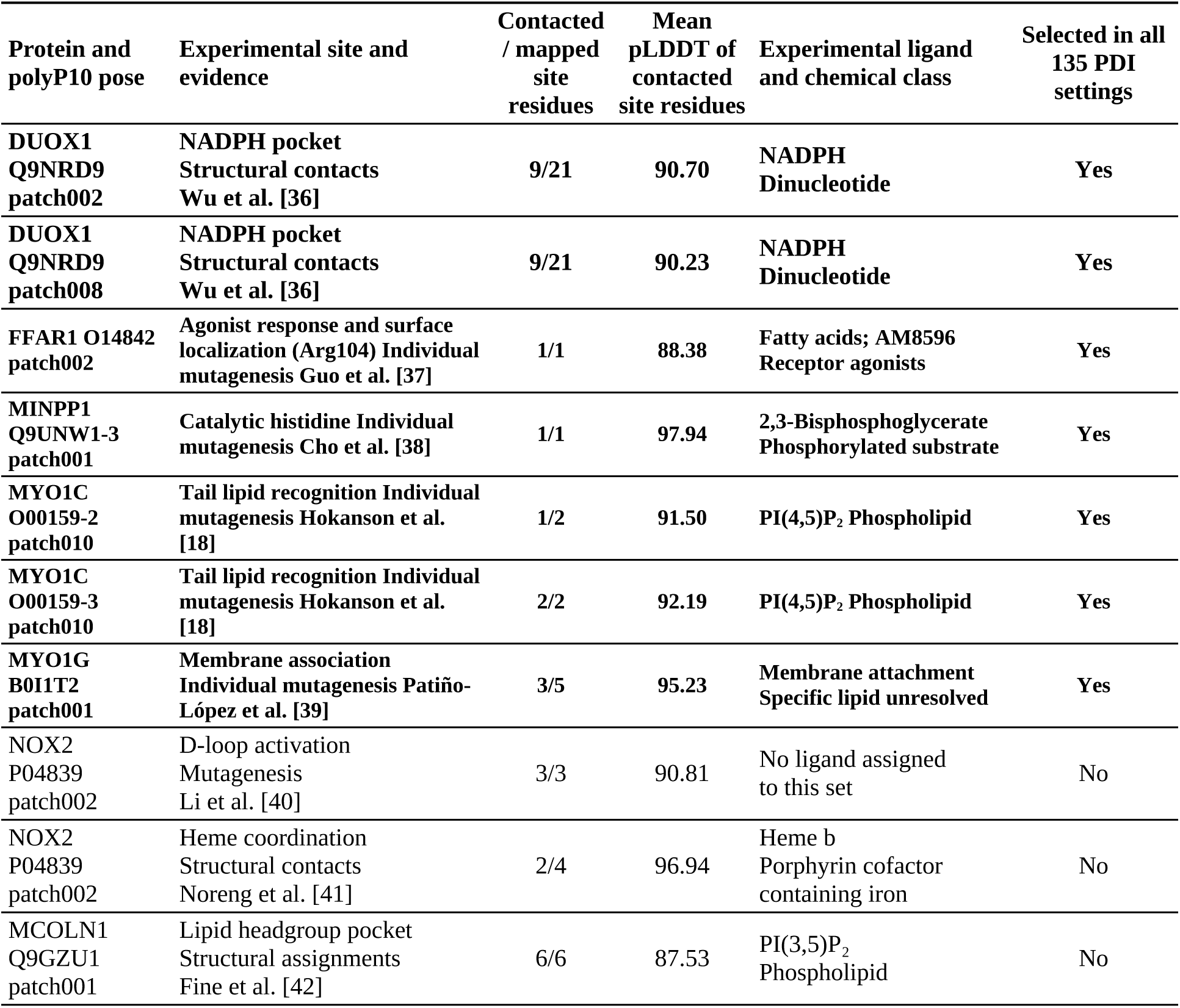
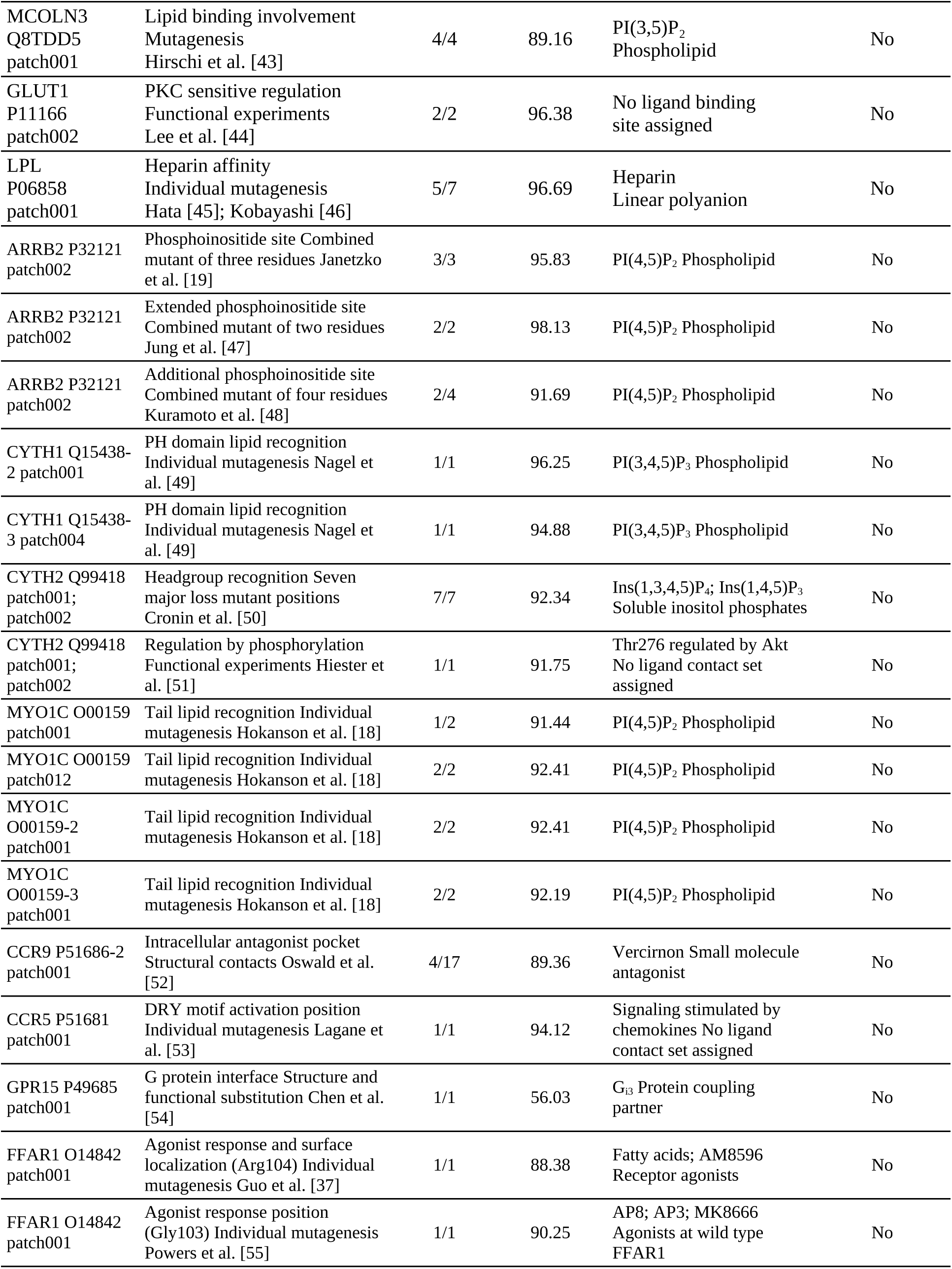
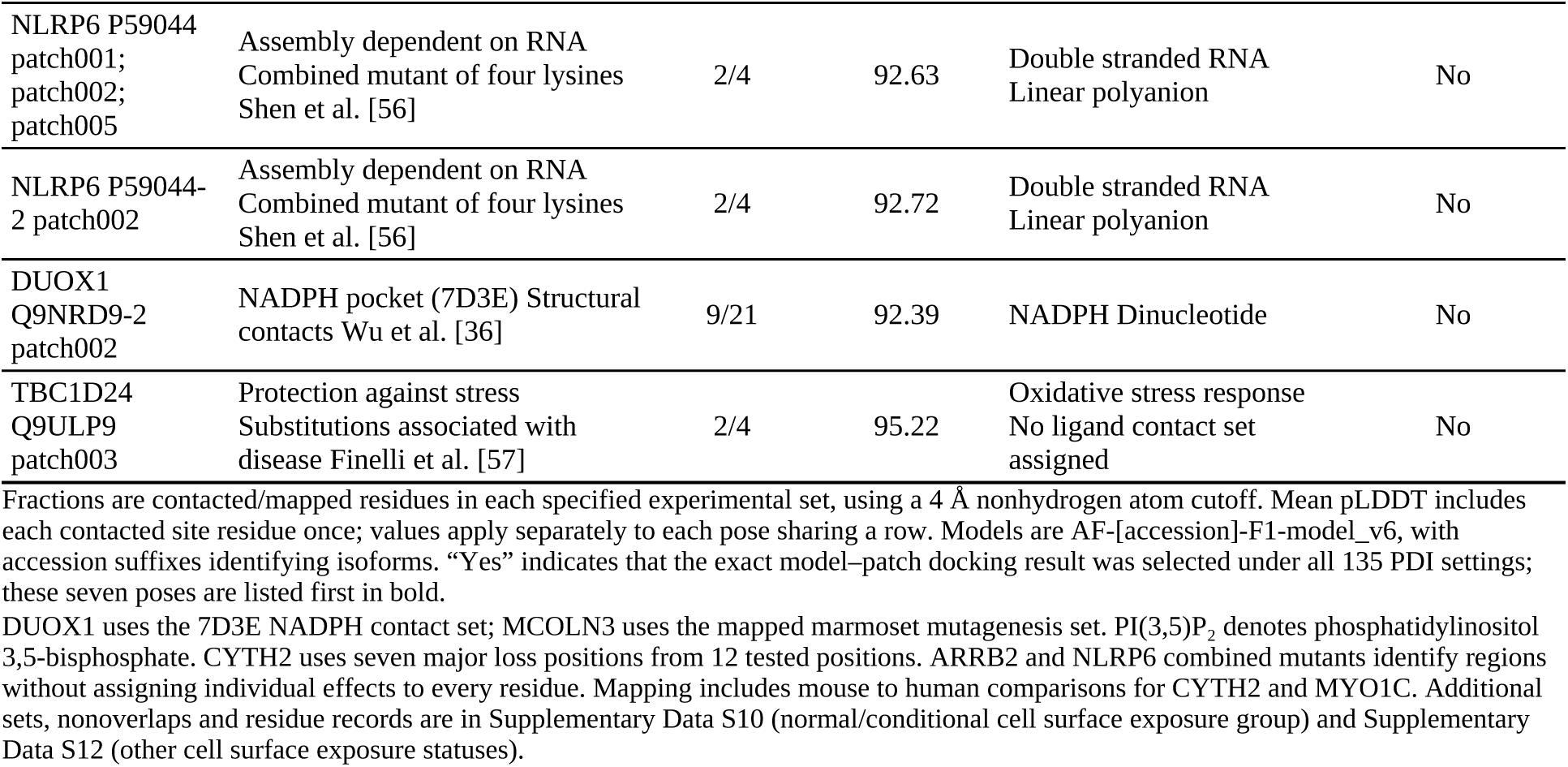
PolyP10 pose overlaps with experimentally identified functional sites.

Seven of the 31 overlapping poses were also selected under all 135 PDI settings, spanning six AlphaFold models and five UniProt protein entries. The six models were canonical DUOX1, FFAR1 and MYO1G, MINPP1 isoform 3, and MYO1C isoforms 2 and 3; DUOX1 contributed two poses (Fig. 5a–g; Supplementary Table S5; Supplementary Data S13).

**Figure 5.**
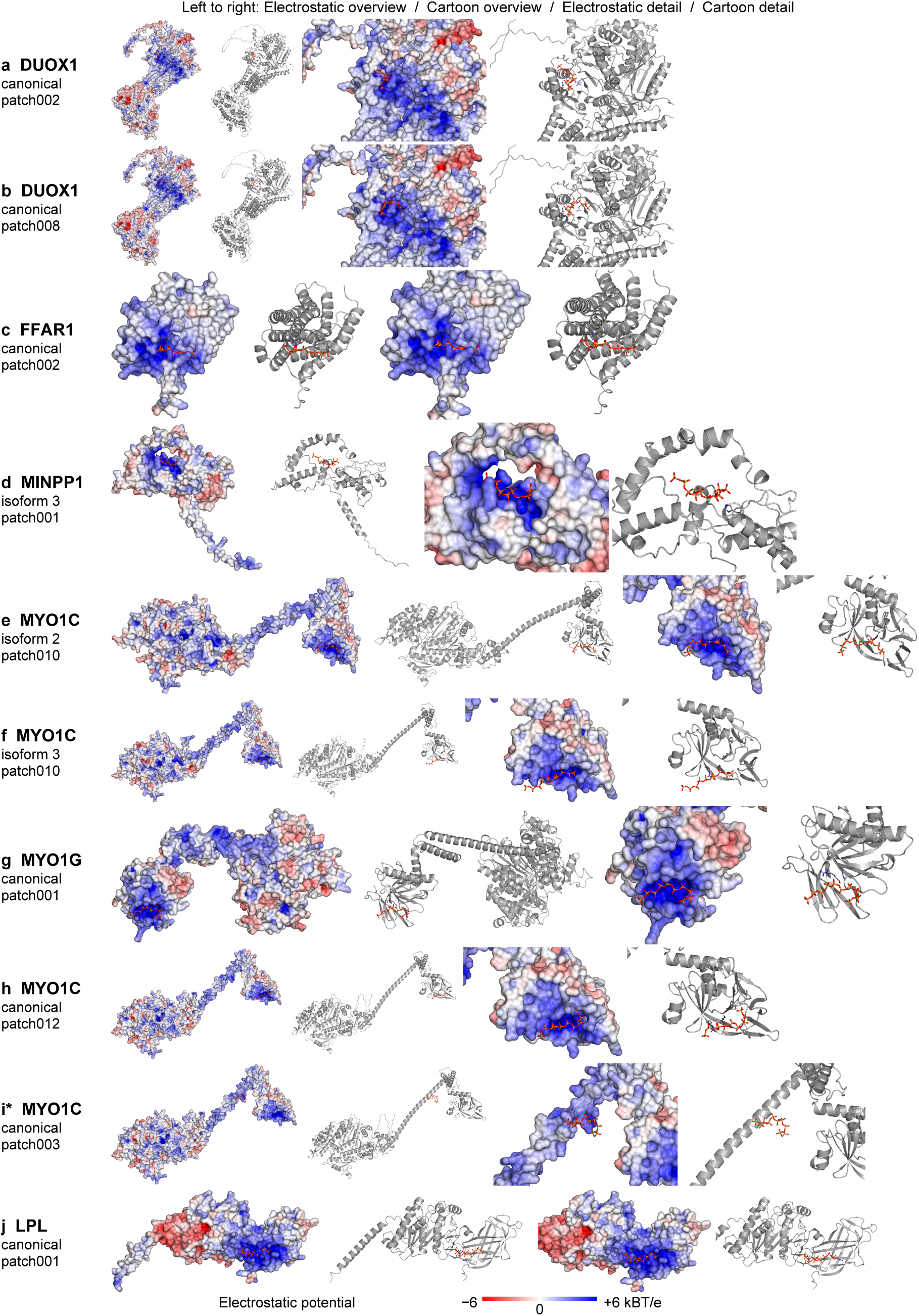
PolyP10 docking poses overlapped functional sites in selected candidates. Each row shows matched electrostatic surface and cartoon overviews and details of one pose. Light gray cartoons show the protein. Overlapping functional site positions are shown as sticks with darker gray carbon, blue nitrogen and red oxygen. PolyP10 sticks show orange phosphorus and red oxygen. Electrostatic potentials use a shared scale: −6 (red), 0 (white) and +6 (blue) kBT/e. Poses a–g were selected under all 135 PDI settings. In canonical MYO1C, h shows the overlapping patch012 pose; i* shows patch003, selected under all 135 PDI settings without functional site overlap. LPL (j) provided the separate heparin affinity comparison. Functional site overlap used nonhydrogen atom distances ≤4 Å (Table 2; Supplementary Data S6, S10 and S12). Images were prepared with PyMOL 3.1.0 (https://pymol.org/).

Both consistently selected DUOX1 poses, patch002 and patch008, overlapped the experimentally resolved cytosolic NADPH pocket, including its phosphate recognition residues Arg1036, Arg1424 and Arg1495 [36]. Each overlapped 9/21 pocket residues in the low calcium structure 7D3E (Table 2; Fig. 5a,b). NADPH supplies electrons for oxidase activity, linking the predicted interfaces to NADPH recognition.

The consistently selected MYO1C patch010 poses contacted one of two mapped membrane binding positions in isoform 2 and both in isoform 3 (Table 2; Fig. 5e,f). Substitutions at the corresponding mouse residues impaired phosphatidylinositol 4,5-bisphosphate [PI(4,5)P₂] dependent membrane binding and cellular localization [18]. The MYO1G patch001 pose contacted three of five positions implicated in membrane association [39] (Table 2; Fig. 5g).

The consistently selected FFAR1 patch002 pose contacted Arg104 (Table 2; Fig. 5c), where substitution impaired agonist signaling and cell surface localization [37].

The MINPP1 isoform 3 patch001 pose contacted catalytic His89 (Fig. 5d); substitution of the corresponding histidine impaired phosphatase activity in the experimental full length protein [38]. Interpretation of this overlap depends on the activity of this truncated isoform (Table 2).

Canonical MYO1C was a seventh AlphaFold model with both consistent selection and functional site overlap, but through different poses: patch003 and patch013 were consistently selected, whereas patch001 and patch012 overlapped the membrane binding positions (Fig. 1c; Supplementary Table S5). The overlapping patch012 pose and consistently selected patch003 pose are shown separately (Fig. 5h,i*).

### The LPL pose linked distributed polyP contacts to heparin affinity

PolyP and other anionic polymers share repeated negative charges that can engage positively charged protein surfaces (Supplementary Notes, “LPL contacts overlapped heparin affinity determinants”).

The literature review of the 49 UniProt protein entries in the normal/conditional cell surface exposure group identified five with experimental heparin affinity or recognition by a specific aptamer (Supplementary Table S9). Mutagenesis identified heparin affinity determinants in LPL [45, 46], while LIPI and LIPH were retained on heparin columns [58]. CCR5 and GLUT1 showed target dependent cell binding by selected RNA and DNA aptamers, respectively [59, 60]. Each of these five entries contributed a high scoring docking result from a model classified as YES.

LIPI’s canonical and isoform 2 docking results remained in the top 185 under all 135 PDI settings. The LPL, LIPH, CCR5 and GLUT1 docking results remained in the top 185 under 79, 106, 113 and 113 settings, respectively (Supplementary Data S2; Supplementary Notes, “LIPI contacts spanned the phosphate chain”).

Individual heparin affinity determinants had been mapped experimentally for LPL. The canonical LPL patch001 result had PDI100 80.63, AD4 score −24.12 kcal/mol and PRI 0.871.

The LPL pose contacted three of five N-terminal heparin affinity determinants from Hata et al. [45], Arg306, Lys307 and Arg309, and both C-terminal determinants from Kobayashi et al. [46], Arg432 and Lys434 (Fig. 6b). Residue numbers refer to the precursor model; adding 27 to the mature chain positions accounts for the signal peptide.

**Figure 6.**
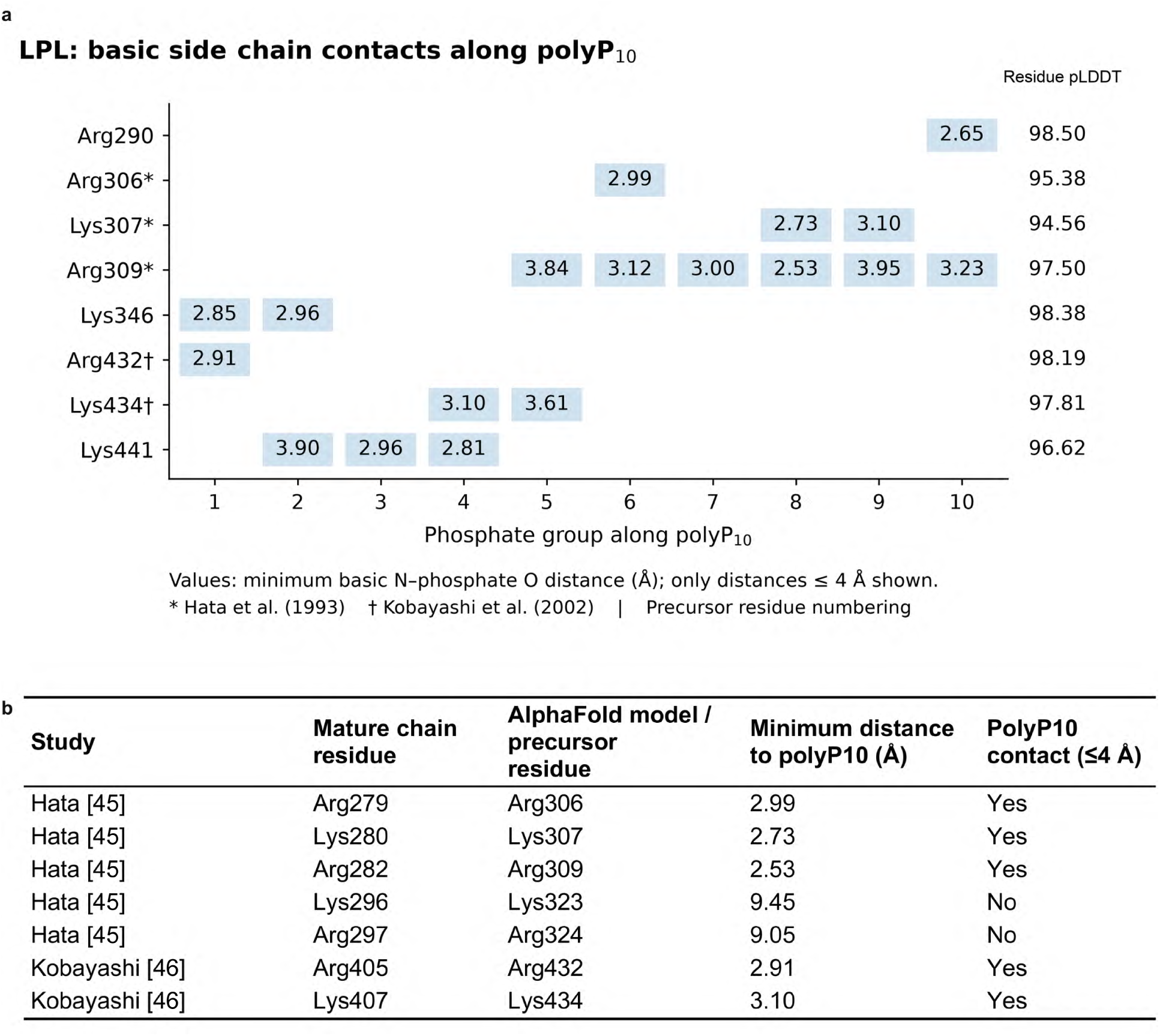
The saved LPL polyP10 pose contacted heparin affinity determinants. (a) Contacts along AF-P06858-F1-model_v6, patch001. Rows show eight contacting Lys/Arg residues in precursor numbering; columns follow the ten phosphate groups. Values are minimum distances in Å from Lys NZ or Arg NE/NH1/NH2 to an oxygen assigned to each phosphate; only distances ≤4 Å before rounding are shown. The final column gives residue pLDDT. Asterisks identify Hata overlaps and daggers Kobayashi overlaps. Shared bridging oxygens may contribute to adjacent columns; the nonbridging oxygen check also retained coverage of all ten phosphates. (b) Minimum nonhydrogen atom distances from seven experimentally determined heparin affinity determinants to the saved polyP10 pose. “Yes” marks distances ≤4 Å. Mature chain and precursor residue numbers are shown separately (precursor = mature chain +27). The first five determinants were mapped by Hata et al. [45], the last two by Kobayashi et al. [46]. Values are rounded to two decimals. Atom specific calculations for panel a are in Supplementary Data S11; the residue distances in panel b are in Supplementary Data S9 and S10. Source coordinates are in Supplementary_Structures.zip.

The predicted interface therefore included five of seven individually identified heparin affinity determinants (71.4%) across the N-terminal and C-terminal regions (Fig. 5j). Their mean pLDDT was 96.69 (range 94.56–98.19), and the 5/7 overlap persisted at both 3.5 and 5 Å (Supplementary Data S9). Supplementary Table S6 separately lists paired substitutions and individual substitutions with no affinity effect; construct details are given in Supplementary Notes, “LPL contacts overlapped heparin affinity determinants”.

Eight of the 11 LPL residues contacting polyP were Lys/Arg residues, each with a basic side chain nitrogen within 4 Å of a polyP oxygen (Fig. 6a).

Together, these nitrogens contacted all ten phosphate groups, including when shared bridging oxygens were excluded; nine phosphates retained contacts at 3.5 Å (Supplementary Data S11). The eight basic contact residues had mean pLDDT 97.12 (Supplementary Methods, “Resolving LPL contacts along the phosphate chain”; Supplementary Data S11).

The pose was spatially separate from the catalytic triad (Supplementary Data S9); experimental substitutions at the catalytic triad residues abolished activity without changing heparin affinity [61]. This arrangement links the distributed polyP contacts to the heparin recognition surface and provides the clearest experimental context for a shared polyanion recognition mechanism and possible competition at the LPL surface (Supplementary Notes, “LPL contacts overlapped heparin affinity determinants”).

## Discussion

This study nominated human membrane protein candidates for noncovalent polyphosphate (polyP) binding by combining surface electrostatics, docking of a ten phosphate chain (polyP10) and AlphaFold model confidence. Stringent selection yielded 185 high scoring docking results across 128 AlphaFold models representing 82 UniProt protein entries. The selected candidates included predicted interfaces at experimentally characterized recognition and regulatory sites. These correspondences suggested that polyP could influence native ligand responses or protein localization through occupation of existing recognition and regulatory sites.

### Surface screening nominated polyP binding candidates

Three dimensional surface recognition extended the search for polyP binding candidates beyond contiguous basic sequences. Experiments on histidine and lysine repeats established noncovalent polyP association, including ionic interactions sensitive to lysine protonation and ionic strength [12, 13]. A recent human protein screen used lysine rich sequences to identify polyP interactions [15]. PPK1 contains a positively charged tunnel proposed to engage polyP, and ligand bound class I PPK2 shows polyP in a channel formed by separated structural elements [7, 8]. Here, analysis of continuous positive patches [25] under near physiological solution conditions guided docking of an explicit phosphate chain. This approach allowed basic residues separated in sequence to contribute to the same predicted interface when brought together by protein folding. It complemented sequence based screening by examining whether surface geometry and local interactions could accommodate the repeated negative charges of polyP.

The integrated selection criteria focused discovery on candidates with strong combined electrostatic, docking and structural confidence evidence. The Patch Docking Index (PDI100) combined electrostatic intensity and docking favorability, while the pLDDT Reliability Index (PRI) reduced the priority of docking results from model regions with lower structural confidence. This penalty changed which results were selected, making structural confidence an active component of prioritization. The PDI100 ≥80 cutoff retained the top 0.95% of completed docking results. The stringent screen was designed to minimize false positive nominations; it was never intended to recover every polyP binding protein. PDI100 therefore orders docking results for candidate nomination without estimating binding affinity or binding probability, and nonselection leaves binding unresolved.

Consistent selection across alternative PDI settings strengthened the computational basis for a subset of candidates. At a fixed selection size, 45 docking results across 34 AlphaFold models representing 24 UniProt protein entries remained selected under all 135 tested parameter combinations. These results remained selected when the relative contributions of docking, electrostatics and structural confidence changed, so their selection did not depend on a single combination within the ranges examined. Similar scoring tendencies across related isoform models provided a further check of internal consistency (Table 1; Supplementary Table S1). Together, the explicit selection criteria and sensitivity analysis provide a reproducible approach for extending polyP candidate discovery to other protein sets.

### Functional site overlaps suggested routes to regulation

Consistent selection showed that a docking result remained prioritized across the tested scoring settings; functional site overlap placed predicted interfaces from 18 UniProt protein entries at residues experimentally implicated in protein function. Seven poses across six AlphaFold models representing five UniProt protein entries satisfied both criteria, providing a focused set of candidate interfaces for noncovalent polyP mediated regulation (Table 2; Fig. 5a–g). A seventh AlphaFold model, canonical MYO1C, met the two criteria through different poses (Figs. 1c and 5h,i*).

Shared recognition of phosphate groups offers one explanation for these functional site correspondences. In dual oxidase 1 (DUOX1) and the MCOLN1/TRPML1 channel, saved polyP10 poses overlapped recognition sites for NADPH and the regulatory phospholipid PI(3,5)P₂, respectively (Table 2) [36, 42, 62]. These chemically distinct ligands both present phosphate groups to basic residues, providing a common basis for accommodation of a phosphate chain. PolyP occupation could therefore affect access to a native ligand or the response normally coupled to its binding. The MCOLN1 comparison illustrates why these possibilities extend beyond simple obstruction: PI(3,5)P₂ and PI(4,5)P₂ occupy a shared pocket but exert opposing effects on channel activity [42]. A shared interface can thus support different functional responses, making the coupling between polyP recognition and protein activity a central biological question.

Phosphate recognition chemistry also provides a plausible connection between the individual interfaces and the enrichment of oxidoreductases and translocases. Both classes remained overrepresented after restricting the comparison to canonical AlphaFold models and after accounting for protein length and completed docking result count. Most selected oxidoreductases had annotations for phosphate containing cofactors, while the selected translocases had ATP binding annotations (Fig. 4i,j; Supplementary Table S8). A screen centered on strongly positive surfaces could preferentially recover proteins with native anion recognition sites. The enrichment, together with the DUOX1 overlap, therefore suggests that recurring recognition chemistry contributed to selection across proteins with different functions.

The membrane association overlaps suggested that polyP could also influence where proteins act. In the class I myosins MYO1C and MYO1G, the contacted positions were experimentally linked to membrane attachment [18, 39]. Related overlaps in β-arrestin 2 (ARRB2) involved phosphoinositide recognition regions (Table 2) [19, 47, 48]. Phosphoinositides regulate the assembly and persistence of receptor–β-arrestin complexes, linking lipid recognition to the spatial and temporal organization of signaling [19]. These comparisons place possible polyP effects at recruitment and retention interfaces, where altered occupancy could change protein localization and partner engagement without directly occupying a catalytic pocket.

### The LPL pose overlapped heparin affinity determinants

Lipoprotein lipase (LPL) provided a complementary case from the broader functional site overlap group, connecting the polyP based screen to an experimentally characterized surface for another anionic polymer. Its saved polyP10 pose contacted five of seven experimentally determined heparin affinity determinants from the two studies used for the comparison, distributed across the N-terminal and C-terminal regions [45, 46]. Neither heparin nor these determinants was used to generate the pose or calculate PDI100. Contacts distributed along the phosphate chain (Fig. 6a) further suggested that this surface could accommodate the repeated negative charges of an extended polymer.

Experimental polyP binding to thrombin exosite II, at a surface partially overlapping its heparin binding site, provides a precedent for this shared recognition [9]. A published proposal of polyP recognition at an RNA binding region of SARS-CoV-2 polymerase offers a further comparison [10]; the present NLRP6 overlaps likewise involved a basic region implicated in RNA dependent assembly [56]. These comparisons suggest that a protein surface capable of engaging multiple anionic groups can accommodate chemically distinct polymers. Polymer chemistry and geometry would then determine the particular contacts and consequences of occupation.

For LPL, occupation of a noncatalytic surface could influence the location of lipolysis. LPL hydrolyzes triglycerides in circulating lipoproteins at the capillary lumen, making its delivery to this site integral to its biological activity [63]. Heparan sulfate proteoglycans retain LPL through electrostatic interactions, whereas glycosylphosphatidylinositol anchored high density lipoprotein binding protein 1 (GPIHBP1) transports LPL to the lumen and stabilizes the enzyme [16, 63].

GPIHBP1’s acidic domain shields the basic LPL surface, enabling release from heparan sulfate retention during transport [63]. The position of the polyP10 pose away from the catalytic triad suggested that polyP could affect delivery of LPL to its site of action through surface retention or partner engagement.

### Limitations

The screen applied stringent computational criteria informed by structural evidence for polyP recognition [7, 8] to minimize false positive nominations. It was never intended to recover every polyP binding protein or minimize false negatives; proteins outside the selected set may also bind polyP.

The screen represented local interactions of polyP10 with rigid AlphaFold models under specified solution conditions. Electrostatic calculations included solvent dielectric response and ionic screening; native membranes, glycans, cofactors and binding partners were absent from the predicted complexes. These components can alter access to or occupy a predicted interface, and a pose within the membrane interior may be inaccessible to polyP. Cell surface exposure labels for proteins or isoforms do not establish access to each saved polyP10 pose. The PRI penalty reduced the priority of uncertain docking regions but did not alter their coordinates or establish confidence at every final ligand contact.

Functional interpretation depended on the available experimental residue sets and their construct context. Some evidence came from another species, isoform or protein fragment; the catalytic histidine overlap in truncated MINPP1 isoform 3, for example, does not establish retention of the activity measured for the full length enzyme. Candidates without mapped residue sets could not be assessed by this comparison. PolyP binding and functional responses were not measured for the nominated candidates. Association with a structural surface may leave function unchanged, while regulatory effects would depend on the interface and biological context.

### Future directions

Comparisons across polyP chain lengths and protein conformations can distinguish contacts that persist across structural representations from those specific to the screened model. Incorporating native membranes and relevant molecular partners would clarify how access to a predicted interface changes when surrounding structures or competing ligands are represented.

Biological investigation can distinguish effects on native ligand responses from effects on protein localization. Comparisons between polyP and native anionic ligands at a nominated interface would clarify whether shared recognition produces similar or distinct functional responses. For membrane association interfaces, the corresponding question is whether polyP changes recruitment or partner dependent retention. LPL provides a setting in which binding, catalytic activity and localization can be considered separately. These studies would determine whether polyP association affects redox regulation, membrane signaling or lipid metabolism, and which interfaces mediate any response.

### Conclusions

This study established a new reproducible computational workflow and nominated candidates for noncovalent polyP binding among human membrane proteins. By connecting surface electrostatics to explicit ligand poses and experimentally characterized sites, the workflow provided a basis for studying both candidate recognition and its possible functional consequences. The findings support shared recognition of anionic groups as one explanation for the predicted interfaces and point to native ligand responses and protein localization as distinct routes through which polyP could influence protein function.

## Supporting information

Supplementary Structures

## Data availability

Screening results, candidate lists and contact analyses are supplied in Supplementary_Data.zip. Supplementary_Structures.zip contains the saved polyP10 poses for all 19,481 completed docking searches and the 128 AlphaFold models for the 185 selected poses. The Supplementary Information index and the file guides identify the records supporting each analysis.

## Author contributions

J.W. developed the hypothesis, designed the computational workflow and subsequent analyses, performed the screening and analyses, prepared the figures, organized the supplementary files and wrote the manuscript. Z.J. provided access to computational resources through Queen’s University’s Centre for Advanced Computing, contributed scientific insights and provided critiques, feedback and advice on manuscript drafting and revision.

