## Supplementary Information for "A structural screen for candidate polyphosphate binding surfaces in human membrane associated proteins"

Jianlin Wu and Zongchao Jia

Implementation details, supporting analyses and experimental context for the predicted polyP–protein contacts. Complete candidate records are provided in the tables and separate data files.

### Tables, figure and data files

Table S1. Scores were consistent across canonical and alternative isoform models............................................8
Table S2. Oxidoreductase and translocase enrichment persisted across sampling controls...............................10
Table S3. Consistent selection within oxidoreductase and translocase candidates............................................10
Table S4. Alternative isoforms changed the highest scores for NOX5 and DUOX1..........................................11
Table S5. Seven AlphaFold models combined consistent selection and functional site overlap........................12
Table S6. Additional LPL substitutions were separate from the seven affinity determinants............................16
Table S7. All 82 UniProt protein entries with high scoring docking results......................................................16
Table S8. Oxidoreductase and translocase candidates.......................................................................................18
Table S9. Heparin affinity and aptamer recognition in five candidate UniProt protein entries..........................20
Figure S1. Docking differed from positive patch properties.................................................................................9

#### Description of additional supplementary files

Supplementary Data labels identify the records below; filenames are unchanged. S1–S6 and S8–S13 are in their corresponding folders in Supplementary_Data.zip. S7 identifies inputs and provenance records in Supplementary_Code.zip. File guide.txt supplies the full paths.

**Data S1.** All 23,383 positive patch records, including incomplete docking outputs. Files: grand_patch_table_PDI100_polyP10.csv and grand_patch_table_PDI100_polyP10.xlsx.

**Data S2.** All 185 high scoring results: exact model and patch identifiers, full precision scores, cell surface exposure, and selection counts and rank summaries across 135 settings. Files: Data_S2_V11.7.csv.

**Data S3.** The 135 PDI combinations and their selection summaries. Files: Data_S3_V11.7.csv.

**Data S4.** All 219 functional annotation comparisons and sampling controls. Files: functional_keyword_enrichment_all219.csv.

**Data S5.** The 1,175 paired canonical and alternative isoform score summaries. Files: isoform_pairs_reconstructed.csv.

**Data S6.** Nonhydrogen atom contacts and exact model and patch identifiers for all 185 selected poses. Files: contacts_185_4A.csv.

**Data S7.** Screening inputs and provenance records. Paths below are relative to Supplementary code/ in Supplementary_Code.zip.

**Retrieval records.** Screening workflow/Inputs/ contains surface_candidates_uid.list (UniProt accessions), pdb_list.txt (retrieved model filenames), and alphafold_summary.csv.xlsx (retrieval summary and source URLs).

**Screening records.** Screening workflow/Inputs/Original patch records.tar.gz contains the original patch and residue lists. Functional analyses/sources/ contains vina_patch_boxes_20260219_070756.csv (box records), vina_patch_boxes_20260219_070756_PRI_only.csv (PRI records), and vina_results_ad4_polyP10.csv (saved docking scores).

**Annotations and exposure review.** Functional analyses/sources/surface_candidates_annotation_snapshot.csv retains the annotation snapshot. PDI selection/inputs/ contains Entry_Audit_128.csv and curated74.csv. The final per-result exposure labels are in Data S2.

**Ligand inputs.** Screening workflow/source_polyP10/ contains polyP10.sdf and polyP10.pdbqt.

**Data S8.** Oxidoreductase, translocase and NOX/DUOX records, with scores and selection counts. Files: Data_S8_V11.7.csv and NOX_DUOX_family.csv.

**Data S9.** LPL and LIPI residue contact distances using all residue nonhydrogen atoms. Files: LPL_LIPI_residue_contacts.csv.

**Data S10.** Contacts, experimental sets, sources and sequence maps for the 74 exposure group poses; 41 pose–set comparisons and 250 residue rows include noncontacts. Files: PolyP_Experimental_Site_Overlap_74_Poses.xlsx, TRPML3_ortholog_mapping.json, TRPML1_mouse_ortholog_mapping.json and NCC_construct_mapping.json.

**Data S11.** LPL basic nitrogen–phosphate oxygen maps for main Fig. 6a, phosphate connectivity, contact cutoff checks and local pLDDT. Files: LPL_LIPI_basic_nitrogen_phosphate_distances.csv and LPL_LIPI_phosphate_atoms.csv.

**Data S12.** Contacts, experimental sets, sources and sequence maps for the other 111 poses. Together, S10 and S12 cover all 185 poses. Files: PolyP_Experimental_Site_Overlap_111_Poses.xlsx and priority_mapping_corrections.json.

**Data S13.** Evidence group membership for all 128 models and the exact poses supporting selection consistency and functional site overlap. Files: PolyP_Data_S13_Model_Evidence_Groups_V13.5.8.csv.

Saved structures. Supplementary_Structures.zip contains Candidate structures/, with Candidate file index.csv, Protein models/ (128 AlphaFold models) and Saved docking poses/ (19,481 saved polyP10 PDBQT poses). The index links all 185 selected poses to their models and patches. Data S9 and S11 were reconstructed from these unchanged coordinates and checked against the reported contacts.

**Source data.** The Figure S1 Patch properties folder in Supplementary_Data.zip contains composition_docking_diagnostics.csv and within_vs_between_model_variation.csv.

**Code.** Supplementary_Code.zip contains the screening and analysis scripts and Run instructions.txt. The instructions identify the supplied entry points, inputs, retained source records and additional dependencies.

### Supplementary Methods

#### Preparing electrostatic maps

Electrostatic mapping identified positive protein surfaces that could attract anionic polyP. PPK1 contains a positively charged tunnel proposed to engage polyP, and a ligand bound class I PPK2 structure shows polyP positioned by basic residues in a folded channel; these structures provided the biological rationale for this step [1, 2]. PDB2PQR 3.7.1 [3] used PROPKA [4, 5] at pH 7.4 and the PARSE force field to assign protonation states, atomic charges and radii. The resulting PQR files supplied these quantities and atomic coordinates to APBS; pdb2pqr_array.slurm ran the preparation in batches on Queen’s CAC Frontenac.

APBS [3] represented the protein and surrounding solution as continuous dielectric media and included dissolved ions through the Poisson–Boltzmann equation. Calculations used 310.15 K, 0.14 M monovalent ionic strength, protein and solvent dielectric constants of 4.0 and 74.2, and a 1.4 Å solvent probe, as specified in the main Methods. The script inputgen_call.py generated and adjusted the APBS settings files. The batch script apbs.slurm ran APBS 1.3 to calculate a three dimensional electrostatic potential map for each prepared model.

#### Linking positive patches to docking boxes

PEP-Patch mapped the APBS potentials onto a triangular protein surface mesh. It interpolated the potential at each mesh vertex and grouped connected vertices above the positive potential cutoff into patches. The main Methods specify the +2.7 kT/e potential cutoff and >0.7 nm² area threshold. The patch list output recorded each patch’s identifier, sign, vertex count, area and summed vertex potential; the residue list output identified its constituent residues. The colored surface mesh was retained for visual inspection.

The docking box file linked each positive patch to its AlphaFold model, center coordinates, dimensions, patch extent and applied padding. The separate PRI file recorded the same model and patch identifier, the pLDDT Reliability Index (PRI), the mean and standard deviation of residue weights, and the included residue and atom counts. These identifiers joined the electrostatic, structural confidence and docking outputs without merging patches or isoforms (Supplementary Data S7).

In the master spreadsheet, UniProt_ID identifies the UniProt protein entry, Entry_ID identifies the AlphaFold model, and patch_id identifies the model and positive patch together. The fields patch_area and patch_value_total give the patch area in nm² and the unweighted sum of its vertex potentials. Their ratio, patch_value_per_area, is electrostatic intensity E. The field dock_best_score_ad4 contains the saved pose’s AD4 score in kcal/mol; docking favorability D is its negative. Dock01 and Elec01 contain the scaled 0–1 scores used to calculate PDI100. Supplementary Data S1 contains the master table; the main Methods define PRI and the PDI100 equations.

##### Searching polyP10 poses with AD4 maps

Meeko [6] prepared receptor PDBQT files containing coordinates, partial charges and AutoDock atom types. Each patch specific Grid Parameter File specified the receptor, docking box center, grid dimensions and spacing. AutoGrid4 calculated an interaction energy map for every atom type in the polyP10 ligand. AutoDock Vina 1.2.5 [7, 8] used these maps to evaluate ligand positions, orientations and permitted bond rotations, retaining the pose with the best AD4 score from each search.

PolyP10 limited the conformational search while retaining a flexible chain of repeated phosphate groups. Increasing chain length adds rotatable bonds and greatly expands the possible conformations. AD4 also penalizes the loss of torsional freedom upon binding, so favorable noncovalent interactions must compensate for that penalty. These considerations motivated the ten phosphate ligand; the main Methods give the search settings and scoring terms.

##### Defining completed and high scoring results

A completed docking result required finite AD4 score, E, PRI and PDI100 values. This definition retained 19,481 results from 5,648 AlphaFold models and 2,925 UniProt protein entries, among 23,383 positive patch records. One record with a numerical PDI100 but no AD4 score was excluded. Data S1 retains all patch records, including the 3,902 incomplete outputs. Missing output was not classified as nonbinding.

The PDI100 ≥80 cutoff selected 185 high scoring docking results from 128 AlphaFold models and 82 UniProt protein entries. No value equaled exactly 80. The normal/conditional cell surface exposure group contained 74 results from 56 models and 49 entries; the other 111 results represented 72 models and 38 entries. Five entries had models in both groups. Cell surface exposure described the represented protein or isoform, without establishing access to its saved polyP10 contact site. All 185 results remained eligible for selection consistency and functional site analyses.

Soluble extracellular evidence was kept separate from cell surface exposure. For example, reported MINPP1 activity in human plasma supported a soluble pool, not a stable cell surface ectodomain, so that annotation did not place canonical MINPP1 in the normal/conditional cell surface exposure group. TAAR3P was retained with uncertain protein existence because the human locus is annotated as a pseudogene. Data S2 preserves these statuses for each exact model and patch.

The normalization references remained fixed during all PDI comparisons. Docking favorability D used 19,481 values, with median 14.88 and interquartile range (IQR) 4.90 kcal/mol. Electrostatic intensity E used all 23,383 positive patches, with median 3,272.4149 and IQR 561.3476. Recalculating PDI100 from the stored inputs reproduced the recorded scores to within $3.41 \times 10^{-13}$. E remained a relative electrostatic intensity measure, not physical charge density.

##### Comparing scores across isoform models

The isoform comparison tested whether canonical and alternative models gave similar score patterns. For each AlphaFold model, we averaged D, E, PRI, Base and PDI100 over its completed docking results. Canonical models followed the identifier pattern AF–[UniProt main accession]–F1; named alternative isoform models shared the main accession. Within each UniProt protein entry, the canonical model mean was paired with the median of the alternative model means. This produced 1,175 pairs from 1,175 canonical and 2,443 alternative models, without assuming matching patches across isoforms (Data S5).

Spearman correlations measured the association between the paired score summaries. We resampled the 1,175 pairs with replacement 2,000 times, using the same resampled indices for all five scores. The 2.5th and 97.5th percentiles gave the 95% bootstrap intervals in Supplementary Table S1. This comparison assessed consistency among related computational models.

#### Isolating PRI's effect on selection

Two comparisons tested PRI at a fixed cutoff and at a fixed selection size. For the cutoff comparison, removing only the PRI multiplier left Base ×100; we counted results at or above 80 before and after applying PRI. For the fixed size comparison, we ranked all 19,481 completed results separately by AD4 score, E, Base and PDI100, then selected the top 185 from each ranking. The source scores are in Supplementary Data S1. AD4 scores were ordered from most negative upward; the other scores were ordered from largest downward.

Within each selected set, we counted results with PRI $<0.75$ and results shared with the PDI100 selection. Base versus PDI100 isolated the PRI penalty. AD4 or E alone versus PDI100 compared a single component with the complete index. Main Figure 4a,b shows the cutoff comparison and low PRI counts in the fixed size selections.

#### Tracking selection across 135 PDI settings

The sensitivity analysis tested dependence on PDI settings chosen manually rather than fitted to experimental polyP binding measurements. Main Figure 4f gives the parameter values used to generate the 135 combinations of docking weight, sigmoid scale, PRI reference and penalty strength. Their original values were $w_D = 0.6$, $s = 1$, PRI reference = 0.75 and $\lambda = 3$; the electrostatic weight was $1 - w_D$. Each combination reranked all 19,481 completed results using the same normalization references, raw D, E and PRI values, model identities and saved poses. The same sigmoid scale was applied to D and E within each combination.

Each combination selected its top 185 results by score; ties retained the source row order. The numerical PDI100 ≥80 cutoff was not reapplied after changing the settings. We tracked each of the original 185 high scoring results by its exact model and patch identifier, so a different patch or isoform could not substitute for it. A consistently selected docking result remained in the top 185 under all 135 tested combinations.

Data S2 gives each original result's selection count and minimum, median and maximum ranks. Data S3 gives the settings and the number of original results retained by each combination, including counts by cell surface exposure group. Its above80_count field is a separate numerical cutoff count, not the selection rule. The analysis followed the original candidates; newly high ranking results did not enlarge that group.

#### Testing enrichment with sampling controls

Functional enrichment was assessed at the UniProt protein entry level, counting each main accession once. We compared the 82 entries with high scoring docking results against the other 2,843 entries with completed docking. Two sided Fisher exact tests covered 219 annotations with at least ten annotated and ten unannotated entries in the 2,925 entry reference set. Fold enrichment divided the annotation frequency among the 82 selected entries by its frequency among all 2,925 reference entries. Benjamini–Hochberg adjustment [9] was applied across the 219 P values; the data files label these adjusted values q.

Two additional analyses tested whether enrichment persisted after accounting for unequal sampling. Longer proteins, additional isoform models and more completed searches can increase the opportunity to obtain a high scoring result. The canonical model analysis compared 64 selected entries with 2,533 nonselected entries among the 2,597 entries with completed canonical model docking. The length and search count analysis retained all 2,925 entries. Both used the original UniProt keywords and sequence lengths, without refreshing annotations between comparisons.

For the length and search count analysis, entries were divided into three sequence length groups (≤372, 373–675 and ≥676 amino acids) and five completed search groups (1, 2–3, 4–7, 8–15 and ≥16), giving 15 groups. Within each group, a hypergeometric distribution described the annotation count expected among selected entries under random selection without replacement. Combining these distributions gave one two sided test per annotation. Each test summed probabilities at least as far from the expected count as the observed count; Benjamini–Hochberg adjustment again covered all 219 annotations. Data S4 retains every tested annotation, including nonsignificant results.

Comparisons within the normal/conditional cell surface exposure group and among consistently selected results were descriptive. They used the same source keywords and exact model and patch memberships. For main Fig. 4k, oxidoreductases, translocases and other entries formed nonoverlapping groups of 8, 9 and 65 among the 82 selected entries, and 6, 8 and 35 among the 49 exposure group entries. No additional enrichment tests were applied to these subsets (Data S2 and S8).

#### Separating patch size from composition

Patch analyses tested whether simpler surface properties explained docking favorability. Spearman correlations compared D with positive patch area and with the count or percentage of histidine, arginine and lysine residues. To account for patch size, we regressed the ranks of D and of each composition measure on ranked patch area, then correlated the residuals. For within model comparisons, we subtracted each AlphaFold model’s mean rank before correlating the remaining variation.

Intervals came from 2,000 bootstrap resamples of UniProt protein entries, keeping all associated patch results together. These exploratory intervals were not adjusted for multiple testing. Separately, we divided each score’s total sum of squares into variation between model means and variation among results within each model. This described where score variation occurred, without assigning a biological cause. Numerical estimates are supplied in the Figure S1 source tables listed in the file index.

#### Mapping experimental sites to saved poses

Residue contacts were calculated for all 185 selected polyP10 poses using the original ligand PDBQT and matching AlphaFold PDB coordinates (Data S6; Supplementary_Structures.zip). For each protein residue, we calculated its shortest distance to polyP10 between nonhydrogen atoms. An unrounded distance ≤4.000 Å defined contact. The calculation used the saved coordinates without redocking or structural alignment. Residue pLDDT came from the AlphaFold PDB B-factor field and contributed once per residue to each reported mean.

Experimental residue sets were assembled from UniProt annotations [10], primary studies and deposited structures, then mapped to the exact model sequences. The review distinguished catalytic residues, structural ligand contacts, substitutions affecting binding and substitutions affecting other functions. Residues changed together in a combined mutant were treated as one experimental set, not as independently tested determinants. Neutral substitutions were kept separate from substitutions with a reported effect. Fractions used all mapped residues in the specified set as their denominator, including noncontacts.

Usable experimental sets were available for 113 poses from 74 AlphaFold models and 44 UniProt protein entries. A missing residue set was distinguished from a mapped set with no overlap. Data S10 contains the 74 poses in the normal/conditional cell surface exposure group, and Data S12 contains the other 111 poses; together they cover the full 185 pose population. The datasets preserve species, constructs, residue identities, distances and nonoverlaps. Structural states, overlapping sets and subsets were not counted as independent experiments. The overlap fractions describe the specified comparisons; they were not subjected to a spatial enrichment test.

Sequence mapping used each study's species, construct and numbering. Mature LPL residues mapped to precursor positions by +27; CCR9 isoform 2 required −12. The reviewed SLC12A3 construct lacked canonical Gln95, requiring +1 at the ATP site positions. RXFP2's +36 correspondence was established by sequence matching, while seven SSTR2 isoform 2 tail contacts had no canonical tail counterpart. Data S10 records these mappings and unmapped positions, rather than applying a universal offset.

For MCOLN3, the marmoset sequence aligned without gaps to the human model, with 538 of 553 positions identical. All four experimentally implicated binding positions retained their residue identities and numbers. Human MCOLN1 structural and functional sets were kept separate from the mapped mouse contact sets from closed and open states [11, 12, 13].

For DUOX1, we calculated the complete NADPH contact sets from structures 7D3E and 7D3F using the same ≤4 Å nonhydrogen atom criterion. The sets contained 21 residues at low calcium and 20 at high calcium. The two DUOX1 copies within each structure gave identical sets and were counted once. Experimental numbering matched the precursor model; the construct's L1178F substitution lay outside these sets. The three phosphate recognition arginines highlighted in the structural study formed a separate subset of the complete pocket [14].

Main Figure 5 was rendered in PyMOL 3.1.0 using the saved AlphaFold models and polyP10 poses. Opaque solvent excluded surfaces were generated from each protein with a 1.4 Å probe. Surface colors came from the existing APBS map for the protein alone, sampled one probe radius (1.4 Å) above the molecular surface. All surface views used the same linear color scale: red at −6, white at 0 and blue at +6 kBT/e, with endpoint colors retained outside this range. Corresponding cartoon and surface views used the same orientation and framing; the saved coordinates were unchanged (Supplementary_Structures.zip).

##### Resolving LPL contacts along the phosphate chain

The detailed LPL comparison used AF-P06858-F1-model_v6, patch001. Experimental residues were mapped from mature LPL to the 475 residue precursor by adding the 27 residue signal peptide and checking amino acid identity. The heparin affinity set comprised five individually identified determinants from Hata et al. [15] and two from Kobayashi et al. [16]. The latter were mainly tested in an isolated C-terminal construct. Combined mutants and individually tested residues without an affinity effect remained separate in Supplementary Table S6. Distances to catalytic Ser159, Asp183 and His268 were also measured [17].

The polyP10 ligand contained ten phosphorus and 31 oxygen atoms. Phosphorus–oxygen neighbors within 1.9 Å defined ten connected phosphate groups and nine shared bridging oxygens. Phosphate numbering followed the chain from the end selected by atom serial order; it did not imply a biological direction. For main Fig. 6a, we measured the shortest distance from Lys NZ or Arg NE/NH1/NH2 to an oxygen in each phosphate group. A second calculation excluded shared bridging oxygens, and contact cutoffs of 3.5, 4 and 5 Å tested distance sensitivity.

Data S11 supplies these basic nitrogen–phosphate oxygen distances, phosphate assignments and local pLDDT values. Data S9 gives LPL and LIPI residue contact distances based on any residue nonhydrogen atom. The two datasets therefore describe different atom selections; main Fig. 6a uses Data S11. The saved source coordinates are in Supplementary_Structures.zip.

##### Combining evidence within each AlphaFold model

We joined selection consistency and functional site overlap by exact AlphaFold model identifier. A model met the selection criterion if at least one of its high scoring docking results remained selected under all 135 PDI settings. It met the overlap criterion if at least one of its high scoring saved poses contacted a mapped experimental functional site residue at ≤4 Å. Both properties had to occur in the same model, but could come from different poses. Canonical and alternative isoform models were counted separately. For the same pose intersection, both criteria had to be met by one saved docking result, identified by its AlphaFold model and positive patch.

Both comparisons used Data S2, S10 and S12 without adding a docking or scoring threshold. Of the 128 models, 34 had consistently selected results and 23 had functional site overlap; seven had both. Experimental sets were available for 74 models. The other 54 lacked a usable mapped set, while 51 had mapped sets without detected overlap. Data S13 retains all memberships and the exact poses supporting each property; Supplementary Table S5 lists the seven model intersection.

#### Supplementary Notes

##### Scores were consistent across isoform models

Canonical and alternative isoform models showed positive score associations for all five components (Supplementary Table S1). PRI had the highest correlation, followed by PDI100 and docking favorability D; electrostatic intensity E showed the weakest association. These comparisons used score summaries for each model, so they describe consistency across isoforms without requiring the same positive patches or docking poses.

Supplementary Table S1. Scores were consistent across canonical and alternative isoform models.

| Score compared | Spearman ρ | 95% bootstrap interval |
|---|---|---|
| Docking favorability D = −AD4 score | 0.760 | 0.726–0.791 |
| Electrostatic intensity E | 0.559 | 0.513–0.602 |
| pLDDT Reliability Index (PRI) | 0.822 | 0.795–0.846 |
| Base score | 0.739 | 0.705–0.771 |
| Patch Docking Index (PDI100) | 0.790 | 0.758–0.817 |

For 1,175 UniProt protein entries, canonical model means were paired with medians of alternative isoform model means. Each model mean used its completed docking results. Intervals came from 2,000 paired bootstrap resamples (Data S5).

##### PRI prioritized confidently modeled regions

PRI changed both how many results exceeded the cutoff and which results were selected. At the fixed cutoff of 80, applying PRI reduced the selected count from 312 to 185, removing 127 results (40.7%). At a fixed selection size of 185, the number with PRI <0.75 fell from 86 under Base to 18 under PDI100, reducing their share from 46.5% to 9.7%. This second comparison isolated the penalty’s effect on membership rather than selection size (main Fig. 4a,b).

The single component rankings produced different selections. Among the top 185 results, AD4 alone selected 142 with PRI <0.75 and E alone selected 84. The AD4, E and Base selections shared 20, eight and 112 results, respectively, with PDI100. Median PRI increased from 0.694 under AD4 alone to 0.807 under PDI100. Together, these comparisons show how the integrated score prioritized favorable docking results from more confidently modeled regions.

##### Patch properties weakly tracked docking favorability

Basic residue content only weakly tracked docking favorability (Supplementary Fig. S1). Across 19,481 completed results, D correlated with His/Arg/Lys count at ρ = 0.203, their percentage at 0.058, and patch area at 0.261. Accounting for patch area reduced the count association to 0.021 (95% bootstrap interval, −0.0004 to 0.0422); the within model association was 0.088. Counting basic residues therefore did not reproduce the docking ranking. Source estimates are in the Figure S1 source tables (file index).

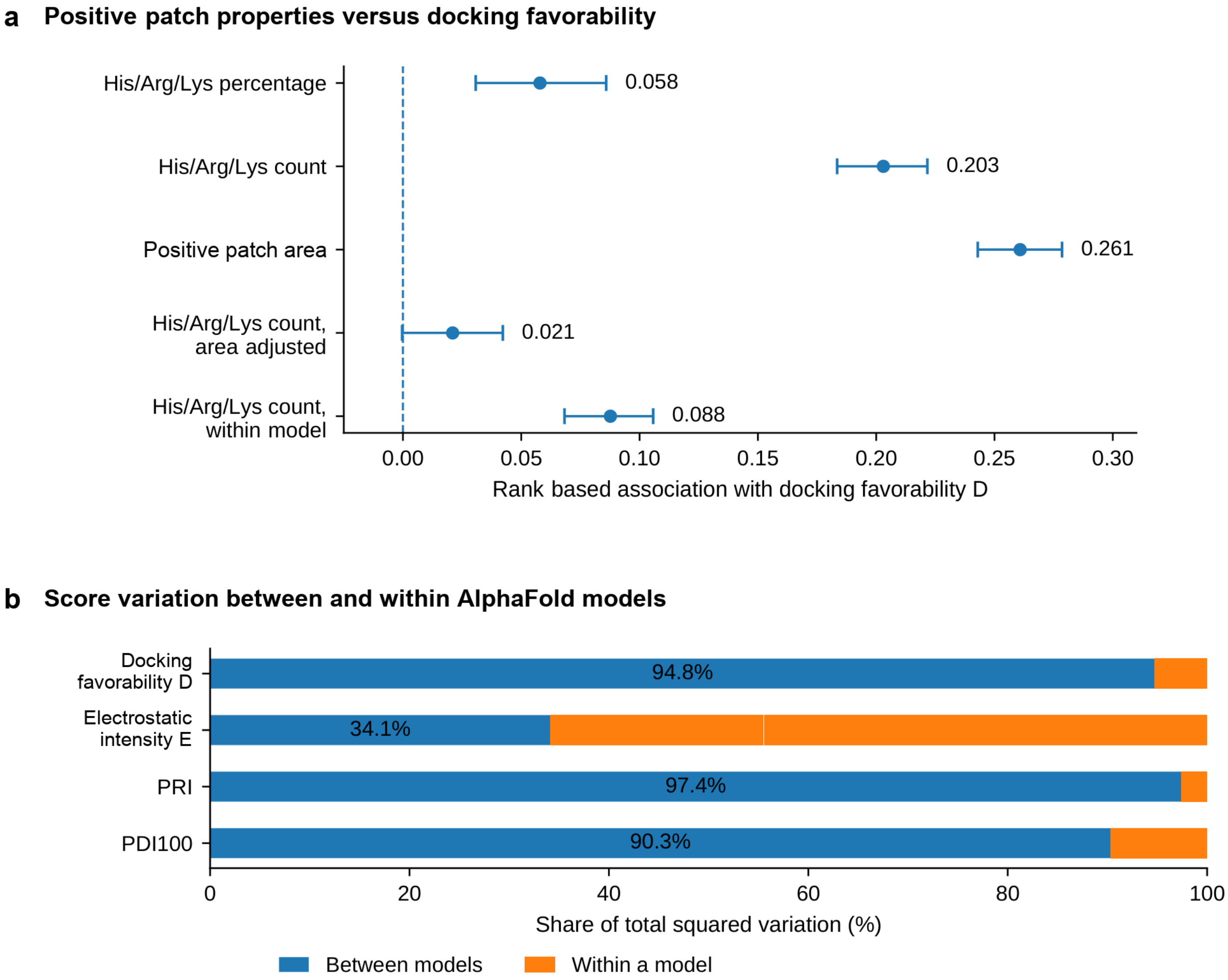

Supplementary Figure S1. Docking differed from positive patch properties. (a) Rank associations with docking favorability D. His/Arg/Lys denotes histidine, arginine and lysine in the original positive patch. Points show estimates; 95% intervals use 2,000 bootstrap resamples of UniProt protein entries and are not adjusted for multiple testing. Area adjusted estimates account for ranked patch area; within model estimates remove model mean ranks. (b) Percentage of each score's squared variation between model means; the remainder occurred within models. The analyses used 19,481 completed docking results. These descriptive comparisons concern patch properties and score variation, not binding accuracy.

E and D were also nearly uncorrelated (ρ = 0.00035), but they described different regions and quantities.

E summarized the original positive patch; AD4 evaluated the final polyP10 pose within the larger docking box using several interaction terms. In LIPI, the inspected pose contacted a surface outside the starting patch. The near zero association thus did not isolate the electrostatic contribution to binding. Most variation in D, PRI and PDI100 occurred between AlphaFold models, whereas E varied more among patches within a model (Supplementary Fig. S1).

#### Oxidoreductases and translocases remained enriched

Oxidoreductase and translocase enrichment persisted in the all model, canonical model and length and search count comparisons (Supplementary Table S2). This consistency showed that the associations were retained after accounting for the measured sampling differences. ATP binding and lipid transport annotations were overrepresented in the initial tests but did not remain significant after length and search count adjustment. Heparin binding and G protein coupled receptor annotations were more frequent among selected entries, but their initial Fisher tests were not significant after adjustment (P = 0.104 and 0.933). Data S4 includes all 219 annotations.

Supplementary Table S2. Oxidoreductase and translocase enrichment persisted across sampling controls.

| Functional annotation | Entries with completed docking n/2,925 | Entries with high scoring docking results n/82 | Fold enrichment | All model analysis adjusted P | Canonical model analysis adjusted P | Length and search count analysis adjusted P |
|---|---|---|---|---|---|---|
| Oxidoreductase | 34 | 8 | 8.39 | **0.000149** | **0.000449** | **0.000133** |
| Translocase | 48 | 9 | 6.69 | **0.000173** | **0.00435** | **0.0317** |
| ATP binding | 255 | 19 | 2.66 | **0.00152** | **0.0184** | 0.089 |
| Lipid transport | 69 | 9 | 4.65 | **0.00271** | **0.0184** | 0.0632 |
| Heparin binding | 15 | 3 | 7.13 | 0.104 | 0.0721 | 0.35 |
| G protein coupled receptor | 601 | 21 | 1.25 | 0.933 | 1 | 0.721 |

Two sided tests used Benjamini–Hochberg adjustment across 219 annotations in each analysis; bold values are <0.05. All model and canonical model tests compared 82 with 2,843 and 64 with 2,533 UniProt protein entries, respectively. Fold enrichment uses all 2,925 completed docking entries as the reference. The length and search count analysis compared entries within 15 groups. Full results are in Data S4.

FAD, NADP and flavoprotein annotations remained enriched in the length and search count analysis, with adjusted P values of 0.0000335, 0.0000335 and 0.0000434, respectively. These annotations occurred in six, seven and six selected entries, all within the eight oxidoreductases. They therefore added biochemical context to the oxidoreductase association rather than identifying three independent candidate classes.

#### Selection consistency varied within functional classes

Four of eight oxidoreductase entries and four of nine translocase entries contributed consistently selected docking results. Together they accounted for 15 of the 45 consistently selected results; the other 30 came from 16 of the remaining 65 selected entries. Supplementary Table S3 distinguishes each entry’s highest scoring result from all its consistently selected results. This distinction matters when the best original score and the most consistently selected pose differ.

Supplementary Table S3. Consistent selection within oxidoreductase and translocase candidates.

| Protein name (symbol) | Functional annotation | Highest PDI100 | Selection count for highest scoring result (out of 135) | Consistently selected / high scoring docking results | Normal/ conditional cell surface exposure group consistently selected / high scoring docking results | Cell surface exposure |
|---|---|---|---|---|---|---|
| NADPH oxidase 1 (NOX1) | Oxidoreductase | 91.96 | 135/135 | 4/4 | 2/2 | Yes |
| NADPH oxidase 2 (CYBB/NOX2) | Oxidoreductase | 80.72 | 81/135 | 0/1 | 0/1 | Yes |
| NADPH oxidase 3 (NOX3) | Oxidoreductase | 82.51 | 103/135 | 0/1 | 0/1 | Yes |
| NADPH oxidase 4 (NOX4) | Oxidoreductase | 84.59 | 135/135 | 2/2 | 1/1 | Conditional |
| Dual oxidase 1 (DUOX1) | Oxidoreductase | 86.90 | 106/135 | 2/3 | 2/2 | Review |
| Dual oxidase 2 (DUOX2) | Oxidoreductase | 87.80 | 135/135 | 1/2 | 1/2 | Yes |
| Nitric oxide synthase 3 (NOS3) | Oxidoreductase | 81.43 | 81/135 | 0/1 | 0/0 | No |
| NADPH oxidase 5 (NOX5) | Oxidoreductase | 83.64 | 104/135 | 0/1 | 0/0 | Review |
| ABC type organic anion transporter ABCA8 | Translocase | 81.15 | 73/135 | 0/1 | 0/1 | Yes |

| Protein name (symbol) | Functional annotation | Highest PDI100 | Selection count for highest scoring result (out of 135) | Consistently selected / high scoring docking results | Normal/ conditional cell surface exposure group consistently selected / high scoring docking results | Cell surface exposure |
|---|---|---|---|---|---|---|
| ATP binding cassette subfamily B member 6 (ABCB6) | Translocase | 84.02 | 135/135 | 1/1 | 1/1 | Yes |
| ATP binding cassette subfamily C member 2 (ABCC2) | Translocase | 88.09 | 135/135 | 2/6 | 2/6 | Yes |
| ATP binding cassette subfamily C member 11 (ABCC11) | Translocase | 81.93 | 69/135 | 0/2 | 0/2 | Yes |
| ATP binding cassette subfamily G member 4 (ABCG4) | Translocase | 82.41 | 108/135 | 0/2 | 0/2 | Yes |
| Phospholipid transporting ATPase IB (ATP8A2) | Translocase | 81.62 | 107/135 | 0/1 | 0/1 | Yes |
| Probable phospholipid transporting ATPase IIA (ATP9A) | Translocase | 82.30 | 108/135 | 0/1 | 0/1 | Conditional |
| Phospholipid transporting ATPase IG (ATP11C) | Translocase | 83.55 | 135/135 | 1/2 | 1/2 | Yes |
| Multidrug resistance associated protein 1 (ABCC1) | Translocase | 87.62 | 135/135 | 2/16 | 0/0 | Review |

The highest PDI100, its selection count and cell surface exposure status describe the same representative result. The two result count columns give consistently selected/high scoring docking results overall and within the normal/conditional cell surface exposure group. Consistently selected results remained in the top 185 under all 135 PDI settings; 0/0 means no results in the specified group (Data S8).

All six oxidoreductases in the normal/conditional cell surface exposure group belonged to the NOX/DUOX family and had canonical model results above 80. Across the NOX/DUOX family, the highest scoring results for NOX5 and DUOX1 came from alternative isoform models with unresolved cell surface exposure (Supplementary Table S4). DUOX1's highest scoring alternative model result was selected under 106 settings, while two canonical model results were selected under all 135. Scores and selection counts therefore need to remain attached to their exact model and patch.

Supplementary Table S4. Alternative isoforms changed the highest scores for NOX5 and DUOX1.

| Protein name | Highest canonical model PDI100 | Highest PDI100 across all models | Highest PDI100 in the normal/ conditional cell surface exposure group | Cell surface exposure of the highest scoring model | Selection count for highest scoring result (out of 135) |
|---|---|---|---|---|---|
| NADPH oxidase 1 (NOX1) | 91.96 | 91.96 | 91.96 | Yes | 135/135 |
| NADPH oxidase 2 (CYBB/NOX2) | 80.72 | 80.72 | 80.72 | Yes | 81/135 |
| NADPH oxidase 3 (NOX3) | 82.51 | 82.51 | 82.51 | Yes | 103/135 |
| NADPH oxidase 4 (NOX4) | 84.59 | 84.59 | 84.59 | Conditional | 135/135 |
| NADPH oxidase 5 (NOX5) | 73.55 | 83.64 | — | Review | 104/135 |
| Dual oxidase 1 (DUOX1) | 85.77 | 86.90 | 85.77 | Review | 106/135 |
| Dual oxidase 2 (DUOX2) | 87.80 | 87.80 | 87.80 | Yes | 135/135 |

Each score is the maximum within the stated model group; maxima may describe different results. The final column tracks the highest scoring result across all models. A dash means no high scoring result in the normal/conditional cell surface exposure group. The highest scoring NOX5 and DUOX1 results came from AF-Q96PH1-2-F1 and AF-Q9NRD9-2-F1, respectively, with unresolved cell surface exposure (Data S8).

Functional categories followed the source UniProt keywords. For example, ABCC8 carried an ATP binding annotation but was not assigned to the translocase category because that keyword was absent from the retrieval export. All eight translocase entries in the normal/conditional cell surface exposure group carried ATP binding annotations and seven carried lipid transport annotations. These shared keywords describe related functions and should not be read as separate groups of proteins (Supplementary Table S8).

#### Seven models combined selection consistency and site overlap

Seven AlphaFold models, representing five UniProt protein entries, combined at least one consistently selected docking result with at least one pose overlapping an experimental functional site (Supplementary Table S5). Six of these models contributed seven saved poses that were consistently selected and overlapped a functional site. These were canonical DUOX1 patch002 and patch008, canonical FFAR1 patch002, canonical MYO1G patch001, MINPP1 isoform 3 patch001, and MYO1C isoforms 2 and 3 patch010. Canonical MYO1C met the criteria through different poses: patch003 and patch013 were consistently selected, whereas patch001 and patch012 overlapped the membrane binding positions. Data S13 records membership and the supporting poses for all 128 models.

Supplementary Table S5. Seven AlphaFold models combined consistent selection and functional site overlap.

| Protein and AlphaFold model | Consistently selected docking results (patch IDs) | Docking results with functional site overlap (patch IDs) | Mapped experimental site |
|---|---|---|---|
| DUOX1 (canonical)<br>AF-Q9NRD9-F1 | patch002, patch008 | patch002, patch008 | NADPH recognition pocket |
| FFAR1 (canonical)<br>AF-O14842-F1 | patch002 | patch001, patch002 | Receptor activation and agonist response |
| MINPP1 (isoform 3)<br>AF-Q9UNW1-3-F1 | patch001 | patch001 | Mapped catalytic histidine; truncated isoform |
| MYO1C (isoform 2)<br>AF-O00159-2-F1 | patch003, patch010, patch017 | patch001, patch010 | Phosphoinositide dependent membrane attachment |
| MYO1C (isoform 3)<br>AF-O00159-3-F1 | patch004, patch010 | patch001, patch010 | Phosphoinositide dependent membrane attachment |
| MYO1C (canonical)<br>AF-O00159-F1 | patch003, patch013 | patch001, patch012 | Phosphoinositide dependent membrane attachment |
| MYO1G (canonical)<br>AF-B0I1T2-F1 | patch001 | patch001 | Membrane association |

Models are AlphaFold version 6; append "-model_v6" to each identifier. Consistent selection means top 185 membership under all 135 PDI settings. Functional site overlap uses the 4 Å criterion and the experimental sets in main Table 2. The two properties may involve different poses within one model. MINPP1 isoform 3 is truncated relative to the experimental protein.

#### PolyP poses overlapped functional sites

The site comparisons linked predicted contacts to experimentally studied protein functions. 31 poses overlapped a mapped residue set, spanning 23 AlphaFold models and 18 UniProt protein entries. Main Table 2 gives the residue level summaries. The notes below explain the experimental endpoints and the distinctions needed to interpret them; Data S10 and S12 retain full sets, nonoverlaps and sequence maps.

##### DUOX1 poses overlapped the NADPH pocket

Both consistently selected canonical DUOX1 poses, patch002 and patch008, overlapped the NADPH pocket. Each contacted Arg1036, Arg1424 and Arg1495, which recognize the ribose phosphate of NADPH in the experimental structure [14]. NADPH supplies electrons for oxidase activity.

Against the complete pocket, both poses contacted 9/21 residues in the low calcium structure 7D3E. In the high calcium structure 7D3F, patch002 contacted 7/20 and patch008 contacted 8/20. The isoform 2 patch002 pose likewise contacted 9/21 and 7/20 mapped residues, including all three phosphate recognition arginines in its own sequence. PolyP binding at this surface could affect NADPH recognition or access.

##### NOX2 contacts included activation and heme sites

The NOX2 patch002 pose overlapped two distinct functional features. It contacted all three D-loop positions at which individual substitutions impaired oxidase activation—Lys195, Arg198 and Arg199—but neither of the two C-terminal positions tested in the same study [18]. It also contacted His101 and His209, two of the four histidines coordinating the enzyme's heme groups in the experimental structure [19]. Heme coordination supports electron transfer, whereas the D-loop evidence concerns activation.

##### MCOLN contacts overlapped phosphoinositide recognition

The MCOLN1 and MCOLN3 poses contacted sites involved in recognizing PI(3,5)$P_2$, a phosphoinositide that regulates these lysosomal ion channels. The canonical MCOLN1 patch001 pose contacted all six lipid contact residues assigned in the human structure [12]. It also contacted Tyr355, one of two positions tested separately for channel activation, but not Arg403. Mapped mouse structures gave overlaps of 6/6 closed state contacts and 6/7 open state contacts [13]. Structural contact and activation evidence were therefore retained as separate sets.

The MCOLN3 patch001 pose contacted all four positions where individual substitutions reduced binding signals for a short chain PI(3,5)$P_2$ analog [11]. Binding was measured by isothermal titration calorimetry, which detects heat changes during binding. The experiments used marmoset MCOLN3 carrying N138Q, a substitution introduced to remove a potential sugar attachment site; all four compared residue identities and numbers matched the human model. Heat signals from the four binding site mutants were too small to estimate affinities reliably, so the evidence identifies reduced binding signals rather than quantified affinity losses. Together, these comparisons place the predicted polyP interfaces at phosphoinositide recognition regions relevant to channel regulation.

##### GLUT1 contacts overlapped a phosphorylation site

The GLUT1 patch002 pose contacted Ser226 and Arg223, linking the predicted interface to regulated glucose uptake. Ser226 is the protein kinase C (PKC) phosphorylation site required for the stimulated uptake response; substitutions at Arg223 impaired phosphorylation and that response [20]. Both residues had pLDDT 96.38 in the AlphaFold model. PolyP association at these sites could affect PKC dependent regulation of glucose uptake.

##### ARRB2 contacts overlapped receptor recruitment sites

The ARRB2 patch002 pose overlapped phosphoinositide recognition regions involved in recruiting β-arrestin 2 to activated receptors. It contacted all three positions in the combined mutant studied by Janetzko et al. [21] and both positions in the combined mutant studied by Jung et al. [22]. Those experiments measured impaired recruitment or receptor internalization. The pose also contacted two of four positions in a further combined mutant, Lys230 and Arg332; Arg332 had additional individual substitution evidence for reduced recruitment [23]. The combined mutants identify functional regions without assigning an independent effect to every residue. PolyP association there could affect phosphoinositide dependent receptor regulation.

##### CYTH contacts overlapped membrane recruitment sites

CYTH1 and CYTH2 activate ARF proteins, molecular switches involved in membrane trafficking. Phosphoinositide binding recruits these regulators to membranes. The CYTH1 isoform 2 patch001 and isoform 3 patch004 poses each contacted the position corresponding to Arg281 in the experimental protein [24]. Substitution at this position impaired PI(3,4,5)$P_3$ binding and membrane association. Sequence mapping placed it at Arg280 in isoform 2 and Arg222 in isoform 3.

In the mouse CYTH2 pleckstrin homology domain, individual substitutions at seven of 12 tested positions markedly reduced binding to soluble inositol phosphates representing phosphoinositide headgroups [25]. Both human CYTH2 poses contacted all seven corresponding positions. They also contacted Thr276, whose phosphorylation by Akt contributes to CYTH2 dependent activation of ARF6 [26]. Wild type CYTH2 increased ARF6 activation, whereas T276A did not produce a significant increase. Lipid recognition and phosphorylation dependent regulation thus provide separate functional contexts for the same predicted interfaces.

##### Myosin contacts overlapped membrane attachment regions

MYO1C poses overlapped a membrane binding region in the motor protein's tail. Individual substitutions at two basic positions in mouse MYO1C impaired PI(4,5)$P_2$ dependent membrane binding and cellular localization [27]. Their human counterparts were Lys927/Arg938 in the canonical model, Lys892/Arg903 in isoform 2, and Lys908/Arg919 in isoform 3. At least one pose in each model contacted both positions. Main Table 2 distinguishes these full overlaps from poses contacting one position; the experimental comparison concerned the same two site set.

The MYO1G patch001 pose contacted three of five positions individually implicated in membrane association: Lys877, Arg887 and Lys898 [28]. That study established the region's contribution to membrane attachment without assigning lipid specificity to each position. These mapped experiments linked both interfaces to membrane attachment and specifically tested PI(4,5)$P_2$ dependent binding for MYO1C.

##### Receptor contacts overlapped intracellular regulatory sites

The CCR9 isoform 2 patch001 pose contacted four of 17 positions in the intracellular pocket for vercirnon, an antagonist that disrupts productive G protein coupling [29]. One of those contacts, Thr69, also belonged to a separate five position set tested for antagonist binding. The structural pocket and mutation dependent affinity evidence therefore had different denominators. Data S10 retains the construct specific numbering and the engineered A255/native V255 mismatch.

The CCR5 patch001 pose contacted Arg126 in the DRY motif, where substitution affected G protein activation [30]. The GPR15 patch001 pose contacted Lys231 at the G protein interface, where substitution reduced G protein activation [31]. Lys231 had pLDDT 56.03, indicating low local confidence in this part of the AlphaFold model. These comparisons concerned receptor–G protein regulation, distinct from the CCR5 and GLUT1 aptamer recognition evidence in Supplementary Table S9.

Both FFAR1 poses contacted Arg104, where the R104P substitution impaired signaling and cell surface localization [32]. Patch001 also contacted Gly103; substitution at this position changed basal activity and responses to some agonists in a separate study [33]. These endpoints link the predicted interfaces to receptor activation and trafficking.

##### NLRP6 contacts overlapped an RNA dependent assembly region

Three canonical NLRP6 poses and one isoform 2 pose contacted Lys352 and Lys356, two of four lysines tested together in a mutant that impaired assembly dependent on double stranded RNA [34]. The experimental study linked this basic region to phase separation and inflammasome activation. Because the four positions were changed together, the overlap identifies an assembly related region rather than two individually established RNA contacts. PolyP association with this region could affect RNA dependent assembly.

##### PolyP contacted His89 in MINPP1 isoform 3

The MINPP1 isoform 3 patch001 pose contacted His89, the mapped counterpart of a catalytic histidine whose substitution impaired phosphatase activity with 2,3-bisphosphoglycerate [35]. His89 had pLDDT 97.94. The contact nominates a catalytic site interaction, but the model represents a 284 residue isoform lacking a substantial C-terminal segment of the experimental full length protein. Its functional interpretation therefore depends on whether the truncated form retains the relevant activity.

##### TBC1D24 contacts overlapped stress protection positions

The TBC1D24 patch003 pose contacted Arg40 and Arg242, two of four positions where disease associated substitutions reduced the protein's protection against oxidative stress in neuronal cells [36]. The measured endpoint was stress protection, not ligand binding, so these positions were treated as functional evidence without assigning a ligand recognition pocket.

##### Contact cutoff checks preserved several site overlaps

Distance sensitivity was tested on the same saved coordinates at 3.5 and 5 Å. The LPL heparin affinity overlap remained 5/7 at both cutoffs; MCOLN1 structural lipid contacts remained 6/6, MCOLN3 binding positions 4/4, DUOX1 phosphate recognition arginines 3/3, NOX2 D-loop positions 3/3, and GLUT1 regulatory positions 2/2. These results show which specific overlaps were insensitive to the tested cutoff changes (Data S10).

Other contacts depended on the cutoff. For each canonical DUOX1 pose, overlap with the full 21 residue low calcium NADPH set increased from 7/21 at 3.5 Å to 9/21 at 4 Å and 10/21 at 5 Å. NOX2 heme site overlap was 0/4 at 3.5 Å and 2/4 at 4 and 5 Å. Reporting these changes separately preserves the distinction between broad pocket proximity and the closer contacts retained across cutoffs.

#### LPL contacts supported shared polymer recognition

##### LPL contacts overlapped heparin affinity determinants

LPL enabled a residue level comparison of predicted polyP contacts with heparin affinity determinants. PolyP and heparin differ chemically but both present repeated negative charges to protein surfaces. Ferrucci et al. proposed that polyP binds at an RNA binding region of SARS-CoV-2 polymerase [37]. This proposal prompted comparison with LPL's heparin binding site. Individual LPL substitutions had identified heparin affinity determinants. The LPL patch001 result scored PDI100 80.63 and was selected under 79 of the 135 PDI settings.

The pose contacted five of seven individually identified heparin affinity determinants. Hata et al.'s five position set contributed three contacts, precursor Arg306, Lys307 and Arg309 [15]. Kobayashi et al.'s two position set contributed Arg432 and Lys434, corresponding to mature Arg405 and Lys407 [16]. The latter study mainly tested these positions in a C-terminal construct, with Arg405 also examined in full length LPL. Main Figure 6b shows all seven distances. Supplementary Table S6 keeps combined substitutions and individual substitutions without affinity effects outside this seven residue denominator.

Supplementary Table S6. Additional LPL substitutions were separate from the seven affinity determinants.

| Mature chain → AlphaFold model residue | Distance to polyP10 (Å) | Experimental evidence | Inclusion in the seven residue comparison |
|---|---|---|---|
| Lys413 → Lys440; Lys414 → Lys441 | 7.10; 2.74 | The K413A/K414A double mutant reduced affinity in the C-terminal construct (Kobayashi). | A paired mutation does not identify two independently positive residues; neither enters the 5/7 set. |
| Lys381, Lys422, Lys428, Lys430 (mature numbering) | — | Individual substitutions did not alter affinity in the C-terminal construct (Kobayashi). | These tested residues are not part of the denominator of positive residues. |

Residue numbering is given for mature LPL and the AlphaFold precursor. Distances use the saved pose (Data S9); a dash means no distance was tabulated here.

Basic contacts extended along the polyP10 chain. Eight of the eleven contacting LPL residues were Lys/Arg residues, and every phosphate had an oxygen within 4 Å of a basic side chain nitrogen. All ten phosphates retained such contacts when shared bridging oxygens were excluded; nine retained them at 3.5 Å. The eight basic residues had mean pLDDT 97.12 (range 94.56–98.50). Main Figure 6a and Data S11 show this distributed contact arrangement.

The pose lay away from catalytic Ser159, Asp183 and His268, at minimum distances of 29.04, 24.74 and 28.62 Å. Experimental substitutions at these catalytic residues abolished activity without changing heparin affinity [17]. The predicted interface therefore aligned with heparin recognition rather than the catalytic triad, supporting a specific hypothesis of shared surface binding and possible competition between polyP and heparin.

##### LIPI contacts spanned the phosphate chain

LIPI provided a complementary case: recombinant LIPI showed heparin column retention [38], and its canonical and isoform 2 patch002 docking results remained selected under all 135 PDI settings, with PDI100 values of 85.91 and 86.82. The inspected canonical pose contacted eleven residues, including six Lys/Arg residues whose basic nitrogens contacted all ten phosphates at 4 Å. The contacting residues had mean pLDDT 97.84, with all above 96 (Data S9 and S11).

The LIPI pose was separated from the proposed catalytic triad: minimum distances were 11.01 Å to Ser159, 13.94 Å to Asp183 and 14.86 Å to His253. Ser159 had mutational evidence for catalytic activity, whereas Asp183 and His253 were assigned by sequence similarity [38]. Heparin column retention established whole protein recognition for LIPI without mapping the binding residues. Its consistently selected docking results therefore supplied different evidence from LPL’s overlap with individually identified heparin affinity determinants.

#### Complete high scoring candidate catalog

Supplementary Table S7 lists all 82 UniProt protein entries with high scoring docking results. Each row shows the result with the highest original PDI100 for that entry; its selection count and cell surface exposure status refer to that same result. Counts of results and models include all high scoring records for the entry. Data S2 retains all 185 exact model and patch records, including additional isoforms and poses.

Supplementary Table S7. All 82 UniProt protein entries with high scoring docking results.

| Gene | UniProt accession | High scoring docking results / AlphaFold models | Normal/conditional cell surface exposure group results / models | Highest PDI100 | Selection count for highest scoring result (out of 135) | Cell surface exposure |
|---|---|---|---|---|---|---|
| GPR26 | Q8NDV2 | 1 / 1 | 1 / 1 | 93.35 | 135/135 | Yes |
| CYTH4 | Q9UIA0 | 2 / 1 | 0 / 0 | 92.84 | 135/135 | No |

| Gene | UniProt accession | High scoring docking results / AlphaFold models | Normal/conditional cell surface exposure group results / models | Highest PDI100 | Selection count for highest scoring result (out of 135) | Cell surface exposure |
|---|---|---|---|---|---|---|
| MYO1C | O00159 | 13 / 3 | 0 / 0 | 92.76 | 123/135 | No |
| MYO1G | B0I1T2 | 1 / 1 | 0 / 0 | 92.18 | 135/135 | No |
| NOX1 | Q9Y5S8 | 4 / 2 | 2 / 1 | 91.96 | 135/135 | Yes |
| PDPK1 | O15530 | 3 / 1 | 0 / 0 | 91.80 | 128/135 | No |
| MINPP1 | Q9UNW1 | 2 / 2 | 0 / 0 | 91.47 | 135/135 | Review |
| TAAR3P | Q9P1P4 | 1 / 1 | 0 / 0 | 90.31 | 135/135 | Uncertain† |
| LIPI | Q6XZB0 | 7 / 5 | 2 / 2 | 89.10 | 135/135 | Review |
| GPR183 | P32249 | 1 / 1 | 1 / 1 | 88.23 | 135/135 | Yes |
| GNA12 | Q03113 | 1 / 1 | 0 / 0 | 88.18 | 135/135 | No |
| ABCC2 | Q92887 | 6 / 1 | 6 / 1 | 88.09 | 135/135 | Yes |
| DUOX2 | Q9NRD8 | 2 / 1 | 2 / 1 | 87.80 | 135/135 | Yes |
| FFAR1 | O14842 | 2 / 1 | 2 / 1 | 87.80 | 135/135 | Yes |
| ABCC1 | P33527 | 16 / 5 | 0 / 0 | 87.62 | 135/135 | Review |
| NOD2 | Q9HC29 | 2 / 2 | 0 / 0 | 87.10 | 135/135 | No |
| RGS6 | P49758 | 16 / 15 | 0 / 0 | 87.06 | 135/135 | No |
| DUOX1 | Q9NRD9 | 3 / 2 | 2 / 1 | 86.90 | 106/135 | Review |
| SLC12A4 | Q9UP95 | 6 / 5 | 6 / 5 | 86.31 | 127/135 | Yes |
| GPR87 | Q9BY21 | 1 / 1 | 1 / 1 | 86.25 | 101/135 | Yes |
| SSTR2 | P30874 | 1 / 1 | 1 / 1 | 86.04 | 101/135 | Yes |
| OPN5 | Q6U736 | 3 / 1 | 3 / 1 | 85.87 | 135/135 | Yes |
| SLC16A7 | O60669 | 2 / 1 | 2 / 1 | 85.85 | 135/135 | Yes |
| ABCC8 | Q09428 | 3 / 1 | 3 / 1 | 85.43 | 135/135 | Yes |
| SLC44A5 | Q8NCS7 | 2 / 1 | 2 / 1 | 85.20 | 108/135 | Yes |
| LTB4R | Q15722 | 1 / 1 | 1 / 1 | 85.18 | 104/135 | Yes |
| MCOLN1 | Q9GZU1 | 1 / 1 | 1 / 1 | 85.04 | 104/135 | Conditional |
| NLRP6 | P59044 | 4 / 2 | 0 / 0 | 84.61 | 99/135 | No |
| NOX4 | Q9NPH5 | 2 / 2 | 1 / 1 | 84.59 | 135/135 | Conditional |
| MYO6 | Q9UM54 | 1 / 1 | 0 / 0 | 84.56 | 97/135 | No |
| TBCD | Q9BTW9 | 4 / 2 | 0 / 0 | 84.48 | 135/135 | No |
| ABCB6 | Q9NP58 | 1 / 1 | 1 / 1 | 84.02 | 135/135 | Yes |
| ADCY10 | Q96PN6 | 5 / 4 | 0 / 0 | 83.99 | 127/135 | No |
| NOX5 | Q96PH1 | 1 / 1 | 0 / 0 | 83.64 | 104/135 | Review |
| P2RY13 | Q9BPV8 | 1 / 1 | 0 / 0 | 83.61 | 132/135 | Review |
| ATP11C | Q8NB49 | 2 / 1 | 2 / 1 | 83.55 | 135/135 | Yes |
| CLCN3 | P51790 | 3 / 1 | 0 / 0 | 83.48 | 96/135 | Review |
| CCR1 | P32246 | 1 / 1 | 1 / 1 | 83.46 | 100/135 | Yes |
| CYTH1 | Q15438 | 2 / 2 | 0 / 0 | 83.44 | 109/135 | No |
| PLPP3 | O14495 | 1 / 1 | 1 / 1 | 83.33 | 81/135 | Yes |
| SLC2A1 | P11166 | 1 / 1 | 1 / 1 | 83.20 | 113/135 | Yes |
| SLC12A5 | Q9H2X9 | 1 / 1 | 1 / 1 | 83.19 | 72/135 | Yes |
| MCOLN3 | Q8TDD5 | 1 / 1 | 1 / 1 | 83.17 | 114/135 | Conditional |
| PLEK2 | Q9NYT0 | 2 / 1 | 0 / 0 | 83.03 | 124/135 | No |
| PLD2 | O14939 | 2 / 2 | 0 / 0 | 83.01 | 112/135 | No |
| GPR15 | P49685 | 1 / 1 | 1 / 1 | 82.84 | 80/135 | Yes |
| LIPH | Q8WWY8 | 1 / 1 | 1 / 1 | 82.84 | 106/135 | Yes |
| TBC1D24 | Q9ULP9 | 1 / 1 | 0 / 0 | 82.80 | 126/135 | No |
| SLCO4C1 | Q8IVM8 | 1 / 1 | 1 / 1 | 82.78 | 108/135 | Yes |
| RASA3 | Q14644 | 2 / 2 | 0 / 0 | 82.73 | 121/135 | No |
| CCR3 | P51677 | 1 / 1 | 1 / 1 | 82.69 | 128/135 | Yes |

| Gene | UniProt accession | High scoring docking results / AlphaFold models | Normal/conditional cell surface exposure group results / models | Highest PDI100 | Selection count for highest scoring result (out of 135) | Cell surface exposure |
|---|---|---|---|---|---|---|
| NOX3 | Q9HBY0 | 1 / 1 | 1 / 1 | 82.51 | 103/135 | Yes |
| ABCG4 | Q9H172 | 2 / 2 | 2 / 2 | 82.41 | 108/135 | Yes |
| CCR5 | P51681 | 1 / 1 | 1 / 1 | 82.39 | 113/135 | Yes |
| SLC12A3 | P55017 | 3 / 2 | 1 / 1 | 82.32 | 64/135 | Yes |
| ATP9A | O75110 | 1 / 1 | 1 / 1 | 82.30 | 108/135 | Conditional |
| EXOC1 | Q9NV70 | 1 / 1 | 0 / 0 | 82.14 | 85/135 | No |
| RGS7 | P49802 | 3 / 3 | 0 / 0 | 82.06 | 105/135 | No |
| ABCC11 | Q96J66 | 2 / 1 | 2 / 1 | 81.93 | 69/135 | Yes |
| SLC12A1 | Q13621 | 2 / 2 | 2 / 2 | 81.87 | 85/135 | Yes |
| SLC44A1 | Q8WWI5 | 1 / 1 | 1 / 1 | 81.83 | 89/135 | Yes |
| OR4N4 | Q8N0Y3 | 1 / 1 | 0 / 0 | 81.71 | 112/135 | Review |
| ATP8A2 | Q9NTI2 | 1 / 1 | 1 / 1 | 81.62 | 107/135 | Yes |
| BDKRB1 | P46663 | 1 / 1 | 1 / 1 | 81.47 | 88/135 | Yes |
| OR4S1 | Q8NGB4 | 1 / 1 | 0 / 0 | 81.45 | 85/135 | Review |
| NOS3 | P29474 | 1 / 1 | 0 / 0 | 81.43 | 81/135 | No |
| RXFP2 | Q8WXD0 | 1 / 1 | 1 / 1 | 81.33 | 67/135 | Yes |
| GJB6 | O95452 | 1 / 1 | 1 / 1 | 81.32 | 56/135 | Conditional |
| ADTRP | Q96IZ2 | 2 / 1 | 0 / 0 | 81.31 | 67/135 | Review |
| ADGRA2 | Q96PE1 | 1 / 1 | 0 / 0 | 81.27 | 71/135 | Review |
| CYTH2 | Q99418 | 2 / 1 | 0 / 0 | 81.26 | 77/135 | No |
| PTAFR | P25105 | 1 / 1 | 1 / 1 | 81.19 | 82/135 | Yes |
| ABCA8 | O94911 | 1 / 1 | 1 / 1 | 81.15 | 73/135 | Yes |
| DMD | P11532 | 1 / 1 | 0 / 0 | 80.80 | 77/135 | No |
| CYBB | P04839 | 1 / 1 | 1 / 1 | 80.72 | 81/135 | Yes |
| ARRB2 | P32121 | 1 / 1 | 0 / 0 | 80.72 | 44/135 | No |
| LPL | P06858 | 1 / 1 | 1 / 1 | 80.63 | 79/135 | Yes |
| CXCR5 | P32302 | 1 / 1 | 1 / 1 | 80.61 | 81/135 | Yes |
| XKRX | Q6PP77 | 1 / 1 | 1 / 1 | 80.10 | 41/135 | Yes |
| DCSTAMP | Q9H295 | 1 / 1 | 1 / 1 | 80.06 | 69/135 | Yes |
| DCST1 | Q5T197 | 1 / 1 | 0 / 0 | 80.04 | 48/135 | Review |
| CCR9 | P51686 | 1 / 1 | 1 / 1 | 80.04 | 59/135 | Yes |

PDI100, selection count and cell surface exposure refer to the same highest scoring result. Selection count is top 185 membership across 135 PDI settings. The two result/model columns cover all high scoring records and those in the normal/conditional cell surface exposure group; 0/0 means none. Yes/Conditional indicate normal/conditional cell surface exposure; No indicates intracellular or inward facing localization; Review indicates unresolved exposure. Status can differ among isoforms and does not establish pose accessibility. † TAAR3P is a pseudogene with uncertain protein existence (Data S2).

Supplementary Table S8. Oxidoreductase and translocase candidates.

| Protein name (symbol; UniProt ID) | Highest scoring result AlphaFold model / positive patch | Cell surface exposure | Highest PDI100 | Selection count for highest scoring result (out of 135) | Consistently selected / high scoring docking results | Other functional annotations recorded in UniProt |
|---|---|---|---|---|---|---|
| **A. Oxidoreductases** | | | | | | |
| NADPH oxidase 1 (NOX1) Q9Y5S8 | AF-Q9Y5S8-F1 patch003 | Yes | **91.96** | **135/135** | **4/4** | FAD; NADP; Flavoprotein |
| NADPH oxidase 2 (CYBB/NOX2) P04839 | AF-P04839-F1 patch002 | Yes | 80.72 | 81/135 | 0/1 | FAD; NADP; Flavoprotein |

| **Protein name (symbol; UniProt ID)** | **Highest scoring result AlphaFold model / positive patch** | **Cell surface exposure** | **Highest PDI100** | **Selection count for highest scoring result (out of 135)** | **Consistently selected / high scoring docking results** | **Other functional annotations recorded in UniProt** |
|---|---|---|---|---|---|---|
| NADPH oxidase 3 (NOX3) Q9HBY0 | AF-Q9HBY0-F1 patch002 | Yes | 82.51 | 103/135 | 0/1 | None recorded |
| NADPH oxidase 4 (NOX4) Q9NPH5 | AF-Q9NPH5-F1 patch003 | Conditional | **84.59** | **135/135** | **2/2** | NADP |
| Dual oxidase 1 (DUOX1) Q9NRD9 | AF-Q9NRD9-2-F1 patch002 | Review | **86.90** | **106/135** | **2/3** | FAD; NADP; Flavoprotein |
| Dual oxidase 2 (DUOX2) Q9NRD8 | AF-Q9NRD8-F1 patch002 | Yes | **87.80** | **135/135** | **1/2** | FAD; NADP; Flavoprotein |
| Nitric oxide synthase 3 (NOS3) P29474 | AF-P29474-F1 patch002 | No | 81.43 | 81/135 | 0/1 | FAD; NADP; Flavoprotein |
| NADPH oxidase 5 (NOX5) Q96PH1 | AF-Q96PH1-2-F1 patch001 | Review | 83.64 | 104/135 | 0/1 | FAD; NADP; Flavoprotein |
| **B. Translocases** | | | | | | |
| ABC type organic anion transporter ABCA8 O94911 | AF-O94911-F1 patch009 | Yes | 81.15 | 73/135 | 0/1 | ATP binding; Lipid transport |
| ATP binding cassette subfamily B member 6 (ABCB6) Q9NP58 | AF-Q9NP58-F1 patch007 | Yes | **84.02** | **135/135** | **1/1** | ATP binding |
| ATP binding cassette subfamily C member 2 (ABCC2) Q92887 | AF-Q92887-F1 patch010 | Yes | **88.09** | **135/135** | **2/6** | ATP binding; Lipid transport |
| ATP binding cassette subfamily C member 11 (ABCC11) Q96J66 | AF-Q96J66-F1 patch004 | Yes | 81.93 | 69/135 | 0/2 | ATP binding; Lipid transport |
| ATP binding cassette subfamily G member 4 (ABCG4) Q9H172 | AF-Q9H172-F1 patch004 | Yes | 82.41 | 108/135 | 0/2 | ATP binding; Lipid transport |
| Phospholipid transporting ATPase IB (ATP8A2) Q9NTI2 | AF-Q9NTI2-1-F1 patch008 | Yes | 81.62 | 107/135 | 0/1 | ATP binding; Lipid transport |
| Probable phospholipid transporting ATPase IIA (ATP9A) O75110 | AF-O75110-F1 patch003 | Conditional | 82.30 | 108/135 | 0/1 | ATP binding; Lipid transport |
| Phospholipid transporting ATPase IG (ATP11C) Q8NB49 | AF-Q8NB49-F1 patch009 | Yes | **83.55** | **135/135** | **1/2** | ATP binding; Lipid transport |
| Multidrug resistance associated protein 1 (ABCC1) P33527 | AF-P33527-8-F1 patch008 | Review | **87.62** | **135/135** | **2/16** | ATP binding; Lipid transport |

Each row represents one UniProt protein entry with a PDI100 ≥80 result. The displayed model, patch, score, cell surface exposure and selection count describe its highest scoring result. “Consistently selected / high scoring docking results” counts consistently selected docking results among all high scoring docking results for that entry. Consistent selection means top 185 membership among 19,481 completed results under all 135 settings. Bold values identify entries with at least one consistently selected docking result, which may differ from the displayed result.
Models are AlphaFold version 6. YES/CONDITIONAL denote normal/conditional cell surface exposure; NO, intracellular or inward facing localization; REVIEW, unresolved exposure. Cell surface exposure labels apply to the displayed model and may differ across isoforms (Supplementary Table S4). “None recorded” means that none of the listed additional keywords appeared in the source export. Full records are in Data S2 and S8.

Supplementary Table S9. Heparin affinity and aptamer recognition in five candidate UniProt protein entries.

| Protein (UniProt ID) | Ligand | Experimental support | Primary study |
|---|---|---|---|
| LPL P06858 | Heparin | Mutagenesis and heparin affinity chromatography identified individual affinity determinants. | Hata et al. [15]; Kobayashi et al. [16] |
| LIPI Q6XZB0 | Heparin | Recombinant human mPA-PLA1β from conditioned medium was retained on a heparin column. | Hiramatsu et al. [38] |
| LIPH Q8WWY8 | Heparin | Recombinant human mPA-PLA1α from conditioned medium was retained on a heparin column. | Hiramatsu et al. [38] |
| CCR5 P51681 | RNA aptamer G-3 | Cell binding assays with controls lacking the receptor and CCR5 knockdown supported recognition dependent on the receptor. | Zhou et al. [39] |
| GLUT1 (SLC2A1) P11166 | DNA aptamer A5 | Selection using isogenic cells and reduced binding in cells with GLUT1 knocked out supported recognition dependent on the target. | Rosch et al. [40] |

The LIPI heparin affinity study did not separately test the isoform represented by the isoform 2 model. Aptamer evidence concerns target dependent cell binding, whereas the lipase studies measured heparin affinity.
