## Supplementary Code for "A structural screen for candidate polyphosphate binding surfaces in human membrane associated proteins": reconstruct_lpl_lipi_contacts.py

```
#!/usr/bin/env python3
"""Reconstruct LPL/LIPI contact tables from the original saved coordinates.

Python 3 standard library only. No docking, alignment, coordinate changes or
external services are used. Distances and cutoff tests use unrounded values.
Run: python3 reconstruct_lpl_lipi_contacts.py --coordinates PATH --output PATH
PATH may contain the four original coordinate files in nested subdirectories.
"""

import argparse
import csv
import hashlib
import json
import math
from collections import defaultdict
from pathlib import Path


SOURCES = {
    "LPL": {
        "model": "AF-P06858-F1-model_v6", "patch": "patch001",
        "model_sha256": "bfebd6e791bede2789ae642fc7b457acc281ed9d6130d90372387e7dbb670014",
        "pose_sha256": "3cdae882c86100bc4b1de6e83417e2e690453670b4196de75a474cdd84bddccb",
    },
    "LIPI": {
        "model": "AF-Q6XZB0-F1-model_v6", "patch": "patch002",
        "model_sha256": "e0c3ad43cbbd7c068f7ca6a6d735d237d69538d7573d600e72dac04a44e9c554",
        "pose_sha256": "da20c50811fd1e9bdc4591872972097d7b66a01e1604df88dcbb1da01059c047",
    },
}

# Transcribed from the existing manuscript Figure 6a and 6b. These are checks,
# never input to the reconstructed numerical results.
FIG6A = {
    (290, 10): 2.65, (306, 6): 2.99,
    (307, 8): 2.73, (307, 9): 3.10,
    (309, 5): 3.84, (309, 6): 3.12, (309, 7): 3.00,
    (309, 8): 2.53, (309, 9): 3.95, (309, 10): 3.23,
    (346, 1): 2.85, (346, 2): 2.96, (432, 1): 2.91,
    (434, 4): 3.10, (434, 5): 3.61,
    (441, 2): 3.90, (441, 3): 2.96, (441, 4): 2.81,
}
FIG6A_PLDDT = {
    290: 98.50, 306: 95.38, 307: 94.56, 309: 97.50,
    346: 98.38, 432: 98.19, 434: 97.81, 441: 96.62,
}
FIG6B = {306: 2.99, 307: 2.73, 309: 2.53, 323: 9.45,
         324: 9.05, 432: 2.91, 434: 3.10}
CUTOFFS = (3.5, 4.0, 5.0)
BASIC_NAMES = {"LYS": {"NZ"}, "ARG": {"NE", "NH1", "NH2"}}


def sha256(path):
    return hashlib.sha256(path.read_bytes()).hexdigest()


def locate(root, name, expected):
    candidates = [root / name] if (root / name).is_file() else sorted(root.rglob(name))
    matches = [p for p in candidates if sha256(p) == expected]
    if not matches:
        raise ValueError(f"No original hash-matched coordinate file found: {name}")
    return matches[0]


def read_atoms(path):
    atoms = []
    model_count = 0
    for line in path.read_text().splitlines():
        if line.startswith("MODEL "):
            model_count += 1
        if not line.startswith(("ATOM  ", "HETATM")):
            continue
        element = line[76:78].strip() if path.suffix == ".pdb" else line.split()[-1]
        if element.startswith("H") or element == "D":
            continue
        atom = {
            "serial": int(line[6:11]), "name": line[12:16].strip(),
            "resname": line[17:20].strip(), "chain": line[21:22].strip(),
            "resno": int(line[22:26]), "icode": line[26:27].strip(),
            "xyz": tuple(float(line[i:i + 8]) for i in (30, 38, 46)),
            "plddt": float(line[60:66]), "element": element,
        }
        if line[16:17].strip():
            raise ValueError(f"Unexpected alternate atom location: {path}")
        atoms.append(atom)
    assert model_count <= 1, f"Multiple coordinate models in {path}"
    assert len({a["serial"] for a in atoms}) == len(atoms)
    return atoms


def distance(a, b):
    return math.dist(a["xyz"], b["xyz"])


def nearest(a_atoms, b_atoms):
    # Serial numbers break exact numerical ties deterministically.
    return min((distance(a, b), a["serial"], b["serial"], a["name"], b["name"])
               for a in a_atoms for b in b_atoms)


def phosphate_groups(ligand):
    phosphorus = [a for a in ligand if a["element"] == "P"]
    oxygen = [a for a in ligand if a["element"] in ("O", "OA")]
    assert len(phosphorus) == 10 and len(oxygen) == 31 and len(ligand) == 41
    groups = {p["serial"]: [o for o in oxygen if distance(p, o) <= 1.9]
              for p in phosphorus}
    assert all(len(v) == 4 for v in groups.values())
    membership = defaultdict(list)
    for p_serial, os in groups.items():
        for o in os:
            membership[o["serial"]].append(p_serial)
    assert len(membership) == 31
    assert all(len(v) in (1, 2) for v in membership.values())
    bridges = {s for s, v in membership.items() if len(v) == 2}
    assert len(bridges) == 9
    adjacent = {p: set() for p in groups}
    for s in bridges:
        a, b = membership[s]
        adjacent[a].add(b)
        adjacent[b].add(a)
    endpoints = sorted(s for s, v in adjacent.items() if len(v) == 1)
    assert len(endpoints) == 2
    assert all(len(v) in (1, 2) for v in adjacent.values())
    order = [endpoints[0]]
    while len(order) < 10:
        nxt = adjacent[order[-1]] - set(order)
        assert len(nxt) == 1
        order.append(next(iter(nxt)))
    return groups, bridges, order, membership


def write_csv(path, rows):
    assert rows
    path.parent.mkdir(parents=True, exist_ok=True)
    with path.open("w", newline="") as f:
        w = csv.DictWriter(f, fieldnames=list(rows[0]))
        w.writeheader()
        w.writerows(rows)


def check(checks, name, expected, actual, decimals=None):
    tested = round(actual, decimals) if decimals is not None else actual
    ok = tested == expected
    checks.append({"check": name, "expected": expected, "actual": actual, "pass": ok})
    return ok


def reconstruct(coordinate_root, output):
    residue_rows, group_rows, basic_rows, connectivity_rows = [], [], [], []
    checks, source_records, summaries = [], [], {}
    calculations = {}
    for gene, cfg in SOURCES.items():
        model_file = cfg["model"] + ".pdb"
        pose_file = cfg["model"] + "_" + cfg["patch"] + "__polyP10__ad4.pdbqt"
        model_path = locate(coordinate_root, model_file, cfg["model_sha256"])
        pose_path = locate(coordinate_root, pose_file, cfg["pose_sha256"])
        for kind, path in (("model", model_path), ("saved pose", pose_path)):
            source_records.append({"protein": gene, "kind": kind, "file": path.name,
                                   "source_path": str(path.resolve()), "sha256": sha256(path)})
        protein, ligand = read_atoms(model_path), read_atoms(pose_path)
        groups, bridges, order, membership = phosphate_groups(ligand)
        ligand_by_serial = {a["serial"]: a for a in ligand}
        residues = defaultdict(list)
        for atom in protein:
            residues[(atom["chain"], atom["resno"], atom["icode"], atom["resname"])].append(atom)
        assert len({r[1] for r in residues}) == len(residues), "Unique residue numbering needed for checks"
        rec, bmap, bmap_nonbridge = {}, {}, {}
        for (chain, number, icode, resname), aa in sorted(residues.items()):
            assert len({a["plddt"] for a in aa}) == 1, "pLDDT differs within residue"
            base = {"protein": gene, "model": cfg["model"], "patch": cfg["patch"],
                    "chain": chain, "residue_number": number, "insertion_code": icode,
                    "residue": resname, "pLDDT": aa[0]["plddt"]}
            md, ps, ls, pn, ln = nearest(aa, ligand)
            rec[number] = {"distance": md, "plddt": aa[0]["plddt"], "resname": resname}
            rr = {**base, "minimum_distance_to_polyP10_A": md,
                  "closest_protein_atom": pn, "closest_protein_atom_serial": ps,
                  "closest_ligand_atom": ln, "closest_ligand_atom_serial": ls}
            for cutoff in CUTOFFS:
                rr[f"contact_at_{cutoff:g}_A"] = md <= cutoff
            residue_rows.append(rr)
            basic = [a for a in aa if a["name"] in BASIC_NAMES.get(resname, set())]
            for phosphate, p_serial in enumerate(order, 1):
                os = groups[p_serial]
                nonbridge = [o for o in os if o["serial"] not in bridges]
                d, ps, ls, pn, ln = nearest(aa, os)
                group_rows.append({**base, "phosphate_number": phosphate,
                                   "phosphorus_atom_serial": p_serial,
                                   "minimum_distance_to_phosphate_oxygen_A": d,
                                   "closest_protein_atom": pn, "closest_protein_atom_serial": ps,
                                   "closest_oxygen_atom_serial": ls,
                                   "closest_oxygen_is_bridging": ls in bridges,
                                   "contact_at_4_A": d <= 4.0})
                if basic:
                    bd, bps, bls, bpn, _ = nearest(basic, os)
                    nd, nps, nls, npn, _ = nearest(basic, nonbridge)
                    bmap[number, phosphate] = bd
                    bmap_nonbridge[number, phosphate] = nd
                    br = {**base, "phosphate_number": phosphate,
                          "phosphorus_atom_serial": p_serial,
                          "minimum_basic_N_to_phosphate_O_A": bd,
                          "closest_basic_nitrogen": bpn, "closest_basic_nitrogen_serial": bps,
                          "closest_oxygen_atom_serial": bls,
                          "closest_oxygen_is_bridging": bls in bridges,
                          "minimum_basic_N_to_nonbridging_O_A": nd,
                          "closest_basic_nitrogen_for_nonbridging_O": npn,
                          "closest_nonbridging_oxygen_serial": nls}
                    for cutoff in CUTOFFS:
                        br[f"any_oxygen_contact_at_{cutoff:g}_A"] = bd <= cutoff
                        br[f"nonbridging_oxygen_contact_at_{cutoff:g}_A"] = nd <= cutoff
                    basic_rows.append(br)
        for phosphate, p_serial in enumerate(order, 1):
            for oxygen in sorted(groups[p_serial], key=lambda a: a["serial"]):
                os = oxygen["serial"]
                connectivity_rows.append({"protein": gene, "model": cfg["model"], "patch": cfg["patch"],
                    "phosphate_number": phosphate, "phosphorus_atom_serial": p_serial,
                    "oxygen_atom_serial": os, "oxygen_is_bridging": os in bridges,
                    "phosphorus_oxygen_distance_A": distance(ligand_by_serial[p_serial], oxygen),
                    "phosphates_sharing_this_oxygen": ";".join(str(order.index(p) + 1) for p in sorted(membership[os]))})
        contacts = {r: v for r, v in rec.items() if v["distance"] <= 4.0}
        basic_contacts = {r: v for r, v in contacts.items() if v["resname"] in BASIC_NAMES}
        summary = {"protein_residues": len(rec), "protein_nonhydrogen_atoms": len(protein),
                   "ligand_phosphorus_atoms": 10, "ligand_oxygen_atoms": 31,
                   "bridging_oxygen_atoms": len(bridges), "phosphorus_serial_order": order,
                   "contacting_residues_at_4_A": sorted(contacts),
                   "basic_contacting_residues_at_4_A": sorted(basic_contacts),
                   "contacting_residue_mean_pLDDT": sum(v["plddt"] for v in contacts.values()) / len(contacts),
                   "basic_contacting_residue_mean_pLDDT": sum(v["plddt"] for v in basic_contacts.values()) / len(basic_contacts)}
        for cutoff in CUTOFFS:
            for label, bm in (("any_oxygen", bmap), ("nonbridging_oxygen", bmap_nonbridge)):
                summary[f"phosphates_with_basic_N_{label}_contact_at_{cutoff:g}_A"] = len({p for (r, p), d in bm.items() if d <= cutoff})
        summaries[gene] = summary
        calculations[gene] = {"residues": rec, "basic_map": bmap, "nonbridge_map": bmap_nonbridge}
        check(checks, f"{gene}: contacting residues at 4 A", 11, len(contacts))
        check(checks, f"{gene}: basic contacting residues at 4 A", 8 if gene == "LPL" else 6, len(basic_contacts))
        check(checks, f"{gene}: phosphates contacted by basic N at 4 A", 10,
              summary["phosphates_with_basic_N_any_oxygen_contact_at_4_A"])

    lpl, lipi = calculations["LPL"], calculations["LIPI"]
    shown = {k: v for k, v in lpl["basic_map"].items() if v <= 4.0}
    check(checks, "Figure 6a: complete displayed cell membership", sorted(FIG6A), sorted(shown))
    for key, expected in FIG6A.items():
        check(checks, f"Figure 6a: residue {key[0]}, phosphate {key[1]} distance", expected, shown[key], 2)
    for r, expected in FIG6A_PLDDT.items():
        check(checks, f"Figure 6a: residue {r} pLDDT", expected, lpl["residues"][r]["plddt"], 2)
    for r, expected in FIG6B.items():
        check(checks, f"Figure 6b: residue {r} distance", expected, lpl["residues"][r]["distance"], 2)
    for cutoff in CUTOFFS:
        check(checks, f"LPL: heparin determinants within {cutoff:g} A", 5,
              sum(lpl["residues"][r]["distance"] <= cutoff for r in FIG6B))
    for r, expected in {440: 7.10, 441: 2.74, 159: 29.04, 183: 24.74, 268: 28.62}.items():
        check(checks, f"LPL SI: residue {r} distance", expected, lpl["residues"][r]["distance"], 2)
    for r, expected in {159: 11.01, 183: 13.94, 253: 14.86}.items():
        check(checks, f"LIPI SI: residue {r} distance", expected, lipi["residues"][r]["distance"], 2)
    check(checks, "LPL: all phosphate contacts excluding bridging O at 4 A", 10,
          summaries["LPL"]["phosphates_with_basic_N_nonbridging_oxygen_contact_at_4_A"])
    check(checks, "LPL: phosphate contacts at 3.5 A, any oxygen", 9,
          summaries["LPL"]["phosphates_with_basic_N_any_oxygen_contact_at_3.5_A"])
    check(checks, "LPL: phosphate contacts at 3.5 A, excluding bridging O", 9,
          summaries["LPL"]["phosphates_with_basic_N_nonbridging_oxygen_contact_at_3.5_A"])
    check(checks, "LPL: eight basic residue mean pLDDT", 97.12,
          summaries["LPL"]["basic_contacting_residue_mean_pLDDT"], 2)
    observed_basic_plddt = [lpl["residues"][r]["plddt"]
                           for r in summaries["LPL"]["basic_contacting_residues_at_4_A"]]
    check(checks, "LPL: basic residue minimum pLDDT", 94.56, min(observed_basic_plddt), 2)
    check(checks, "LPL: basic residue maximum pLDDT", 98.50, max(observed_basic_plddt), 2)
    check(checks, "LIPI: contacting residue mean pLDDT", 97.84,
          summaries["LIPI"]["contacting_residue_mean_pLDDT"], 2)
    check(checks, "LIPI: every contacting residue pLDDT above 96", True,
          all(v["plddt"] > 96 for v in lipi["residues"].values() if v["distance"] <= 4.0))

    output.mkdir(parents=True, exist_ok=True)
    write_csv(output / "Data S9" / "LPL_LIPI_residue_contacts.csv", residue_rows)
    write_csv(output / "Data S9" / "LPL_LIPI_residue_phosphate_distances.csv", group_rows)
    write_csv(output / "Data S11" / "LPL_LIPI_basic_nitrogen_phosphate_distances.csv", basic_rows)
    write_csv(output / "Data S11" / "LPL_LIPI_phosphate_atoms.csv", connectivity_rows)
    write_csv(output / "checks" / "Figure6_and_SI_checks.csv", checks)
    (output / "checks" / "reconstruction_record.json").write_text(json.dumps({
        "description": "Deterministic reconstruction from original saved coordinates; not recovery of historical CSV files.",
        "sources": source_records, "summary": summaries,
        "distance_definition": "Euclidean distance in angstroms from unchanged PDB/PDBQT coordinates.",
        "cutoffs": "All thresholds apply to unrounded distances; 4 A means <= 4.000 A.",
        "phosphate_definition": "P-O distances <=1.9 A; chain traversal starts at the terminal P with the lower atom serial.",
        "basic_atoms": {k: sorted(v) for k, v in BASIC_NAMES.items()},
        "pLDDT_definition": "PDB B-factor field; one value per residue; consistency across atoms checked.",
        "output_rows": {"residue_contacts": len(residue_rows), "residue_phosphate_distances": len(group_rows),
                        "basic_nitrogen_phosphate_distances": len(basic_rows), "phosphate_atoms": len(connectivity_rows)},
        "checks": checks, "all_checks_pass": all(c["pass"] for c in checks),
    }, indent=2))
    print(json.dumps({"output": str(output), "all_checks_pass": all(c["pass"] for c in checks),
                      "checks": len(checks), "failed_checks": [c for c in checks if not c["pass"]],
                      "summary": summaries}, indent=2))
    if not all(c["pass"] for c in checks):
        raise SystemExit("Mismatch found. Do not use these tables until reviewed.")


if __name__ == "__main__":
    parser = argparse.ArgumentParser(description=__doc__)
    parser.add_argument("--coordinates", type=Path, required=True)
    parser.add_argument("--output", type=Path, required=True)
    args = parser.parse_args()
    reconstruct(args.coordinates, args.output)
```
