## Supplementary Code for "A structural screen for candidate polyphosphate binding surfaces in human membrane associated proteins": reproduce_analysis.py

```
"""Reproduce functional comparisons and score diagnostics from the supplied source data.
Requires Python 3.10+, numpy and scipy. No docking runs or source-file edits.
Usage: python reproduce_analysis.py --replicates 2000
Outputs tables and summary next to this script's parent folder.
"""
from pathlib import Path
from collections import defaultdict, Counter
import argparse, csv, hashlib, json, math, re
import numpy as np
from scipy import stats

ROOT=Path(__file__).resolve().parents[1]
SRC=ROOT/'sources'
OUT=ROOT/'tables'

def load(name):
    with (SRC/name).open(encoding='utf-8-sig',newline='') as f:
        return list(csv.DictReader(f))

def num(v):
    try:return float(v)
    except (ValueError,TypeError):return float('nan')

def write(name, records, keys=None):
    if not records: return
    with (OUT/name).open('w',encoding='utf-8-sig',newline='') as f:
        w=csv.DictWriter(f,fieldnames=keys or list(records[0]),extrasaction='ignore')
        w.writeheader();w.writerows(records)

def bh(p):
    p=np.asarray(p,float); order=np.argsort(p)
    qq=np.minimum.accumulate((p[order]*len(p)/np.arange(1,len(p)+1))[::-1])[::-1]
    ans=np.empty(len(p));ans[order]=np.minimum(qq,1);return ans

def main(replicates):
    OUT.mkdir(exist_ok=True)
    rows=load('master_input.csv'); heads=list(rows[0])
    numeric=[k for k in heads if k not in ['UniProt_ID','Entry_ID','Protein_Name','patch_id','patch_residues']]
    for i,r in enumerate(rows):
        r['source_csv_row']=i+2
        for k in numeric:r[k]=num(r[k])
        r['D']=-r['dock_best_score_ad4'];r['E']=r['patch_value_per_area']
        r['base_accession']=r['Entry_ID'].split('-')[1]
        aa=re.findall(r'([A-Z]{3})(-?\d+)',r['patch_residues']);c=Counter(t[0] for t in aa)
        r['n_patch_residues']=len(aa);r['n_HRK']=c['HIS']+c['ARG']+c['LYS']
        r['n_KR']=c['ARG']+c['LYS'];r['n_DE']=c['ASP']+c['GLU']
        r['pct_KR']=100*r['n_KR']/len(aa) if aa else np.nan
        if aa:assert abs(r['pct_HRK']-100*r['n_HRK']/len(aa))<1e-9
    done=[r for r in rows if all(math.isfinite(r[k]) for k in ['D','E','PRI','PDI100'])]
    hi=[r for r in done if r['PDI100']>80]
    cinput=load('curated_input.csv'); cm={r['patch_id']:r for r in cinput}
    cur=[r for r in done if r['patch_id'] in cm]
    assert (len(rows),len(done),len(hi),len(cur))==(23383,19481,185,74)
    byid={r['patch_id']:r for r in rows}
    for r in cinput:
        for k in heads:
            if k in numeric:assert math.isclose(num(r[k]),byid[r['patch_id']][k],rel_tol=1e-11,abs_tol=1e-11)
            else:assert r[k]==byid[r['patch_id']][k]
    pr={re.sub(r'__patch(\d+)$',lambda m:f'_patch{int(m[1]):03d}',r['patch_name']):r
        for r in load('vina_patch_boxes_20260219_070756_PRI_only.csv')}
    bx={r['structure_id']+f"_patch{int(r['patch_nr']):03d}":r
        for r in load('vina_patch_boxes_20260219_070756.csv')}
    vr={r['patch_id']:num(r['best_score']) for r in load('vina_results_ad4_polyP10.csv') if r['status']=='OK'}
    meta={r['Entry']:r for r in load('surface_candidates_annotation_snapshot.csv')}
    genes=defaultdict(list);models=defaultdict(list)
    for r in done:
        pid=r['patch_id'];b=bx[pid];p=pr[pid]
        assert r['dock_best_score_ad4']==vr[pid]
        assert abs(r['PRI']-num(p['pri']))<1e-12
        assert int(b['n_residues'])==r['n_patch_residues']
        assert abs(num(b['size_x'])-2*(num(b['R_patch_A'])+65))<1e-8
        r['box_size_x']=num(b['size_x']);r['box_R_patch_A']=num(b['R_patch_A'])
        r['pri_n_residues_in_box']=num(p['pri_n_residues_in_box'])
        r['unpenalized_PDI']=100*r['Dock01']**.6*r['Elec01']**.4
        r['PRI_penalty_multiplier']=math.exp(-3*max(0,.75-r['PRI']))
        assert abs(r['PDI100']-r['unpenalized_PDI']*r['PRI_penalty_multiplier'])<1e-10
        r['gene_symbol']=meta[r['base_accession']]['Gene Names'].split(' ')[0]
        r['source_keywords']=meta[r['base_accession']]['Keywords']
        r['Extracellular_Eligibility']=cm.get(pid,{}).get('Extracellular_Eligibility','NOT_CURATED_HERE')
        genes[r['base_accession']].append(r);models[r['Entry_ID']].append(r)
    bg=set(genes);hits={r['base_accession'] for r in hi}
    kw={g:{k.strip() for k in meta[g]['Keywords'].split(';') if k.strip()} for g in bg}
    keywords=Counter(k for g in bg for k in kw[g])
    can=[r for r in done if r['Entry_ID']==f"AF-{r['base_accession']}-F1"]
    cb={r['base_accession'] for r in can};ch={r['base_accession'] for r in can if r['PDI100']>80}
    length_q=np.quantile([num(meta[g]['Length']) for g in sorted(bg)],[1/3,2/3])
    bins=[1,3,7,15,np.inf]
    strata=defaultdict(set);strata_length=defaultdict(set)
    for g in bg:
        b=next(i for i,e in enumerate(bins) if len(genes[g])<=e)
        strata[b].add(g);strata_length[(b,int(np.searchsorted(length_q,num(meta[g]['Length']))))].add(g)
    def conditional(k,ss):
        prob=np.array([1.]);expected=0.;obs=sum(k in kw[g] for g in hits)
        for ids in ss.values():
            n=len(ids);draw=len(ids&hits);succ=sum(k in kw[g] for g in ids)
            expected+=draw*succ/n
            prob=np.convolve(prob,stats.hypergeom.pmf(np.arange(draw+1),n,succ,draw))
        pv=prob[np.abs(np.arange(len(prob))-expected)>=abs(obs-expected)-1e-10].sum()
        return min(1.,float(pv)),expected
    enrichment=[]
    for k,n in sorted(keywords.items()):
        if not 10<=n<=len(bg)-10:continue
        a=sum(k in kw[g] for g in hits);b=len(hits)-a;c=n-a;d=len(bg)-len(hits)-c
        st=stats.fisher_exact([[a,b],[c,d]],alternative='two-sided')
        ca=sum(k in kw[g] for g in ch);cn=sum(k in kw[g] for g in cb)
        ct=stats.fisher_exact([[ca,len(ch)-ca],[cn-ca,len(cb)-len(ch)-cn+ca]],alternative='two-sided')
        pp,pe=conditional(k,strata);pl,ple=conditional(k,strata_length)
        enrichment.append(dict(keyword=k,high_with=a,high_total=len(hits),background_with=n,background_total=len(bg),
            fold_enrichment=(a/len(hits))/(n/len(bg)),odds_ratio=float(st.statistic),p=float(st.pvalue),
            canonical_high_with=ca,canonical_high_total=len(ch),canonical_bg_with=cn,canonical_bg_total=len(cb),
            canonical_fold=(ca/len(ch))/(cn/len(cb)) if cn else np.nan,canonical_p=float(ct.pvalue),
            opportunity_p=pp,opportunity_expected=pe,opportunity_length_p=pl,opportunity_length_expected=ple))
    for pk,qk in [('p','q'),('canonical_p','canonical_q'),('opportunity_p','opportunity_q'),('opportunity_length_p','opportunity_length_q')]:
        for r,q in zip(enrichment,bh([r[pk] for r in enrichment])):r[qk]=float(q)
    assert len(enrichment)==219
    write('functional_keyword_enrichment_all219.csv',sorted(enrichment,key=lambda r:r['q']))
    gs=[]
    for g,rs in genes.items():
        best=max(rs,key=lambda r:r['PDI100']);ca=[r for r in rs if r['Entry_ID']==f'AF-{g}-F1']
        cs=[r for r in rs if r['patch_id'] in cm]
        gs.append(dict(UniProt_ID=g,Gene=meta[g]['Gene Names'].split(' ')[0],Protein_Name=best['Protein_Name'],
            n_models=len({r['Entry_ID'] for r in rs}),n_complete_patch_runs=len(rs),best_PDI=best['PDI100'],
            best_patch=best['patch_id'],canonical_best_PDI=max([r['PDI100'] for r in ca],default=''),
            n_PDI_above80=sum(r['PDI100']>80 for r in rs),n_curated_patch_runs=len(cs),source_length=int(meta[g]['Length']),
            Keywords=meta[g]['Keywords'],curated_status=';'.join(sorted({cm[r['patch_id']]['Extracellular_Eligibility'] for r in cs}))))
    gs.sort(key=lambda r:-r['best_PDI']);write('all2925_protein_record_summary.csv',gs)
    nox={'NOX1','CYBB','NOX3','NOX4','NOX5','DUOX1','DUOX2'}
    write('NOX_DUOX_family.csv',sorted([r for r in gs if r['Gene'] in nox],key=lambda r:r['Gene']))
    sels={k:{r['patch_id'] for r in sorted(done,key=lambda r:(-r[k],r['patch_id']))[:185]} for k in ['D','E','PDI100','pct_HRK']}
    ranking=[]
    for mode,s in sels.items():
        rr=[r for r in done if r['patch_id'] in s]
        d=dict(selection=mode,n_selected=185)
        for k in ['D','E','PRI','PDI100']:d['median_'+('PDI' if k=='PDI100' else k)]=float(np.median([r[k] for r in rr]))
        d['n_PRI_below_075']=sum(r['PRI']<.75 for r in rr);d['overlap_PDI_top185']=len(s&sels['PDI100']);ranking.append(d)
    write('equal_budget_rankings.csv',ranking)
    for k in ['D','E','PRI']:
        vals=np.sort([r[k] for r in done])
        for r in done:r[k+'_empirical_percentile']=100*np.searchsorted(vals,r[k],side='right')/len(vals)
    extra=['source_csv_row','gene_symbol','unpenalized_PDI','PRI_penalty_multiplier','D_empirical_percentile',
       'E_empirical_percentile','PRI_empirical_percentile','n_patch_residues','n_HRK','n_KR','n_DE','box_size_x',
       'box_R_patch_A','pri_n_residues_in_box','Extracellular_Eligibility','source_keywords']
    demoted=[r for r in done if r['unpenalized_PDI']>80 and r['PDI100']<=80]
    for name,rs in [('high_PDI185_annotated.csv',hi),('curated74_annotated.csv',cur),('PRI_demoted127.csv',demoted)]:
        write(name,sorted(rs,key=lambda r:-r['PDI100']),heads+extra)
    keys=['D','n_HRK','pct_HRK','patch_area','n_KR','pct_KR','n_patch_residues']
    X=np.array([[r[k] for k in keys] for r in done],float)
    mn,mi=np.unique([r['Entry_ID'] for r in done],return_inverse=True)
    gn,gi=np.unique([r['base_accession'] for r in done],return_inverse=True)
    def coeff(x,m):
        R=stats.rankdata(x,axis=0);C=np.corrcoef(R,rowvar=False)
        def partial(i,j,k):return (C[i,j]-C[i,k]*C[j,k])/math.sqrt((1-C[i,k]**2)*(1-C[j,k]**2))
        count=np.bincount(m,minlength=len(mn));cent=np.zeros_like(R)
        for j in range(R.shape[1]):
            mean=np.divide(np.bincount(m,weights=R[:,j],minlength=len(mn)),count,out=np.zeros(len(mn)),where=count>0)
            cent[:,j]=R[:,j]-mean[m]
        W=np.corrcoef(cent,rowvar=False)
        return np.array([C[0,1],C[0,2],C[0,3],partial(0,1,3),partial(0,2,3),W[0,1],W[0,2],W[0,3],C[0,4],C[0,5],partial(0,4,3),partial(0,5,3)])
    names=['HRK count','HRK percent','Patch area','HRK count, area-adjusted','HRK percent, area-adjusted',
        'HRK count, within-model','HRK percent, within-model','Patch area, within-model',
        'KR count','KR percent','KR count, area-adjusted','KR percent, area-adjusted']
    c=coeff(X,mi);rng=np.random.default_rng(20260908);groups=[np.flatnonzero(gi==i) for i in range(len(gn))];boot=[]
    for b in range(replicates):
        sel=rng.integers(0,len(groups),len(groups));idx=np.concatenate([groups[j] for j in sel]);boot.append(coeff(X[idx],mi[idx]))
    ci=np.quantile(boot,[.025,.975],axis=0)
    cr=[dict(analysis=n,association_with='raw_D_negated_AD4',estimate=float(v),cluster_bootstrap_95CI_low=float(lo),
        cluster_bootstrap_95CI_high=float(hi_),n_patches=len(done),n_base_clusters=len(gn),bootstrap_replicates=replicates,
        interpretation='Exploratory association; no experimental binding outcome; CI unadjusted for multiplicity.')
        for n,v,lo,hi_ in zip(names,c,*ci)]
    write('composition_docking_diagnostics.csv',cr)
    vr=[]
    for k in ['D','E','PRI','PDI100']:
        vals=np.array([r[k] for r in done]);means=np.bincount(mi,weights=vals)/np.bincount(mi)
        within=float(np.sum((vals-means[mi])**2)/np.sum((vals-vals.mean())**2))
        ranges=np.array([np.ptp([r[k] for r in rs]) for rs in models.values() if len(rs)>1])
        vr.append(dict(metric=k,within_model_sum_squares_fraction=within,between_model_sum_squares_fraction=1-within,
            n_multi_patch_models=len(ranges),median_within_model_range=float(np.median(ranges)),
            n_zero_range=int(sum(ranges<1e-12)),interpretation='Descriptive variance partition, not explained mechanistic variance.'))
    write('within_vs_between_model_variation.csv',vr)
    summary=dict(n_original=len(rows),n_complete=len(done),n_models=len(models),n_base=len(bg),n_high=len(hi),n_curated=len(cur),
        n_high_models=len({r['Entry_ID'] for r in hi}),n_high_base=len(hits),n_canonical_base=len(cb),n_canonical_high_base=len(ch),
        n_base_without_PRI_high=sum(r['unpenalized_PDI']>80 for r in done),n_below_after_PRI=len(demoted),
        n_high_PRI_penalized=sum(r['PRI']<.75 for r in hi),n_keyword_tests=len(enrichment),canonical_length_tertiles=length_q.tolist(),
        high_min_D_percentile=min(r['D_empirical_percentile'] for r in hi),high_min_E_percentile=min(r['E_empirical_percentile'] for r in hi),
        n_high_without_PRI_removed_fraction=len(demoted)/(len(demoted)+len(hi)),
        n_curated_unique_models=len({r['Entry_ID'] for r in cur}),n_curated_unique_base=len({r['base_accession'] for r in cur}),
        n_curated_single_high_patch_models=sum(n==1 for n in Counter(r['Entry_ID'] for r in cur).values()))
    (ROOT/'summary.json').write_text(json.dumps(summary,indent=2))
    print(json.dumps(summary,indent=2))

if __name__=='__main__':
    p=argparse.ArgumentParser();p.add_argument('--replicates',type=int,default=2000);a=p.parse_args()
    if a.replicates<100:raise ValueError('Use at least 100 bootstrap replicates; release used 2000.')
    main(a.replicates)
```
