## Supplementary Code for "A structural screen for candidate polyphosphate binding surfaces in human membrane associated proteins": expand_sensitivity.py

```
"""Evaluate selection of the 185 high scoring docking results across the 135 PDI settings.

Numerical reanalysis only: no docking, coordinates, annotations or score tuning.
Run with Python 3, numpy and pandas. Paths are relative to this script.
"""
from pathlib import Path
from itertools import product
import hashlib
import json
import platform
import shutil
import sys

import numpy as np
import pandas as pd

HERE = Path(__file__).resolve().parent
ROOT = HERE.parents[2] if len(HERE.parents) > 2 else HERE.parent
INPUT = HERE / "inputs"
INPUT.mkdir(exist_ok=True)
BUNDLED = (INPUT / "curated74.csv").exists()
SOURCE = INPUT if BUNDLED else HERE.parent / "source"
RAW_ORIGINAL = Path("C:/Users/Administrator/Desktop/delete/grand_patch_table_PDI100_polyP10.csv")
RAW = INPUT / RAW_ORIGINAL.name
if not RAW.exists():
    shutil.copy2(RAW_ORIGINAL, RAW)


def sha(path):
    return hashlib.sha256(Path(path).read_bytes()).hexdigest()


def write_json(name, data):
    (HERE / name).write_text(json.dumps(data, indent=2, ensure_ascii=False, allow_nan=False), encoding="utf-8")


def write_csv(name, frame):
    frame.to_csv(HERE / name, index=False, encoding="utf-8-sig", float_format="%.17g")


def scope_counts(frame):
    stable = frame[frame.selected_in_all_135]
    return {
        "results": len(frame), "models": frame.Entry_ID.nunique(), "entries": frame.base_accession.nunique(),
        "stable_results": len(stable), "models_with_stable_result": stable.Entry_ID.nunique(),
        "entries_with_stable_result": stable.base_accession.nunique(),
    }


def main():
    source_paths = [RAW, SOURCE / "curated74.csv", SOURCE / "Entry_Audit_128.csv"]
    before = {str(p): sha(p) for p in source_paths}
    data = pd.read_csv(RAW)
    data.insert(0, "source_csv_row", np.arange(2, len(data) + 2))
    assert data.patch_id.notna().all() and data.patch_id.is_unique
    required = ["dock_best_score_ad4", "patch_value_per_area", "PRI", "PDI100"]
    completed = data[np.isfinite(data[required].to_numpy(float)).all(axis=1)].copy()
    completed["completed_row_order"] = np.arange(1, len(completed) + 1)
    completed["base_accession"] = completed.UniProt_ID.str.split("-").str[0]
    assert completed.base_accession.equals(completed.Entry_ID.str.split("-").str[1])
    high_mask = completed.PDI100.to_numpy() >= 80
    high_indices = np.flatnonzero(high_mask)
    curated = pd.read_csv(SOURCE / "curated74.csv")
    keys74 = set(curated.patch_id)
    assert len(keys74) == 74
    nested_mask = completed.patch_id.isin(keys74).to_numpy()
    assert (len(data), len(completed), high_mask.sum(), nested_mask.sum()) == (23383, 19481, 185, 74)
    assert not (completed.PDI100 == 80).any()
    assert set(curated.patch_id).issubset(set(completed.loc[high_mask, "patch_id"]))
    source74 = data.set_index("patch_id").loc[curated.patch_id]
    assert np.allclose(source74[required], curated[required], rtol=0, atol=1e-10)

    D = -completed.dock_best_score_ad4.to_numpy()
    E = completed.patch_value_per_area.to_numpy()
    PRI = completed.PRI.to_numpy()
    allE = data.patch_value_per_area.to_numpy()
    Dmed, Dq1, Dq3 = np.median(D), np.percentile(D, 25), np.percentile(D, 75)
    Emed, Eq1, Eq3 = np.nanmedian(allE), np.nanpercentile(allE, 25), np.nanpercentile(allE, 75)
    Dz, Ez = (D - Dmed) / (Dq3 - Dq1), (E - Emed) / (Eq3 - Eq1)
    counts = np.zeros(len(completed), dtype=int)
    ranks185 = np.empty((185, 135), dtype=int)
    scores185 = np.empty((185, 135), dtype=float)
    settings, top_rows, all_ranks = [], [], []
    baseline_error = None
    for j, (wd, scale, ref, penalty) in enumerate(product([.4, .5, .6, .7, .8], [.75, 1, 1.25], [.7, .75, .8], [1.5, 3, 4.5])):
        dock = 1 / (1 + np.exp(-np.clip(Dz / scale, -50, 50)))
        elec = 1 / (1 + np.exp(-np.clip(Ez / scale, -50, 50)))
        score = 100 * dock ** wd * elec ** (1 - wd) * np.exp(-penalty * np.maximum(ref - PRI, 0))
        order = np.argsort(-score, kind="stable")
        ranks = np.empty(len(order), dtype=int)
        ranks[order] = np.arange(1, len(order) + 1)
        top = order[:185]
        counts[top] += 1
        ranks185[:, j], scores185[:, j] = ranks[high_indices], score[high_indices]
        settings.append({"setting_id": j + 1, "weight_D": wd, "weight_E": 1 - wd,
                         "sigmoid_scale": scale, "PRI_reference": ref, "lambda": penalty,
                         "top_count": 185, "original185_selected": int(high_mask[top].sum()),
                         "exposure_class74_selected": int(nested_mask[top].sum()),
                         "other111_selected": int((high_mask[top] & ~nested_mask[top]).sum()),
                         "above80_count": int((score >= 80).sum()),
                         "score_at_rank185": float(score[order[184]]),
                         "tie_at_rank185_boundary": bool(score[order[184]] == score[order[185]])})
        for ix in top:
            top_rows.append({"setting_id": j + 1, "rank": int(ranks[ix]),
                             "patch_id": completed.iloc[ix].patch_id, "score": float(score[ix])})
        if (wd, scale, ref, penalty) == (.6, 1, .75, 3):
            baseline_error = float(np.max(np.abs(score - completed.PDI100.to_numpy())))
            assert baseline_error < 1e-10
            assert set(top) == set(high_indices)
    assert j + 1 == 135

    keep = ["source_csv_row", "completed_row_order", "base_accession", "UniProt_ID", "Entry_ID", "Protein_Name",
            "patch_id", "dock_best_score_ad4", "patch_value_per_area", "PRI", "Dock01", "Elec01", "PDI100"]
    out = completed.loc[high_mask, keep].copy().reset_index(drop=True)
    out["in_exposure_class74"] = out.patch_id.isin(keys74)
    out["selections_of_135"] = counts[high_indices]
    out["minimum_rank"] = ranks185.min(axis=1)
    out["median_rank"] = np.median(ranks185, axis=1)
    out["maximum_rank"] = ranks185.max(axis=1)
    out["selected_in_all_135"] = out.selections_of_135 == 135
    out["original_rank"] = np.argsort(np.argsort(-completed.PDI100.to_numpy(), kind="stable"), kind="stable")[high_indices] + 1
    audit = pd.read_csv(SOURCE / "Entry_Audit_128.csv")
    assert audit.Entry_ID.is_unique
    assert set(audit.Entry_ID) == set(out.Entry_ID)
    assert curated.groupby("Entry_ID").Extracellular_Eligibility.nunique().eq(1).all()
    label74 = curated.groupby("Entry_ID").Extracellular_Eligibility.first().to_dict()
    audit = audit.set_index("Entry_ID")
    out["Gene"] = out.Entry_ID.map(audit.Gene)
    out["historical_exposure_decision"] = out.Entry_ID.map(audit.Decision)
    out["exposure_decision"] = out.Entry_ID.map(label74).fillna(out.historical_exposure_decision)
    out["exposure_group"] = np.where(out.in_exposure_class74, "normal_or_conditional_class74", "other_location_evidence111")
    out["publication_group"] = np.where(out.in_exposure_class74, "Normal/conditional exposure group", "Other model annotations")
    label_map = {"YES": "Yes", "CONDITIONAL": "Conditional", "NO": "No", "REVIEW": "Review", "EXCLUDE": "Existence uncertain"}
    out["publication_model_annotation"] = out.exposure_decision.map(label_map)
    out["exposure_qualification"] = "Model-level classification; pose accessibility not established."
    minpp = out.Entry_ID.eq("AF-Q9UNW1-F1")
    out.loc[minpp, "exposure_qualification"] = "Historical CONDITIONAL for a soluble extracellular pool; excluded from the fixed74 by its cell-surface scope exception. Pose accessibility not established."
    out.loc[out.exposure_decision.eq("EXCLUDE"), "exposure_qualification"] = "Source audit identifies TAAR3P as a pseudogene with uncertain protein existence. Its AlphaFold model and stable computational ranking do not establish expression of a human membrane protein."
    assert out.loc[out.Entry_ID.eq("AF-Q6U736-F1"), "exposure_decision"].eq("YES").all()
    assert out.loc[minpp, "in_exposure_class74"].eq(False).all()
    corrections = [{"Entry_ID": k, "historical": audit.loc[k, "Decision"], "corrected74": v,
                    "source": "corrected74 CSV", "scope": "all exact high-scoring results of this model"}
                   for k, v in label74.items() if audit.loc[k, "Decision"] != v]
    assert len(corrections) == 1 and corrections[0]["Entry_ID"] == "AF-Q6U736-F1"
    assert out.groupby("Entry_ID").in_exposure_class74.nunique().eq(1).all()
    assert np.array_equal(out.groupby("Entry_ID").size().sort_index(), audit["Tab 2 patches"].sort_index())

    # Existing Table5's source-export annotation memberships, not a fresh query.
    ox = {"Q9Y5S8", "P04839", "Q9HBY0", "Q9NPH5", "Q9NRD9", "Q9NRD8", "P29474", "Q96PH1"}
    tr = {"O94911", "Q9NP58", "Q92887", "Q96J66", "Q9H172", "Q9NTI2", "O75110", "Q8NB49", "P33527"}
    assert ox.isdisjoint(tr) and (ox | tr).issubset(set(out.base_accession))
    out["functional_group"] = np.where(out.base_accession.isin(ox), "oxidoreductase", np.where(out.base_accession.isin(tr), "translocase", "other"))
    nested = out[out.in_exposure_class74].copy()
    stable74 = nested[nested.selected_in_all_135]
    assert (len(stable74), stable74.Entry_ID.nunique(), stable74.base_accession.nunique()) == (19, 15, 14)
    grid = pd.DataFrame(settings)
    overlap = grid.original185_selected
    hist, edges = np.histogram(overlap, bins=np.arange(100, 191, 10))
    assert (int(overlap.min()), float(overlap.median()), int(overlap.max())) == (105, 139, 185)
    assert hist.tolist() == [1, 14, 22, 34, 25, 17, 13, 8, 1]
    assert nested.groupby("functional_group").base_accession.nunique().to_dict() == {"oxidoreductase": 6, "translocase": 8, "other": 35}
    assert stable74.groupby("functional_group").base_accession.nunique().to_dict() == {"oxidoreductase": 4, "translocase": 3, "other": 7}

    groups, exposures, reps, model_rows = [], [], [], []
    scopes = {"all185": out, "exposure_class74": nested, "other111": out[~out.in_exposure_class74]}
    for scope, frame in scopes.items():
        for category, group in frame.groupby("functional_group", sort=True):
            record = {"scope": scope, "functional_group": category, **scope_counts(group)}
            record["percent_entries_with_stable_result"] = 100 * record["entries_with_stable_result"] / record["entries"]
            groups.append(record)
        for category, group in frame.groupby("exposure_decision", sort=True):
            exposures.append({"scope": scope, "exposure_decision": category, **scope_counts(group)})
        for accession, group in frame.groupby("base_accession", sort=False):
            # Highest original PDI; original row order resolves ties.
            representative = group.sort_values("PDI100", ascending=False, kind="stable").iloc[0].to_dict()
            representative.update({"scope": scope, **scope_counts(group)})
            reps.append(representative)
    for model, group in out.groupby("Entry_ID", sort=False):
        r = group.iloc[0]
        model_rows.append({"Entry_ID": model, "base_accession": r.base_accession, "Gene": r.Gene,
                           "historical_exposure_decision": r.historical_exposure_decision, "exposure_decision": r.exposure_decision,
                           "in_exposure_class74": bool(r.in_exposure_class74), "exposure_qualification": r.exposure_qualification,
                           "publication_model_annotation": r.publication_model_annotation,
                           **scope_counts(group)})
    long_ranks = pd.DataFrame({"patch_id": np.repeat(out.patch_id.to_numpy(), 135),
                               "setting_id": np.tile(np.arange(1, 136), 185),
                               "rank": ranks185.ravel(), "score": scores185.ravel()})
    long_ranks["selected_top185"] = long_ranks["rank"] <= 185
    write_csv("all185_sensitivity.csv", out)
    write_csv("exposure_class74_sensitivity.csv", nested)
    write_csv("stable_all185.csv", out[out.selected_in_all_135].sort_values("PDI100", ascending=False, kind="stable"))
    write_csv("all185_ranks_135_settings.csv", long_ranks)
    write_csv("settings135.csv", grid)
    write_csv("top185_each_setting.csv", pd.DataFrame(top_rows))
    write_csv("protein_representatives.csv", pd.DataFrame(reps))
    write_csv("functional_group_counts.csv", pd.DataFrame(groups))
    write_csv("exposure_counts.csv", pd.DataFrame(exposures))
    write_csv("model_exposure_audit128.csv", pd.DataFrame(model_rows))
    write_csv("functional_annotation_membership82.csv", out[["base_accession", "Gene", "functional_group"]].drop_duplicates())
    write_csv("histogram_original185_overlap.csv", pd.DataFrame({"bin_left": edges[:-1], "bin_right": edges[1:], "count": hist}))
    summary = {
        "status": "calculation complete",
        "question": "Track all original185 exact high-scoring docking results using the unchanged established135-setting ranking calculation.",
        "fixed_reference": {"positive_patch_records": len(data), "completed_results": len(completed),
                            "completed_models": completed.Entry_ID.nunique(), "completed_entries": completed.base_accession.nunique(),
                            "D_median": float(Dmed), "D_IQR": float(Dq3-Dq1), "E_median": float(Emed), "E_IQR": float(Eq3-Eq1),
                            "normalization_refitted": False, "original_score_reconstruction_max_abs_error": baseline_error},
        "scopes": {scope: scope_counts(frame) for scope, frame in scopes.items()},
        "overlap": {"median": float(overlap.median()), "minimum": int(overlap.min()), "maximum": int(overlap.max()), "histogram": hist.tolist()},
        "functional_groups": groups, "exposure_groups": exposures,
        "historical_exposure_corrections": corrections,
        "historical_scope_exception": {"Entry_ID": "AF-Q9UNW1-F1", "historical_decision": "CONDITIONAL", "fixed74_member": False,
                                       "source": "Fixed74 membership and model audit", "action": "Preserve soluble-pool qualification; do not recode as NO or REVIEW."},
        "entry_counts_warning": "One UniProt entry may have models in both exposure populations or multiple decision classes. Do not sum distinct entry counts across these groups.",
        "original_row_order": "CSV data row order; one-based source_csv_row includes header; numpy stable descending sort.",
        "score_ties_at_selection_boundary": int(grid.tie_at_rank185_boundary.sum()),
        "limitations": ["Model-level exposure labels do not establish saved-pose accessibility.", "The fixed74 class is retained as a nested descriptive population; no high-scoring result is excluded from all185 analysis.", "Original functional memberships come from the final V11.6.1 Results table; no annotation refresh or new inferential test.", "Settings measure sensitivity of the existing score ranking, not experimental validation or binding probability."],
    }
    write_json("summary.json", summary)
    assert all(sha(p) == digest for p, digest in before.items())
    write_json("rerun_provenance.json", {
        "source_sha256": before, "source_original_csv_path": str(RAW_ORIGINAL),
        "source_original_csv_sha256": sha(RAW_ORIGINAL) if RAW_ORIGINAL.exists() else None,
        "source_copy_exact": sha(RAW) == sha(RAW_ORIGINAL) if RAW_ORIGINAL.exists() else "original source unavailable on this machine; use bundled input hash",
        "established_code_function": "build_results_figures.py::fig7 (read only; formula and grid reproduced)",
        "functional_membership_source": "Fixed original memberships encoded in this script: 8 oxidoreductases and 9 translocases",
        "software": {"python": platform.python_version(), "numpy": np.__version__, "pandas": pd.__version__},
        "analysis_script_sha256": sha(__file__),
        "output_sha256": {p.name: sha(p) for p in HERE.glob("*.csv")},
    })
    print(json.dumps({"scopes": summary["scopes"], "functional_groups": groups, "exposure_groups": exposures, "reconstruction_error": baseline_error}, indent=2))


if __name__ == "__main__":
    sys.stdout.reconfigure(encoding="utf-8")
    main()
```
