## Supplementary Code for "A structural screen for candidate polyphosphate binding surfaces in human membrane associated proteins": export_publication_tables.py

```
"""Export publication CSVs without altering precision of the working CSV cells."""
from pathlib import Path
import csv
import math
import shutil


def read(path):
    with Path(path).open(encoding="utf-8-sig", newline="") as stream:
        return list(csv.DictReader(stream))


def write(path, rows):
    with Path(path).open("w", encoding="utf-8-sig", newline="") as stream:
        writer = csv.DictWriter(stream, fieldnames=list(rows[0]))
        writer.writeheader()
        writer.writerows(rows)


def export(source, destination):
    source, destination = Path(source), Path(destination)
    rows = read(source / "all185_sensitivity.csv")
    reps = read(source / "protein_representatives.csv")
    exclude = {"historical_exposure_decision", "exposure_decision", "exposure_group"}
    rename = {"publication_model_annotation": "model_annotation", "exposure_qualification": "annotation_qualification", "in_exposure_class74": "in_normal_conditional74"}
    clean = [{rename.get(k, k): v for k, v in row.items() if k not in exclude} for row in rows]
    write(destination / "Data_S2_V11.7.csv", clean)
    grid = read(source / "settings135.csv")
    grid_rename = {"exposure_class74_selected": "normal_conditional74_selected", "other111_selected": "other_model_annotations111_selected"}
    write(destination / "Data_S3_V11.7.csv", [{grid_rename.get(k, k): v for k, v in row.items()} for row in grid])
    raw = read(source / "inputs/grand_patch_table_PDI100_polyP10.csv")
    complete = [r for r in raw if all(r[k] and math.isfinite(float(r[k])) for k in ["dock_best_score_ad4", "patch_value_per_area", "PRI", "PDI100"])]
    fields = ["patch_id", "Entry_ID", "PDI100", "selections_of_135", "minimum_rank", "median_rank", "maximum_rank",
              "results", "models", "stable_results", "models_with_stable_result", "publication_model_annotation", "exposure_qualification"]
    output = []
    for row in reps:
        if row["scope"] != "all185" or row["functional_group"] not in ["oxidoreductase", "translocase"]:
            continue
        item = {k: row[k] for k in ["base_accession", "Gene", "Protein_Name", "functional_group"]}
        item.update({"all185_" + k: row[k] for k in fields})
        nested = [r for r in reps if r["scope"] == "exposure_class74" and r["base_accession"] == row["base_accession"]]
        assert len(nested) <= 1
        item["has_normal_conditional74_result"] = bool(nested)
        for key in fields:
            item["normal_conditional74_" + key] = nested[0][key] if nested else (0 if key in ["results", "models", "stable_results", "models_with_stable_result"] else "")
        canonical = [r for r in complete if r["Entry_ID"] == "AF-" + row["base_accession"] + "-F1"]
        best = max(canonical, key=lambda r: float(r["PDI100"])) if canonical else {}
        item["best_completed_canonical_PDI100"] = best.get("PDI100", "")
        item["best_completed_canonical_patch_id"] = best.get("patch_id", "")
        output.append(item)
    assert len(output) == 17
    write(destination / "Data_S8_V11.7.csv", output)
    shutil.copyfile(source / "all185_ranks_135_settings.csv", destination / "Data_S2_ranks_V11.7.csv")


if __name__ == "__main__":
    here = Path(__file__).resolve().parent
    export(here, here)
```
