## Supplementary Code for "A structural screen for candidate polyphosphate binding surfaces in human membrane associated proteins": calculate_contacts.py.recovered.txt

"""Exact saved-pose residue contacts from the captured 74-pose Drive snapshot.

No docking, coordinate transform, residue renumbering, confidence filtering,
or replacement model download is performed. Input files remain unchanged.
"""
from pathlib import Path
from collections import Counter, defaultdict
import csv, hashlib, json, math, re
import numpy as np

HERE=Path(__file__).resolve().parent
CUTOFF=4.0
ONE=dict(zip(['ALA','ARG','ASN','ASP','CYS','GLN','GLU','GLY','HIS','ILE','LEU','LYS','MET','PHE','PRO','SER','THR','TRP','TYR','VAL'],'ARNDCQEGHILKMFPSTWYV'))
THREE={k:k.capitalize() for k in ONE}

def digest(path):return hashlib.sha256(path.read_bytes()).hexdigest()

def parse(path,ligand=False):
    text=path.read_text(encoding='ascii')
    lines=text.splitlines()
    models=[s[6:].strip() for s in lines if s.startswith('MODEL ')]
    if len(models)>1:raise ValueError(f'Multiple models in {path.name}')
    if ligand and models!=['1']:raise ValueError(f'Expected ligand MODEL1: {path.name}: {models}')
    records=[];serials=set();identities=set();all_elements=Counter();scores=[]
    for s in lines:
        if s.startswith('REMARK VINA RESULT:'):scores.append(float(s.split()[3]))
        record=s[:6].strip()
        if record not in ['ATOM','HETATM']:continue
        if not ligand and record!='ATOM':raise ValueError(f'Unexpected receptor HETATM: {path.name}')
        if s[16].strip():raise ValueError(f'Alternate location needs explicit resolution: {path.name}')
        name=s[12:16].strip()
        if ligand:
            atype=s.split()[-1]
            element='H' if atype in ['H','HD','HS'] else 'O' if atype=='OA' else atype.upper()
        else:element=s[76:78].strip().upper()
        if not element:raise ValueError(f'Missing element: {path.name}: {s}')
        all_elements[element]+=1
        if element in ['H','D']:continue
        serial=int(s[6:11]);chain=s[21].strip();num=int(s[22:26]);icode=s[26].strip();resname=s[17:20].strip()
        identity=(chain,num,icode,resname,name)
        # Ligand atom labels repeat (e.g. O/P); serials identify saved ligand atoms.
        if serial in serials or (not ligand and identity in identities):raise ValueError(f'Duplicate atom: {path.name} {identity}')
        serials.add(serial);identities.add(identity)
        xyz=tuple(float(s[i:i+8]) for i in [30,38,46])
        b=float(s[60:66]);occ=float(s[54:60])
        if not all(math.isfinite(x) for x in (*xyz,b,occ)):raise ValueError('Nonfinite atom data')
        if not ligand and (resname not in ONE or not 0<=b<=100):raise ValueError(f'Invalid protein residue/confidence: {path.name}')
        if occ!=1.0:raise ValueError(f'Unexpected occupancy: {path.name}')
        records.append({'serial':serial,'name':name,'element':element,'chain':chain,'number':num,'icode':icode,'resname':resname,'xyz':xyz,'plddt':b})
    if not records:raise ValueError(f'Empty file: {path.name}')
    if ligand:
        if Counter(a['element'] for a in records)!=Counter({'O':31,'P':10}):raise ValueError(f'Ligand heavy-atom composition: {path.name}')
        if len(scores)!=1:raise ValueError(f'Unexpected score records: {path.name}')
    else:
        for key,group in group_residues(records).items():
            if len({a['plddt'] for _,a in group})!=1:raise ValueError(f'Nonuniform residue pLDDT: {path.name} {key}')
            if sum(a['name']=='CA' for _,a in group)!=1:raise ValueError(f'Missing/duplicate CA: {path.name} {key}')
    return records,{'model':models[0] if models else 'single unnumbered','heavy_atoms':len(records),'hydrogens_excluded':all_elements['H']+all_elements['D'],'saved_score':scores[0] if scores else None}

def group_residues(atoms):
    out=defaultdict(list)
    for i,a in enumerate(atoms):out[(a['chain'],a['number'],a['icode'],a['resname'])].append((i,a))
    return out

def band(value):
    return 'Very high (>=90)' if value>=90 else 'Confident (70-<90)' if value>=70 else 'Low (50-<70)' if value>=50 else 'Very low (<50)'

def write_csv(name,rows):
    with (HERE/name).open('w',encoding='utf-8-sig',newline='') as f:
        w=csv.DictWriter(f,fieldnames=list(rows[0]));w.writeheader();w.writerows(rows)

def main():
    snapshot=json.loads((HERE/'input_snapshot.json').read_text())
    files=snapshot['files'];by_name={f['title']:f for f in files}
    if len(by_name)!=130:raise ValueError('Expected130unique files')
    for f in files:
        p=HERE/f['local_relative_path']
        assert p.stat().st_size==int(f['size'])
        assert digest(p)==f['sha256']
    poses=sorted([f for f in files if f['title'].endswith('.pdbqt')],key=lambda f:(f['folder_title'].lower(),f['title']))
    assert len(poses)==74
    contacts=[];summaries=[];all_residues=[];validation=[];model_cache={}
    for seq,f in enumerate(poses,1):
        m=re.fullmatch(r'(AF-([A-Z0-9]+)(?:-\d+)?-F\d+-model_v\d+)_(patch\d+)__polyP10__ad4\.pdbqt',f['title'])
        if not m:raise ValueError(f'Unparsed pose: {f["title"]}')
        model,accession,patch=m.groups();pose_id=f'{model}_{patch}'
        rf=by_name[model+'.pdb']
        assert rf['folder_id']==f['folder_id'],'Model/ligand are in different source folders'
        if model not in model_cache:model_cache[model]=parse(HERE/rf['local_relative_path'])
        rec,rec_info=model_cache[model];lig,lig_info=parse(HERE/f['local_relative_path'],True)
        rxyz=np.array([a['xyz'] for a in rec],dtype=np.float64)
        lxyz=np.array([a['xyz'] for a in lig],dtype=np.float64)
        delta=rxyz[:,None,:]-lxyz[None,:,:]
        d2=np.einsum('ijk,ijk->ij',delta,delta)
        best_lig=np.argmin(d2,axis=1)
        atom_min=d2[np.arange(len(rec)),best_lig]
        group=group_residues(rec);pose_contacts=[];all_distances=[]
        for key,ratoms in sorted(group.items()):
            chain,num,icode,resname=key
            ri=min((i for i,a in ratoms),key=lambda i:atom_min[i]);li=int(best_lig[ri])
            pa=rec[ri];la=lig[li];sq=float(atom_min[ri]);dist=math.sqrt(sq);confidence=pa['plddt']
            row={'pose_id':pose_id,'protein':f['folder_title'],'uniprot_base':accession,'model_id':model,'patch':patch,'chain':chain,
                 'residue_number':num,'insertion_code':icode,'residue_name':resname,'residue_label':f'{THREE[resname]}{num}{icode}',
                 'minimum_distance_A':dist,'pLDDT':confidence,'confidence_band':band(confidence),'pLDDT_below_70':'Yes' if confidence<70 else 'No',
                 'pLDDT_below_50':'Yes' if confidence<50 else 'No','nearest_protein_atom':pa['name'],'nearest_protein_atom_serial':pa['serial'],
                 'nearest_ligand_atom':la['name'],'nearest_ligand_atom_serial':la['serial']}
            all_residues.append({**row,'within_4A':sq<=CUTOFF*CUTOFF})
            all_distances.append(dist)
            if sq<=CUTOFF*CUTOFF:pose_contacts.append(row)
        contacts.extend(pose_contacts)
        n=len(pose_contacts)
        vals=[r['pLDDT'] for r in pose_contacts]
        names=[r['residue_label'] for r in pose_contacts]
        if not n:raise ValueError(f'No contact residues: {pose_id}')
        summaries.append({'pose_id':pose_id,'protein':f['folder_title'],'uniprot_base':accession,'model_id':model,'patch':patch,
            'contact_count':n,'mean_contact_pLDDT':float(np.mean(vals)),'median_contact_pLDDT':float(np.median(vals)),
            'minimum_contact_pLDDT':min(vals),'maximum_contact_pLDDT':max(vals),
            'n_pLDDT_ge90':sum(v>=90 for v in vals),'n_pLDDT_70_to90':sum(70<=v<90 for v in vals),
            'n_pLDDT_50_to70':sum(50<=v<70 for v in vals),'n_pLDDT_lt50':sum(v<50 for v in vals),
            'n_pLDDT_lt70':sum(v<70 for v in vals),'fraction_pLDDT_ge70':sum(v>=70 for v in vals)/n,
            'lys_arg_count':sum(r['residue_name'] in ['LYS','ARG'] for r in pose_contacts),
            'minimum_distance_A':min(r['minimum_distance_A'] for r in pose_contacts),
            'saved_AD4_score_kcal_mol':lig_info['saved_score'],'contact_residues':', '.join(names),
            'contacts_with_pLDDT':'; '.join(f"{r['residue_label']} ({r['pLDDT']:.2f})" for r in pose_contacts),
            'ligand_filename':f['title'],'receptor_filename':rf['title'],'ligand_Drive_URL':f['url'],'receptor_Drive_URL':rf['url']})
        validation.append({'pose_id':pose_id,'protein_model_record':rec_info['model'],'ligand_model_record':lig_info['model'],
            'receptor_heavy_atoms':rec_info['heavy_atoms'],'receptor_residues':len(group),'ligand_heavy_atoms':lig_info['heavy_atoms'],
            'ligand_hydrogens_excluded':lig_info['hydrogens_excluded'],'minimum_residue_distance_A':min(all_distances),
            'residues_within_3_5A':sum(x<=3.5 for x in all_distances),'residues_within_4A':n,'residues_within_5A':sum(x<=5 for x in all_distances),
            'nearest_residue_to_cutoff_difference_A':min(abs(x-4) for x in all_distances),
            'input_validation':'Matched model; single pose; correct composition; uniform residue pLDDT'})
    assert len(summaries)==74 and len(model_cache)==56 and len({s['uniprot_base'] for s in summaries})==49
    assert len({(r['pose_id'],r['chain'],r['residue_number'],r['insertion_code']) for r in contacts})==len(contacts)
    stats={'pose_count':74,'model_count':56,'base_protein_count':49,'contact_observations':len(contacts),
      'distinct_model_residues':len({(r['model_id'],r['chain'],r['residue_number'],r['insertion_code']) for r in contacts}),
      'poses_with_any_contact_pLDDT_below70':sum(s['n_pLDDT_lt70']>0 for s in summaries),
      'poses_with_any_contact_pLDDT_below50':sum(s['n_pLDDT_lt50']>0 for s in summaries),
      'poses_all_contacts_pLDDT_ge70':sum(s['n_pLDDT_lt70']==0 for s in summaries),
      'poses_all_contacts_pLDDT_ge90':sum(s['n_pLDDT_ge90']==s['contact_count'] for s in summaries),
      'minimum_contact_count':min(s['contact_count'] for s in summaries),'maximum_contact_count':max(s['contact_count'] for s in summaries),
      'contact_pLDDT_bands':dict(Counter(r['confidence_band'] for r in contacts)),
      'closest_residue_to_4A_boundary_A':min(v['nearest_residue_to_cutoff_difference_A'] for v in validation)}
    # Previously saved LPL/LIPI are controls, not inputs to the new calculation.
    controls={}
    for accession,n,mean in [('P06858',11,95.94363636363637),('Q6XZB0',11,97.8409090909091)]:
        s=next(s for s in summaries if s['model_id']==f'AF-{accession}-F1-model_v6')
        assert s['contact_count']==n and abs(s['mean_contact_pLDDT']-mean)<1e-10
        controls[accession]={'contact_count':s['contact_count'],'mean_contact_pLDDT':s['mean_contact_pLDDT'],'minimum_contact_pLDDT':s['minimum_contact_pLDDT']}
    # Threshold behavior is checked on squared distances, before display rounding.
    assert (3.999999**2<=16) and (4.0**2<=16) and not (4.000001**2<=16)
    sources=[{'filename':f['title'],'file_type':'Ligand pose' if f['title'].endswith('.pdbqt') else 'AlphaFold model',
      'protein_folder':f['folder_title'],'Drive_file_id':f['id'],'Drive_URL':f['url'],'bytes':int(f['size']),
      'modified_UTC':f['modified_time'],'retrieved_UTC':snapshot['retrieved_utc'],'sha256':f['sha256']} for f in files]
    methods=[
      {'topic':'Population','definition':'All 74 saved polyP10 poses in the current Drive docking-pose folder, paired to 56 exact AlphaFold models representing 49 base UniProt proteins. No confidence-based exclusion.'},
      {'topic':'Contact definition','definition':'One row per pose/protein-residue pair when any protein non-hydrogen atom is at most 4.000 Å from any ligand non-hydrogen atom. Inclusive cutoff applied before rounding.'},
      {'topic':'Residue numbering','definition':'Original AlphaFold model numbering, chain and insertion code. No signal-peptide offset or cross-isoform remapping. Exact model/isoform/version retained.'},
      {'topic':'pLDDT source','definition':'B-factor field in the matched original AlphaFold PDB. Every atom in each residue was checked for the same pLDDT; one value per residue enters summaries.'},
      {'topic':'Confidence bands','definition':'Very high: pLDDT >= 90; confident: 70 <= pLDDT < 90; low: 50 <= pLDDT < 70; very low: pLDDT < 50. Threshold flags are descriptive, not removal rules.'},
      {'topic':'Summary weighting','definition':'Mean/median/minimum/maximum use each contacting residue once per pose. Counts across poses are contact observations; the same model residue can appear for more than one patch.'},
      {'topic':'Geometric provenance','definition':'Minimum-distance rows retain the nearest protein atom and saved ligand atom serial. Coordinates were neither redocked nor aligned, relaxed or transformed.'},
      {'topic':'Confidence interpretation','definition':'pLDDT describes local model confidence, not docking-pose correctness or experimental binding. High local confidence does not establish relative domain orientation.'},
      {'topic':'Source folder','definition':'https://drive.google.com/drive/folders/1vD_rMLr8Pxp0Fd4OeSwJuSvO5G-H_LJO'},
      {'topic':'AlphaFold confidence reference','definition':'https://pmc.ncbi.nlm.nih.gov/articles/PMC8728224/'},
      {'topic':'AlphaFold pLDDT guidance','definition':'https://alphafold.ebi.ac.uk/faq | https://www.ebi.ac.uk/training/online/courses/alphafold/inputs-and-outputs/evaluating-alphafolds-predicted-structures-using-confidence-scores/plddt-understanding-local-confidence/'},
      {'topic':'Validation','definition':'All downloaded byte sizes and saved hashes were checked. Exact file matches, unique residue and atom identifiers, finite coordinates, one ligand model, 10 P + 31 O heavy atoms and uniform valid pLDDT were required. LPL/LIPI reproduce V9 controls.'},
      {'topic':'Precision','definition':'Coordinates retain source 0.001 Å precision and pLDDT source 0.01 precision. Calculations use float64; workbook distances display 3 decimals and confidence 2 decimals without changing stored values.'},
      {'topic':'Independent verification','definition':'A separate parser and scalar distance calculation reproduced all 854 contact memberships, residue pLDDT values and nearest atom identifiers exactly. Maximum distance difference was below 4.45e-16 Å; all 74 pose summaries agreed within 1.43e-14.'},
      {'topic':'Retained-roster cross-check','definition':'All 74 exact model/patch keys match grand_patch_table_PDI100_polyP10_extracellular_curated_FIXED.xlsx, sheet Extracell >80 PDI, E2:E75. No extra, missing or duplicate keys. Source: https://drive.google.com/file/d/1UCO9bH4ErQEwZ5ZcgG391BYUjAKyJpOv/view ; SHA-256: 4a6ee1e507697eca5c59126826fe4754ec4e99c8fe374425564ded722233b542.'},
      {'topic':'Coordinate snapshot','definition':f"All 130 coordinate files were downloaded from the current Drive folder on {snapshot['retrieved_utc']}. The Sources sheet records each original URL, filename, byte size and SHA-256 hash."},
      {'topic':'Population totals','definition':f"{len(contacts)} pose-residue contact observations correspond to {stats['distinct_model_residues']} distinct model/chain/residue identifiers. Multiple poses of one model can reuse a residue; neither quantity is a count of independent proteins."},
    ]
    payload={'summary':summaries,'contacts':contacts,'sources':sources,'methods':methods,'statistics':stats,'validation':validation,'controls':controls}
    (HERE/'workbook_payload.json').write_text(json.dumps(payload,indent=2),encoding='utf-8')
    write_csv('pose_summary.csv',summaries);write_csv('residue_contacts_4A.csv',contacts);write_csv('all_residue_minimum_distances.csv',all_residues);write_csv('input_validation.csv',validation)
    (HERE/'calculation_summary.json').write_text(json.dumps({'statistics':stats,'controls':controls,'numpy_version':np.__version__,'script_sha256':digest(Path(__file__))},indent=2),encoding='utf-8')
    print(json.dumps({'statistics':stats,'controls':controls},indent=2))

if __name__=='__main__':main()
