## Supplementary Code for "A structural screen for candidate polyphosphate binding surfaces in human membrane associated proteins": prepare_inputs.py

```
"""Freeze exact model sequences and reproduce the existing saved-pose contacts.

Original coordinate files and prior results are read-only. No docking is run.
"""
from pathlib import Path
from collections import Counter, defaultdict
from concurrent.futures import ThreadPoolExecutor
import csv, datetime, hashlib, json, re, runpy, subprocess
import numpy as np
import openpyxl

ROOT = Path(__file__).resolve().parents[2]
HERE = Path(__file__).resolve().parent
INPUT = HERE / 'inputs'
INPUT.mkdir(exist_ok=True)
UNI = INPUT / 'uniprot'
UNI.mkdir(exist_ok=True)
MASTER = Path('/Users/davidwu/Downloads/grand_patch_table_PDI100_polyP10_extracellular_curated_FIXED.xlsx')
OLDWB = Path('/Users/davidwu/Downloads/PolyP_Experimental_Site_Overlap.xlsx')
HISTORY = ROOT / 'artifacts/overlap72/history/calculate_contacts.py.recovered.txt'
historical = runpy.run_path(str(HISTORY))
parse, group = historical['parse'], historical['group_residues']
ONE = historical['ONE']

def sha(data): return hashlib.sha256(data).hexdigest()
def save(name, data): (INPUT/name).write_text(json.dumps(data, indent=2)+'\n')
def csvsave(name, records):
    with (INPUT/name).open('w',newline='') as f:
        w=csv.DictWriter(f,fieldnames=list(records[0])); w.writeheader(); w.writerows(records)

old = json.loads((HERE/'independent_input_audit/file_inventory.json').read_text())['files']
new = json.loads((ROOT/'artifacts/pre_exposure_download/local_download_readiness.json').read_text())['files']
downloaded = {r['filename']:r for r in json.loads((HERE/'drive_downloads/download_manifest.json').read_text())}
files = {}
for r in old:
    p = Path(downloaded[r['filename']]['local_path']) if r['filename'] in downloaded else Path(r['local_paths'][0])
    b = p.read_bytes()
    files[r['filename']] = dict(filename=r['filename'],local_path=str(p),drive_url=r['drive_url'],byte_size=len(b),sha256=sha(b),cohort='Original 56 models')
for r in new:
    p=Path(r['staged_path']); b=p.read_bytes()
    assert len(b)==r['byte_size'] and sha(b)==r['sha256']
    files[r['filename']]=dict(filename=r['filename'],local_path=str(p),drive_url='',byte_size=len(b),sha256=sha(b),cohort='Additional 72 models')
inventory=json.loads((ROOT/'artifacts/pre_exposure_download/final_drive_inventory.json').read_text())
def nested_files(obj):
    if isinstance(obj,dict):
        if obj.get('title','').endswith(('.pdb','.pdbqt')) and obj.get('url'): yield obj
        for v in obj.values(): yield from nested_files(v)
    elif isinstance(obj,list):
        for v in obj: yield from nested_files(v)
for r in nested_files(inventory):
    if r['title'] in files: files[r['title']]['drive_url']=r['url']
assert len(files)==313 and sum(f.endswith('.pdb') for f in files)==128
wb=openpyxl.load_workbook(MASTER,read_only=True,data_only=True)
vals=list(wb['>80 PDI'].values)
rows=[dict(zip(vals[0],r),source_row=i) for i,r in enumerate(vals[1:],2) if r[0]]
assert len(rows)==185 and len({r['Entry_ID'] for r in rows})==128
positive=json.loads((ROOT/'artifacts/overlap72/current_study_table_combined.json').read_text())
positive_poses={r['pose_id'] for r in positive if r['numerator']>0}
assert len(positive_poses)==31
models={}; receptors={}
for r in rows:
    mid=r['patch_id'].rsplit('_patch',1)[0]
    assert mid+'.pdb' in files and r['patch_id']+'__polyP10__ad4.pdbqt' in files
    if mid in models: continue
    f=files[mid+'.pdb']; p=Path(f['local_path']); rec,meta=parse(p); residues=list(group(rec))
    assert len({x[0] for x in residues})==1 and [x[1] for x in residues]==list(range(1,len(residues)+1))
    assert not any(x[2] for x in residues)
    sequence=''.join(ONE[k[3]] for k in residues)
    seqres=''.join(ONE[aa] for line in p.read_text().splitlines() if line.startswith('SEQRES') for aa in line[19:70].split())
    assert sequence==seqres,(mid,'ATOM/SEQRES differ')
    accession=re.fullmatch(r'AF-(.+)-F\d+-model_v\d+',mid).group(1); base=accession.split('-')[0]
    models[mid]=dict(model_id=mid,accession=accession,uniprot_base=base,protein=r['Protein_Name'],sequence=sequence,length=len(sequence),sequence_sha256=sha(sequence.encode()),cohort=f['cohort'],source_pdb_sha256=f['sha256'],source_pdb_url=f['drive_url'],chain=residues[0][0],atom_seqres_exact=True)
    receptors[mid]=rec

def get_uniprot(base):
    cached=ROOT/'artifacts/overlap72/literature_batch1'/f'{base}_uniprot.json'
    target=UNI/f'{base}.json'
    if not target.exists():
        if cached.exists(): target.write_bytes(cached.read_bytes())
        else: subprocess.run(['curl','--fail','--silent','--show-error','--location','--retry','2','--output',str(target),f'https://rest.uniprot.org/uniprotkb/{base}.json'],check=True)
    j=json.loads(target.read_text()); assert j['primaryAccession']==base
    return base,j
uniprot=dict(ThreadPoolExecutor(max_workers=5).map(get_uniprot,sorted({m['uniprot_base'] for m in models.values()})))
mapping=[]
for mid,m in models.items():
    j=uniprot[m['uniprot_base']]; can=j['sequence']['value']; features=[]
    if '-' in m['accession']:
        iso=[i for c in j.get('comments',[]) if c.get('commentType')=='ALTERNATIVE PRODUCTS' for i in c.get('isoforms',[]) if m['accession'] in i.get('isoformIds',[])]
        assert len(iso)==1,(mid,'isoform missing')
        if iso[0]['isoformSequenceStatus']=='Described':
            fdict={f['featureId']:f for f in j['features'] if f.get('type')=='Alternative sequence'}
            features=sorted([fdict[k] for k in iso[0]['sequenceIds']],key=lambda x:x['location']['start']['value'])
        else: assert iso[0]['isoformSequenceStatus']=='Displayed'
    seq=''; positions=[]; start=1
    for f in features:
        a=f['location']['start']['value']; b=f['location']['end']['value']; assert a>=start
        seq+=can[start-1:a-1]; positions.extend(range(start,a))
        alt=f.get('alternativeSequence',{}); original=alt.get('originalSequence')
        assert original is None or original==can[a-1:b]
        repl=alt.get('alternativeSequences',[]); assert len(repl)<=1
        replacement=repl[0] if repl else ''; seq+=replacement; positions.extend([None]*len(replacement)); start=b+1
    seq+=can[start-1:]; positions.extend(range(start,len(can)+1))
    assert seq==m['sequence'],(mid,'exact UniProt isoform differs',len(seq),m['length'])
    m['uniprot_exact_match']=True; m['canonical_positions']=positions; m['uniprot_version']=j.get('entryAudit',{})
    m['uniprot_locations']=[c for c in j.get('comments',[]) if c.get('commentType')=='SUBCELLULAR LOCATION']
    m['uniprot_topology']=[f for f in j.get('features',[]) if f.get('type') in ['Topological domain','Transmembrane','Intramembrane','Signal']]
    for i,aa in enumerate(seq,1): mapping.append(dict(model_id=mid,chain=m['chain'],sequence_position=i,pdb_residue_number=i,insertion_code='',aa=aa,canonical_position=positions[i-1]))
fasta=''.join('>'+mid+'\n'+'\n'.join(m['sequence'][i:i+80] for i in range(0,m['length'],80))+'\n' for mid,m in sorted(models.items()))
(INPUT/'PolyP_128_exact_models.fasta').write_text(fasta)
save('models.json',list(sorted(models.values(),key=lambda m:m['model_id'])))
csvsave('residue_number_map.csv',mapping)
save('source_files.json',list(files.values()))

contacts=[]; summaries=[]
for r in rows:
    pid=r['patch_id']; mid=pid.rsplit('_patch',1)[0]; rec=receptors[mid]; m=models[mid]
    f=files[pid+'__polyP10__ad4.pdbqt']; lig,meta=parse(Path(f['local_path']),True)
    assert abs(meta['saved_score']-r['dock_best_score_ad4'])<=.010000001,(pid,'score mismatch')
    xyz=np.array([a['xyz'] for a in rec],dtype=np.float64); lxyz=np.array([a['xyz'] for a in lig],dtype=np.float64)
    delta=xyz[:,None,:]-lxyz[None,:,:]; d2=np.einsum('ijk,ijk->ij',delta,delta); atommin=d2.min(axis=1)
    these=[]
    for key,atoms in group(rec).items():
        i=min((i for i,a in atoms),key=lambda i:atommin[i]); sq=float(atommin[i]); chain,num,icode,resname=key
        if sq>16.: continue
        cr=dict(pose_id=pid,model_id=mid,chain=chain,residue_number=num,insertion_code=icode,aa=ONE[resname],residue_label=ONE[resname]+str(num),minimum_distance_A=float(np.sqrt(sq)),pLDDT=rec[i]['plddt'],sequence_position=num)
        contacts.append(cr); these.append(cr)
    assert these
    summaries.append(dict(pose_id=pid,model_id=mid,protein=m['protein'],uniprot_base=m['uniprot_base'],cohort=m['cohort'],PDI100=r['PDI100'],saved_AD4_score=meta['saved_score'],contact_count=len(these),contact_residues='; '.join(x['residue_label'] for x in these),functional_overlap=pid in positive_poses,pose_sha256=f['sha256'],pose_url=f['drive_url']))
oldbook=openpyxl.load_workbook(OLDWB,read_only=True,data_only=True)
oldcounts={r[1]:int(r[3]) for r in list(oldbook['All74screen'].values)[1:] if r[1]}
assert len(oldcounts)==74
assert all(x['contact_count']==oldcounts[x['pose_id']] for x in summaries if x['pose_id'] in oldcounts)
newdata=json.loads((ROOT/'artifacts/overlap72/new111_contact_payload.json').read_text())
key=lambda r:(r['pose_id'],r['chain'],r['residue_number'],r['insertion_code'])
prior={key(r):r for r in newdata['contacts']}; now={key(r):r for r in contacts if r['pose_id'] not in oldcounts}
assert set(prior)==set(now)
maxdiff=max(abs(prior[k]['minimum_distance_A']-now[k]['minimum_distance_A']) for k in prior)
assert maxdiff<1e-12
save('contacts.json',contacts); save('pose_summary.json',summaries)
csvsave('contacts_185_4A.csv',contacts); csvsave('pose_manifest_185.csv',summaries)
audit=dict(time_utc=datetime.datetime.now(datetime.timezone.utc).isoformat(),models=len(models),poses=len(summaries),base_proteins=len(uniprot),residues=len(mapping),contacts=len(contacts),max_length=max(m['length'] for m in models.values()),fasta_sha256=sha(fasta.encode()),exact_isoform_matches=len(models),ATOM_SEQRES_matches=len(models),old74_contact_counts_match=True,new111_contact_identities_match=True,new111_maximum_distance_difference=maxdiff,master_sha256=sha(MASTER.read_bytes()),contact_parser_sha256=sha(HISTORY.read_bytes()),contact_rule='Minimum protein-ligand heavy-atom distance squared <=16 in original saved coordinates; same recovered historical parser and criteria; no pLDDT exclusion; no docking rerun.')
save('input_audit.json',audit); print(json.dumps(audit,indent=2))
```
