## Supplementary Code for "A structural screen for candidate polyphosphate binding surfaces in human membrane associated proteins": build_combined_figure.py

```
raw = pd.read_csv(SRC/'inputs/grand_patch_table_PDI100_polyP10.csv')
d = raw[raw[['dock_best_score_ad4','patch_value_per_area','PRI','PDI100']].notna().all(axis=1)].copy()
d['D']=-d.dock_best_score_ad4; d['E']=d.patch_value_per_area
d['Base']=d.Dock01**.6*d.Elec01**.4
d['accession']=d.Entry_ID.str.split('-').str[1]
metrics=['D','E','PRI','Base','PDI100']
means=d.groupby(['accession','Entry_ID'],sort=False)[metrics].mean().reset_index()
means['canonical']=means.Entry_ID.eq('AF-'+means.accession+'-F1')
canonical=means[means.canonical].set_index('accession')[metrics]
alternatives=means[~means.canonical].groupby('accession')[metrics].median()
pairs=canonical.join(alternatives,lsuffix='_canonical',rsuffix='_alternative',how='inner')
pairs.to_csv(ROOT/'research/isoform_pairs_reconstructed.csv')
rhos={k:float(np.corrcoef(pairs[k+'_canonical'].rank(),pairs[k+'_alternative'].rank())[0,1]) for k in ['D','E','PRI','Base','PDI100']}
assert [round(v,3) for v in rhos.values()]==[.760,.559,.822,.739,.790]
```
