## Supplementary Code for "A structural screen for candidate polyphosphate binding surfaces in human membrane associated proteins": audit_memberships.py

```
"""Read-only set audit for manuscript V11.8.1 figure; no new biological analysis."""
import csv, hashlib, json, re, zipfile
from collections import defaultdict
from pathlib import Path
import xml.etree.ElementTree as ET

ROOT=Path(__file__).resolve().parents[3]
OUT=Path(__file__).resolve().parent
NS={'x':'http://schemas.openxmlformats.org/spreadsheetml/2006/main'}
RID='{http://schemas.openxmlformats.org/officeDocument/2006/relationships}id'
SOURCES={
 'original_workbook':Path('/Users/davidwu/Downloads/PolyP_Experimental_Site_Overlap.xlsx'),
 'additional_workbook':ROOT/'outputs/01a0d2e4-aa8c-71c2-8a4d-f518fa65f4d0/PolyP_Experimental_Site_Overlap_72_Additional_Models.xlsx',
 'manuscript_results':ROOT/'tmp/manuscript_11_8_1/results.txt',
 'candidate_manifest':ROOT/'artifacts/topology128/results/All185_pose_locations.csv',
}

def read_sheet(path,name,header_row):
    with zipfile.ZipFile(path) as z:
        strings=[]
        if 'xl/sharedStrings.xml' in z.namelist():
            strings=[''.join(s.itertext()) for s in ET.fromstring(z.read('xl/sharedStrings.xml')).findall('x:si',NS)]
        wb=ET.fromstring(z.read('xl/workbook.xml'))
        rid=next(s.attrib[RID] for s in wb.find('x:sheets',NS) if s.attrib['name']==name)
        rels=ET.fromstring(z.read('xl/_rels/workbook.xml.rels'))
        target=next(r.attrib['Target'] for r in rels if r.attrib['Id']==rid)
        target=target.lstrip('/') if target.startswith('/') else 'xl/'+target
        xml=ET.fromstring(z.read(target))
        rows={}
        for row in xml.findall('x:sheetData/x:row',NS):
            cells={}
            for c in row.findall('x:c',NS):
                col=re.match(r'[A-Z]+',c.attrib['r']).group(); idx=0
                for ch in col: idx=idx*26+ord(ch)-64
                v=c.find('x:v',NS); typ=c.attrib.get('t')
                if typ=='s': value=strings[int(v.text)]
                elif typ=='inlineStr': value=''.join(c.find('x:is',NS).itertext())
                elif v is None: value=None
                elif typ in ('str','e'): value=v.text
                elif typ=='b': value=v.text=='1'
                else:
                    f=float(v.text); value=int(f) if f.is_integer() else f
                cells[idx]=value
            rows[int(row.attrib['r'])]=cells
        heads=rows[header_row]
        return [{**{h:cells.get(i) for i,h in heads.items() if h},'_excel_row':r,'_workbook':str(path),'_sheet':name}
                for r,cells in sorted(rows.items()) if r>header_row and any(v is not None for v in cells.values())]

def base_id(pose): return re.match(r'^AF-(\w+)',pose).group(1)
def model_id(pose): return pose.rsplit('_patch',1)[0]

manifest=list(csv.DictReader(SOURCES['candidate_manifest'].open()))
assert len(manifest)==185
by_id=defaultdict(list)
for r in manifest: by_id[r['uniprot_base']].append(r)
A=set(by_id)
text=SOURCES['manuscript_results'].read_text()
lines=text.split('Best–worst\nrank / 19,481\n')[1].split('\nFunction annotation')[0].splitlines()
assert len(lines)==45*7
stable=[dict(zip(['gene','uniprot','model','annotation','patch','pdi100','rank_range'],lines[i:i+7])) for i in range(0,len(lines),7)]
for r in stable:r['pose_id']=r['model']+'-model_v6_'+r['patch']
D={r['uniprot'] for r in stable}; stable_poses={r['pose_id'] for r in stable}
original=read_sheet(SOURCES['original_workbook'],'Site overlap',5)
additional=read_sheet(SOURCES['additional_workbook'],'Site overlap',5)
assert len(original)==41 and len(additional)==316
rows=original+additional
mapped=[r for r in rows if isinstance(r.get('Mapped site residues'),(int,float)) and r['Mapped site residues']>0]
positive=[r for r in mapped if r['Overlapping residues']>0]
B={base_id(r['Exact saved pose ID']) for r in mapped}
C={base_id(r['Exact saved pose ID']) for r in positive}
assert (len(A),len(B),len(C),len(D),len(B&D),len(C&D))==(82,44,18,24,13,5)
assert C<=B<=A and D<=A
assert len({r['Exact saved pose ID'] for r in mapped})==113
assert len({model_id(r['Exact saved pose ID']) for r in mapped})==74
assert len({r['Exact saved pose ID'] for r in positive})==31
assert len({model_id(r['Exact saved pose ID']) for r in positive})==23
positive_poses={r['Exact saved pose ID'] for r in positive}
assert {base_id(p) for p in (positive_poses&stable_poses)}==C&D

regions={
 'neither_level2_nor_mapped_site': A-(D|B),
 'level2_without_mapped_site': D-B,
 'mapped_site_without_overlap_outside_level2': B-(C|D),
 'mapped_site_without_overlap_in_level2': (B&D)-C,
 'site_overlap_outside_level2': C-D,
 'site_overlap_in_level2': C&D,
}
assert sum(map(len,regions.values()))==82
lookup={r['uniprot_base']:{'gene':r['gene'],'protein':r['protein']} for r in manifest}
# Preserve familiar aliases used in the manuscript alongside current gene symbols.
aliases={'P11166':'GLUT1','P04839':'NOX2'}
proteins=[]
for uid in sorted(A,key=lambda u:lookup[u]['gene']):
    ps=sorted(p for p in positive_poses if base_id(p)==uid)
    ms=sorted({r['Exact saved pose ID'] for r in mapped if base_id(r['Exact saved pose ID'])==uid})
    ss=sorted(p for p in stable_poses if base_id(p)==uid)
    proteins.append({
      'uniprot':uid,'gene':lookup[uid]['gene'],'manuscript_alias':aliases.get(uid,lookup[uid]['gene']),
      'protein_name':lookup[uid]['protein'],'level1':True,'level2':uid in D,
      'mapped_experimental_site':uid in B,'predicted_site_overlap':uid in C,
      'region':next(k for k,v in regions.items() if uid in v),
      'level1_pose_count':len(by_id[uid]),'level2_pose_count':len(ss),'mapped_pose_count':len(ms),'positive_pose_count':len(ps),
      'level2_pose_ids':'; '.join(ss),'mapped_pose_ids':'; '.join(ms),'positive_pose_ids':'; '.join(ps),
      'same_pose_stable_and_overlap':'; '.join(sorted(set(ss)&set(ps))),
      'site_source_rows':'; '.join(sorted({f"{Path(r['_workbook']).name} | {r['_sheet']}!row{r['_excel_row']}" for r in mapped if base_id(r['Exact saved pose ID'])==uid})),
    })
with (OUT/'protein_set_membership.csv').open('w',newline='') as f:
    w=csv.DictWriter(f,fieldnames=list(proteins[0]));w.writeheader();w.writerows(proteins)
sitecols=['_workbook','_sheet','_excel_row','Exact saved pose ID','Study-defined site','Mapped site residues','Overlapping residues','Experimental site-set ID']
with (OUT/'site_membership_evidence.csv').open('w',newline='') as f:
    w=csv.DictWriter(f,fieldnames=sitecols,extrasaction='ignore');w.writeheader();w.writerows(rows)

def describe(ids):
    return [{'uniprot':u,'gene':lookup[u]['gene'],'manuscript_alias':aliases.get(u,lookup[u]['gene'])} for u in sorted(ids,key=lambda u:lookup[u]['gene'])]

out={
 'manuscript':'V11.8.1','unit':'Unique base UniProt entries; alternative isoforms grouped',
 'definitions':{
   'level1':'At least one original high-scoring docking result with PDI100 >=80.',
   'level2':'At least one exact model-and-patch result remained in the top185 under all135 tested PDI parameter combinations, as listed in manuscript Table3.',
   'mapped_experimental_site':'At least one actual Site overlap comparison with Mapped site residues >0 in the original or additional workbook.',
   'predicted_site_overlap':'At least one actual Site overlap comparison with Overlapping residues >0 (a predicted polyP pose overlaps a mapped experimental site).',
 },
 'counts':{'level1':len(A),'level2':len(D),'mapped_experimental_site':len(B),'predicted_site_overlap':len(C),'level2_and_mapped_experimental_site':len(D&B),'level2_and_predicted_site_overlap':len(D&C),'mapped_poses':113,'mapped_models':74,'positive_poses':31,'positive_models':23,'level2_poses':45},
 'regions':{k:{'count':len(v),'members':describe(v)} for k,v in regions.items()},
 'sets':{'level1':describe(A),'level2':describe(D),'mapped_experimental_site':describe(B),'predicted_site_overlap':describe(C),'level2_and_mapped_experimental_site':describe(D&B),'level2_and_predicted_site_overlap':describe(D&C)},
 'cautions':['44 is evidence coverage, not44 validated polyP binders.','18 denotes predicted contact with experimentally characterised sites; polyP regulation itself is not experimentally established.','38 of82 lack a usable mapped site set in the reviewed sources; this is missing evidence, not a biological negative.','26 of44 have a usable mapped site set but no overlap for their assessed saved poses.','Do not use Reviewed sets >0 from screening sheets: it includes reviewed sets with zero residues mapped to an exact isoform and yields45 proteins/121poses instead of44/113. ADGRA2 is the extra protein; affected zero-mapped pose records also occur for ADCY10 and RGS6.'],
 'source_files':{k:{'path':str(p),'sha256':hashlib.sha256(p.read_bytes()).hexdigest()} for k,p in SOURCES.items()},
 'checks':{'union_regions_equals82':True,'all_regions_disjoint':all(not regions[a]&regions[b] for i,a in enumerate(regions) for b in list(regions)[i+1:]),'18_nested_in44_nested_in82':True,'24_nested_in82':True,'all_five_have_same_pose_stable_and_overlap':True,'original_and_additional_workbook_direct_read':True},
}
(OUT/'membership_evidence.json').write_text(json.dumps(out,indent=2,ensure_ascii=False)+'\n')
print(json.dumps(out['counts'],indent=2));print({k:len(v) for k,v in regions.items()});print('44intersect24',describe(B&D))
```
