## Supplementary Code for "A structural screen for candidate polyphosphate binding surfaces in human membrane associated proteins": audit_model_memberships.py

```
"""Regroup previously audited saved-pose memberships at exact AlphaFold-model grain."""
from pathlib import Path
from collections import defaultdict
import csv, hashlib, json, re

ROOT=Path(__file__).resolve().parents[3]
OUT=Path(__file__).resolve().parent
REVIEW=ROOT/'outputs/overlap_chart_v11_8_1/revision'
paths={
 'previous_membership':REVIEW/'protein_set_membership.csv',
 'previous_evidence':REVIEW/'membership_evidence.json',
 'site_membership_evidence':REVIEW/'site_membership_evidence.csv',
 'candidate_manifest':ROOT/'artifacts/topology128/results/All185_pose_locations.csv',
}
prev=list(csv.DictReader(paths['previous_membership'].open()))
old_audit=json.loads(paths['previous_evidence'].read_text())
manifest=list(csv.DictReader(paths['candidate_manifest'].open()))
sites=list(csv.DictReader(paths['site_membership_evidence'].open()))
def pose_set(col):return {p for r in prev for p in r[col].split('; ') if p}
def model(pose):return pose.rsplit('_patch',1)[0]
def models(poses):return {model(p) for p in poses}
sp=pose_set('level2_pose_ids'); ep=pose_set('mapped_pose_ids'); pp=pose_set('positive_pose_ids')
by_model=defaultdict(list)
for r in manifest:by_model[r['model_id']].append(r)
A=set(by_model);D=models(sp);B=models(ep);C=models(pp);J=models(sp&pp)
assert len(manifest)==185 and len({r['pose_id'] for r in manifest})==185
assert (len(A),len(D),len(B),len(C),len(D&B),len(D&C),len(J))==(128,34,74,23,18,7,6)
assert C<=B<=A and D<=A
assert (len(sp),len(ep),len(pp),len(sp&pp))==(45,113,31,7)
regions={
 'neither_level2_nor_mapped_site': A-(D|B),
 'level2_without_mapped_site': D-B,
 'mapped_site_without_overlap_outside_level2': B-(C|D),
 'mapped_site_without_overlap_in_level2': (B&D)-C,
 'site_overlap_outside_level2': C-D,
 'site_overlap_in_level2': C&D,
}
assert sum(map(len,regions.values()))==128
aliases={'P11166':'GLUT1','P04839':'NOX2'}
rows=[]
for m in sorted(A):
 base=by_model[m][0]['uniprot_base'];gene=by_model[m][0]['gene']
 acc=m[3:].removesuffix('-F1-model_v6');suffix=acc[len(base):]
 label='canonical' if not suffix else 'isoform '+suffix.lstrip('-')
 stable=sorted(p for p in sp if model(p)==m);mapped=sorted(p for p in ep if model(p)==m);positive=sorted(p for p in pp if model(p)==m)
 joint=sorted(set(stable)&set(positive))
 refs=sorted({f"{Path(r['_workbook']).name} | {r['_sheet']}!row{r['_excel_row']}" for r in sites if model(r['Exact saved pose ID'])==m and float(r['Mapped site residues'] or 0)>0})
 rows.append({
  'Entry_ID':m.removesuffix('-model_v6'),'model_id':m,'exact_accession':acc,'base_uniprot':base,'gene':gene,
  'manuscript_alias':aliases.get(base,gene),'model_isoform_label':label,'display_label':f'{aliases.get(base,gene)} ({label})',
  'level1':True,'level2':m in D,'mapped_experimental_site':m in B,'predicted_site_overlap':m in C,
  'same_pose_level2_and_overlap':m in J,'level2_and_overlap_only_on_different_poses':m in ((D&C)-J),
  'region':next(k for k,v in regions.items() if m in v),
  'level1_pose_count':len(by_model[m]),'level2_pose_count':len(stable),'mapped_pose_count':len(mapped),'positive_pose_count':len(positive),'same_pose_joint_count':len(joint),
  'level1_pose_ids':'; '.join(sorted(r['pose_id'] for r in by_model[m])),
  'level2_pose_ids':'; '.join(stable),'mapped_pose_ids':'; '.join(mapped),'positive_pose_ids':'; '.join(positive),
  'same_pose_level2_and_overlap_ids':'; '.join(joint),'site_source_rows':'; '.join(refs),
 })
with (OUT/'model_set_membership.csv').open('w',newline='') as f:
 w=csv.DictWriter(f,fieldnames=list(rows[0]));w.writeheader();w.writerows(rows)
lookup={r['model_id']:r for r in rows}
def describe(ms):
 cols=['Entry_ID','model_id','exact_accession','base_uniprot','gene','manuscript_alias','model_isoform_label','display_label','level2_pose_ids','positive_pose_ids','same_pose_level2_and_overlap_ids']
 return [{k:lookup[m][k] for k in cols} for m in sorted(ms)]
out={
 'manuscript':'V11.8.1','unit':'Exact AlphaFold model (Entry_ID; canonical and named isoform models counted separately); all model files version6',
 'method':'Regroup the already-audited exact pose IDs by their model identifier; no redocking or reinterpretation of experimental evidence.',
 'definitions':{
  'level1':'Exact model with at least one original saved docking result scoring PDI100 >=80.',
  'level2':'Exact model with at least one exact model-and-patch result selected under all135 PDI parameter combinations, from manuscript Table3.',
  'mapped_experimental_site':'Exact model with at least one saved-pose comparison containing >0 mapped experimental site residues.',
  'predicted_site_overlap':'Exact model with at least one saved pose contacting >0 residues of a mapped experimental set.',
  'level2_and_predicted_site_overlap':'Same exact model belongs to both sets; its qualifying patches may differ.',
  'same_pose_level2_and_overlap':'Stricter subset: one exact model-and-patch result meets both criteria.',
 },
 'counts':{'level1':len(A),'level2':len(D),'mapped_experimental_site':len(B),'predicted_site_overlap':len(C),'level2_and_mapped_experimental_site':len(D&B),'level2_and_predicted_site_overlap':len(D&C),'models_with_same_pose_level2_and_overlap':len(J),'models_with_joint_membership_only_on_different_poses':len((D&C)-J),'level1_poses':185,'level2_poses':len(sp),'mapped_poses':len(ep),'positive_poses':len(pp),'same_pose_level2_and_overlap_poses':len(sp&pp)},
 'regions':{k:{'count':len(v),'models':describe(v)} for k,v in regions.items()},
 'sets':{'level1':describe(A),'level2':describe(D),'mapped_experimental_site':describe(B),'predicted_site_overlap':describe(C),'level2_and_mapped_experimental_site':describe(D&B),'level2_and_predicted_site_overlap':describe(D&C),'same_pose_level2_and_overlap':describe(J),'joint_membership_only_on_different_poses':describe((D&C)-J)},
 'cautions':[
  'Counts now represent exact structural models, not unique UniProt protein entries and not saved poses.',
  'Canonical and named isoform models are kept separate. Experimental evidence is mapped from the recorded source species/construct; it does not imply that every modeled isoform was itself tested experimentally.',
  'The model-level intersection contains7 models. Six have a common exact pose satisfying both criteria; canonical MYO1C has stable patch003/patch013 and overlapping patch001/patch012, with no shared exact patch.',
  'Do not label the7-model intersection as7 jointly qualifying poses or7 models with a single jointly qualifying pose.',
  '44/18 protein-entry counts become74/23 exact-model counts. The evidence coverage definition remains mapped site residues >0, not reviewed sets >0.',
  'The54 models without usable mapped-site evidence are not biological negatives; the51 models with mapped evidence and no site overlap are nonoverlaps for the assessed saved poses and specified experimental sets.',
 ],
 'source_files':{k:{'path':str(p),'sha256':hashlib.sha256(p.read_bytes()).hexdigest()} for k,p in paths.items()},
 'upstream_workbook_sources':{k:v for k,v in old_audit['source_files'].items() if k.endswith('workbook')},
 'checks':{'expected_manuscript_population_counts_reproduced':True,'all_regions_disjoint':all(not regions[a]&regions[b] for i,a in enumerate(regions) for b in list(regions)[i+1:]),'regions_sum128':True,'23_nested_in74_nested_in128':True,'34_nested_in128':True,'same_pose_joint_subset_of_model_joint':J<=D&C},
}
(OUT/'model_membership_evidence.json').write_text(json.dumps(out,indent=2,ensure_ascii=False)+'\n')
print(json.dumps(out['counts'],indent=2));print({k:len(v) for k,v in regions.items()});print('different-pose exception:',describe((D&C)-J))
```
