## Supplementary Code for "A structural screen for candidate polyphosphate binding surfaces in human membrane associated proteins": build_v12.py

```
with open(ROOT.parent/'overlap_chart_v11_8_1/model_level/model_set_membership.csv') as f: rows=list(csv.DictReader(f))
field_map={'model_id':'AlphaFold_model','base_uniprot':'UniProt_entry','gene':'Gene','model_isoform_label':'Model_isoform','level1':'High_scoring_model','level2':'Consistently_selected_model','mapped_experimental_site':'Experimental_sites_mapped','predicted_site_overlap':'Functional_site_overlap','same_pose_level2_and_overlap':'Both_properties_in_same_pose','level2_and_overlap_only_on_different_poses':'Both_properties_only_in_different_poses','level1_pose_ids':'High_scoring_pose_IDs','level2_pose_ids':'Consistently_selected_pose_IDs','mapped_pose_ids':'Poses_with_mapped_sites','positive_pose_ids':'Poses_with_site_overlap','same_pose_level2_and_overlap_ids':'Poses_meeting_both_properties'}
with open(OUT/'PolyP_Data_S13_Model_Evidence_Groups_V12.csv','w',newline='') as f:
 writer=csv.DictWriter(f,fieldnames=list(field_map.values()));writer.writeheader()
 for r in rows:writer.writerow({new:r[old] for old,new in field_map.items()})
```
