## Supplementary Code for "A structural screen for candidate polyphosphate binding surfaces in human membrane associated proteins": Run instructions.txt

POLYP ANALYSIS CODE

Extract Supplementary_Data.zip, Supplementary_Code.zip and
Supplementary_Structures.zip into the same folder. Keep the resulting
"Supplementary data", "Supplementary code" and "Candidate structures" folders
together. Use a working copy and run the commands below from "Supplementary
code". Regenerated outputs do not
replace the submitted tables in "Supplementary data".

Software for the saved data analyses
-----------------------------------
Tested with Python 3.12.14, NumPy 2.3.5, pandas 2.2.3, SciPy 1.18.1 and
openpyxl 3.1.5. Install these packages in your Python environment:

python3 -m pip install numpy==2.3.5 pandas==2.2.3 scipy==1.18.1 openpyxl==3.1.5

The LPL/LIPI contact reconstruction uses only the Python standard library. The screening
workflow has separate external software requirements, listed below.

Prepare the shared inputs once
------------------------------
To avoid distributing identical tables several times, the package stores one
master CSV in Supplementary data and one curated74.csv in PDI selection.
curated74.csv contains the 74 high scoring results in the normal/conditional
cell surface exposure group.
Run this command once to place exact copies at the paths expected by the
analysis scripts (shown on one line):

python3 -c "from shutil import copyfile; master='../Supplementary data/S1 All screening results/grand_patch_table_PDI100_polyP10.csv'; copyfile(master, 'PDI selection/inputs/grand_patch_table_PDI100_polyP10.csv'); copyfile(master, 'Functional analyses/sources/master_input.csv'); copyfile('PDI selection/inputs/curated74.csv', 'Functional analyses/sources/curated_input.csv')"

This only copies supplied files; it performs no scientific calculation.

1. Reproduce selection under the 135 PDI settings
------------------------------------------------
python3 "PDI selection/expand_sensitivity.py"
python3 "PDI selection/export_publication_tables.py"

After preparation, "PDI selection/inputs" contains the original master CSV,
curated74.csv and Entry_Audit_128.csv. Entry_Audit_128.csv records the cell
surface exposure review for the 128 selected AlphaFold models. The first
command writes the working selection and rank tables beside the script; the
second exports Data_S2_V11.7.csv, Data_S2_ranks_V11.7.csv, Data_S3_V11.7.csv
and Data_S8_V11.7.csv there.
This analysis uses saved docking scores and does not rerun docking.

2. Reproduce functional comparisons and diagnostic analyses
----------------------------------------------------------
python3 "Functional analyses/code/reproduce_analysis.py" --replicates 2000

After preparation, "Functional analyses/sources" contains the six inputs:
master_input.csv; curated_input.csv; surface_candidates_annotation_snapshot.csv;
vina_patch_boxes_20260219_070756.csv;
vina_patch_boxes_20260219_070756_PRI_only.csv; vina_results_ad4_polyP10.csv.
Outputs appear in "Functional analyses/tables", with a summary beside it.
These include the 219 keyword comparisons, NOX/DUOX records, composition
diagnostics and within_vs_between_model_variation.csv for Supplementary
Figure S1. The original script sets its bootstrap seed internally.

3. Rebuild the original master calculation table
-----------------------------------------------
First extract the supplied patch records:

python3 -m tarfile -e "Screening workflow/Inputs/Original patch records.tar.gz" "Screening workflow/Inputs"

Then run this command (shown on one line):

python3 "Screening workflow/final csv/final_csv.py" --alphafold_summary "Screening workflow/Inputs/alphafold_summary.csv.xlsx" --peppatch_dir "Screening workflow/Inputs/patch_records" --vina_csv "Functional analyses/sources/vina_results_ad4_polyP10.csv" --pri_csv "Functional analyses/sources/vina_patch_boxes_20260219_070756_PRI_only.csv" --output_dir "Rebuilt outputs/Master table"

The archive contains 8,678 patch lists and their 8,678 residue lists. The
command rebuilds grand_patch_table_PDI100_polyP10.csv from these records,
the original metadata and saved PRI/docking summaries. All 23,383 rows were
reproduced in testing; differences were limited to floating point rounding.
The separately supplied master workbook is not used as a calculation input.

4. Reconstruct the LPL and LIPI contact tables
-------------------------------------------
python3 "Contact calculations/reconstruct_lpl_lipi_contacts.py" --coordinates "../Candidate structures" --output "Rebuilt outputs/Contact tables"

The script finds the supplied canonical LPL patch001 and canonical LIPI
patch002 model/pose pairs and checks their original hashes. It writes the
Data S9 and Data S11 tables and numerical checks under the output directory.
All 59 manuscript checks passed in testing. These tables are calculated directly from the saved coordinates. No docking, alignment or coordinate modification is performed.

5. Original cluster screening workflow
-------------------------------------
"Screening workflow" contains the original Python and Slurm files for:
PDB2PQR -> inputgen -> APBS -> PEP-Patch -> patch coordinates/PRI -> Meeko
-> AutoGrid4 -> Vina -> final CSV assembly.

The original records specify PDB2PQR 3.7.1, APBS 1.3 and AutoDock Vina 1.2.5.
Additional tools are PROPKA through PDB2PQR, PEP-Patch, Meeko's
mk_prepare_receptor.py, AutoGrid4, GNU Parallel, Bash and Slurm. The patch
coordinate and PRI scripts require NumPy and pandas. The original Vina
Slurm file requests 1.2.6, whereas the saved completed run records report
1.2.5.

The screening scripts were configured for the CAC cluster with Slurm and
project-specific software environments. They retain the original /global/...
paths, environment activation commands, module names and resource settings.
Configure these for the intended computing system. The package uses
descriptive folder names for the original numbered cluster directories.

Inputs includes 4,062 starting UniProt IDs, metadata/download URLs and the
list of 8,808 model filenames. The saved docking box, PRI and Vina CSVs are
shared from "Functional analyses/sources". Candidate structures supplies
128 candidate models and all 19,481 original saved docking poses. A full
screening run uses the listed AlphaFold version 6 models and generates
prepared PQR files, APBS inputs/maps, receptor PDBQT files and AutoGrid maps. Preserve the supplied polyP10 input files and
autogrid/run/boron-silicon-atom_par.dat when configuring those stages.

The commands in sections 1–4 operate on the saved analysis inputs. Vina
was run without a fixed random seed; the supplied saved poses and scores
are the records used for the reported results.

SUPPLEMENTARY LABELS

Supplementary Data S7 identifies the original retrieval, patch, PRI, docking-score, annotation and ligand inputs in this archive. The Supplementary Information file index and Supplementary data/File guide.txt map each filename and path. Scripts and retained source are supplied under Code availability.

RETAINED SOURCE FOR ADDITIONAL TABLES

"Retained source" contains original code for the paired isoform table,
contacts across all 185 high scoring docking results, and model evidence
groups. "Source provenance.json" identifies complete files and verbatim
extracts, their original line ranges, file hashes and required inputs.

The table-processing sources retain the author's original local paths
and input records. Configure input locations and output directories for
the intended working environment. Source provenance.json identifies the
source files, excerpt ranges, original inputs and supplied counterparts.
The commands in sections 1–4 describe the saved data analyses in this package.

Isoform pairs
-------------
build_combined_figure.py contains the original paired-table construction
and correlation-check extracts. Its master CSV is supplied in Supplementary
data, and its saved output matches isoform_pairs_reconstructed.csv exactly.

All selected contacts
---------------------
prepare_inputs.py and calculate_contacts.py.recovered.txt retain the
original all-185 contact calculation and its parser. The saved output
matches contacts_185_4A.csv exactly. The original run also used coordinate
manifests, workbooks, prior contact records and frozen UniProt records.
The provenance file records these input specifications. This source
requires NumPy and openpyxl.

Model evidence groups
---------------------
audit_memberships.py and audit_model_memberships.py retain the original
grouping steps. build_v12.py contains only the input-table and export
extracts. The original exported table matches the supplied model evidence
groups CSV exactly. These steps used frozen workbook records, a candidate
manifest, the consistently selected pose list and intermediate membership
tables. Their original dependencies are listed in the provenance file.

Source provenance.json records source-file hashes, excerpt ranges and
matching saved-table hashes.
