## Supplementary Code for "A structural screen for candidate polyphosphate binding surfaces in human membrane associated proteins": pdb2pqr_array.slurm

```
#!/bin/bash
#SBATCH --job-name=pdb2pqr
#SBATCH --time=01:00:00
#SBATCH --ntasks=1
#SBATCH --cpus-per-task=100
#SBATCH --mem=35G
#SBATCH --account=def-hpcg1856_cpu
#SBATCH --output=%x_%j.out
#SBATCH --error=%x_%j.err

set -euo pipefail

# --- paths ---
IN_DIR=/global/home/hpc6167/ondemand/david_poly_project/0_pdb
OUT_DIR=/global/home/hpc6167/ondemand/david_poly_project/1_pdb2pqr
LIST_FILE="$IN_DIR/pdb_list.txt"
LOG_DIR="$OUT_DIR/logs"

mkdir -p "$OUT_DIR" "$LOG_DIR"

# --- software ---
module load python/3.13.2
source /global/home/hpc6167/ondemand/david_poly_project/0_programs/venvs/pdb2pqr_py313/bin/activate

# GNU Parallel (module name may vary; this tries common names)
module load parallel 2>/dev/null || module load gnu-parallel 2>/dev/null || true

# --- make list if missing ---
if [ ! -f "$LIST_FILE" ]; then
  (cd "$IN_DIR" && ls -1 *.pdb > "$LIST_FILE")
fi

# --- worker function: one PDB -> one PQR (+ optional pdb output) + one log ---
run_one() {
  local pdb="$1"
  local base="${pdb%.pdb}"

  pdb2pqr \
    --ff=PARSE \
    --drop-water \
    --titration-state-method=propka \
    --with-ph 7.4 \
    --whitespace \
    --pdb-output "$OUT_DIR/${base}_pdb2pqr.pdb" \
    "$IN_DIR/$pdb" "$OUT_DIR/${base}.pqr" \
    > "$LOG_DIR/${base}.pdb2pqr.log" 2>&1
}

export -f run_one
export IN_DIR OUT_DIR LOG_DIR

# Run up to 100 structures at once (one per CPU), then refill as they finish
parallel --jobs "${SLURM_CPUS_PER_TASK}" run_one :::: "$LIST_FILE"
```
