## Supplementary Code for "A structural screen for candidate polyphosphate binding surfaces in human membrane associated proteins": inputgen.slurm

```
#!/bin/bash -l
#SBATCH --job-name=inputgen_serial
#SBATCH --time=02:00:00
#SBATCH --ntasks=1
#SBATCH --cpus-per-task=16
#SBATCH --mem-per-cpu=1G
#SBATCH --account=def-hpcg1856_cpu
#SBATCH --output=%x_%j.out
#SBATCH --error=%x_%j.err

set -euo pipefail

ROOT=/global/project/hpcg1856/david_poly_project
PQR_DIR=$ROOT/1_pdb2pqr
OUT_IN_DIR=$ROOT/2_inputgen
OUT_DX_DIR=$ROOT/3_apbs

mkdir -p "$OUT_IN_DIR" "$OUT_DX_DIR"

source $ROOT/0_programs/venvs/pdb2pqr_py313/bin/activate

python3 $ROOT/2_inputgen/inputgen_call.py \
  --pqr-dir "$PQR_DIR" \
  --out-in-dir "$OUT_IN_DIR" \
  --out-dx-dir "$OUT_DX_DIR" \
  --jobs "${SLURM_CPUS_PER_TASK}" \
  --apply-settings \
  --overwrite
```
