## Supplementary Code for "A structural screen for candidate polyphosphate binding surfaces in human membrane associated proteins": inputgen_call.py

```
#!/usr/bin/env python3
"""
Generate APBS .in files from .pqr using pdb2pqr.psize + pdb2pqr.inputgen,
then normalize APBS settings, enforce mg-auto (serial-friendly), and rewrite
'write pot dx' output stems.

This version is for running MANY single-core APBS jobs (job array), not MPI per-structure.
"""

from __future__ import annotations

import argparse
import os
import re
from concurrent.futures import ProcessPoolExecutor, as_completed
from pathlib import Path, PurePosixPath
from typing import Optional, Tuple, List

from pdb2pqr import psize, inputgen


PROJECT_ROOT = Path("/global/project/hpcg1856/david_poly_project")
DEFAULT_PQR_DIR = PROJECT_ROOT / "1_pdb2pqr"
DEFAULT_OUT_IN_DIR = PROJECT_ROOT / "2_inputgen"
DEFAULT_OUT_DX_DIR = PROJECT_ROOT / "3_apbs"

CFAC = 1.7
FADD = 25.0
SPACE = 0.40
GMEMFAC = 200.0
GMEMCEIL = 12000.0
OFRAC = 0.1
REDFAC = 0.25

DEFAULT_METHOD = "mg-auto"
ISTRNG = 0.14
ASYNCFLAG = False
POTDX = True

TARGET = {
    "pbe": "npbe",
    "bcfl": "sdh",
    "pdie": 4.0,
    "sdie": 74.2,
    "srfm": "smol",
    "srad": 1.4,
    "sdens": 15.0,
    "temp": 310.15,
    "chgm": "spl2",
    "calcenergy": "no",
    "calcforce": "no",
}

RE_PBE = re.compile(r"^\s*(lpbe|npbe)\s*$", re.IGNORECASE)
RE_METHOD = re.compile(r"^\s*mg-(auto|para)\s*$", re.IGNORECASE)
RE_PDIME = re.compile(r"^\s*pdime\b.*$", re.IGNORECASE)
RE_WRITE_POT = re.compile(r"^\s*write\s+pot\s+\S+\s+\S+\s*$", re.IGNORECASE)
RE_PRINT_ENERGY = re.compile(r"^\s*print\s+.*energy.*$", re.IGNORECASE)

RE_KV = {
    "bcfl": re.compile(r"^\s*bcfl\b.*$", re.IGNORECASE),
    "pdie": re.compile(r"^\s*pdie\b.*$", re.IGNORECASE),
    "sdie": re.compile(r"^\s*sdie\b.*$", re.IGNORECASE),
    "srfm": re.compile(r"^\s*srfm\b.*$", re.IGNORECASE),
    "srad": re.compile(r"^\s*srad\b.*$", re.IGNORECASE),
    "sdens": re.compile(r"^\s*sdens\b.*$", re.IGNORECASE),
    "temp": re.compile(r"^\s*temp\b.*$", re.IGNORECASE),
    "chgm": re.compile(r"^\s*chgm\b.*$", re.IGNORECASE),
    "calcenergy": re.compile(r"^\s*calcenergy\b.*$", re.IGNORECASE),
    "calcforce": re.compile(r"^\s*calcforce\b.*$", re.IGNORECASE),
}


def _rewrite_mol_pqr_line_abs(in_file: Path, pqr_file: Path) -> None:
    pqr_abs = pqr_file.resolve().as_posix()
    lines = in_file.read_text(encoding="utf-8", errors="replace").splitlines(True)

    out_lines: List[str] = []
    replaced = False
    for line in lines:
        if line.lstrip().startswith("mol pqr ") and not replaced:
            indent = line[: len(line) - len(line.lstrip())]
            out_lines.append(f"{indent}mol pqr {pqr_abs}\n")
            replaced = True
        else:
            out_lines.append(line)

    in_file.write_text("".join(out_lines), encoding="utf-8")


def _apply_settings_writepot_serial(
    in_file: Path,
    out_dx_dir: Optional[PurePosixPath],
) -> None:
    lines = in_file.read_text(encoding="utf-8", errors="replace").splitlines(True)
    new_lines: List[str] = []

    inside_elec = False

    for line in lines:
        s = line.strip().lower()

        if s == "elec":
            inside_elec = True
            new_lines.append(line)
            continue
        if inside_elec and s == "end":
            inside_elec = False
            new_lines.append(line)
            continue

        # remove noisy print energy lines
        if RE_PRINT_ENERGY.match(line):
            continue

        # force npbe
        if RE_PBE.match(line):
            indent = line[: len(line) - len(line.lstrip())]
            new_lines.append(f"{indent}{TARGET['pbe']}\n")
            continue

        # force mg-auto (serial-friendly)
        if inside_elec and RE_METHOD.match(line):
            indent = line[: len(line) - len(line.lstrip())]
            new_lines.append(f"{indent}mg-auto\n")
            continue

        # remove pdime if present (not used for mg-auto serial runs)
        if inside_elec and RE_PDIME.match(line):
            continue

        # normalize key-value lines
        replaced = False
        for k, rx in RE_KV.items():
            if rx.match(line):
                indent = line[: len(line) - len(line.lstrip())]
                v = TARGET[k]
                if isinstance(v, float):
                    if k in ("pdie", "sdie"):
                        new_lines.append(f"{indent}{k} {v:.4f}\n")
                    elif k == "temp":
                        new_lines.append(f"{indent}{k} {v:.2f}\n")
                    else:
                        new_lines.append(f"{indent}{k} {v:.2f}\n")
                else:
                    new_lines.append(f"{indent}{k} {v}\n")
                replaced = True
                break
        if replaced:
            continue

        # rewrite write pot dx output stem to OUT_DX_DIR/<stem>
        if out_dx_dir is not None and RE_WRITE_POT.match(line):
            indent = line[: len(line) - len(line.lstrip())]
            stem = in_file.stem
            out_stem = (out_dx_dir / stem).as_posix()
            new_lines.append(f"{indent}write pot dx {out_stem}\n")
            continue

        new_lines.append(line)

    in_file.write_text("".join(new_lines), encoding="utf-8")


def _process_one(
    pqr_file: str,
    out_in_dir: str,
    overwrite: bool,
    apply_settings: bool,
    out_dx_dir: Optional[str],
) -> Tuple[str, bool, str]:
    pqr_path = Path(pqr_file)
    out_in_dir_p = Path(out_in_dir)
    out_in = out_in_dir_p / f"{pqr_path.stem}.in"

    if out_in.exists() and not overwrite:
        return (pqr_path.name, True, "skip_exists")

    size = psize.Psize(
        cfac=CFAC,
        fadd=FADD,
        space=SPACE,
        gmemfac=GMEMFAC,
        gmemceil=GMEMCEIL,
        ofrac=OFRAC,
        redfac=REDFAC,
    )
    size.run_psize(str(pqr_path))

    apbs_input = inputgen.Input(
        pqrpath=str(pqr_path),
        size=size,
        method=DEFAULT_METHOD,   # mg-auto
        asyncflag=ASYNCFLAG,
        istrng=ISTRNG,
        potdx=POTDX,
    )
    apbs_input.print_input_files(str(out_in))

    _rewrite_mol_pqr_line_abs(out_in, pqr_path)

    if apply_settings:
        out_posix = PurePosixPath(out_dx_dir) if out_dx_dir else None
        _apply_settings_writepot_serial(out_in, out_posix)

    return (pqr_path.name, True, "ok")


def main() -> None:
    ap = argparse.ArgumentParser()
    ap.add_argument("--pqr-dir", default=str(DEFAULT_PQR_DIR))
    ap.add_argument("--out-in-dir", default=str(DEFAULT_OUT_IN_DIR))
    ap.add_argument("--out-dx-dir", default=str(DEFAULT_OUT_DX_DIR))
    ap.add_argument("--jobs", type=int, default=0)
    ap.add_argument("--overwrite", action="store_true")
    ap.add_argument("--apply-settings", action="store_true")
    args = ap.parse_args()

    pqr_dir = Path(args.pqr_dir)
    out_in_dir = Path(args.out_in_dir)
    out_in_dir.mkdir(parents=True, exist_ok=True)

    pqr_files = sorted(pqr_dir.glob("*.pqr"))
    if not pqr_files:
        raise SystemExit(f"No .pqr files found in: {pqr_dir}")

    jobs = args.jobs if args.jobs > 0 else int(os.environ.get("SLURM_CPUS_PER_TASK", "1"))

    print(f"METHOD: {DEFAULT_METHOD}")
    print(f"FILES: {len(pqr_files)}  JOBS(inputgen workers): {jobs}")

    failures: List[Tuple[str, str]] = []
    done = 0

    with ProcessPoolExecutor(max_workers=jobs) as ex:
        futs = [
            ex.submit(
                _process_one,
                str(p),
                str(out_in_dir),
                args.overwrite,
                args.apply_settings,
                args.out_dx_dir,
            )
            for p in pqr_files
        ]
        for fut in as_completed(futs):
            if fut.exception() is not None:
                failures.append(("<unknown>", str(fut.exception())))
            else:
                name, ok, msg = fut.result()
                if not ok:
                    failures.append((name, msg))
            done += 1
            if done % 500 == 0:
                print(f"Processed {done}/{len(pqr_files)}")

    if failures:
        rep = out_in_dir / "inputgen_failures.txt"
        with rep.open("w", encoding="utf-8") as f:
            for name, err in failures:
                f.write(f"{name}\n{err}\n{'-'*60}\n")
        print(f"Done with failures: {len(failures)} -> {rep}")
    else:
        print("Done. No failures.")


if __name__ == "__main__":
    main()
```
