## Supplementary Code for "A structural screen for candidate polyphosphate binding surfaces in human membrane associated proteins": apbs.slurm

```
#!/bin/bash -l
#SBATCH --job-name=apbs_pool
#SBATCH --time=5:50:59
#SBATCH --ntasks=15
#SBATCH --cpus-per-task=1
#SBATCH --mem-per-cpu=120G
#SBATCH --account=def-hpcg1856_cpu
#SBATCH --output=%x_%j.out
#SBATCH --error=%x_%j.err
#SBATCH --exclude=frnt076,frnt135

set -euo pipefail
export OMP_NUM_THREADS=1

module purge || true
module load StdEnv/2020 gcc/9.3.0 openmpi/4.0.3
module load apbs/1.3

APBS_BIN="$(command -v apbs)"
echo "APBS_BIN=$APBS_BIN"

ROOT=/global/project/hpcg1856/david_poly_project
IN_DIR=$ROOT/2_inputgen
OUT_DIR=$ROOT/3_apbs
LOG_DIR=$OUT_DIR/logs
mkdir -p "$LOG_DIR"

mapfile -t INFILES < <(ls -1 "$IN_DIR"/*.in | sort)

# Return 0 if any NON-EMPTY "safe" .dx or .dx.gz exists for this base
dx_exists() {
  local base="$1"
  shopt -s nullglob
  local f

  for f in \
    "$OUT_DIR/${base}.dx" \
    "$OUT_DIR/${base}_"*.dx \
    "$OUT_DIR/${base}-"*.dx \
    "$OUT_DIR/${base}."*.dx \
    "$OUT_DIR/${base}+"*.dx \
    "$OUT_DIR/${base}.dx.gz" \
    "$OUT_DIR/${base}_"*.dx.gz \
    "$OUT_DIR/${base}-"*.dx.gz \
    "$OUT_DIR/${base}."*.dx.gz \
    "$OUT_DIR/${base}+"*.dx.gz
  do
    if [[ -s "$f" ]]; then
      shopt -u nullglob
      return 0
    fi
  done

  shopt -u nullglob
  return 1
}

# Launch up to 500 concurrent APBS runs
for infile in "${INFILES[@]}"; do
  base="$(basename "$infile" .in)"

  # === SKIP LOGIC: ONLY OUTPUT-BASED ===
  if dx_exists "$base"; then
    continue
  fi
  # =====================================

  # write a unique log per run to avoid overwriting old logs
  logfile="$LOG_DIR/${base}.apbs.log"

  srun --exclusive -N1 -n1 "$APBS_BIN" "$infile" >"$logfile" 2>&1 &

  while (( $(jobs -pr | wc -l) >= 15 )); do
   sleep 1
 done

done

wait
echo "All APBS runs finished."
```
