## Supplementary Code for "A structural screen for candidate polyphosphate binding surfaces in human membrane associated proteins": find_coordination.py

```
# -*- coding: utf-8 -*-
import os
import re
import gzip
from pathlib import Path
from datetime import datetime

import numpy as np
import pandas as pd

# =========================
# USER SETTINGS (EDIT THESE)
# =========================
PEPPATCH_OUT_DIR = Path("/global/project/hpcg1856/david_poly_project/4_peppatch/csv")
PDB_DIR          = Path("/global/project/hpcg1856/david_poly_project/0_pdb")
OUT_DIR          = Path("/global/project/hpcg1856/david_poly_project/5_patch_coordination/coordination")

MARGIN_A         = 65.0  # Angstrom
WRITE_PATCH_CONFIGS = False
USE_CA_ONLY         = False
HEAVY_ONLY          = True
SKIP_MULTICHAIN_PDB = True

# =========================
# INTERNALS
# =========================
_RES_ANYWHERE_RE = re.compile(r"([A-Za-z]{3})\s*([-]?\d+)", re.IGNORECASE)

def now_stamp():
    return datetime.now().strftime("%Y%m%d_%H%M%S")

def open_run_log(out_dir: Path) -> Path:
    out_dir.mkdir(parents=True, exist_ok=True)
    log_path = out_dir / f"build_vina_boxes_{now_stamp()}.log"
    with open(log_path, "a", encoding="utf-8") as f:
        f.write(f"[START] {datetime.now().isoformat(timespec='seconds')}\n")
        f.write(f"PEPPATCH_OUT_DIR = {PEPPATCH_OUT_DIR}\n")
        f.write(f"PDB_DIR          = {PDB_DIR}\n")
        f.write(f"OUT_DIR          = {OUT_DIR}\n")
        f.write(f"MARGIN_A         = {MARGIN_A}\n")
        f.write(f"USE_CA_ONLY      = {USE_CA_ONLY}\n")
        f.write(f"HEAVY_ONLY       = {HEAVY_ONLY}\n")
        f.write(f"SKIP_MULTICHAIN_PDB = {SKIP_MULTICHAIN_PDB}\n")
        f.write(f"WRITE_PATCH_CONFIGS = {WRITE_PATCH_CONFIGS}\n\n")
        f.flush()
    return log_path

def log_append(log_path: Path, msg: str):
    with open(log_path, "a", encoding="utf-8") as f:
        f.write(msg.rstrip() + "\n")
        f.flush()

def log_full_list(log_path: Path, header: str, names: list[str]):
    """
    Print full structure name lists into the same log.
    """
    log_append(log_path, f"{header}: {len(names)}")
    if not names:
        return
    # One per line for readability/grepping
    for n in names:
        log_append(log_path, f"  {n}")

def parse_peppatch_tokens(residue_string: str):
    s = str(residue_string)
    for sep in [",", ";", "|", "\t", "\n", "\r"]:
        s = s.replace(sep, " ")
    out = []
    for m in _RES_ANYWHERE_RE.finditer(s):
        res3 = m.group(1).upper()
        resi = int(m.group(2))
        out.append((res3, resi))
    return out

def pick_pdb_path(base_id: str) -> Path | None:
    p1 = PDB_DIR / f"{base_id}.pdb"
    if p1.exists():
        return p1
    p2 = PDB_DIR / f"{base_id}.pdb.gz"
    if p2.exists():
        return p2
    return None

def open_text_maybe_gz(path: Path):
    if path.suffix == ".gz":
        return gzip.open(path, mode="rt", encoding="utf-8", errors="ignore", newline=None)
    return open(path, "r", encoding="utf-8", errors="ignore")

def read_pdb_index(pdb_path: Path, use_ca_only=False, heavy_only=True):
    idx = {}
    chains = set()

    with open_text_maybe_gz(pdb_path) as f:
        for line in f:
            if not (line.startswith("ATOM") or line.startswith("HETATM")):
                continue

            atomname = line[12:16].strip()
            resname  = line[17:20].strip().upper()
            chain    = (line[21].strip() or "?")
            chains.add(chain)

            try:
                resseq = int(line[22:26])
            except ValueError:
                continue

            try:
                x = float(line[30:38]); y = float(line[38:46]); z = float(line[46:54])
            except ValueError:
                continue

            if use_ca_only and atomname != "CA":
                continue
            if heavy_only and atomname.startswith("H"):
                continue

            key = (resname, resseq)
            idx.setdefault(key, []).append((x, y, z))

    for k, v in list(idx.items()):
        idx[k] = np.asarray(v, dtype=float)

    return idx, chains

def compute_patch_box(coords_xyz: np.ndarray, margin_a: float):
    center = coords_xyz.mean(axis=0)
    d = np.linalg.norm(coords_xyz - center, axis=1)
    r_patch = float(d.max())
    half = r_patch + margin_a
    size = np.array([2*half, 2*half, 2*half], dtype=float)
    return center, size, r_patch

def positive_patch_set(patch_list_csv: Path):
    pl = pd.read_csv(patch_list_csv)
    if not {"nr", "type"}.issubset(pl.columns):
        raise ValueError(f"Expected columns ['nr','type'] in {patch_list_csv.name}, got {list(pl.columns)}")
    mask = pl["type"].astype(str).str.strip().str.lower().eq("positive")
    return set(pl.loc[mask, "nr"].astype(int).tolist()), len(pl)

def main():
    OUT_DIR.mkdir(parents=True, exist_ok=True)
    log_path = open_run_log(OUT_DIR)

    combined_csv = OUT_DIR / f"vina_patch_boxes_{now_stamp()}.csv"
    cfg_dir = OUT_DIR / f"vina_patch_box_txt_{now_stamp()}" if WRITE_PATCH_CONFIGS else None
    if cfg_dir:
        cfg_dir.mkdir(parents=True, exist_ok=True)

    residue_csvs = sorted(PEPPATCH_OUT_DIR.glob("*_residue_list.csv"))
    log_append(log_path, f"Found {len(residue_csvs)} residue_list.csv files")

    rows = []
    n_missing_pdb = 0
    n_missing_patchlist = 0
    n_read_fail = 0
    n_skipped_multichain = 0
    n_patch_skipped_not_positive = 0
    n_patch_empty = 0

    # NEW: structure-level counts
    n_struct_has_pos = 0
    n_struct_no_pos = 0
    struct_no_pos = []
    struct_has_pos = []

    # NEW: name lists
    missing_patchlist_structs = []
    missing_pdb_structs = []
    read_fail_patchlist_structs = []
    read_fail_residue_structs = []
    read_fail_pdb_structs = []
    skipped_multichain_structs = []

    for i, res_csv in enumerate(residue_csvs, start=1):
        base = res_csv.name.replace("_residue_list.csv", "")
        patch_csv = PEPPATCH_OUT_DIR / f"{base}_patch_list.csv"

        log_append(log_path, f"[{i}/{len(residue_csvs)}] {base}")

        if not patch_csv.exists():
            n_missing_patchlist += 1
            missing_patchlist_structs.append(base)
            log_append(log_path, f"  ERROR: missing patch_list.csv: {patch_csv}")
            continue

        try:
            pos_set, n_total_patches_in_list = positive_patch_set(patch_csv)
            log_append(log_path, f"  patch_list: total={n_total_patches_in_list}, positive={len(pos_set)}")
        except Exception as e:
            n_read_fail += 1
            read_fail_patchlist_structs.append(base)
            log_append(log_path, f"  ERROR: failed reading patch_list {patch_csv.name}: {repr(e)}")
            continue

        # NEW: record whether structure has any positive patch
        if len(pos_set) == 0:
            n_struct_no_pos += 1
            struct_no_pos.append(base)
        else:
            n_struct_has_pos += 1
            struct_has_pos.append(base)

        pdb_path = pick_pdb_path(base)
        if pdb_path is None:
            n_missing_pdb += 1
            missing_pdb_structs.append(base)
            log_append(log_path, f"  ERROR: missing PDB (.pdb or .pdb.gz) for: {base}")
            continue

        try:
            rl = pd.read_csv(res_csv)
            if not {"nr", "residues"}.issubset(rl.columns):
                n_read_fail += 1
                read_fail_residue_structs.append(base)
                log_append(log_path, f"  ERROR: unexpected columns in {res_csv.name}: {list(rl.columns)}")
                continue
        except Exception as e:
            n_read_fail += 1
            read_fail_residue_structs.append(base)
            log_append(log_path, f"  ERROR: failed reading residue_list {res_csv.name}: {repr(e)}")
            continue

        try:
            pdb_idx, chains = read_pdb_index(pdb_path, use_ca_only=USE_CA_ONLY, heavy_only=HEAVY_ONLY)
        except Exception as e:
            n_read_fail += 1
            read_fail_pdb_structs.append(base)
            log_append(log_path, f"  ERROR: failed reading PDB {pdb_path.name}: {repr(e)}")
            continue

        if SKIP_MULTICHAIN_PDB and len(chains) > 1:
            n_skipped_multichain += 1
            skipped_multichain_structs.append(base)
            log_append(
                log_path,
                f"  WARN: multi-chain PDB detected (chains={sorted(chains)}). Skipping to avoid wrong coordinates."
            )
            continue

        kept = 0
        for _, r in rl.iterrows():
            try:
                patch_nr = int(r["nr"])
            except Exception:
                continue

            if patch_nr not in pos_set:
                n_patch_skipped_not_positive += 1
                continue

            res_keys = parse_peppatch_tokens(r["residues"])
            if not res_keys:
                n_patch_empty += 1
                log_append(log_path, f"  WARN: positive patch {patch_nr} has no parsable residues")
                continue

            coords_list = []
            for key in res_keys:
                if key in pdb_idx:
                    coords_list.append(pdb_idx[key])

            if not coords_list:
                n_patch_empty += 1
                log_append(log_path, f"  WARN: positive patch {patch_nr} residues not found in PDB numbering")
                continue

            coords = np.vstack(coords_list)
            center, size, r_patch = compute_patch_box(coords, MARGIN_A)
            kept += 1

            rows.append({
                "structure_id": base,
                "pdb_path": str(pdb_path),
                "patch_list_csv": str(patch_csv),
                "residue_list_csv": str(res_csv),
                "patch_nr": patch_nr,
                "n_residues": len(res_keys),
                "n_atoms_used": int(coords.shape[0]),
                "center_x": float(center[0]),
                "center_y": float(center[1]),
                "center_z": float(center[2]),
                "size_x": float(size[0]),
                "size_y": float(size[1]),
                "size_z": float(size[2]),
                "R_patch_A": float(r_patch),
                "pad_A": float(MARGIN_A),
            })

            if cfg_dir:
                cfg_path = cfg_dir / f"{base}__patch_{patch_nr:04d}.box.txt"
                with open(cfg_path, "w", encoding="utf-8") as f:
                    f.write(f"center_x = {center[0]:.3f}\n")
                    f.write(f"center_y = {center[1]:.3f}\n")
                    f.write(f"center_z = {center[2]:.3f}\n")
                    f.write(f"size_x = {size[0]:.3f}\n")
                    f.write(f"size_y = {size[1]:.3f}\n")
                    f.write(f"size_z = {size[2]:.3f}\n")

        log_append(log_path, f"  OK: kept {kept} positive patches (others ignored)")

    out_df = pd.DataFrame(rows)
    out_df.to_csv(combined_csv, index=False)

    log_append(log_path, "\n[SUMMARY]")
    log_append(log_path, f"Total residue CSVs:               {len(residue_csvs)}")
    log_append(log_path, f"Structures w/ >=1 positive patch:  {n_struct_has_pos}")
    log_append(log_path, f"Structures w/ 0 positive patches:  {n_struct_no_pos}")

    # Full inline lists in log
    log_full_list(log_path, "STRUCTURES_HAS_POSITIVE_PATCH", struct_has_pos)
    log_full_list(log_path, "STRUCTURES_NO_POSITIVE_PATCH", struct_no_pos)

    log_append(log_path, f"Missing patch_list.csv:           {n_missing_patchlist}")
    log_full_list(log_path, "MISSING_PATCHLIST_STRUCTURES", missing_patchlist_structs)

    log_append(log_path, f"Missing PDBs:                     {n_missing_pdb}")
    log_full_list(log_path, "MISSING_PDB_STRUCTURES", missing_pdb_structs)

    log_append(log_path, f"Read failures (total):            {n_read_fail}")
    log_full_list(log_path, "READ_FAIL_PATCHLIST_STRUCTURES", read_fail_patchlist_structs)
    log_full_list(log_path, "READ_FAIL_RESIDUECSV_STRUCTURES", read_fail_residue_structs)
    log_full_list(log_path, "READ_FAIL_PDB_STRUCTURES", read_fail_pdb_structs)

    log_append(log_path, f"Skipped multi-chain PDBs:          {n_skipped_multichain}")
    log_full_list(log_path, "SKIPPED_MULTICHAIN_STRUCTURES", skipped_multichain_structs)

    log_append(log_path, f"Skipped non-positive patches:      {n_patch_skipped_not_positive}")
    log_append(log_path, f"Positive patches skipped empty:    {n_patch_empty}")
    log_append(log_path, f"Boxes written (positive only):     {len(out_df)}")
    log_append(log_path, f"Combined CSV:                      {combined_csv}")
    if cfg_dir:
        log_append(log_path, f"Box TXT dir:                       {cfg_dir}")
    log_append(log_path, f"[END] {datetime.now().isoformat(timespec='seconds')}")

    print(f"Done. Combined CSV: {combined_csv}")
    print(f"Run log: {log_path}")

if __name__ == "__main__":
    main()
```
