## Supplementary Code for "A structural screen for candidate polyphosphate binding surfaces in human membrane associated proteins": find_coordination.slurm

```
#!/bin/bash
#SBATCH --job-name=find_coord
#SBATCH --output=/global/project/hpcg1856/david_poly_project/5_patch_coordination/logs/%x_%j.out
#SBATCH --error=/global/project/hpcg1856/david_poly_project/5_patch_coordination/logs/%x_%j.err
#SBATCH --time=05:50:00
#SBATCH --cpus-per-task=1
#SBATCH --mem=60G

set -euo pipefail

# Ensure log dir exists
mkdir -p /global/project/hpcg1856/david_poly_project/5_patch_coordination/logs

# Clean module environment, then load Python + SciPy stack (includes numpy/pandas)
module purge
module load StdEnv/2023
module load python scipy-stack

# Sanity check (optional but helpful)
python -c "import numpy, pandas; print('numpy', numpy.__version__, 'pandas', pandas.__version__)"

# Run your script
python /global/project/hpcg1856/david_poly_project/5_patch_coordination/find_coordination.py
```
