## Supplementary Code for "A structural screen for candidate polyphosphate binding surfaces in human membrane associated proteins": PRI_calculation.py

```
# af_patch_PRI_calculation.py
# PRI = max(0, s_mean - s_sd / sqrt(N))
# where s_i is mapped from residue pLDDT x_i by:
#   s_i = 0                  if x_i <= 50
#   s_i = (x_i - 50) / 40    if 50 < x_i < 90
#   s_i = 1                  if x_i >= 90
#
# Heavy-atom inclusion rule for residue-in-box:
#   A residue is included if ANY heavy atom (non-H) from that residue lies inside the Vina box.
#
# This version:
#   - Reads the patch-box CSV as strings
#   - Converts ALL numeric columns to numeric (fixes "numbers stored as text")
#   - Writes a PRI-only CSV (minimal, mergeable by patch_name)

import os
import math
import argparse
from pathlib import Path

import numpy as np
import pandas as pd


# Minimum required columns for PRI computation
REQUIRED_COLS = [
    "structure_id", "pdb_path", "patch_nr",
    "center_x", "center_y", "center_z",
    "size_x", "size_y", "size_z",
]

# Columns that should be numeric if present
NUMERIC_COLS_INT = ["patch_nr", "n_residues", "n_atoms_used"]
NUMERIC_COLS_FLOAT = [
    "center_x", "center_y", "center_z",
    "size_x", "size_y", "size_z",
    "R_patch_A", "pad_A",
]


def resolve_input_csv(path_str: str) -> Path:
    """Allow --boxes to be either a CSV file path or a directory containing a CSV."""
    p = Path(path_str)
    if p.is_file():
        return p
    if p.is_dir():
        csvs = sorted(p.glob("*.csv"))
        if not csvs:
            raise FileNotFoundError(f"No .csv found in directory: {p}")
        return csvs[0]
    raise FileNotFoundError(f"Input path not found: {p}")


def _clean_numeric_series(s: pd.Series) -> pd.Series:
    """
    Clean typical 'number stored as text' issues:
      - leading/trailing whitespace
      - thousands separators like "1,234.5"
      - empty strings -> NaN via to_numeric(..., errors="coerce")
    """
    s = s.astype(str).str.strip()
    s = s.str.replace(",", "", regex=False)
    # Sometimes Excel writes 'nan'/'NaN' as text
    s = s.replace({"": np.nan, "nan": np.nan, "NaN": np.nan, "None": np.nan})
    return pd.to_numeric(s, errors="coerce")


def coerce_numeric_columns(df: pd.DataFrame) -> pd.DataFrame:
    """
    Convert known numeric columns to numeric dtypes (even if stored as text).
    Also: tries to convert ANY other column that looks numeric (high conversion success rate).
    """
    # First: explicitly convert the known numeric columns
    for c in NUMERIC_COLS_FLOAT:
        if c in df.columns:
            df[c] = _clean_numeric_series(df[c]).astype(float)

    for c in NUMERIC_COLS_INT:
        if c in df.columns:
            v = _clean_numeric_series(df[c])
            # patch_nr etc should be integers; allow "1.0" -> 1
            v = v.round()
            df[c] = v.astype("Int64")  # nullable integer

    # Second: optionally convert other numeric-looking columns (safe heuristic)
    # Exclude obvious text/path columns
    exclude = {"structure_id", "pdb_path", "patch_list_csv", "residue_list_csv"}
    for c in df.columns:
        if c in exclude or c in NUMERIC_COLS_FLOAT or c in NUMERIC_COLS_INT:
            continue
        if df[c].dtype != object and not pd.api.types.is_string_dtype(df[c]):
            continue

        # Try numeric conversion; if most values convert, keep it numeric
        v = _clean_numeric_series(df[c])
        non_null = df[c].replace({"": np.nan}).notna().sum()
        if non_null == 0:
            continue
        converted = v.notna().sum()
        # If >= 95% of non-null values became numeric, treat column as numeric
        if converted / non_null >= 0.95:
            df[c] = v.astype(float)

    return df


def is_heavy_atom(atom_name: str, element: str) -> bool:
    """Heavy atom = non-hydrogen. Robust if element field is blank."""
    if element:
        return element.upper() != "H"
    atom_name = atom_name.strip()
    return not atom_name.startswith("H")


def parse_pdb_heavy_atoms_with_plddt(pdb_path: str):
    """
    Parse an AlphaFold-style PDB:
      - residue-level pLDDT is stored in the B-factor field on ATOM lines
      - we compute per-residue pLDDT by averaging B-factors across atoms of that residue (robust)
      - keep heavy atoms for box inclusion

    Returns:
      coords: (n_atoms, 3) float32 heavy-atom coordinates
      atom_res_idx: (n_atoms,) int32 mapping each heavy atom -> residue index
      res_plddt: (n_residues,) float32 residue pLDDT
    """
    pdb_path = str(pdb_path)
    if not os.path.isfile(pdb_path):
        raise FileNotFoundError(pdb_path)

    res_index = {}          # (chain, resseq, icode) -> residue idx
    res_plddt_sum = []
    res_plddt_count = []

    xs, ys, zs = [], [], []
    atom_res_idxs = []

    with open(pdb_path, "r", encoding="utf-8", errors="replace") as f:
        for line in f:
            if not (line.startswith("ATOM") or line.startswith("HETATM")):
                continue

            atom_name = line[12:16].strip()
            chain_id = line[21].strip() or "_"
            resseq_raw = line[22:26].strip()
            icode = line[26].strip() or ""

            try:
                x = float(line[30:38])
                y = float(line[38:46])
                z = float(line[46:54])
            except ValueError:
                continue

            element = line[76:78].strip() if len(line) >= 78 else ""

            # B-factor field (AlphaFold pLDDT per residue copied onto atoms)
            try:
                bfac = float(line[60:66])
            except ValueError:
                bfac = np.nan

            try:
                resseq = int(resseq_raw)
            except ValueError:
                resseq = resseq_raw

            key = (chain_id, resseq, icode)
            if key not in res_index:
                idx = len(res_index)
                res_index[key] = idx
                res_plddt_sum.append(0.0)
                res_plddt_count.append(0)

            idx = res_index[key]

            if not np.isnan(bfac):
                res_plddt_sum[idx] += float(bfac)
                res_plddt_count[idx] += 1

            if is_heavy_atom(atom_name, element):
                xs.append(x)
                ys.append(y)
                zs.append(z)
                atom_res_idxs.append(idx)

    if len(res_index) == 0:
        return (
            np.zeros((0, 3), dtype=np.float32),
            np.zeros((0,), dtype=np.int32),
            np.zeros((0,), dtype=np.float32),
        )

    res_plddt = np.empty((len(res_index),), dtype=np.float32)
    for i in range(len(res_index)):
        if res_plddt_count[i] > 0:
            res_plddt[i] = res_plddt_sum[i] / res_plddt_count[i]
        else:
            res_plddt[i] = np.nan

    coords = np.column_stack([xs, ys, zs]).astype(np.float32, copy=False)
    atom_res_idx = np.array(atom_res_idxs, dtype=np.int32)
    return coords, atom_res_idx, res_plddt


def plddt_to_s(plddt: np.ndarray) -> np.ndarray:
    """Piecewise mapping pLDDT -> s in [0,1]."""
    x = plddt.astype(np.float32)
    s = np.empty_like(x, dtype=np.float32)
    s[x <= 50.0] = 0.0
    s[x >= 90.0] = 1.0
    mid = (x > 50.0) & (x < 90.0)
    s[mid] = (x[mid] - 50.0) / 40.0
    s[np.isnan(x)] = np.nan
    return s


def compute_pri_for_box(coords, atom_res_idx, res_plddt, cx, cy, cz, sx, sy, sz):
    """
    Residue-in-box: include residue if ANY heavy atom from residue is inside the box.
    Box bounds: [c - size/2, c + size/2] for each axis.
    """
    xmin, xmax = cx - sx / 2.0, cx + sx / 2.0
    ymin, ymax = cy - sy / 2.0, cy + sy / 2.0
    zmin, zmax = cz - sz / 2.0, cz + sz / 2.0

    if coords.shape[0] == 0:
        return dict(
            pri=np.nan, pri_status="NO_ATOMS_PARSED",
            pri_n_atoms_in_box=0, pri_n_residues_in_box=0,
            pri_s_mean=np.nan, pri_s_sd=np.nan,
        )

    x, y, z = coords[:, 0], coords[:, 1], coords[:, 2]
    mask = (x >= xmin) & (x <= xmax) & (y >= ymin) & (y <= ymax) & (z >= zmin) & (z <= zmax)
    n_atoms_in_box = int(mask.sum())

    if n_atoms_in_box == 0:
        return dict(
            pri=np.nan, pri_status="NO_RESIDUES_IN_BOX",
            pri_n_atoms_in_box=0, pri_n_residues_in_box=0,
            pri_s_mean=np.nan, pri_s_sd=np.nan,
        )

    res_ids = np.unique(atom_res_idx[mask])
    plddt_sel = res_plddt[res_ids]
    plddt_sel = plddt_sel[~np.isnan(plddt_sel)]
    N = int(plddt_sel.shape[0])

    if N == 0:
        return dict(
            pri=np.nan, pri_status="ONLY_NAN_PLDDT_IN_BOX",
            pri_n_atoms_in_box=n_atoms_in_box, pri_n_residues_in_box=0,
            pri_s_mean=np.nan, pri_s_sd=np.nan,
        )

    s = plddt_to_s(plddt_sel)
    s_mean = float(np.nanmean(s))
    s_sd = float(np.nanstd(s, ddof=1)) if N > 1 else 0.0  # sample SD (N-1)
    pri = max(0.0, s_mean - s_sd / math.sqrt(N))

    return dict(
        pri=pri, pri_status="OK",
        pri_n_atoms_in_box=n_atoms_in_box, pri_n_residues_in_box=N,
        pri_s_mean=s_mean, pri_s_sd=s_sd,
    )


def main():
    ap = argparse.ArgumentParser(description="Compute PRI for Vina patch boxes (AlphaFold PDBs).")
    ap.add_argument("--boxes", required=True, help="Path to patch box CSV file OR directory containing it.")
    ap.add_argument("--out_dir", required=True, help="Output directory for the PRI CSV.")
    ap.add_argument("--pdb_root", default="", help="Fallback root dir if pdb_path is missing.")
    args = ap.parse_args()

    in_csv = resolve_input_csv(args.boxes)
    out_dir = Path(args.out_dir)
    out_dir.mkdir(parents=True, exist_ok=True)

    # Read everything as string first -> then convert numerics (fixes Excel/text issues)
    df = pd.read_csv(in_csv, dtype=str)

    # Strip key text/path columns
    for c in ["structure_id", "pdb_path", "patch_list_csv", "residue_list_csv"]:
        if c in df.columns:
            df[c] = df[c].astype(str).str.strip().str.strip('"')

    # Convert numeric columns reliably (even if stored as text)
    df = coerce_numeric_columns(df)

    # Validate required columns exist
    missing = [c for c in REQUIRED_COLS if c not in df.columns]
    if missing:
        raise ValueError(f"Missing required columns in CSV: {missing}\nFound columns: {list(df.columns)}")

    # Validate required numeric columns converted (no NaN)
    must_be_numeric = ["patch_nr", "center_x", "center_y", "center_z", "size_x", "size_y", "size_z"]
    bad = {c: int(df[c].isna().sum()) for c in must_be_numeric if c in df.columns}
    bad = {k: v for k, v in bad.items() if v > 0}
    if bad:
        raise ValueError(
            "Some required numeric fields could not be converted to numbers (NaN after conversion):\n"
            + "\n".join([f"  {k}: {v} bad rows" for k, v in bad.items()])
        )

    # Build patch_name key
    patch_nr_int = df["patch_nr"].astype("Int64")
    patch_nr_str = patch_nr_int.astype("string").str.zfill(2)
    df["patch_name"] = df["structure_id"].astype(str) + "__patch" + patch_nr_str.astype(str)

    # Results arrays aligned to df rows
    results = {
        "pri": np.full(len(df), np.nan, dtype=float),
        "pri_status": np.array([""] * len(df), dtype=object),
        "pri_n_atoms_in_box": np.zeros(len(df), dtype=int),
        "pri_n_residues_in_box": np.zeros(len(df), dtype=int),
        "pri_s_mean": np.full(len(df), np.nan, dtype=float),
        "pri_s_sd": np.full(len(df), np.nan, dtype=float),
    }

    missing_pdb = 0
    parse_fail = 0

    # Parse each structure once (cache by pdb_path)
    parse_cache = {}

    for structure_id, g in df.groupby("structure_id", sort=False):
        pdb_path = None
        if "pdb_path" in g.columns and g["pdb_path"].notna().any():
            pdb_path = str(g["pdb_path"].dropna().iloc[0]).strip().strip('"')
        elif args.pdb_root:
            pdb_path = str(Path(args.pdb_root) / f"{structure_id}.pdb")

        if not pdb_path or not os.path.isfile(pdb_path):
            missing_pdb += len(g)
            for idx in g.index:
                results["pri_status"][idx] = "PDB_MISSING"
            continue

        if pdb_path in parse_cache:
            coords, atom_res_idx, res_plddt = parse_cache[pdb_path]
        else:
            try:
                coords, atom_res_idx, res_plddt = parse_pdb_heavy_atoms_with_plddt(pdb_path)
                parse_cache[pdb_path] = (coords, atom_res_idx, res_plddt)
            except Exception:
                parse_fail += len(g)
                for idx in g.index:
                    results["pri_status"][idx] = "PDB_PARSE_FAILED"
                continue

        for idx, row in g.iterrows():
            out = compute_pri_for_box(
                coords, atom_res_idx, res_plddt,
                float(row["center_x"]), float(row["center_y"]), float(row["center_z"]),
                float(row["size_x"]), float(row["size_y"]), float(row["size_z"]),
            )
            results["pri"][idx] = out["pri"]
            results["pri_status"][idx] = out["pri_status"]
            results["pri_n_atoms_in_box"][idx] = out["pri_n_atoms_in_box"]
            results["pri_n_residues_in_box"][idx] = out["pri_n_residues_in_box"]
            results["pri_s_mean"][idx] = out["pri_s_mean"]
            results["pri_s_sd"][idx] = out["pri_s_sd"]

    # PRI-only output (no duplicated box columns)
    pri_df = pd.DataFrame({
        "patch_name": df["patch_name"].astype(str).values,
        "pri": results["pri"],
        "pri_status": results["pri_status"],
        "pri_n_atoms_in_box": results["pri_n_atoms_in_box"].astype(int),
        "pri_n_residues_in_box": results["pri_n_residues_in_box"].astype(int),
        "pri_s_mean": results["pri_s_mean"],
        "pri_s_sd": results["pri_s_sd"],
    })

    out_csv = out_dir / f"{in_csv.stem}_PRI_only.csv"
    pri_df.to_csv(out_csv, index=False)

    summary_path = out_dir / f"{in_csv.stem}_PRI_summary.txt"
    with open(summary_path, "w", encoding="utf-8") as f:
        f.write(f"Input CSV: {in_csv}\n")
        f.write(f"Output CSV: {out_csv}\n")
        f.write(f"Rows: {len(df)}\n")
        f.write(f"PDB missing rows: {missing_pdb}\n")
        f.write(f"PDB parse failed rows: {parse_fail}\n")
        f.write("PRI status counts:\n")
        f.write(pri_df["pri_status"].value_counts(dropna=False).to_string())
        f.write("\n")

    print(f"[OK] Wrote: {out_csv}")
    print(f"[OK] Wrote: {summary_path}")


if __name__ == "__main__":
    main()
```
