## Supplementary Code for "A structural screen for candidate polyphosphate binding surfaces in human membrane associated proteins": PRI_calculation.slurm

```
#!/bin/bash
#SBATCH --job-name=PRI
#SBATCH --time=05:00:00
#SBATCH --cpus-per-task=1
#SBATCH --mem=16G
#SBATCH --output=/global/project/hpcg1856/david_poly_project/5_patch_coordination/PRI/pri_%j.out
#SBATCH --error=/global/project/hpcg1856/david_poly_project/5_patch_coordination/PRI/pri_%j.err


set -euo pipefail

module purge
module load python/3.13.2
source /global/project/hpcg1856/david_poly_project/venvs/pri/bin/activate

BOXES="/global/project/hpcg1856/david_poly_project/5_patch_coordination/coordination/vina_patch_boxes_20260219_070756.csv"
OUTDIR="/global/project/hpcg1856/david_poly_project/5_patch_coordination/PRI"
SCRIPT="/global/project/hpcg1856/david_poly_project/5_patch_coordination/PRI_calculation.py"

mkdir -p "$OUTDIR"

python3 "$SCRIPT" --boxes "$BOXES" --out_dir "$OUTDIR"
```
