## Supplementary Code for "A structural screen for candidate polyphosphate binding surfaces in human membrane associated proteins": pep_patch.py

```
# -*- coding: utf-8 -*-
from __future__ import annotations

import gzip
import os
import shutil
import subprocess
import sys
from dataclasses import dataclass
from pathlib import Path
from typing import Optional, List, Tuple

# ============================================================
# PATHS (CAC)
# ============================================================
APBS_DIR = Path("/global/project/hpcg1856/david_poly_project/3_apbs")     # .dx.gz
PDB_DIR  = Path("/global/project/hpcg1856/david_poly_project/0_pdb")      # .pdb

OUT_DIR  = Path("/global/project/hpcg1856/david_poly_project/4_peppatch/csv")
PLY_DIR  = Path("/global/project/hpcg1856/david_poly_project/4_peppatch/ply")

# temp folder INSIDE 4_peppatch
TMP_ROOT = Path("/global/project/hpcg1856/david_poly_project/4_peppatch/_tmp_dx")

PEP_PATCH_EXE = Path("pep_patch_electrostatic")  # relies on PATH inside the conda env "peppatch"

# ============================================================
# PEP-PATCH SETTINGS (KEEP THESE THE SAME)
# ============================================================
PROBE_RADIUS = "0.14"
PATCH_CUTOFF_POS, PATCH_CUTOFF_NEG = "2.7", "-2.7"
INTEGRAL_CUTOFF_POS, INTEGRAL_CUTOFF_NEG = "0.3", "-0.3"
SURFACE_TYPE = "sas"
N_PATCHES = "0"
SIZE_CUTOFF_NM2 = "0.7"
CHECK_CDRS = False

PLY_CMAP = "coolwarm_r"
PLY_CLIM_MIN, PLY_CLIM_MAX = "-6", "6"

# ============================================================
# BEHAVIOR / NAMING
# ============================================================
COPY_BUF = 8 * 1024 * 1024
SKIP_IF_OUTPUT_EXISTS = True

PE0_SUFFIX = "-PE0"
ALLOW_FALLBACK_NO_PE0 = True

# If pep_patch crashes with the known "no patches" bug, we treat it as:
#   patches = 0  (write header-only CSVs, and skip PLY)
NO_PATCH_BUG_MARKER = "ValueError: max() arg is an empty sequence"

# ============================================================
# OUTPUT TEMP ROOTS (NO SUBFOLDERS)
# ============================================================
LOG_DIR = Path("/global/project/hpcg1856/david_poly_project/4_peppatch/logs")
CSV_TMP_DIR = OUT_DIR / "_tmp"
PLY_TMP_DIR = PLY_DIR / "_tmp"
LOG_TMP_DIR = LOG_DIR / "_tmp"


@dataclass
class Job:
    stem: str
    pdb_path: Path
    dx_path: Optional[Path]
    dx_gz_path: Optional[Path]
    needs_decompress: bool


def ensure_dirs() -> None:
    OUT_DIR.mkdir(parents=True, exist_ok=True)
    PLY_DIR.mkdir(parents=True, exist_ok=True)
    LOG_DIR.mkdir(parents=True, exist_ok=True)

    CSV_TMP_DIR.mkdir(parents=True, exist_ok=True)
    PLY_TMP_DIR.mkdir(parents=True, exist_ok=True)
    LOG_TMP_DIR.mkdir(parents=True, exist_ok=True)

    TMP_ROOT.mkdir(parents=True, exist_ok=True)


def outputs_for(stem: str, base_dir: Path) -> Tuple[Path, Path, Path]:
    out_patch = base_dir / f"{stem}_patch_list.csv"
    out_res   = base_dir / f"{stem}_residue_list.csv"
    out_log   = base_dir / f"{stem}_peppatch.log"
    return out_patch, out_res, out_log


def ply_outputs_for(stem: str, base_dir: Path) -> Tuple[Path, Path, Path]:
    ply_pos  = base_dir / f"{stem}-pos.ply"
    ply_neg  = base_dir / f"{stem}-neg.ply"
    ply_log  = base_dir / f"{stem}_peppatch_ply.log"
    return ply_pos, ply_neg, ply_log


def _is_nonempty_file(p: Path) -> bool:
    return p.exists() and p.stat().st_size > 0


def outputs_exist(stem: str) -> bool:
    out_patch, out_res, _ = outputs_for(stem, OUT_DIR)
    # Important: treat 0-byte files as NOT existing (they are usually from crashes)
    return _is_nonempty_file(out_patch) and _is_nonempty_file(out_res)


def ply_outputs_exist(stem: str) -> bool:
    ply_pos, ply_neg, _ = ply_outputs_for(stem, PLY_DIR)
    return _is_nonempty_file(ply_pos) and _is_nonempty_file(ply_neg)


def patch_list_has_any_patches(patch_csv: Path) -> bool:
    """
    True if patch_list.csv has at least 1 data row (not just the header).
    """
    if not _is_nonempty_file(patch_csv):
        return False
    try:
        with patch_csv.open("r", encoding="utf-8", errors="replace") as f:
            header = f.readline()
            if not header:
                return False
            second = f.readline()
            return bool(second and second.strip())
    except Exception:
        return False


def candidate_dx_names(stem: str) -> List[str]:
    names = [f"{stem}{PE0_SUFFIX}.dx", f"{stem}{PE0_SUFFIX}.dx.gz"]
    if ALLOW_FALLBACK_NO_PE0:
        names += [f"{stem}.dx", f"{stem}.dx.gz"]
    return names


def find_dx_for_stem(stem: str) -> Tuple[Optional[Path], Optional[Path]]:
    dx = None
    dx_gz = None

    for name in candidate_dx_names(stem):
        p = APBS_DIR / name
        if p.exists() and name.endswith(".dx") and not name.endswith(".dx.gz"):
            dx = p
            break

    if dx is None:
        for name in candidate_dx_names(stem):
            if name.endswith(".dx.gz"):
                p = APBS_DIR / name
                if p.exists():
                    dx_gz = p
                    break

    return dx, dx_gz


def decompress_gz_to_tmp(dx_gz: Path, out_dx: Path) -> Path:
    # Writes directly under _tmp_dx/<stem>.dx (NO slurm subfolders)
    if out_dx.exists():
        out_dx.unlink()
    with gzip.open(dx_gz, "rb") as src, out_dx.open("wb") as dst:
        shutil.copyfileobj(src, dst, length=COPY_BUF)
    return out_dx


def pep_patch_cmd_csv(pdb: Path, dx: Path, out_csv: Path, out_res_csv: Path) -> List[str]:
    """
    Newer pep_patch_electrostatic signature expects:
      parm trajs... plus --dx, and supports -ro/--resout.

    We pass the PDB twice (parm + one traj) so it behaves like a single-structure run.
    """
    cmd = [
        str(PEP_PATCH_EXE),
        str(pdb),
        str(pdb),
        "--dx", str(dx),
        "--probe_radius", PROBE_RADIUS,
        "-o", str(out_csv),
        "-ro", str(out_res_csv),
        "-c", PATCH_CUTOFF_POS, PATCH_CUTOFF_NEG,
        "-ic", INTEGRAL_CUTOFF_POS, INTEGRAL_CUTOFF_NEG,
        "--surface_type", SURFACE_TYPE,
        "-n", N_PATCHES,
        "-s", SIZE_CUTOFF_NM2,
    ]
    if CHECK_CDRS:
        cmd.append("--check_cdrs")
    return cmd


def pep_patch_cmd_ply(pdb: Path, dx: Path, ply_base: Path) -> List[str]:
    cmd = [
        str(PEP_PATCH_EXE),
        str(pdb),
        str(pdb),
        "--dx", str(dx),
        "--probe_radius", PROBE_RADIUS,
        "--ply_out", str(ply_base),
        "--ply_cmap", PLY_CMAP,
        "--ply_clim", PLY_CLIM_MIN, PLY_CLIM_MAX,
        "-c", PATCH_CUTOFF_POS, PATCH_CUTOFF_NEG,
        "-ic", INTEGRAL_CUTOFF_POS, INTEGRAL_CUTOFF_NEG,
        "--surface_type", SURFACE_TYPE,
        "-n", N_PATCHES,
        "-s", SIZE_CUTOFF_NM2,
    ]
    if CHECK_CDRS:
        cmd.append("--check_cdrs")
    return cmd


def _write_header_only_patch_csv(p: Path) -> None:
    p.parent.mkdir(parents=True, exist_ok=True)
    p.write_text("nr,type,npoints,area,value,main_residue\n", encoding="utf-8")


def _write_header_only_residue_csv(p: Path) -> None:
    p.parent.mkdir(parents=True, exist_ok=True)
    p.write_text("nr,residues\n", encoding="utf-8")


def _log_tail_contains_no_patch_bug(log_path: Path) -> bool:
    try:
        t = log_path.read_text(encoding="utf-8", errors="replace")
    except Exception:
        return False
    return (NO_PATCH_BUG_MARKER in t)


def _move_if_exists(src: Path, dst: Path) -> None:
    if not src.exists():
        return
    dst.parent.mkdir(parents=True, exist_ok=True)
    # overwrite-safe on POSIX (Path.replace)
    src.replace(dst)


def _delete_success_temp_files(stem: str) -> None:
    # Delete only this stem's temp outputs (no subfolders, keep other stems)
    for p in CSV_TMP_DIR.glob(f"{stem}_*"):
        try:
            p.unlink()
        except Exception:
            pass
    for p in PLY_TMP_DIR.glob(f"{stem}*"):
        try:
            p.unlink()
        except Exception:
            pass
    for p in LOG_TMP_DIR.glob(f"{stem}*"):
        try:
            p.unlink()
        except Exception:
            pass


def _killed_by_sigkill(rc: int) -> bool:
    # Common patterns:
    # - 137 = 128 + 9 (SIGKILL) :contentReference[oaicite:1]{index=1}
    # - negative return codes sometimes represent signals (platform-dependent)
    if rc == 137:
        return True
    if rc < 0 and abs(rc) == 9:
        return True
    return False


def run_one(job: Job) -> int:
    # FINAL paths
    out_patch, out_res, out_log = outputs_for(job.stem, OUT_DIR)
    ply_pos, ply_neg, ply_log = ply_outputs_for(job.stem, PLY_DIR)

    # TEMP paths (NO subfolders)
    tmp_patch, tmp_res, tmp_csv_log = outputs_for(job.stem, CSV_TMP_DIR)
    tmp_pos, tmp_neg, tmp_ply_log = ply_outputs_for(job.stem, PLY_TMP_DIR)

    # Skip logic: if final exists and either final PLY exists OR 0 patches, skip.
    if SKIP_IF_OUTPUT_EXISTS and outputs_exist(job.stem):
        if ply_outputs_exist(job.stem) or (not patch_list_has_any_patches(out_patch)):
            return 0

    # DX choice (decompress to _tmp_dx/<stem>.dx if needed)
    dx_to_use = job.dx_path
    made_tmp_dx = False
    tmp_dx_path = TMP_ROOT / f"{job.stem}.dx"

    if dx_to_use is None and job.dx_gz_path is not None:
        dx_to_use = decompress_gz_to_tmp(job.dx_gz_path, tmp_dx_path)
        made_tmp_dx = True

    if dx_to_use is None:
        tmp_csv_log.write_text(f"[ERROR] Missing dx for {job.stem}\n", encoding="utf-8")
        # non-OOM failure => move temp artifacts to final
        _move_if_exists(tmp_csv_log, out_log)
        _delete_success_temp_files(job.stem)
        return 2

    # ------------------------------------------------------------------
    # 1) CSV run (always write to TEMP first)
    # ------------------------------------------------------------------
    for p in (tmp_patch, tmp_res, tmp_csv_log):
        if p.exists():
            try:
                p.unlink()
            except Exception:
                pass

    cmd = pep_patch_cmd_csv(job.pdb_path, dx_to_use, tmp_patch, tmp_res)
    with tmp_csv_log.open("w", encoding="utf-8") as logf:
        logf.write(" ".join(cmd) + "\n")
        logf.flush()
        rc = subprocess.call(cmd, stdout=logf, stderr=subprocess.STDOUT)

    # If pep_patch hit the known "no patches bug", convert it to a valid "0 patches" result
    if rc != 0 and _log_tail_contains_no_patch_bug(tmp_csv_log):
        _write_header_only_patch_csv(tmp_patch)
        _write_header_only_residue_csv(tmp_res)
        rc = 0

    # OOM/SIGKILL-like: KEEP EVERYTHING IN TEMP and DO NOT delete tmp dx
    # (This handles the case where the child dies but Python survives.)
    if _killed_by_sigkill(rc):
        return 137

    # If rc==0 but outputs missing/empty, still treat as "finished" and move whatever we have.
    # Also produce header-only CSVs so you get a deterministic "0 patch" output.
    if rc == 0:
        if not _is_nonempty_file(tmp_patch):
            _write_header_only_patch_csv(tmp_patch)
        if not _is_nonempty_file(tmp_res):
            _write_header_only_residue_csv(tmp_res)

    # Decide whether to run PLY: only if patch_list has >=1 data row
    run_ply = (rc == 0) and patch_list_has_any_patches(tmp_patch)

    # ------------------------------------------------------------------
    # 2) PLY run (TEMP)
    # ------------------------------------------------------------------
    rc2 = None
    if run_ply:
        for p in (tmp_pos, tmp_neg, tmp_ply_log):
            if p.exists():
                try:
                    p.unlink()
                except Exception:
                    pass

        ply_base = PLY_TMP_DIR / job.stem  # pep_patch writes "<base>-pos.ply" and "<base>-neg.ply"
        cmd_ply = pep_patch_cmd_ply(job.pdb_path, dx_to_use, ply_base)
        with tmp_ply_log.open("w", encoding="utf-8") as logf:
            logf.write(" ".join(cmd_ply) + "\n")
            logf.flush()
            rc2 = subprocess.call(cmd_ply, stdout=logf, stderr=subprocess.STDOUT)

        # If PLY got SIGKILL/OOM: KEEP EVERYTHING IN TEMP and DO NOT delete tmp dx
        if rc2 is not None and _killed_by_sigkill(rc2):
            return 137

        # If PLY failed (non-OOM), we still move temp outputs to final (your rule).
        # If outputs are empty, they just won't be moved (exists check).

    # ------------------------------------------------------------------
    # FINALIZE for all non-OOM outcomes (success or other failure):
    # move temp -> final, then delete temp + delete tmp dx (if created)
    # ------------------------------------------------------------------
    _move_if_exists(tmp_patch, out_patch)
    _move_if_exists(tmp_res, out_res)
    _move_if_exists(tmp_csv_log, out_log)

    _move_if_exists(tmp_pos, ply_pos)
    _move_if_exists(tmp_neg, ply_neg)
    _move_if_exists(tmp_ply_log, ply_log)

    # delete decompressed dx ONLY if not OOM/SIGKILL path
    if made_tmp_dx and tmp_dx_path.exists():
        try:
            tmp_dx_path.unlink()
        except Exception:
            pass

    # delete this stem's temp outputs (non-OOM only)
    _delete_success_temp_files(job.stem)

    # Return codes (kept simple)
    if rc != 0:
        return 10
    if rc2 is not None and rc2 != 0:
        return 20
    return 0


def list_jobs() -> List[Job]:
    jobs: List[Job] = []
    for pdb in sorted(PDB_DIR.glob("AF-*-model_v6.pdb")):
        stem = pdb.stem
        dx, dx_gz = find_dx_for_stem(stem)
        needs_decompress = dx is None and dx_gz is not None
        jobs.append(Job(stem=stem, pdb_path=pdb, dx_path=dx, dx_gz_path=dx_gz, needs_decompress=needs_decompress))
    return jobs


def get_global_index() -> int:
    task_id = int(os.environ.get("SLURM_ARRAY_TASK_ID", "1"))
    offset = int(os.environ.get("OFFSET", "0"))
    return task_id + offset


def main() -> None:
    ensure_dirs()
    jobs = list_jobs()
    total = len(jobs)

    gi = get_global_index()  # 1..N
    if gi < 1 or gi > total:
        print(f"[INFO] global_index={gi} out of range 1..{total}. Exiting 0.")
        sys.exit(0)

    job = jobs[gi - 1]
    print(f"global_index={gi}/{total} stem={job.stem}")
    rc = run_one(job)
    sys.exit(rc)


if __name__ == "__main__":
    main()
```
