## Supplementary Code for "A structural screen for candidate polyphosphate binding surfaces in human membrane associated proteins": pep_patch.slurm

```
#!/bin/bash -l
#SBATCH --job-name=peppatch_arr
#SBATCH --time=05:00:00
#SBATCH --ntasks=1
#SBATCH --cpus-per-task=1
#SBATCH --mem-per-cpu=50G
#SBATCH --account=def-hpcg1856_cpu
#SBATCH --output=/global/project/hpcg1856/david_poly_project/4_peppatch/logs/%x_%A_%a.out
#SBATCH --error=/global/project/hpcg1856/david_poly_project/4_peppatch/logs/%x_%A_%a.err

set -euo pipefail

ROOT=/global/project/hpcg1856/david_poly_project
mkdir -p "$ROOT/4_peppatch/logs"

# conda env
export PATH="$ROOT/0_programs/miniforge3/condabin:$ROOT/0_programs/miniforge3/bin:$PATH"
source "$ROOT/0_programs/miniforge3/etc/profile.d/conda.sh"
conda activate peppatch

echo "HOST=$(hostname)"
echo "JOB=$SLURM_JOB_ID ARRAY_TASK=${SLURM_ARRAY_TASK_ID:-NA} OFFSET=${OFFSET:-0}"
which python
python -c "import sys; print(sys.executable)"

python "$ROOT/4_peppatch/pep_patch.py"
```
