## Supplementary Code for "A structural screen for candidate polyphosphate binding surfaces in human membrane associated proteins": autogrid.py

```
#!/usr/bin/env python3
# -*- coding: utf-8 -*-

from __future__ import annotations

import argparse
import datetime as dt
import re
import shutil
import subprocess
import time
from pathlib import Path
from typing import List, Optional, Tuple
from shutil import which

# =============================
# PATHS (CAC)
# =============================
GPF_DIR   = Path("/global/project/hpcg1856/david_poly_project/6_meeko/gpf")
PDBQT_DIR = Path("/global/project/hpcg1856/david_poly_project/6_meeko/pdbqt")

LIGAND_PDBQT = Path("/global/project/hpcg1856/david_poly_project/0_polyP_PDBQT/polyP10/polyP10.pdbqt")

OUT_ROOT = Path("/global/project/hpcg1856/david_poly_project/7_autogrid")
RUN_DIR  = OUT_ROOT / "run"          # where autogrid is executed (cwd)
MAPS_DIR = OUT_ROOT / "maps"         # final maps output
LOGS_DIR = OUT_ROOT / "logs"         # final logs output
USED_GPF_DIR = OUT_ROOT / "used_gpf" # final "effective" GPFs used

AUTOGRID_EXE = "autogrid4"

# =============================
# UTIL
# =============================
def ensure_dir(p: Path) -> None:
    p.mkdir(parents=True, exist_ok=True)

def nonempty_file(p: Path) -> bool:
    try:
        return p.is_file() and p.stat().st_size > 0
    except Exception:
        return False

def read_ligand_atom_types_from_pdbqt(ligand_pdbqt: Path) -> List[str]:
    types: List[str] = []
    seen = set()
    for line in ligand_pdbqt.read_text(encoding="utf-8", errors="replace").splitlines():
        if not (line.startswith("ATOM") or line.startswith("HETATM")):
            continue
        toks = line.split()
        if not toks:
            continue
        t = toks[-1]
        if t not in seen:
            seen.add(t)
            types.append(t)
    return types

def patch_and_struct_id(gpf_stem: str) -> Tuple[str, str]:
    """AF-..._patch001 -> (patch_id, struct_id)."""
    m = re.match(r"^(.*)_patch\d+$", gpf_stem)
    if m:
        return gpf_stem, m.group(1)
    return gpf_stem, gpf_stem

def parse_parameter_file_name(gpf_text: str) -> Optional[str]:
    for line in gpf_text.splitlines():
        s = line.strip()
        if not s or s.startswith("#"):
            continue
        if s.startswith("parameter_file "):
            return s.split(None, 1)[1].strip()
    return None

def find_param_file(param_name: str, candidates: List[Path]) -> Optional[Path]:
    p = Path(param_name)
    if p.is_absolute() and p.is_file():
        return p.resolve()
    for d in candidates:
        cand = d / param_name
        if cand.is_file():
            return cand.resolve()
    return None

def rewrite_gpf_for_polyp(
    src_gpf: Path,
    dst_gpf: Path,
    receptor_pdbqt: Path,
    maps_dir: Path,
    staged_param_filename: str,
    ligand_types: List[str],
) -> None:
    """
    Keep all GPF settings unchanged EXCEPT:
      - parameter_file -> staged_param_filename (no path; file is in cwd)
      - receptor -> absolute receptor pdbqt path
      - gridfld/map/elecmap/dsolvmap -> redirected into maps_dir with patch-unique names
      - ligand_types + map lines -> filtered to only ligand_types (polyP types)
    """
    patch_id = src_gpf.stem
    ligand_set = set(ligand_types)

    lines = src_gpf.read_text(encoding="utf-8", errors="replace").splitlines()
    out_lines: List[str] = []

    for raw in lines:
        s = raw.strip()
        if not s or s.startswith("#"):
            out_lines.append(raw)
            continue

        key = s.split(None, 1)[0]

        if key == "parameter_file":
            out_lines.append(f"parameter_file {staged_param_filename}")
            continue

        if key == "receptor":
            out_lines.append(f"receptor {receptor_pdbqt}")
            continue

        if key == "gridfld":
            out_lines.append(f"gridfld {maps_dir / f'{patch_id}.maps.fld'}")
            continue

        if key == "elecmap":
            out_lines.append(f"elecmap {maps_dir / f'{patch_id}.e.map'}")
            continue

        if key == "dsolvmap":
            out_lines.append(f"dsolvmap {maps_dir / f'{patch_id}.d.map'}")
            continue

        if key == "ligand_types":
            out_lines.append("ligand_types " + " ".join(ligand_types))
            continue

        if key == "map":
            # map <path> where filename ends with .<ATYPE>.map
            # keep only if ATYPE in ligand_types, and redirect output path
            parts = s.split(None, 1)
            if len(parts) == 2:
                old_path = parts[1].strip()
                # infer atom type from filename ...<ATYPE>.map
                name = Path(old_path).name
                atype = "X"
                if name.lower().endswith(".map"):
                    stem = name[:-4]
                    segs = stem.split(".")
                    if len(segs) >= 2:
                        atype = segs[-1]
                if atype in ligand_set:
                    out_lines.append(f"map {maps_dir / f'{patch_id}.{atype}.map'}")
                # else: drop this map line
                continue

        out_lines.append(raw)

    dst_gpf.parent.mkdir(parents=True, exist_ok=True)
    dst_gpf.write_text("\n".join(out_lines) + "\n", encoding="utf-8")

# =============================
# MAIN
# =============================
def main() -> int:
    ap = argparse.ArgumentParser()
    ap.add_argument("--offset", type=int, default=0, help="0-based offset added to SLURM_ARRAY_TASK_ID-1")
    ap.add_argument("--timeout_sec", type=int, default=0, help="0 = no timeout")
    ap.add_argument("--skip_if_outputs_exist", action="store_true", help="skip patch if all expected outputs exist")
    args = ap.parse_args()

    ensure_dir(OUT_ROOT)
    ensure_dir(RUN_DIR)
    ensure_dir(MAPS_DIR)
    ensure_dir(LOGS_DIR)
    ensure_dir(USED_GPF_DIR)

    task_id = int(Path("/proc/self/environ").read_bytes().split(b"SLURM_ARRAY_TASK_ID=")[1].split(b"\0")[0].decode()) \
              if b"SLURM_ARRAY_TASK_ID=" in Path("/proc/self/environ").read_bytes() else 0
    if task_id <= 0:
        raise RuntimeError("SLURM_ARRAY_TASK_ID not set (or <=0). Run via sbatch --array=1-N.")

    gpfs = sorted(GPF_DIR.glob("*_patch*.gpf"))
    if not gpfs:
        raise RuntimeError(f"No patch GPFs found in {GPF_DIR}")

    idx = (task_id - 1) + args.offset
    if idx >= len(gpfs):
        print(f"Task {task_id}: idx={idx} out of range (gpfs={len(gpfs)}). Nothing to do.")
        return 0

    autogrid = which(AUTOGRID_EXE)
    if not autogrid:
        raise RuntimeError("autogrid4 not found in PATH (activate your autogrid env in SLURM).")

    if not LIGAND_PDBQT.is_file():
        raise RuntimeError(f"polyP ligand PDBQT not found: {LIGAND_PDBQT}")

    ligand_types = read_ligand_atom_types_from_pdbqt(LIGAND_PDBQT)
    if not ligand_types:
        raise RuntimeError(f"Could not detect ligand atom types from: {LIGAND_PDBQT}")

    gpf = gpfs[idx]
    patch_id, struct_id = patch_and_struct_id(gpf.stem)
    receptor_pdbqt = PDBQT_DIR / f"{struct_id}.pdbqt"
    if not receptor_pdbqt.is_file():
        raise RuntimeError(f"Missing receptor PDBQT for {struct_id}: {receptor_pdbqt}")

    # param file name from the GPF
    param_name = parse_parameter_file_name(gpf.read_text(encoding="utf-8", errors="replace"))
    if not param_name:
        raise RuntimeError(f"Could not find 'parameter_file ...' in {gpf}")

    # Find param source (either in GPF dir or your old meeko tmp location)
    meeko_tmp = Path("/global/project/hpcg1856/david_poly_project/6_meeko/logs/_tmp_meeko")
    param_src = find_param_file(param_name, [GPF_DIR, meeko_tmp, RUN_DIR])
    if not param_src:
        raise RuntimeError(f"parameter file '{param_name}' not found in {GPF_DIR}, {meeko_tmp}, or {RUN_DIR}")

    # Stage param into RUN_DIR (so parameter_file is resolvable via cwd)
    staged_param = RUN_DIR / Path(param_name).name
    if not staged_param.is_file():
        shutil.copy2(param_src, staged_param)

    # Expected outputs
    fld = MAPS_DIR / f"{patch_id}.maps.fld"
    e_map = MAPS_DIR / f"{patch_id}.e.map"
    d_map = MAPS_DIR / f"{patch_id}.d.map"
    type_maps = [MAPS_DIR / f"{patch_id}.{t}.map" for t in ligand_types]
    glg = LOGS_DIR / f"{patch_id}.glg"
    used_gpf = USED_GPF_DIR / f"{patch_id}.gpf"

    if args.skip_if_outputs_exist:
        if nonempty_file(fld) and nonempty_file(e_map) and nonempty_file(d_map) and all(nonempty_file(p) for p in type_maps) and nonempty_file(glg):
            print(f"{patch_id}: SKIP (outputs exist)")
            return 0

    # Build the effective GPF for polyP maps, pointing outputs into MAPS_DIR
    rewrite_gpf_for_polyp(
        src_gpf=gpf,
        dst_gpf=used_gpf,
        receptor_pdbqt=receptor_pdbqt,
        maps_dir=MAPS_DIR,
        staged_param_filename=staged_param.name,   # IMPORTANT: no path
        ligand_types=ligand_types,
    )

    # Run autogrid
    cmd = [autogrid, "-p", str(used_gpf), "-l", str(glg)]
    print(f"{patch_id}: RUN autogrid4", flush=True)

    t0 = time.perf_counter()
    try:
        if args.timeout_sec > 0:
            proc = subprocess.run(cmd, cwd=str(RUN_DIR), capture_output=True, text=True, timeout=args.timeout_sec)
        else:
            proc = subprocess.run(cmd, cwd=str(RUN_DIR), capture_output=True, text=True)
        rc = proc.returncode
    except subprocess.TimeoutExpired:
        rc = 124

    dt_s = time.perf_counter() - t0
    print(f"{patch_id}: DONE rc={rc} time={dt_s:.2f}s", flush=True)

    # Validate outputs (fail fast for missing)
    missing: List[str] = []
    if rc != 0:
        missing.append(f"autogrid_rc={rc}")

    if not nonempty_file(fld):  missing.append("fld")
    if not nonempty_file(e_map): missing.append("e.map")
    if not nonempty_file(d_map): missing.append("d.map")
    for t, p in zip(ligand_types, type_maps):
        if not nonempty_file(p):
            missing.append(f"{t}.map")
    if not nonempty_file(glg):
        missing.append("glg")

    if missing:
        raise RuntimeError(f"{patch_id}: FAIL missing={', '.join(missing)}")

    print(f"{patch_id}: OK", flush=True)
    return 0

if __name__ == "__main__":
    raise SystemExit(main())
```
