## Supplementary Code for "A structural screen for candidate polyphosphate binding surfaces in human membrane associated proteins": autogrid.slurm

```
#!/bin/bash
#SBATCH --job-name=autogrid
#SBATCH --output=/global/project/hpcg1856/david_poly_project/7_autogrid/slurm_logs/slurm_%A_%a.out
#SBATCH --error=/global/project/hpcg1856/david_poly_project/7_autogrid/slurm_logs/slurm_%A_%a.err
#SBATCH --time=05:50:00
#SBATCH --cpus-per-task=1
#SBATCH --mem-per-cpu=12G

set -euo pipefail

mkdir -p /global/project/hpcg1856/david_poly_project/7_autogrid/slurm_logs
mkdir -p /global/project/hpcg1856/david_poly_project/7_autogrid/run
mkdir -p /global/project/hpcg1856/david_poly_project/7_autogrid/maps
mkdir -p /global/project/hpcg1856/david_poly_project/7_autogrid/logs
mkdir -p /global/project/hpcg1856/david_poly_project/7_autogrid/used_gpf

source /global/software/python/anaconda3-2024.06-1/etc/profile.d/conda.sh
conda activate /global/project/hpcg1856/david_poly_project/0_programs/conda_envs/autogrid_py310

# OFFSET passed at submission time, e.g. OFFSET=2999 sbatch --array=1-2999%600 autogrid.slurm
export OFFSET="${OFFSET:-0}"

python /global/project/hpcg1856/david_poly_project/7_autogrid/autogrid.py \
  --offset "${OFFSET}" \
  --skip_if_outputs_exist
```
