## Supplementary Code for "A structural screen for candidate polyphosphate binding surfaces in human membrane associated proteins": final_csv.py

```
#!/usr/bin/env python3
# -*- coding: utf-8 -*-
from __future__ import annotations

import argparse
import math
import re
from pathlib import Path
from typing import Dict, Optional, Tuple, List

import numpy as np
import pandas as pd


# ----------------------------
# IO
# ----------------------------

def read_table(path: Path) -> pd.DataFrame:
    if not path.exists():
        raise FileNotFoundError(f"Not found: {path}")
    suf = path.suffix.lower()
    if suf in [".xlsx", ".xlsm", ".xls"]:
        return pd.read_excel(path)  # requires openpyxl
    return pd.read_csv(path)


def read_csv_safe(path: Path) -> Optional[pd.DataFrame]:
    """
    Read CSV safely.
    If the file is empty (0 bytes or no header/data), pandas raises EmptyDataError.
    In that case, treat it as missing (return None) and continue.
    """
    try:
        return pd.read_csv(path)
    except pd.errors.EmptyDataError:
        return None
    except (pd.errors.ParserError, UnicodeDecodeError):
        # malformed/encoding issue -> treat as missing
        return None


def safe_float(x) -> float:
    try:
        if pd.isna(x):
            return np.nan
        return float(x)
    except Exception:
        return np.nan


# ----------------------------
# Naming / parsing
# ----------------------------

def parse_entry_id_from_base(base: str) -> str:
    # e.g. AF-O75122-3-F1-model_v6  ->  AF-O75122-3-F1
    return re.sub(r"-model_v\d+$", "", base)


def patch_id_from_base_nr(base: str, nr: int) -> str:
    # e.g. ...-model_v6 + 2 -> ...-model_v6_patch002
    return f"{base}_patch{nr:03d}"


def normalize_pri_patch_name(name: str) -> Optional[str]:
    # PRI example: AF-...-model_v6__patch01  -> AF-...-model_v6_patch001
    if not isinstance(name, str) or not name.strip():
        return None
    m = re.match(r"^(.*?)(?:__patch|_patch)(\d+)$", name.strip())
    if not m:
        return None
    base = m.group(1).rstrip("_")
    try:
        n = int(m.group(2))
    except Exception:
        return None
    return f"{base}_patch{n:03d}"


# ----------------------------
# Patch residue composition (count-based)
# ----------------------------

def residue_percentages(residue_str: str) -> Tuple[float, float, float, float]:
    """
    residue_str example (space separated): "PRO421 THR456 HIS928 ARG931 LYS930 ..."
    Count-based (% of listed residues in that patch).
    Returns %HIS, %ARG, %LYS, %HRK (0-100). Missing -> NaN.
    """
    if not isinstance(residue_str, str) or not residue_str.strip():
        return (np.nan, np.nan, np.nan, np.nan)

    toks = residue_str.split()
    total = len(toks)
    if total == 0:
        return (np.nan, np.nan, np.nan, np.nan)

    c_h = c_r = c_k = 0
    for t in toks:
        aa3 = t[:3].upper()
        if aa3 == "HIS":
            c_h += 1
        elif aa3 == "ARG":
            c_r += 1
        elif aa3 == "LYS":
            c_k += 1

    pct_h = 100.0 * c_h / total
    pct_r = 100.0 * c_r / total
    pct_k = 100.0 * c_k / total
    pct_hrk = 100.0 * (c_h + c_r + c_k) / total
    return (pct_h, pct_r, pct_k, pct_hrk)


# ----------------------------
# 0-1 transforms (robust scale + sigmoid)
# ----------------------------

def robust_z(series: pd.Series) -> pd.Series:
    """
    Robust z-like scaling: (x - median) / IQR, IQR=Q3-Q1.
    Handles all-NaN safely by returning zeros.
    """
    x = series.astype(float)
    arr = x.values
    if np.all(~np.isfinite(arr)):
        return pd.Series(np.zeros(len(x)), index=x.index, dtype=float)

    finite = arr[np.isfinite(arr)]
    med = np.nanmedian(finite)
    q1 = np.nanpercentile(finite, 25)
    q3 = np.nanpercentile(finite, 75)
    iqr = q3 - q1
    if not np.isfinite(iqr) or iqr == 0:
        return pd.Series(np.zeros(len(x)), index=x.index, dtype=float)

    return (x - med) / iqr


def sigmoid(z: pd.Series, scale: float = 1.0) -> pd.Series:
    """
    Logistic sigmoid mapping to (0,1). 'scale' controls steepness.
    """
    if scale <= 0:
        raise ValueError("sigmoid_scale must be > 0")
    zz = (z.astype(float) / scale).clip(-50, 50)
    return 1.0 / (1.0 + np.exp(-zz))


# ----------------------------
# PRI penalty
# ----------------------------

def pri_penalty(pri: float, ref: float, lam: float) -> float:
    """
    Penalty-only:
      PRI >= ref => 1
      PRI < ref  => exp(-lam*(ref - PRI))
    Missing PRI => NaN (kept blank).
    """
    if pri is None or pd.isna(pri):
        return np.nan
    pri = float(pri)
    if pri >= ref:
        return 1.0
    return math.exp(-lam * (ref - pri))


# ----------------------------
# AlphaFold summary dedupe helper
# ----------------------------

def first_nonempty(s: pd.Series):
    """
    Return first non-empty value in a Series; else NaN.
    Used to collapse duplicate Entry_ID rows safely.
    """
    for v in s.tolist():
        if pd.notna(v) and str(v).strip() != "":
            return v
    return np.nan


# ----------------------------
# Main
# ----------------------------

def main() -> None:
    p = argparse.ArgumentParser()

    p.add_argument("--alphafold_summary", type=str,
                   default="/global/project/hpcg1856/david_poly_project/0_pdb/summary_csv/alphafold_summary.csv.xlsx")
    p.add_argument("--peppatch_dir", type=str,
                   default="/global/project/hpcg1856/david_poly_project/4_peppatch/csv")
    p.add_argument("--vina_csv", type=str,
                   default="/global/project/hpcg1856/david_poly_project/8_vina/csv/vina_results_ad4_polyP10.csv")
    p.add_argument("--pri_csv", type=str,
                   default="/global/project/hpcg1856/david_poly_project/5_patch_coordination/PRI/vina_patch_boxes_20260219_070756_PRI_only.csv")
    p.add_argument("--output_dir", type=str,
                   default="/global/project/hpcg1856/david_poly_project/9_final_csv")
    p.add_argument("--out_name", type=str,
                   default="grand_patch_table_PDI100_polyP10.csv")

    # locked math knobs (defaults match your decisions)
    p.add_argument("--sigmoid_scale", type=float, default=1.0)
    p.add_argument("--w_dock", type=float, default=0.6)
    p.add_argument("--w_elec", type=float, default=0.4)
    p.add_argument("--pri_ref", type=float, default=0.75)
    p.add_argument("--pri_lambda", type=float, default=3.0)

    # progress prints
    p.add_argument("--progress_every", type=int, default=250)

    args = p.parse_args()

    if abs((args.w_dock + args.w_elec) - 1.0) > 1e-9:
        raise ValueError("w_dock + w_elec must equal 1.0")

    out_dir = Path(args.output_dir)
    out_dir.mkdir(parents=True, exist_ok=True)

    # ---- AlphaFold summary (UID/Protein name mapping) - DEDUPE Entry_ID
    af = read_table(Path(args.alphafold_summary)).copy()
    for col in ["UniProt_ID", "Entry_ID", "Protein_Name"]:
        if col not in af.columns:
            raise ValueError(f"alphafold_summary missing required column: {col}")

    af = af[["Entry_ID", "UniProt_ID", "Protein_Name"]].copy()
    af = af.dropna(subset=["Entry_ID"])
    af["Entry_ID"] = af["Entry_ID"].astype(str)

    af_dedup = (
        af.groupby("Entry_ID", as_index=False)
          .agg({"UniProt_ID": first_nonempty, "Protein_Name": first_nonempty})
    )
    af_meta = af_dedup.set_index("Entry_ID")[["UniProt_ID", "Protein_Name"]].to_dict(orient="index")

    # ---- Vina AD4 (best_score)
    vina = pd.read_csv(Path(args.vina_csv)).copy()
    for col in ["patch_id", "best_score", "status"]:
        if col not in vina.columns:
            raise ValueError(f"vina CSV missing required column: {col}")

    vina_ok = vina[vina["status"] == "OK"].copy()
    if "scoring" in vina_ok.columns:
        vina_ok = vina_ok[vina_ok["scoring"].astype(str).str.lower() == "ad4"]

    # If multiple entries per patch_id, keep best (most negative) score
    vina_best: Dict[str, float] = (
        vina_ok.groupby("patch_id", as_index=True)["best_score"]
        .min()
        .astype(float)
        .to_dict()
    )

    # ---- PRI (patch_name -> patch_id -> pri)
    pri = pd.read_csv(Path(args.pri_csv)).copy()
    for col in ["patch_name", "pri", "pri_status"]:
        if col not in pri.columns:
            raise ValueError(f"PRI CSV missing required column: {col}")

    pri_ok = pri[pri["pri_status"] == "OK"].copy()
    pri_ok["patch_id"] = pri_ok["patch_name"].map(normalize_pri_patch_name)
    pri_ok = pri_ok[pri_ok["patch_id"].notna()].copy()

    # If multiple entries per patch_id, keep max PRI (most reliable)
    pri_best: Dict[str, float] = (
        pri_ok.groupby("patch_id", as_index=True)["pri"]
        .max()
        .astype(float)
        .to_dict()
    )

    # ---- PEP-Patch: scan all *_patch_list.csv and pair with *_residue_list.csv
    pep_dir = Path(args.peppatch_dir)
    patch_list_files = sorted(pep_dir.glob("*_patch_list.csv"))
    if not patch_list_files:
        raise FileNotFoundError(f"No *_patch_list.csv files found in: {pep_dir}")

    rows: List[dict] = []
    n_files = len(patch_list_files)

    for i, pl_path in enumerate(patch_list_files, start=1):
        base = pl_path.name.replace("_patch_list.csv", "")
        entry_id = parse_entry_id_from_base(base)

        # --- read patch_list safely (empty/malformed => treat as no patches)
        pl = read_csv_safe(pl_path)
        if pl is None:
            # treat as "no patches / failed / nothing above threshold"
            if args.progress_every > 0 and (i % args.progress_every == 0 or i == n_files):
                print(f"[progress] {i}/{n_files} scanned; rows={len(rows)} (skipped empty/malformed patch_list)")
            continue

        # required columns in patch_list
        for col in ["nr", "type", "area", "value"]:
            if col not in pl.columns:
                # malformed => treat like missing for this structure
                pl = None
                break
        if pl is None:
            if args.progress_every > 0 and (i % args.progress_every == 0 or i == n_files):
                print(f"[progress] {i}/{n_files} scanned; rows={len(rows)} (skipped malformed patch_list columns)")
            continue

        # ONLY positive patches
        pl = pl[pl["type"].astype(str).str.lower() == "positive"].copy()
        n_pos = int(pl.shape[0])

        # residue list partner (may be missing/empty/malformed -> treat as missing)
        residue_map: Dict[int, str] = {}
        rl_path = pep_dir / f"{base}_residue_list.csv"
        if rl_path.exists():
            rl = read_csv_safe(rl_path)
            if rl is not None and ("nr" in rl.columns) and ("residues" in rl.columns):
                residue_map = dict(zip(rl["nr"].astype(int), rl["residues"].astype(str)))

        # mapping (may be missing -> left blank)
        meta = af_meta.get(entry_id, {})
        uniprot_id = meta.get("UniProt_ID", np.nan)
        protein_name = meta.get("Protein_Name", np.nan)

        for _, r in pl.iterrows():
            nr = int(r["nr"])
            patch_id = patch_id_from_base_nr(base, nr)

            area = safe_float(r.get("area"))
            value_total = safe_float(r.get("value"))

            value_per_area = np.nan
            if np.isfinite(area) and area != 0 and np.isfinite(value_total):
                value_per_area = value_total / area

            residues = residue_map.get(nr, "")
            pct_h, pct_r, pct_k, pct_hrk = residue_percentages(residues)

            dock_score = vina_best.get(patch_id, np.nan)
            pri_val = pri_best.get(patch_id, np.nan)

            rows.append({
                "UniProt_ID": uniprot_id,
                "Entry_ID": entry_id,
                "Protein_Name": protein_name,
                "n_positive_patches_in_structure": n_pos,
                "patch_id": patch_id,
                "patch_area": area,
                "patch_value_total": value_total,
                "patch_value_per_area": value_per_area,
                "PRI": pri_val,
                "patch_residues": residues,
                "pct_HIS": pct_h,
                "pct_ARG": pct_r,
                "pct_LYS": pct_k,
                "pct_HRK": pct_hrk,
                "dock_best_score_ad4": dock_score,
            })

        if args.progress_every > 0 and (i % args.progress_every == 0 or i == n_files):
            print(f"[progress] {i}/{n_files} patch_list files scanned; rows={len(rows)}")

    df = pd.DataFrame(rows)

    # ---- 0-1 scores
    # docking: more negative is better -> strength = -score (bigger = better)
    dock_strength = -df["dock_best_score_ad4"].astype(float)
    df["Dock01"] = sigmoid(robust_z(dock_strength), scale=args.sigmoid_scale)
    df["Elec01"] = sigmoid(robust_z(df["patch_value_per_area"]), scale=args.sigmoid_scale)

    # keep missing as blank (not forced to 0.5)
    df.loc[df["dock_best_score_ad4"].isna(), "Dock01"] = np.nan
    df.loc[df["patch_value_per_area"].isna(), "Elec01"] = np.nan

    # ---- final PDI-100
    penalty = df["PRI"].apply(lambda x: pri_penalty(x, args.pri_ref, args.pri_lambda))
    base_score = (df["Dock01"] ** args.w_dock) * (df["Elec01"] ** args.w_elec)
    df["PDI100"] = 100.0 * base_score * penalty

    # ---- EXACT output schema (NO extras; NO patch_nr)
    out_cols = [
        "UniProt_ID",
        "Entry_ID",
        "Protein_Name",
        "n_positive_patches_in_structure",
        "patch_id",
        "patch_area",
        "patch_value_total",
        "patch_value_per_area",
        "PRI",
        "patch_residues",
        "pct_HIS",
        "pct_ARG",
        "pct_LYS",
        "pct_HRK",
        "dock_best_score_ad4",
        "Dock01",
        "Elec01",
        "PDI100",
    ]
    df = df[out_cols]

    out_path = out_dir / args.out_name
    df.to_csv(out_path, index=False)
    print(f"[done] wrote: {out_path} (rows={len(df)})")


if __name__ == "__main__":
    main()
```
