## Supplementary Code for "A structural screen for candidate polyphosphate binding surfaces in human membrane associated proteins": final_csv.slurm

```
#!/bin/bash
#SBATCH --job-name=final_csv
#SBATCH --ntasks=1
#SBATCH --cpus-per-task=1
#SBATCH --mem=16G
#SBATCH --time=04:00:00
#SBATCH --output=/global/project/hpcg1856/david_poly_project/9_final_csv/logs/final_csv_%j.out
#SBATCH --error=/global/project/hpcg1856/david_poly_project/9_final_csv/logs/final_csv_%j.err


set -euo pipefail

mkdir -p /global/project/hpcg1856/david_poly_project/9_final_csv/logs

module --force purge
module load StdEnv/2023 python/3.13.2 scipy-stack

source /global/project/hpcg1856/david_poly_project/venv_finalcsv/bin/activate

python -c "import numpy, pandas, openpyxl; print('deps OK')"

python /global/project/hpcg1856/david_poly_project/9_final_csv/final_csv.py
```
