## Supplementary Code for "A structural screen for candidate polyphosphate binding surfaces in human membrane associated proteins": meeko.py

```
#!/usr/bin/env python3
# -*- coding: utf-8 -*-
"""
Batch Meeko receptor preparation + GPF generation per patch box (CAC/SLURM array).

Outputs (under /global/project/hpcg1856/david_poly_project/6_meeko):
- receptor PDBQT per structure
- receptor JSON per structure
- template GPF per structure (in logs/gpf_templates/)
- GPF per patch (in gpf/)
- per-patch logs + per-structure logs + overall summary log

Run inside an environment where mk_prepare_receptor.py is on PATH (your meeko_py310).
"""

from __future__ import annotations

import csv
import math
import os
import re
import sys
import time
import shutil
import subprocess
from dataclasses import dataclass
from pathlib import Path
from typing import List, Optional, Tuple, Dict


# =============================
# CAC SETTINGS (ONLY PATH CHANGES + PQR-ONLY + ARRAY SELECTION)
# =============================

# --- INPUTS ---
PQR_DIR = Path("/global/project/hpcg1856/david_poly_project/1_pdb2pqr")  # <-- EDIT IF NEEDED
CSV_PATH = Path("/global/project/hpcg1856/david_poly_project/5_patch_coordination/coordination/vina_patch_boxes_20260219_070756.csv")

# --- OUTPUT ROOT ---
OUT_ROOT = Path("/global/project/hpcg1856/david_poly_project/6_meeko")

OUT_PDBQT_DIR = OUT_ROOT / "pdbqt"
OUT_JSON_DIR  = OUT_ROOT / "json"
OUT_GPF_DIR   = OUT_ROOT / "gpf"
OUT_LOG_DIR   = OUT_ROOT / "logs"

MEEKO_CMD = "mk_prepare_receptor.py"

SIZE_PAD_PER_AXIS_A = 0

SKIP_PATCH_IF_GPF_EXISTS = True
SKIP_STRUCTURE_IF_PDBQT_JSON_EXIST = True

DEFAULT_ALTLOC = "A"
USE_ALLOW_BAD_RES = True
CHARGE_MODEL = "gasteiger"

CACHE_TEMPLATES_FILENAME = "meeko_templates_cache.json"

GPF_SPACING_OVERRIDE: Optional[float] = 0.5


# =============================
# Helpers
# =============================

def ensure_csv_path(csv_path: Path) -> Path:
    if csv_path.exists():
        return csv_path
    if csv_path.suffix.lower() != ".csv":
        candidate = csv_path.with_suffix(".csv")
        if candidate.exists():
            return candidate
    raise FileNotFoundError(f"CSV not found: {csv_path}")


def find_pqr_for_structure(pqr_dir: Path, structure_id: str) -> Optional[Path]:
    """
    CAC version: ONLY accept .pqr (no .pdb fallback).
    Prefers exact match <structure_id>.pqr, otherwise glob.
    """
    exact = pqr_dir / f"{structure_id}.pqr"
    if exact.exists():
        return exact

    hits = sorted(pqr_dir.glob(f"{structure_id}*.pqr"))
    if hits:
        return hits[0]
    return None


@dataclass(frozen=True)
class PatchJob:
    structure_id: str
    patch_nr: int
    center: Tuple[float, float, float]
    size: Tuple[float, float, float]


def read_patch_jobs(csv_path: Path) -> List[PatchJob]:
    jobs: List[PatchJob] = []
    with csv_path.open(newline="", encoding="utf-8") as f:
        reader = csv.DictReader(f)
        required = {"structure_id", "patch_nr", "center_x", "center_y", "center_z", "size_x", "size_y", "size_z"}
        missing = required - set(reader.fieldnames or [])
        if missing:
            raise ValueError(f"CSV missing required columns: {sorted(missing)}")

        for row in reader:
            sid = row["structure_id"].strip()
            patch_nr = int(float(row["patch_nr"]))
            cx = float(row["center_x"]); cy = float(row["center_y"]); cz = float(row["center_z"])
            sx = float(row["size_x"]) + SIZE_PAD_PER_AXIS_A
            sy = float(row["size_y"]) + SIZE_PAD_PER_AXIS_A
            sz = float(row["size_z"]) + SIZE_PAD_PER_AXIS_A
            jobs.append(PatchJob(sid, patch_nr, (cx, cy, cz), (sx, sy, sz)))
    return jobs


def run_cmd(cmd: List[str], log_path: Path) -> Tuple[int, float]:
    t0 = time.perf_counter()
    proc = subprocess.run(cmd, capture_output=True, text=True)
    dt = time.perf_counter() - t0

    log_path.parent.mkdir(parents=True, exist_ok=True)
    with log_path.open("w", encoding="utf-8") as lf:
        lf.write("COMMAND:\n")
        lf.write(" ".join(cmd) + "\n\n")
        lf.write(f"RETURN_CODE: {proc.returncode}\n")
        lf.write(f"RUNTIME_SECONDS: {dt:.2f}\n\n")
        lf.write("STDOUT:\n")
        lf.write(proc.stdout or "")
        lf.write("\n\nSTDERR:\n")
        lf.write(proc.stderr or "")
        lf.write("\n")

    return proc.returncode, dt


def parse_spacing_from_gpf(gpf_text: str) -> float:
    for line in gpf_text.splitlines():
        if line.strip().startswith("spacing "):
            return float(line.strip().split()[1])
    raise ValueError("Could not find 'spacing <float>' in template GPF")


def even_npts_from_size(size_a: float, spacing: float) -> int:
    raw = int(math.ceil(size_a / spacing))
    if raw % 2 == 1:
        raw += 1
    return max(raw, 2)


def write_patch_gpf_from_template(
    template_gpf: Path,
    out_gpf: Path,
    receptor_pdbqt: Path,
    patch_center: Tuple[float, float, float],
    patch_size: Tuple[float, float, float],
    new_base: Path,
) -> None:
    txt = template_gpf.read_text(encoding="utf-8", errors="replace")

    template_spacing = parse_spacing_from_gpf(txt)
    spacing = GPF_SPACING_OVERRIDE if GPF_SPACING_OVERRIDE is not None else template_spacing

    cx, cy, cz = patch_center
    sx, sy, sz = patch_size
    nx = even_npts_from_size(sx, spacing)
    ny = even_npts_from_size(sy, spacing)
    nz = even_npts_from_size(sz, spacing)

    lines = txt.splitlines()
    out_lines: List[str] = []

    for line in lines:
        s = line.strip()
        if s.startswith("receptor "):
            out_lines.append(f"receptor {receptor_pdbqt}")
        elif s.startswith("gridcenter "):
            out_lines.append(f"gridcenter {cx:.6f} {cy:.6f} {cz:.6f}")
        elif s.startswith("npts "):
            out_lines.append(f"npts {nx} {ny} {nz}")
        elif s.startswith("spacing ") and GPF_SPACING_OVERRIDE is not None:
            out_lines.append(f"spacing {spacing}")
        elif s.startswith("gridfld "):
            out_lines.append(f"gridfld {new_base}.maps.fld")
        elif s.startswith("elecmap "):
            out_lines.append(f"elecmap {new_base}.e.map")
        elif s.startswith("dsolvmap "):
            out_lines.append(f"dsolvmap {new_base}.d.map")
        elif s.startswith("map "):
            parts = s.split()
            if len(parts) >= 2:
                fn = parts[1]
                m = re.search(r"\.([A-Za-z0-9]+)\.map$", fn)
                if m:
                    atype = m.group(1)
                    out_lines.append(f"map {new_base}.{atype}.map")
                else:
                    out_lines.append(f"map {new_base}.X.map")
            else:
                out_lines.append(line)
        else:
            out_lines.append(line)

    out_gpf.parent.mkdir(parents=True, exist_ok=True)
    out_gpf.write_text("\n".join(out_lines) + "\n", encoding="utf-8")


# =============================
# Main: one structure per array task
# =============================
def main() -> int:
    pqr_dir = PQR_DIR
    csv_path = ensure_csv_path(CSV_PATH)

    out_pdbqt_dir = OUT_PDBQT_DIR
    out_json_dir  = OUT_JSON_DIR
    out_gpf_dir   = OUT_GPF_DIR
    out_log_dir   = OUT_LOG_DIR

    for d in [out_pdbqt_dir, out_json_dir, out_gpf_dir, out_log_dir]:
        d.mkdir(parents=True, exist_ok=True)

    # === ONLY CHANGE (1): get GLOBAL_ID from argv; fallback to SLURM_ARRAY_TASK_ID ===
    if len(sys.argv) >= 2:
        global_id = int(sys.argv[1])
    else:
        global_id = int(os.environ.get("SLURM_ARRAY_TASK_ID", "0") or "0")

    if global_id <= 0:
        raise RuntimeError("GLOBAL_ID not provided (argv[1]) and SLURM_ARRAY_TASK_ID not set (or <=0).")

    # === ONLY CHANGE (2): use GLOBAL_ID in cache filename (avoid collisions across multiple array submissions) ===
    cache_file = out_log_dir / f"meeko_templates_cache_task{global_id:06d}.json"

    tmp_dir = out_log_dir / "_tmp_meeko"
    tmp_dir.mkdir(parents=True, exist_ok=True)

    templates_dir = out_log_dir / "gpf_templates"
    templates_dir.mkdir(parents=True, exist_ok=True)

    jobs = read_patch_jobs(csv_path)
    total_patches = len(jobs)
    if total_patches == 0:
        print("No patch rows found in CSV; exiting.")
        return 0

    # group patches by structure
    jobs_by_sid: Dict[str, List[PatchJob]] = {}
    for j in jobs:
        jobs_by_sid.setdefault(j.structure_id, []).append(j)
    for sid in jobs_by_sid:
        jobs_by_sid[sid].sort(key=lambda x: x.patch_nr)

    structures = sorted(jobs_by_sid.keys())

    # === ONLY CHANGE (3): select structure by GLOBAL_ID instead of array task id ===
    idx = global_id - 1
    if idx >= len(structures):
        print(f"GLOBAL_ID {global_id} out of range: structures={len(structures)}; nothing to do.")
        return 0

    sid = structures[idx]
    sid_jobs = jobs_by_sid[sid]

    overall_t0 = time.perf_counter()
    overall_log = out_log_dir / f"meeko_overall_{sid}_{time.strftime('%Y%m%d_%H%M%S')}.log"

    prep_ok = prep_fail = 0
    print(f"[1/2] Preparing receptor PDBQT+JSON (+template GPF) for 1 structure: {sid}")

    pdbqt_target = out_pdbqt_dir / f"{sid}.pdbqt"
    json_target  = out_json_dir  / f"{sid}.json"
    template_gpf_target = templates_dir / f"{sid}.gpf"

    if SKIP_STRUCTURE_IF_PDBQT_JSON_EXIST and pdbqt_target.exists() and json_target.exists() and template_gpf_target.exists():
        print(f"[prep] {sid} . SKIP (already has pdbqt+json+template gpf)")
    else:
        pqr_file = find_pqr_for_structure(pqr_dir, sid)
        if pqr_file is None:
            prep_fail += 1
            (out_log_dir / "missing_pqr_structures.txt").open("a", encoding="utf-8").write(f"{sid}\n")
            print(f"[prep] {sid} . MISSING PQR")
        else:
            first_patch = sid_jobs[0]
            cx, cy, cz = first_patch.center
            sx, sy, sz = first_patch.size

            base = tmp_dir / sid
            cmd = [
                MEEKO_CMD,
                "--read_pqr", str(pqr_file),
                "--output_basename", str(base),
                "--write_pdbqt",
                "--write_json",
                "--write_gpf",
                "--box_center", f"{cx:.6f}", f"{cy:.6f}", f"{cz:.6f}",
                "--box_size",   f"{sx:.6f}", f"{sy:.6f}", f"{sz:.6f}",
                "--charge_model", CHARGE_MODEL,
                "--default_altloc", DEFAULT_ALTLOC,
                "--cache_templates", str(cache_file),
            ]
            if USE_ALLOW_BAD_RES:
                cmd.append("--allow_bad_res")

            struct_log = out_log_dir / "structure_prep_logs" / f"{sid}.log"
            rc, dt = run_cmd(cmd, struct_log)

            if rc != 0:
                prep_fail += 1
                print(f"[prep] {sid} . FAIL ({dt:.2f}s)")
            else:
                moved_pdbqt = moved_json = moved_gpf = False

                pdbqt_files = sorted(tmp_dir.glob(f"{sid}*.pdbqt"))
                if pdbqt_files:
                    shutil.move(str(pdbqt_files[0]), str(pdbqt_target))
                    moved_pdbqt = True

                json_files = sorted(tmp_dir.glob(f"{sid}*.json"))
                if json_files:
                    shutil.move(str(json_files[0]), str(json_target))
                    moved_json = True

                gpf_files = sorted(tmp_dir.glob(f"{sid}*.gpf"))
                if gpf_files:
                    shutil.move(str(gpf_files[0]), str(template_gpf_target))
                    moved_gpf = True

                for fp in tmp_dir.glob(f"{sid}*"):
                    try:
                        fp.unlink()
                    except Exception:
                        pass

                if moved_pdbqt and moved_json and moved_gpf:
                    prep_ok += 1
                    print(f"[prep] {sid} . OK ({dt:.2f}s)")
                else:
                    prep_fail += 1
                    print(f"[prep] {sid} . INCOMPLETE OUTPUTS ({dt:.2f}s)")

    has_pdbqt = pdbqt_target.exists()
    has_tgpf = template_gpf_target.exists()
    print("\n[STEP1 CHECK] File availability after preparation:")
    print(f"  Structure: {sid}")
    print(f"  PDBQT present:        {int(has_pdbqt)}")
    print(f"  Template GPF present: {int(has_tgpf)}")
    print(f"  Both present (eligible for patch GPF): {int(has_pdbqt and has_tgpf)}")

    gpf_ok = 0
    gpf_edit_fail = 0
    gpf_skip = 0

    inel_missing_pdbqt = 0
    inel_missing_template = 0
    inel_missing_both = 0

    patch_file_edit_time = 0.0

    eligible_jobs: List[PatchJob] = []
    ineligible_jobs: List[PatchJob] = []

    for job in sid_jobs:
        template_gpf = template_gpf_target
        receptor_pdbqt = pdbqt_target
        has_t = template_gpf.exists()
        has_r = receptor_pdbqt.exists()

        if has_t and has_r:
            eligible_jobs.append(job)
        else:
            ineligible_jobs.append(job)
            if (not has_t) and (not has_r):
                inel_missing_both += 1
            elif not has_r:
                inel_missing_pdbqt += 1
            else:
                inel_missing_template += 1

    eligible_total = len(eligible_jobs)
    print(f"\n[2/2] Generating patch GPF files from templates for {sid}:")
    print(f"  Total patch rows for structure: {len(sid_jobs)}")
    print(f"  Eligible patches (have template GPF + receptor PDBQT): {eligible_total}")
    print(f"  Ineligible patches: {len(ineligible_jobs)} "
          f"(missing_pdbqt={inel_missing_pdbqt}, missing_template={inel_missing_template}, missing_both={inel_missing_both})")

    for job in ineligible_jobs:
        patch_id = f"{sid}_patch{job.patch_nr:03d}"
        patch_log = out_log_dir / "patch_logs" / f"{patch_id}.log"
        msg = []
        if not pdbqt_target.exists():
            msg.append("MISSING receptor PDBQT")
        if not template_gpf_target.exists():
            msg.append("MISSING template GPF")
        patch_log.parent.mkdir(parents=True, exist_ok=True)
        patch_log.write_text("\n".join(msg) + "\n", encoding="utf-8")

    for k, job in enumerate(eligible_jobs, start=1):
        patch_id = f"{sid}_patch{job.patch_nr:03d}"
        out_gpf = out_gpf_dir / f"{patch_id}.gpf"

        if SKIP_PATCH_IF_GPF_EXISTS and out_gpf.exists():
            gpf_skip += 1
            continue

        patch_log = out_log_dir / "patch_logs" / f"{patch_id}.log"
        try:
            t0 = time.perf_counter()
            new_base = out_gpf_dir / patch_id
            write_patch_gpf_from_template(
                template_gpf=template_gpf_target,
                out_gpf=out_gpf,
                receptor_pdbqt=pdbqt_target,
                patch_center=job.center,
                patch_size=job.size,
                new_base=new_base,
            )
            patch_file_edit_time += (time.perf_counter() - t0)

            patch_log.parent.mkdir(parents=True, exist_ok=True)
            patch_log.write_text("OK\n", encoding="utf-8")
            gpf_ok += 1
        except Exception as e:
            patch_log.parent.mkdir(parents=True, exist_ok=True)
            patch_log.write_text(f"FAIL_EDIT: {repr(e)}\n", encoding="utf-8")
            gpf_edit_fail += 1

    overall_dt = time.perf_counter() - overall_t0

    overall_log.parent.mkdir(parents=True, exist_ok=True)
    with overall_log.open("w", encoding="utf-8") as f:
        f.write("=== MEEKO OVERALL SUMMARY ===\n")
        f.write(f"CSV: {csv_path}\n")
        f.write(f"PQR_DIR: {pqr_dir}\n")
        f.write(f"STRUCTURE: {sid}\n\n")

        f.write("STEP1 (receptor prep):\n")
        f.write(f"  OK={prep_ok}\n")
        f.write(f"  FAIL={prep_fail}\n\n")

        f.write("[STEP1 CHECK]\n")
        f.write(f"  PDBQT_present={int(pdbqt_target.exists())}\n")
        f.write(f"  TemplateGPF_present={int(template_gpf_target.exists())}\n")
        f.write(f"  Both_present={int(pdbqt_target.exists() and template_gpf_target.exists())}\n\n")

        f.write("PATCH GPF ELIGIBILITY:\n")
        f.write(f"  TOTAL_PATCH_ROWS_FOR_STRUCTURE={len(sid_jobs)}\n")
        f.write(f"  ELIGIBLE_PATCHES={eligible_total}\n")
        f.write(f"  INELIGIBLE_PATCHES={len(ineligible_jobs)}\n")
        f.write(f"    INELIGIBLE_missing_pdbqt={inel_missing_pdbqt}\n")
        f.write(f"    INELIGIBLE_missing_template={inel_missing_template}\n")
        f.write(f"    INELIGIBLE_missing_both={inel_missing_both}\n\n")

        f.write("PATCH GPF RESULTS (eligible patches only):\n")
        f.write(f"  OK={gpf_ok}\n")
        f.write(f"  FAIL_EDIT={gpf_edit_fail}\n")
        f.write(f"  SKIP_EXISTING={gpf_skip}\n\n")

        f.write(f"TOTAL RUNTIME (s): {overall_dt:.2f}\n")
        f.write(f"PATCH FILE-EDIT TIME (s): {patch_file_edit_time:.2f}\n")

    print("\n=== SUMMARY ===")
    print(f"Structure: {sid}")
    print(f"Structure prep: OK={prep_ok} FAIL={prep_fail}")
    print("Patch GPF eligibility:",
          f"eligible={eligible_total}/{len(sid_jobs)}, "
          f"ineligible={len(ineligible_jobs)} (missing_pdbqt={inel_missing_pdbqt}, missing_template={inel_missing_template}, missing_both={inel_missing_both})")
    print("Patch GPF results (eligible only):",
          f"OK={gpf_ok} FAIL_EDIT={gpf_edit_fail} SKIP={gpf_skip}")
    print(f"Total runtime:  {overall_dt/60.0:.2f} min")
    print(f"Overall log:    {overall_log}")

    return 0


if __name__ == "__main__":
    raise SystemExit(main())
```
