## Supplementary Code for "A structural screen for candidate polyphosphate binding surfaces in human membrane associated proteins": meeko.slurm

```
#!/bin/bash
#SBATCH --job-name=meeko
#SBATCH --output=/global/project/hpcg1856/david_poly_project/6_meeko/logs/session_logs/slurm_%A_%a.out
#SBATCH --error=/global/project/hpcg1856/david_poly_project/6_meeko/logs/session_logs/slurm_%A_%a.err
#SBATCH --time=04:00:00
#SBATCH --cpus-per-task=1
#SBATCH --mem-per-cpu=30G

set -euo pipefail

mkdir -p /global/project/hpcg1856/david_poly_project/6_meeko/logs/session_logs

source "$(conda info --base)/etc/profile.d/conda.sh"
conda activate /global/project/hpcg1856/david_poly_project/0_programs/conda_envs/meeko_py310

# === ONLY CHANGE: OFFSET comes from script argument; compute GLOBAL_ID and pass to Python ===
OFFSET="${1:-0}"
GLOBAL_ID=$((OFFSET + SLURM_ARRAY_TASK_ID))   # 1-based global index

echo "JOB=$SLURM_JOB_ID TASK=$SLURM_ARRAY_TASK_ID OFFSET=$OFFSET GLOBAL_ID=$GLOBAL_ID"

python /global/project/hpcg1856/david_poly_project/6_meeko/meeko.py "$GLOBAL_ID"
```
