## Supplementary Code for "A structural screen for candidate polyphosphate binding surfaces in human membrane associated proteins": vina.py

```
#!/usr/bin/env python3
# -*- coding: utf-8 -*-

"""
CAC/Linux AutoDock Vina runner (AD4 scoring, map-based docking) for per-patch AutoGrid map sets.

Behavior preserved from your local script:
- One patch = one CSV row; append continuously.
- CSV row written ONLY after docking is completely finished:
    rc==0 AND output pose exists AND score parsed.
- Skip logic: skip ONLY patches already in CSV with status OK/RECOVERED.
- Per-patch logs + master log.
- RECOVER: if output pose exists but CSV missing, parse score and write RECOVERED row.
- CSV contains NO PATH columns.

Output layout (your requirement):
/global/project/hpcg1856/david_poly_project/8_vina/
    csv/   (no subfolders)
    log/   (no subfolders)
    pdbqt/ (no subfolders)
"""

import argparse
import csv
import datetime as dt
import fcntl
import os
import re
import shutil
import subprocess
import sys
from contextlib import contextmanager
from typing import Dict, List, Optional, Tuple


# =========================
# USER CONFIG (EDIT HERE)
# =========================

# Ligand (given by you)
LIGAND_PATH = "/global/project/hpcg1856/david_poly_project/0_polyP_PDBQT/polyP10/polyP10.pdbqt"

# Map directory: must contain *.maps.fld and corresponding *.map/*.xyz/*.fld files
# Set this to your AutoGrid output folder on CAC.
MAPS_DIR = "/global/project/hpcg1856/david_poly_project/7_autogrid/maps"  # <-- EDIT if needed

# Output root (given by you) + fixed 3 subfolders
OUT_ROOT = "/global/project/hpcg1856/david_poly_project/8_vina"
OUT_PDBQT_DIRNAME = "pdbqt"
OUT_LOGS_DIRNAME  = "log"
OUT_CSV_DIRNAME   = "csv"

# AutoDock Vina settings
SCORING = "ad4"
CPU_THREADS = 10
EXHAUSTIVENESS = 16
NUM_MODES = 1
VERBOSITY = 1
SEED: Optional[int] = None

# Validate files referenced by .maps.fld before docking (keep)
VALIDATE_MAPS_FLD_REFERENCES = True

LIGAND_TAG = "polyP10"

# =========================
# END USER CONFIG
# =========================


@contextmanager
def locked_open(path: str, mode: str):
    """
    Exclusive lock for safe concurrent appends (SLURM array % concurrency).
    Does NOT change content/logic, only prevents CSV/log corruption.
    """
    os.makedirs(os.path.dirname(path), exist_ok=True)
    with open(path, mode, encoding="utf-8", newline="") as f:
        fcntl.flock(f.fileno(), fcntl.LOCK_EX)
        try:
            yield f
        finally:
            fcntl.flock(f.fileno(), fcntl.LOCK_UN)


def now_ts() -> str:
    return dt.datetime.now().strftime("%Y-%m-%d %H:%M:%S")


def run_id() -> str:
    return dt.datetime.now().strftime("%Y%m%d_%H%M%S")


def ensure_dirs() -> Tuple[str, str, str, str, str]:
    out_pdbqt = os.path.join(OUT_ROOT, OUT_PDBQT_DIRNAME)
    out_logs  = os.path.join(OUT_ROOT, OUT_LOGS_DIRNAME)
    out_csv   = os.path.join(OUT_ROOT, OUT_CSV_DIRNAME)

    os.makedirs(out_pdbqt, exist_ok=True)
    os.makedirs(out_logs, exist_ok=True)
    os.makedirs(out_csv, exist_ok=True)

    results_csv = os.path.join(out_csv, f"vina_results_{SCORING}_{LIGAND_TAG}.csv")
    master_log  = os.path.join(out_logs, f"vina_master_{SCORING}_{LIGAND_TAG}.log")
    return out_pdbqt, out_logs, out_csv, results_csv, master_log


def append_line(file_path: str, line: str) -> None:
    with locked_open(file_path, "a") as f:
        f.write(line)
        f.flush()


def ensure_csv_header(csv_path: str, header: List[str]) -> None:
    # Lock to avoid two array tasks racing to create/write header
    with locked_open(csv_path, "a+") as f:
        f.seek(0, os.SEEK_END)
        if f.tell() == 0:
            w = csv.writer(f)
            w.writerow(header)
            f.flush()


def read_done_from_csv(csv_path: str) -> Dict[str, str]:
    done: Dict[str, str] = {}
    if not os.path.exists(csv_path) or os.path.getsize(csv_path) == 0:
        return done
    # Lock read to avoid parsing while another task is appending
    with locked_open(csv_path, "r") as f:
        r = csv.DictReader(f)
        for row in r:
            pid = (row.get("patch_id") or "").strip()
            status = (row.get("status") or "").strip().upper()
            if pid and status in {"OK", "RECOVERED"}:
                done[pid] = status
    return done


def list_maps_fld(maps_dir: str) -> List[str]:
    files = [os.path.join(maps_dir, n) for n in os.listdir(maps_dir) if n.endswith(".maps.fld")]
    files.sort()
    return files


def parse_maps_fld_for_files(fld_text: str) -> List[str]:
    out = []
    for line in fld_text.splitlines():
        line = line.strip()
        if not line or line.startswith("#"):
            continue
        m = re.match(r"^(map|elecmap|dsolvmap)\s+(.+)$", line)
        if m:
            out.append(m.group(2).strip())
    return out


def validate_maps_fld(maps_dir: str, maps_fld_path: str) -> Tuple[bool, List[str]]:
    with open(maps_fld_path, "r", encoding="utf-8", errors="replace") as f:
        txt = f.read()
    refs = parse_maps_fld_for_files(txt)
    missing = []
    for ref in refs:
        p = ref if ref.startswith("/") else os.path.join(maps_dir, ref)
        if not os.path.isfile(p):
            missing.append(p)
    return (len(missing) == 0), missing


def detect_vina() -> Tuple[str, str]:
    exe = shutil.which("vina")
    if not exe:
        raise RuntimeError("AutoDock Vina not found in PATH. Ensure `vina --help` and `vina --version` work in this shell/env.")
    cp = subprocess.run([exe, "--version"], capture_output=True, text=True)
    ver = (cp.stdout or cp.stderr or "").strip()
    return exe, ver


def parse_score_from_stdout(lines: List[str]) -> Optional[float]:
    # Standard Vina table line: "1   -7.5  0.0  0.0"
    for line in lines:
        s = line.strip()
        m = re.match(r"^1\s+(-?\d+(?:\.\d+)?)\s+\d+(?:\.\d+)?\s+\d+(?:\.\d+)?", s)
        if m:
            try:
                return float(m.group(1))
            except Exception:
                pass
        m2 = re.search(r"(Affinity|Score|Energy)\s*[:=]\s*(-?\d+(?:\.\d+)?)", s, re.IGNORECASE)
        if m2:
            try:
                return float(m2.group(2))
            except Exception:
                pass
    return None


def parse_score_from_pdbqt(pdbqt_path: str) -> Optional[float]:
    try:
        with open(pdbqt_path, "r", encoding="utf-8", errors="replace") as f:
            txt = f.read()
    except Exception:
        return None

    m = re.search(r"REMARK\s+VINA\s+RESULT:\s*([-\d\.]+)", txt)
    if m:
        try:
            return float(m.group(1))
        except Exception:
            return None

    m = re.search(r"REMARK.*RESULT[:\s]+([-\d\.]+)", txt)
    if m:
        try:
            return float(m.group(1))
        except Exception:
            return None

    m = re.search(r"REMARK.*(score|energy|affinity).*?([-\d]+\.\d+|[-\d]+)", txt, re.IGNORECASE)
    if m:
        try:
            return float(m.group(2))
        except Exception:
            return None

    return None


def run_vina(
    vina_exe: str,
    ligand_path: str,
    maps_arg: str,
    out_pose: str,
    patch_log: str,
    extra_note: str
) -> Tuple[int, float, List[str]]:
    cmd = [
        vina_exe,
        "--ligand", ligand_path,
        "--scoring", SCORING,
        "--maps", maps_arg,
        "--out", out_pose,
        "--cpu", str(CPU_THREADS),
        "--exhaustiveness", str(EXHAUSTIVENESS),
        "--num_modes", str(NUM_MODES),
        "--verbosity", str(VERBOSITY),
    ]
    if SEED is not None:
        cmd += ["--seed", str(SEED)]

    start = dt.datetime.now()
    append_line(patch_log, f"[{now_ts()}] NOTE: {extra_note}\n")
    append_line(patch_log, f"[{now_ts()}] CMD: {' '.join(cmd)}\n")

    captured: List[str] = []
    p = subprocess.Popen(
        cmd,
        stdout=subprocess.PIPE,
        stderr=subprocess.STDOUT,
        text=True,
        bufsize=1,
        cwd=os.path.dirname(maps_arg) if os.path.dirname(maps_arg) else None,
    )

    assert p.stdout is not None
    for line in p.stdout:
        append_line(patch_log, line)
        if len(captured) < 8000:
            captured.append(line)

    rc = p.wait()
    runtime_s = (dt.datetime.now() - start).total_seconds()
    append_line(patch_log, f"[{now_ts()}] RC={rc} runtime_s={runtime_s:.1f}\n")
    return rc, runtime_s, captured


def append_csv_row(csv_path: str, row: Dict[str, str], header: List[str]) -> None:
    # Lock to avoid interleaved writes from concurrent array tasks
    with locked_open(csv_path, "a") as f:
        w = csv.DictWriter(f, fieldnames=header)
        w.writerow(row)
        f.flush()


def main() -> int:
    rid = run_id()
    print(f"=== AutoDock Vina batch runner starting (run_id={rid}) ===")

    parser = argparse.ArgumentParser(add_help=True)
    parser.add_argument("--offset", type=int, default=int(os.environ.get("OFFSET", "0")))
    # Your style uses --array=1-N, so task_id is 1-based when provided.
    parser.add_argument("--task_id", type=int, default=int(os.environ.get("SLURM_ARRAY_TASK_ID", "0")))
    args = parser.parse_args()

    if not os.path.isfile(LIGAND_PATH):
        print(f"ERROR: ligand not found: {LIGAND_PATH}")
        return 2
    if not os.path.isdir(MAPS_DIR):
        print(f"ERROR: maps dir not found: {MAPS_DIR}")
        return 2

    vina_exe, vina_ver = detect_vina()
    print(f"Vina: {vina_exe}")
    print(f"Vina version: {vina_ver}")

    out_pdbqt, out_logs, out_csv, results_csv, master_log = ensure_dirs()

    header = [
        "run_id",
        "patch_id",
        "scoring",
        "cpu",
        "exhaustiveness",
        "num_modes",
        "seed",
        "vina_version",
        "best_score",
        "status",
        "finished_time",
    ]
    ensure_csv_header(results_csv, header)

    append_line(master_log, f"[{now_ts()}] START run_id={rid} vina={vina_ver} ligand={LIGAND_PATH} maps_dir={MAPS_DIR}\n")

    done = read_done_from_csv(results_csv)
    maps_flds = list_maps_fld(MAPS_DIR)
    if not maps_flds:
        msg = f"[{now_ts()}] ERROR: no *.maps.fld found under {MAPS_DIR}\n"
        print(msg.strip())
        append_line(master_log, msg)
        return 2

    # ===== SLURM ARRAY MODE (added) =====
    # Each array task runs exactly ONE patch:
    # idx0 = offset + (task_id - 1)  (because task_id is 1-based in your submissions)
    if args.task_id > 0:
        idx0 = args.offset + (args.task_id - 1)
        if idx0 < 0 or idx0 >= len(maps_flds):
            msg = f"[{now_ts()}] ERROR: task_id={args.task_id} offset={args.offset} idx0={idx0} out of range (n={len(maps_flds)})\n"
            print(msg.strip())
            append_line(master_log, msg)
            return 2
        maps_flds = [maps_flds[idx0]]
    # ===== END SLURM ARRAY MODE =====

    total = len(maps_flds)
    ok = fail = skip_done = skip_missing_maps = recovered = 0

    for i, maps_fld in enumerate(maps_flds, start=1):
        patch_id = os.path.basename(maps_fld).replace(".maps.fld", "")
        maps_prefix = os.path.join(MAPS_DIR, patch_id)

        out_pose = os.path.join(out_pdbqt, f"{patch_id}__{LIGAND_TAG}__{SCORING}.pdbqt")
        patch_log = os.path.join(out_logs, f"{patch_id}__{LIGAND_TAG}__{SCORING}.log")

        if patch_id in done:
            skip_done += 1
            print(f"[{i}/{total}] SKIP_DONE: {patch_id} ({done[patch_id]})")
            append_line(master_log, f"[{now_ts()}] SKIP_DONE patch_id={patch_id} status={done[patch_id]}\n")
            continue

        # RECOVER
        if os.path.isfile(out_pose) and os.path.getsize(out_pose) > 0:
            score = parse_score_from_pdbqt(out_pose)
            if score is not None:
                recovered += 1
                print(f"[{i}/{total}] RECOVER: {patch_id} score={score}")
                append_line(master_log, f"[{now_ts()}] RECOVER patch_id={patch_id} score={score}\n")
                row = {
                    "run_id": rid,
                    "patch_id": patch_id,
                    "scoring": SCORING,
                    "cpu": str(CPU_THREADS),
                    "exhaustiveness": str(EXHAUSTIVENESS),
                    "num_modes": str(NUM_MODES),
                    "seed": "" if SEED is None else str(SEED),
                    "vina_version": vina_ver,
                    "best_score": str(score),
                    "status": "RECOVERED",
                    "finished_time": now_ts(),
                }
                append_csv_row(results_csv, row, header)
                done[patch_id] = "RECOVERED"
                continue

        if VALIDATE_MAPS_FLD_REFERENCES:
            ok_maps, missing = validate_maps_fld(MAPS_DIR, maps_fld)
            if not ok_maps:
                skip_missing_maps += 1
                print(f"[{i}/{total}] SKIP_MISSING_MAPS: {patch_id} missing={len(missing)}")
                append_line(master_log, f"[{now_ts()}] SKIP_MISSING_MAPS patch_id={patch_id} missing_count={len(missing)}\n")
                for m in missing[:30]:
                    append_line(master_log, f"    MISSING {m}\n")
                if len(missing) > 30:
                    append_line(master_log, f"    ... ({len(missing)-30} more)\n")
                continue

        print(f"[{i}/{total}] RUN: {patch_id}")
        append_line(master_log, f"[{now_ts()}] RUN patch_id={patch_id}\n")
        append_line(patch_log, f"\n[{now_ts()}] ===== PATCH START patch_id={patch_id} run_id={rid} =====\n")

        # Attempt A: --maps <prefix>
        rcA, rtA, capA = run_vina(
            vina_exe, LIGAND_PATH, maps_prefix, out_pose, patch_log,
            extra_note="Attempt A: --maps <prefix>"
        )

        rc = rcA
        runtime_s = rtA
        captured = capA

        # Attempt B: --maps <.maps.fld> (fallback)
        if rcA != 0:
            append_line(patch_log, f"[{now_ts()}] RETRY: Attempt B using --maps <maps_fld>\n")
            rcB, rtB, capB = run_vina(
                vina_exe, LIGAND_PATH, maps_fld, out_pose, patch_log,
                extra_note="Attempt B: --maps <.maps.fld>"
            )
            rc = rcB
            runtime_s += rtB
            captured = capA + capB

        if rc != 0:
            fail += 1
            print(f"[{i}/{total}] FAIL: {patch_id} (rc={rc})")
            append_line(master_log, f"[{now_ts()}] FAIL patch_id={patch_id} rc={rc} runtime_s={runtime_s:.1f}\n")
            continue

        if not os.path.isfile(out_pose) or os.path.getsize(out_pose) == 0:
            fail += 1
            print(f"[{i}/{total}] FAIL_NO_OUTPUT: {patch_id}")
            append_line(master_log, f"[{now_ts()}] FAIL_NO_OUTPUT patch_id={patch_id} runtime_s={runtime_s:.1f}\n")
            continue

        score = parse_score_from_stdout(captured) or parse_score_from_pdbqt(out_pose)
        if score is None:
            fail += 1
            print(f"[{i}/{total}] FAIL_NO_SCORE: {patch_id}")
            append_line(master_log, f"[{now_ts()}] FAIL_NO_SCORE patch_id={patch_id} runtime_s={runtime_s:.1f}\n")
            continue

        ok += 1
        row = {
            "run_id": rid,
            "patch_id": patch_id,
            "scoring": SCORING,
            "cpu": str(CPU_THREADS),
            "exhaustiveness": str(EXHAUSTIVENESS),
            "num_modes": str(NUM_MODES),
            "seed": "" if SEED is None else str(SEED),
            "vina_version": vina_ver,
            "best_score": str(score),
            "status": "OK",
            "finished_time": now_ts(),
        }
        append_csv_row(results_csv, row, header)
        done[patch_id] = "OK"

        print(f"[{i}/{total}] OK: {patch_id} score={score} (runtime {runtime_s:.1f}s)")
        append_line(master_log, f"[{now_ts()}] OK patch_id={patch_id} score={score} runtime_s={runtime_s:.1f}\n")

    summary = f"[{now_ts()}] SUMMARY total={total} ok={ok} fail={fail} skip_done={skip_done} skip_missing_maps={skip_missing_maps} recovered={recovered}\n"
    print(summary.strip())
    append_line(master_log, summary)

    print("\nOutputs:")
    print(f"  PDBQT : {out_pdbqt}")
    print(f"  LOGS  : {out_logs}")
    print(f"  CSV   : {results_csv}")
    print("=== Done ===")
    return 0


if __name__ == "__main__":
    sys.exit(main())
```
