## Supplementary Code for "A structural screen for candidate polyphosphate binding surfaces in human membrane associated proteins": vina.slurm

```
#!/bin/bash
#SBATCH --job-name=vina_ad4_p10
#SBATCH --output=/global/project/hpcg1856/david_poly_project/8_vina/log/slurm_%A_%a.out
#SBATCH --error=/global/project/hpcg1856/david_poly_project/8_vina/log/slurm_%A_%a.err
#SBATCH --time=05:30:00
#SBATCH --cpus-per-task=10
#SBATCH --mem=15G

set -euo pipefail

mkdir -p /global/project/hpcg1856/david_poly_project/8_vina/{csv,log,pdbqt}

module --force purge
module load StdEnv/2023 gcc/12.3
module load autodock-vina/1.2.6
module load python/3.13

# OFFSET passed at submission time, e.g. OFFSET=2999 sbatch --array=1-2999%600 vina.slurm
export OFFSET="${OFFSET:-0}"

python3 /global/project/hpcg1856/david_poly_project/8_vina/vina.py \
  --offset "${OFFSET}" \
  --task_id "${SLURM_ARRAY_TASK_ID:-0}"
```
