## Supplementary material for "A structural screen for candidate polyphosphate binding surfaces in human membrane associated proteins": File guide.txt

SUPPLEMENTARY DATA

The Supplementary Data labels match the Supplementary Information. Except for S7, each heading below names a folder within Supplementary data/. S7 maps existing files in Supplementary_Code.zip. Open CSV files as tables and XLSX files as workbooks. Model and pose coordinates are supplied separately in Supplementary_Structures.zip; analysis scripts and inputs are in Supplementary_Code.zip.

Shared identifiers and scores

patch_id and pose_id identify one AlphaFold model and positive patch. model_id, AlphaFold_model and model identify the exact protein model; alternative isoforms remain separate. UniProt_ID, base_accession and UniProt_entry identify protein entries as specified in each table.

PDI100 is the Patch Docking Index on its 0–100 scale. PRI is the pLDDT Reliability Index. Dock01 and Elec01 are the normalised docking and electrostatic components. dock_best_score_ad4 is the saved AD4 score in kcal/mol. Distances ending in _A are in angstroms; pLDDT is the AlphaFold residue confidence value on its 0–100 scale. Blank values indicate unavailable or inapplicable fields, not zero.

S1 All screening results

grand_patch_table_PDI100_polyP10.csv
The computational master table: 23,383 positive patch records, including 19,481 completed results with numerical AD4 and PDI100 scores. Each row is one patch on one model. patch_area is in nm²; patch_value_per_area is electrostatic intensity E.

grand_patch_table_PDI100_polyP10.xlsx
The master workbook presents the same 23,383 identified patch records, with separate worksheets for the 185 high scoring results and the 74 results in the normal/conditional cell surface exposure group. It retains one additional numerical PDI100 value on an incomplete record without an AD4 score; that record was excluded from completed docking analyses. Use the CSV for computational records and S2 for exact selection memberships.

S2 Selected results and ranks

Data_S2_V11.7.csv
The 185 original high scoring results. selections_of_135 counts how often that exact model and patch remained in the top 185 under the 135 PDI settings. selected_in_all_135 is true only for an exact pose selected in every setting. minimum_rank, median_rank and maximum_rank summarise ranks across the tested settings. in_normal_conditional74 marks membership in the 74 result exposure group.

model_annotation means cell surface exposure status: Yes, Conditional, No or Review; Existence uncertain identifies uncertain protein existence. annotation_qualification explains the status. These labels describe the protein model, not access to its individual docking pose.

S3 Tested PDI settings

Data_S3_V11.7.csv
The 135 parameter combinations and selection summaries. original185_selected counts original results retained in each top 185 selection. normal_conditional74_selected counts retained results from the 74 result normal/conditional cell surface exposure group; other_model_annotations111_selected counts retained results from the other 111 high scoring results. above80_count is a separate diagnostic count; a score cutoff of 80 was not reapplied as the selection rule for these settings.

S4 Functional enrichment

functional_keyword_enrichment_all219.csv
All 219 tested UniProt annotations, including nonsignificant results. high_with and background_with count entries carrying each annotation; the corresponding total fields give their populations. background_total is the full completed-result reference population. p and q are the unadjusted and Benjamini–Hochberg adjusted values. canonical_ fields use canonical models; opportunity_ fields control completed search count; opportunity_length_ fields control both search count and sequence length.

S5 Isoform score pairs

isoform_pairs_reconstructed.csv
The 1,175 paired source score summaries. _canonical fields are means across completed results for the canonical model. _alternative fields are medians of the corresponding alternative model means. D is the negative AD4 score, E is electrostatic intensity, and Base is the score before the PRI penalty.

S6 Contacts for all selected poses

contacts_185_4A.csv
The 2,086 contacting residue records across all 185 poses. minimum_distance_A is the shortest nonhydrogen atom distance between that residue and polyP10. The contact cutoff is ≤4.000 Å before rounding. model_id, chain, residue_number and insertion_code identify the residue.

S7 Screening inputs and provenance records

S7 is a logical collection in Supplementary_Code.zip. These are existing files, with their original names and locations; there is no separate S7 folder. Paths below start from Supplementary code/.

Screening workflow/Inputs/surface_candidates_uid.list
The 4,062 starting UniProt accession IDs.

Screening workflow/Inputs/pdb_list.txt
The 8,808 retrieved AlphaFold model filenames.

Screening workflow/Inputs/alphafold_summary.csv.xlsx
AlphaFold model metadata, retrieval summary and source download URLs.

Screening workflow/Inputs/Original patch records.tar.gz
The original 8,678 patch lists and 8,678 residue lists, under patch_records/ after extraction.

Functional analyses/sources/vina_patch_boxes_20260219_070756.csv
Patch-specific docking box records.

Functional analyses/sources/vina_patch_boxes_20260219_070756_PRI_only.csv
Saved PRI records and their model/patch identifiers.

Functional analyses/sources/vina_results_ad4_polyP10.csv
Saved docking score records linked to the PDBQT poses in Supplementary_Structures.zip.

Functional analyses/sources/surface_candidates_annotation_snapshot.csv
Frozen annotation snapshot used for the functional comparisons.

PDI selection/inputs/Entry_Audit_128.csv
Cell surface exposure review for the 128 selected models.

PDI selection/inputs/curated74.csv
Original input table for the 74 results in the normal/conditional cell surface exposure group. Data S2 provides the final per-result labels and rank summaries.

Screening workflow/source_polyP10/polyP10.sdf
Screening workflow/source_polyP10/polyP10.pdbqt
Original ligand inputs.

S8 Functional groups

Data_S8_V11.7.csv
The 17 oxidoreductase and translocase entries. all185_ fields describe all high scoring results; normal_conditional74_ fields describe the exposure group. Scores and selection counts refer to the specified representative patch. stable_results counts results selected in all 135 settings.

NOX_DUOX_family.csv
The seven NOX/DUOX entries, including their highest overall and canonical PDI100 scores. n_curated_patch_runs counts results in the 74 result normal/conditional cell surface exposure group; curated_status gives their cell surface exposure labels.

S9 LPL and LIPI residue contacts

LPL_LIPI_residue_contacts.csv
One row per residue, with its shortest nonhydrogen atom distance to polyP10 and contact flags at 3.5, 4 and 5 Å.

These distances were reconstructed from the saved canonical LPL patch001 and canonical LIPI patch002 coordinates, without redocking or changing the coordinates.

S10 Experimental sites for 74 poses

PolyP_Experimental_Site_Overlap_74_Poses.xlsx
The 74 exposure group poses. All74screen gives the complete pose list. Site overlap summarises comparisons; Site residues retains mapped residues, distances and noncontacts. Evidence identifies the studies and experimental sets. Numbering explains sequence mapping; Methods defines the comparisons.

TRPML3_ortholog_mapping.json
TRPML1_mouse_ortholog_mapping.json
NCC_construct_mapping.json
The original ortholog and construct residue-number mapping records referenced by the workbook. These files are supplied beside the workbook with their original filenames.

S11 LPL phosphate contacts

LPL_LIPI_basic_nitrogen_phosphate_distances.csv
Distances from Lys NZ or Arg NE/NH1/NH2 atoms to phosphate oxygens, with separate results excluding shared bridging oxygens and contact flags at 3.5, 4 and 5 Å. LPL records support main Figure 6a; LIPI records support its supplementary comparison.

LPL_LIPI_phosphate_atoms.csv
Phosphate numbering, phosphorus and oxygen atom serials, and shared bridging oxygen assignments. Atom serials refer to the saved ligand files; phosphate numbers follow the reconstructed chain.

These measurements were reconstructed from the same unchanged coordinates as S9, without redocking. S9 uses all residue nonhydrogen atoms; S11 isolates basic side chain nitrogens.

S12 Experimental sites for 111 poses

PolyP_Experimental_Site_Overlap_111_Poses.xlsx
The other 111 poses, with the same comparison sheets as S10 and All111screen as the complete list. Pose contacts supplies residue contacts; Download manifest identifies the saved coordinate files. Its File path starts from the folder containing Supplementary data, Supplementary code and Candidate structures. Source folder and Drive fields identify the original download source; File path locates the supplied copy. S10 and S12 together cover all 185 poses.

priority_mapping_corrections.json
Original held mapping record cited by S12. The PLD2 candidate mappings remain unaccepted; this record does not establish a new mapping.

S13 Model evidence groups

PolyP_Data_S13_Model_Evidence_Groups_V13.5.8.csv
One row per AlphaFold model, covering all 128 models. Consistently_selected_model and Functional_site_overlap describe the two properties. Both_properties_in_same_pose distinguishes a shared supporting pose from Both_properties_only_in_different_poses. The pose-ID fields identify the exact supporting records.

Figure S1 Patch properties

composition_docking_diagnostics.csv
Association estimates and 95% bootstrap intervals for docking favourability and patch properties. HRK denotes histidine, arginine and lysine; n_patches and n_base_clusters give patch and UniProt-entry counts.

within_vs_between_model_variation.csv
Fractions of each score's total squared variation within and between AlphaFold models. Multiply these fractions by 100 for the percentages in Figure S1b.
